# Identification of candidate biomarkers and signaling pathways associated with Alzheimer’s disease using bioinformatics analysis of next generation sequencing data

**DOI:** 10.1101/2024.12.03.626535

**Authors:** Basavaraj Vastrad, Chanabasayya Vastrad

## Abstract

Alzheimer’s disease (AD) is the most common cause of dementia, and one of the most common health problems all over the world. However, the specific molecular mechanisms of AD have not been fully investigated. The current investigation aimed to elucidate potential key candidate genes and signaling pathways in AD. Next generation sequencing (NGS) dataset GSE203206 was downloaded from the Gene Expression Omnibus (GEO) database, which included data from 39 AD samples and 8 normal control samples. Differentially expressed genes (DEGs) were identified using t-tests in the limma R bioconductor package. These DEGs were subsequently investigated by Gene ontology (GO) and pathway enrichment analysis, and a protein-protein interaction (PPI) network and modules were constructed and analyzed. The miRNA-hub gene regulatory network and TF-hub gene regulatory network analysis were performed to identify key miRNAs and TFs. The receiver operating characteristic (ROC) curve analysis was performed to estimate the clinical diagnostic value of the hub genes. A total of 958 DEGs, including 479 up regulated genes and 479 down regulated genes, were screened between AD and normal control samples. GO and pathway enrichment analysis results revealed that the up regulated genes were mainly enriched in response to stimulus, cytoplasm, small molecule binding and signal transduction, whereas down regulated genes were mainly enriched in multicellular organism development, cell junction, ion binding and cardiac conduction. The PPI network contained 4886 nodes and 10342 edges. HSP90AA1, FN1, KIT, YAP1, LSM2, SKP1, EIF5A2, TAF9, DDX39B and CDK7 were identified as the top hub genes. The regulatory network analysis revealed that microRNA (miRNA) hsa-mir-545-3p and hsa-miR-548f-5p, and transcription factor (TF) PLAG1 and MEF2A might be involved in the development of AD. These findings provide new insights into the pathogenesis of AD. The hub genes, miRNAs and TFs have the potential to be used as diagnostic and therapeutic markers.

## Introduction

Alzheimer’s disease (AD) is a severe neurodegenerative disorder characterized by extracellular senile plaques composed of amyloid β (Aβ) [1], intracellular tau aggregates [2], and neuronal loss and neurofibrillary degeneration [3]. The clinical incidence of AD is high, and its main features include slowly destroys memory and thinking skills [4]. AD occurring worldwide with an incidence of about 24 million persons, presenting at before the age of 60, and leading to both physical and psychological substantial burden for individuals and society [5]. At present, it is generally believed that the onset of AD might be caused by various risk factors such as age [6], genetic factors [7], head injuries [8], vascular diseases [9], infections [10], inflammation [11], environmental factors [12], diabetes mellitus [13], obesity [14], hypertension [15] and cardiovascular diseases [16]. The other neurological complications of AD include multiple sclerosis [17], Huntington’s disease [18], schizophrenia [19], autism spectrum disorder [20], amyotrophic lateral sclerosis [21], stroke [22], epilepsy [23], dementia [24], Parkinson’s diseases [25], bipolar disorder [26] and depression [27]. However, the specific regulatory mechanism of AD is still unclear, and further exploration is needed.

Debate on the leading strategy for AD management continues despite great progress in treating AD in current decades. Extensive investigation have shown recent therapeutic approaches in AD included cholinesterase inhibitors [28] and N-methyl-D-aspartate receptor noncompetitive antagonist [29] targeting several crucial signaling pathways. However, AD might also can be caused by many unknown causes, which cannot be well solved by current drug treatment and AD is still a complicated incurable neurodegenerative disease [30]. Thus, it is necessary for us to utilize bioinformatics and next generation sequencing (NGS) technology to explore the molecular pathogenesis or potential treatments of AD.

Bioinformatics methods and NGS technology are widely used to find molecular changes in the occurrence and development of diseases and are effective ways to explore the pathogenesis of diseases [31–32]. The biomarkers and signaling pathways that are being used for the etiological diagnosis of AD are genetic markers. The genetic markers include APOE3 [33], OPRM1 and OPRL1 [34], NRF2 [35], INPP5D [36] and PICALM [37]. The signaling pathways include insulin signaling pathway [38], TREM2 signaling pathway [39], PI3K/Akt signaling pathway [40], Notch signaling pathway [41] and CREB signaling pathway [42]. However, there are few reports on the use of bioinformatics and NGS data analysis to explore the molecular mechanisms of AD.

Bioinformatics and NGS data analysis can be used to screen and identify genes related to diseases. The purpose of this investigation is to identify molecular biomarkers in AD using bioinformatics and NGS methods, in order to provide potential targets. NGS dataset (GSE203206) [43] downloaded from the gene expression omnibus (GEO) (https://www.ncbi.nlm.nih.gov/geo) [44] were used to identify Differential expressed genes (DEGs), gene ontology (GO) and REACTOME pathway enrichment analyses were performed, and a protein-protein interaction (PPI) network was constructed using the STRING database and Cytoscape software, and also modules were isolated and analyzed form PPI network. A miRNA-hub gene regulatory network and TF-hub gene regulatory network were also constructed to predict potential target miRNAs and TFs. The predictive capability of the hub genes analyzed by receiver operating characteristic (ROC) curve and logistic regression analyses. The final results will help us obtain novel treatment targets for AD.

## Materials and Methods

### Next generation sequencing (NGS) data source

The GEO database is a public functional genomics database, from which the (GSE203206) [43] NGS dataset (GPL20301, Illumina HiSeq 4000 (Homo sapiens)) was downloaded. GSE203206 contained 47 samples, of which we chose 39 AD samples and 8 normal control samples. Based on these data, the next step was carried out.

### Identification of DEGs

The limma package [45] of the R bioconductor is used to screen DEGs. We adjusted p-value to correct the false discovery rate caused by the multiple tests and determined it by the Benjamini & Hochberg method [46]. DEGs were selected with threshold of log2FC > 0.7764 or log2FC < -0.664 and adj.P.Val < 0.05. Differentially expressed genes with log2FC > 0.7764 were considered as up-regulated genes, and log2FC < -0.664 as down-regulated genes. And the volcano plot and Heatmap of DEGs were drawn by “ggplot2” and “gplot” R bioconductor package.

### GO and pathway enrichment analyses of DEGs

One online tool, g:Profiler (http://biit.cs.ut.ee/gprofiler/) [47], was applied to carried out the functional annotation for DEGs. GO (http://www.geneontology.org) [48] generally perform enrichment analysis of genomes. And there are mainly biological processes (BP), cellular components (CC) and molecular functions (MF) in the GO enrichment analysis. REACTOME pathway (https://reactome.org/) [49] is a comprehensive database of genomic, chemical, and systemic functional information. Therefore, g:Profiler was used to make enrichment analysis of GO and REACTOME pathway. P-value <0.05 were considered to be significantly enriched.

### Construction of the PPI network and module analysis

Protein-protein interaction (PPI) refers to the dynamic and complex molecular network of interactions that arise between different proteins. The String Database (http://string-db.org) [50] is a database for PPI analysis, providing experimental and predictive interaction information, and subject to visualization using Cytoscape (version 3.10.2) (http://www.cytoscape.org/) [51] software. The Network Analyzer plugin of Cytoscape was used to score each node gene by 4 selected algorithms, including node degree [52], betweenness [53], stress [54] and closeness [55]. The results obtained by PPI network analysis were further module analyzed using Cytoscape Software. During the analysis, PEWCC algorithm [56] was used to identify the most significant module of the PPI network.

### Construction of the miRNA-hub gene regulatory network

Bioinformatics techniques were used to construct an interaction network between hub genes and miRNAs using the miRNet (https://www.mirnet.ca/) [57] online tool. A miRNA-hub gene interaction network was drawn to facilitate the screening of target miRNAs. Based on the analysis of hub genes, target miRNAs associated with hub genes were selected, providing strong support for further investigation of the interactions between hub genes and miRNAs. The miRNA-hub gene interaction network was visualized using Cytoscape software [51].

### Construction of the TF-hub gene regulatory network

Bioinformatics techniques were used to construct an interaction network between hub genes and TFs using the NetworkAnalyst (https://www.networkanalyst.ca/) [58] online tool. A TF - hub gene interaction network was drawn to facilitate the screening of target TFs. Based on the analysis of hub genes, target TFs associated with hub genes were selected, providing strong support for further investigation of the interactions between hub genes and TFs. The TF-hub gene interaction network was visualized using Cytoscape software [51].

### Receiver operating characteristic curve (ROC) analysis

The ROC curve was subsequently established to evaluate the diagnostic accuracy of hub genes. ROC curves were constructed using the "pROC" package in R bioconductor [59] and the area under the curve (AUC) was calculated to evaluate the diagnostic effectiveness of these hub genes in the verification datasets. AUC > 0.8 was considered the ideal diagnostic value.

## Results

### Identification of DEGs

The limma package of the R bioconductor tool that is specialized in analyzing DEGs in the AD group in relative to the normal control group. Besides, the two groups were compared using limma in the GEO NGS dataset GSE203206 and the results were downloaded for further analysis. Up regulated genes of |log FC| > 0.7764, down regulated genes of |log FC| < -0.664 and adj.P.Val < 0.05 were identified as DEGs. Totally 958 DEGs, containing 479 up regulated genes and 479 down regulated genes were detected (Table 1). A volcano plot was drawn to validate the results (Fig. 1). A heatmap of the 958 DEGs is presented in Fig. 2.

**Fig. 1.**
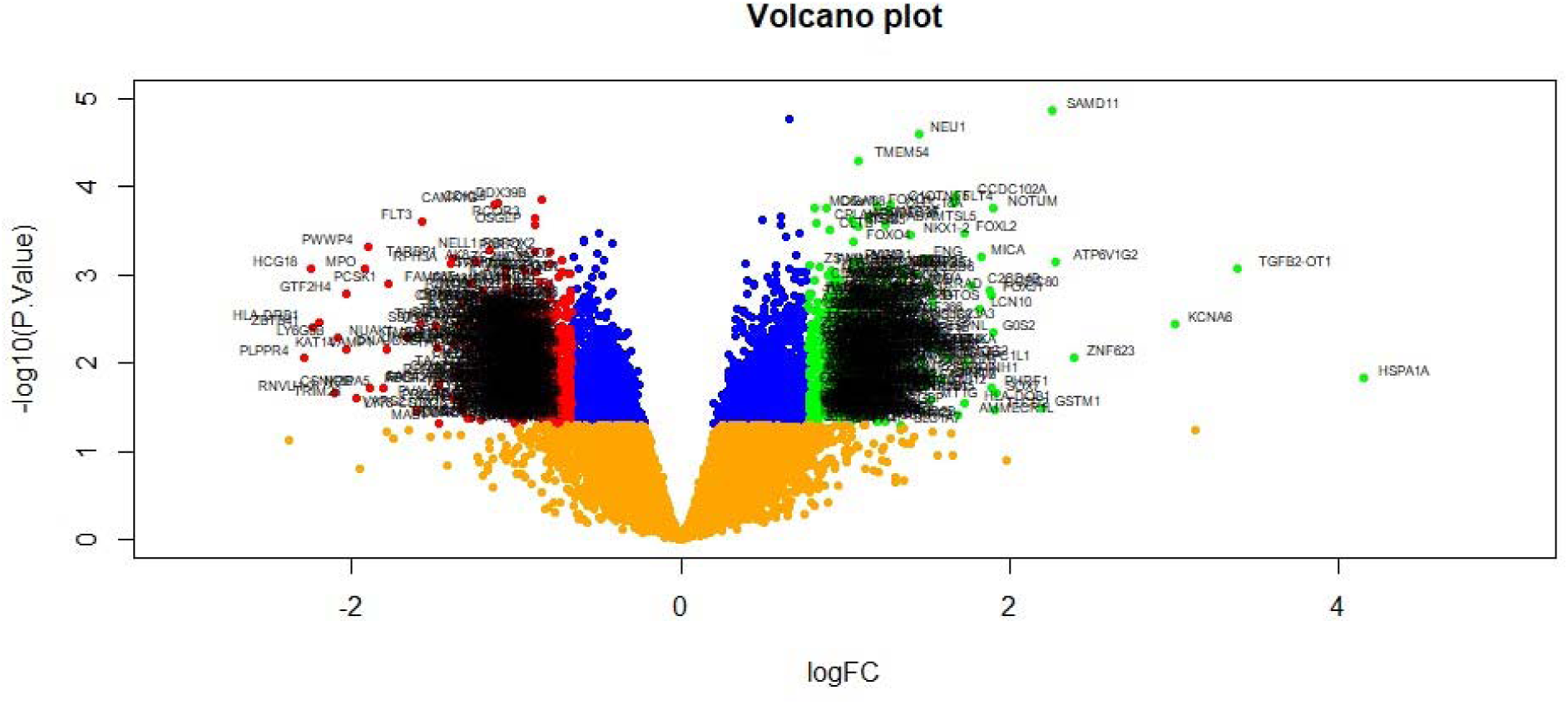
Volcano plot of differentially expressed genes. Genes with a significant change of more than two-fold were selected. Green dot represented up regulated significant genes and red dot represented down regulated significant genes.

**Fig. 2.**
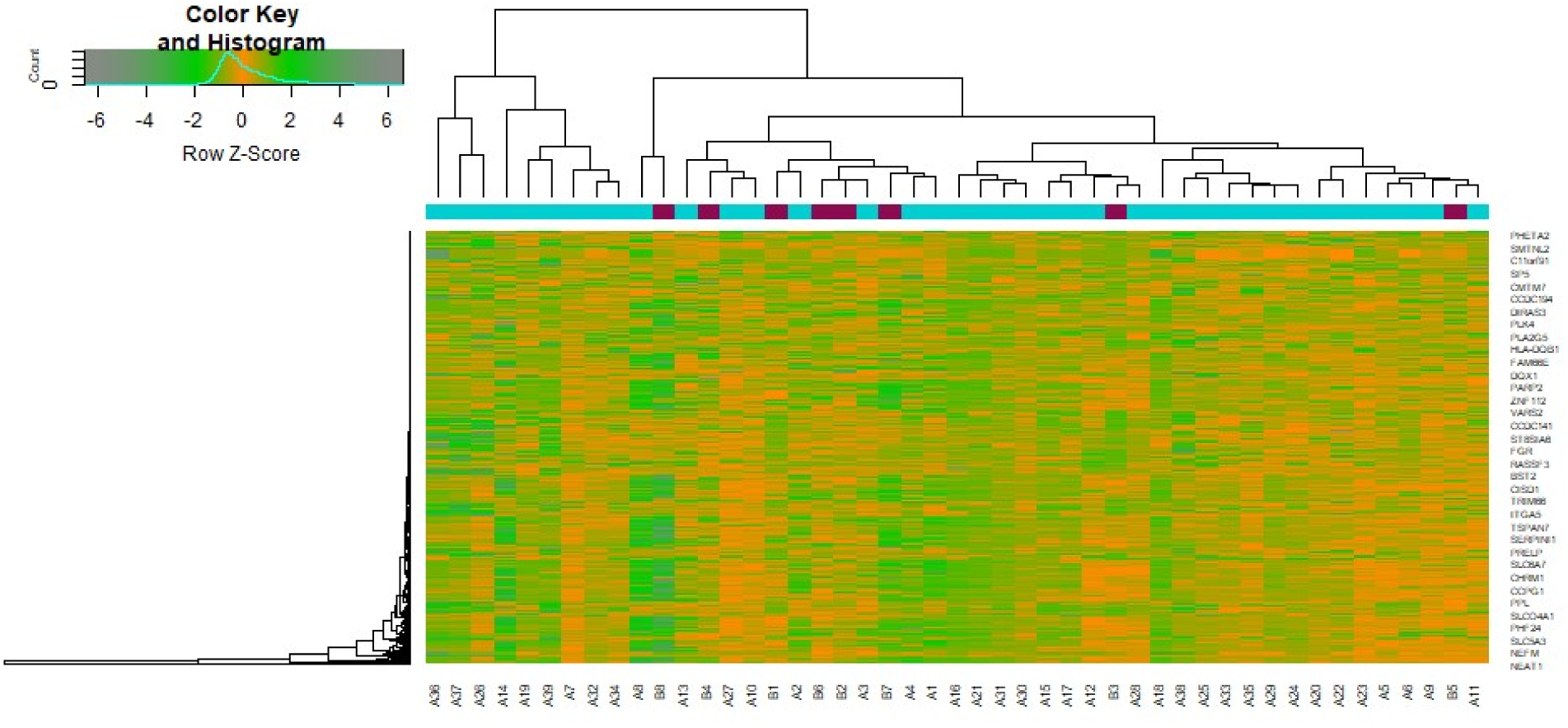
Heat map of differentially expressed genes. Legend on the top left indicate log fold change of genes. (A1 – A39 = AD samples; B1 – B8 = Normal control samples)

**Table 1.**
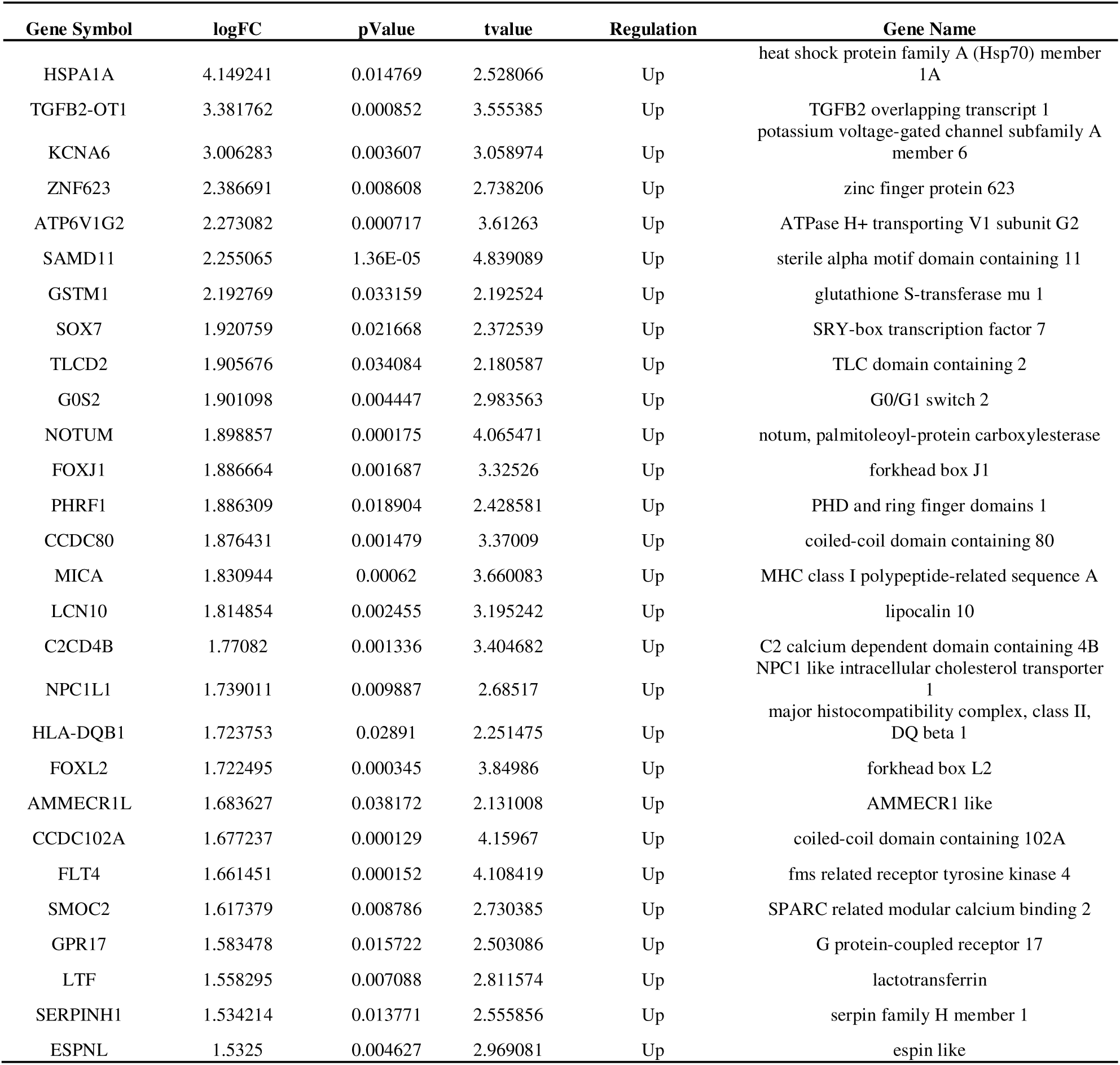
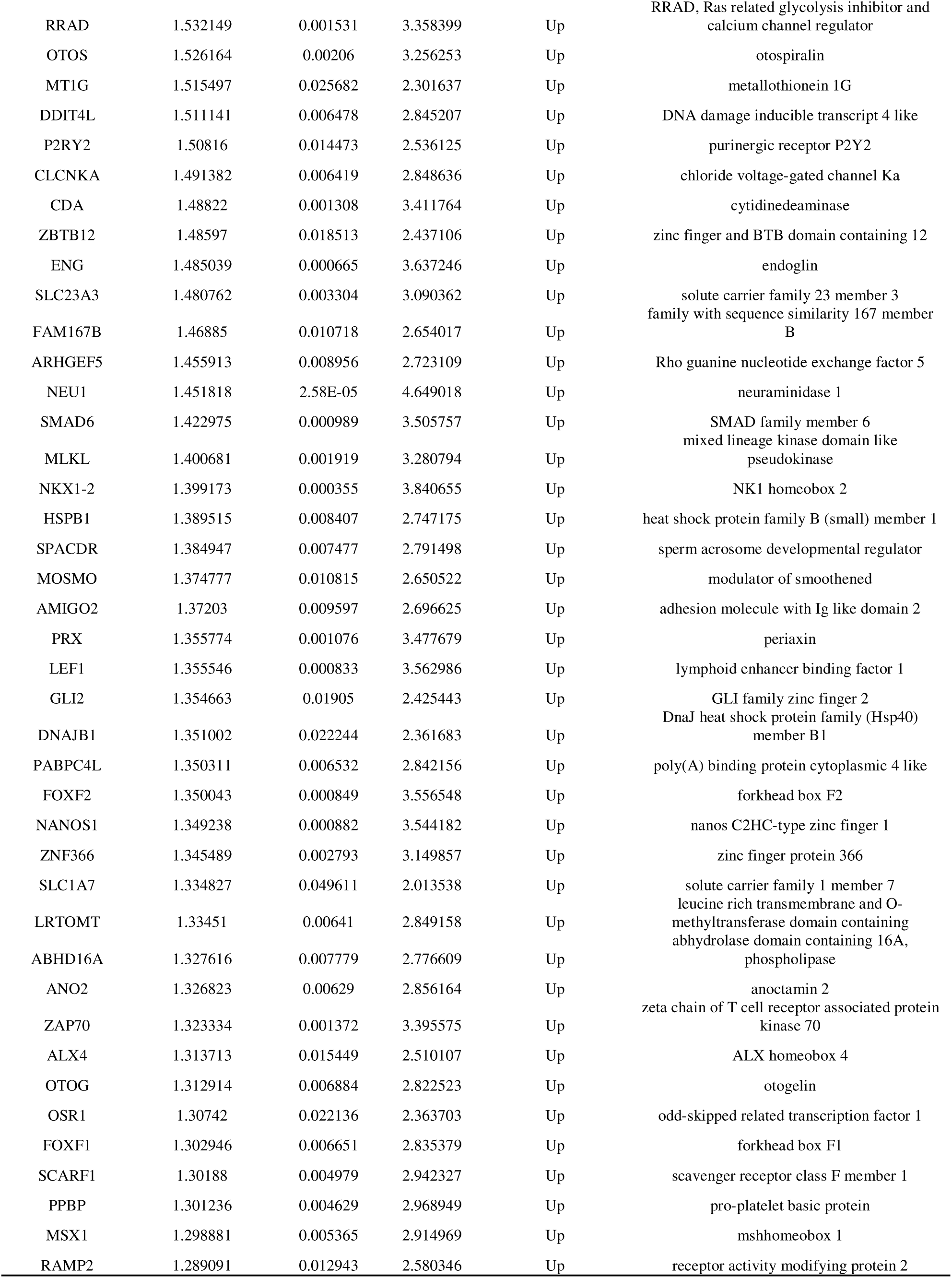
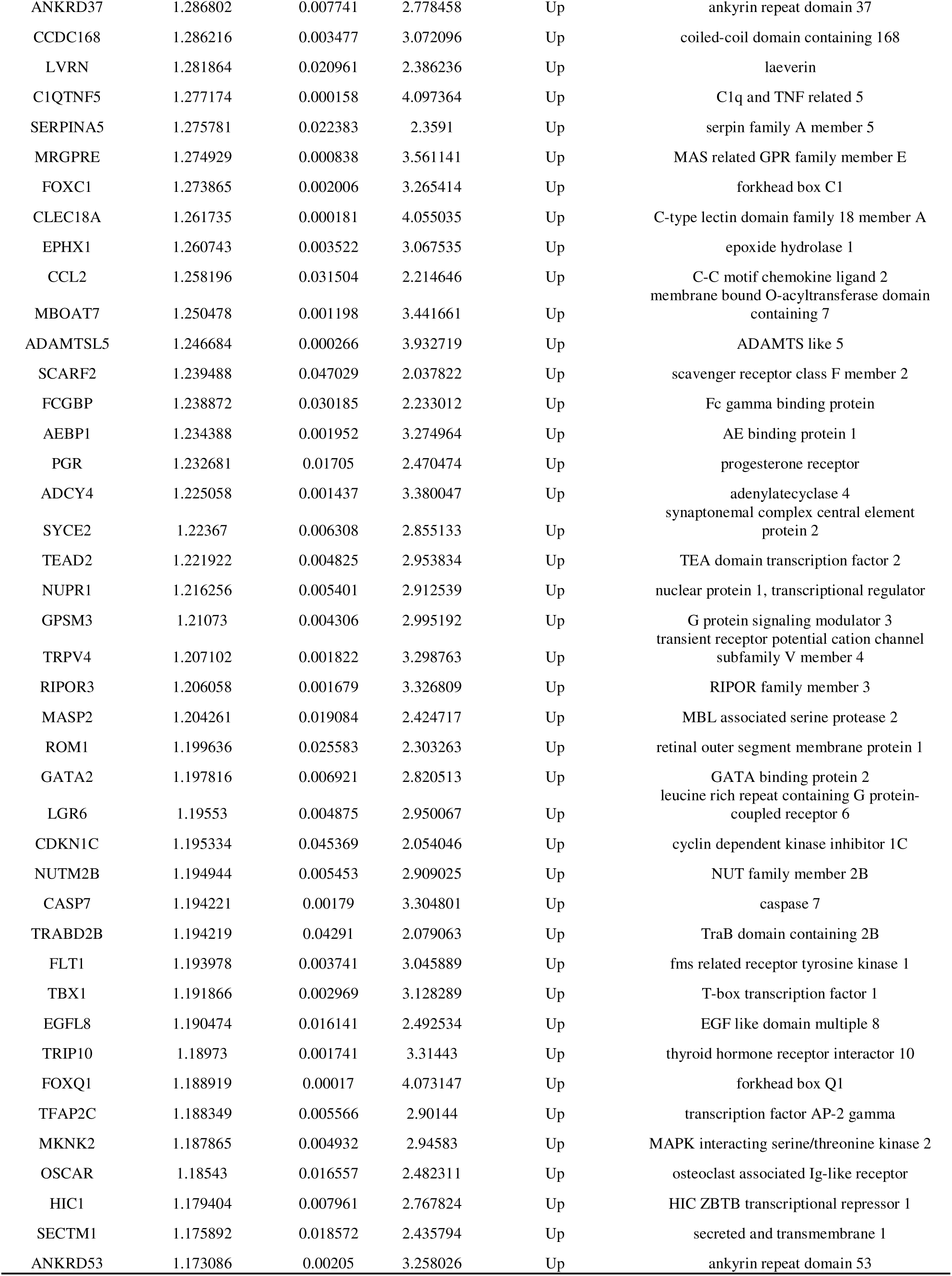
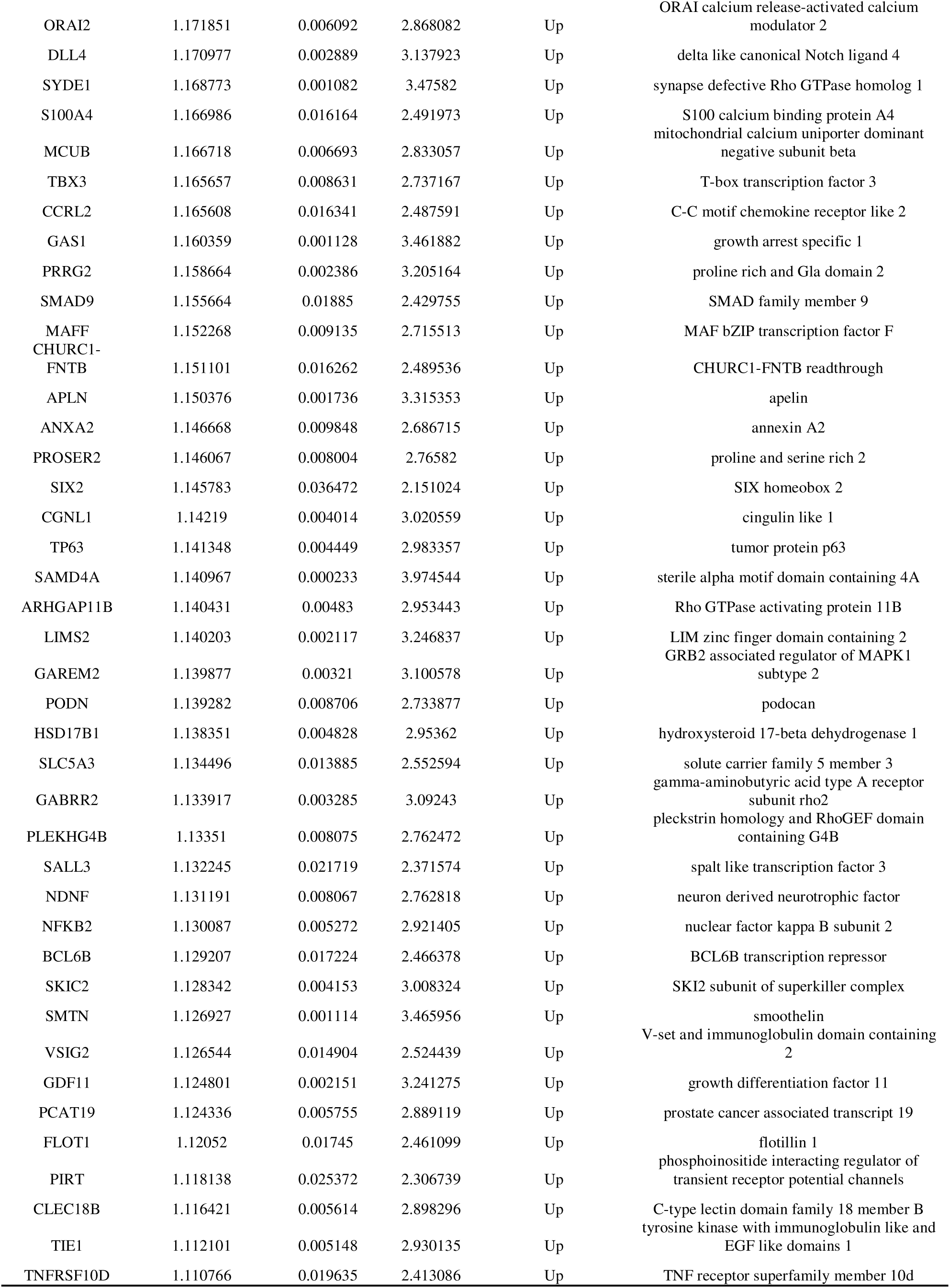
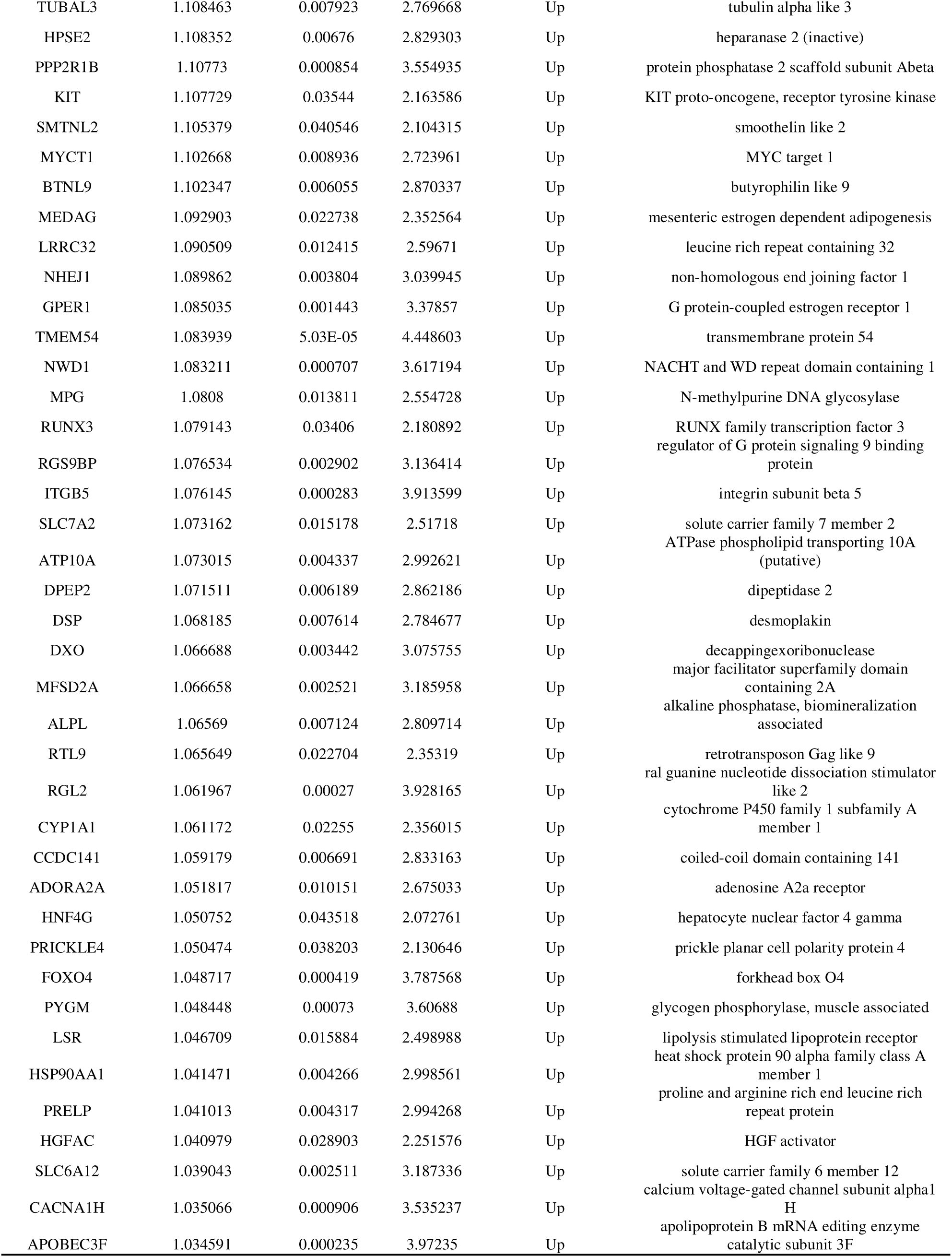
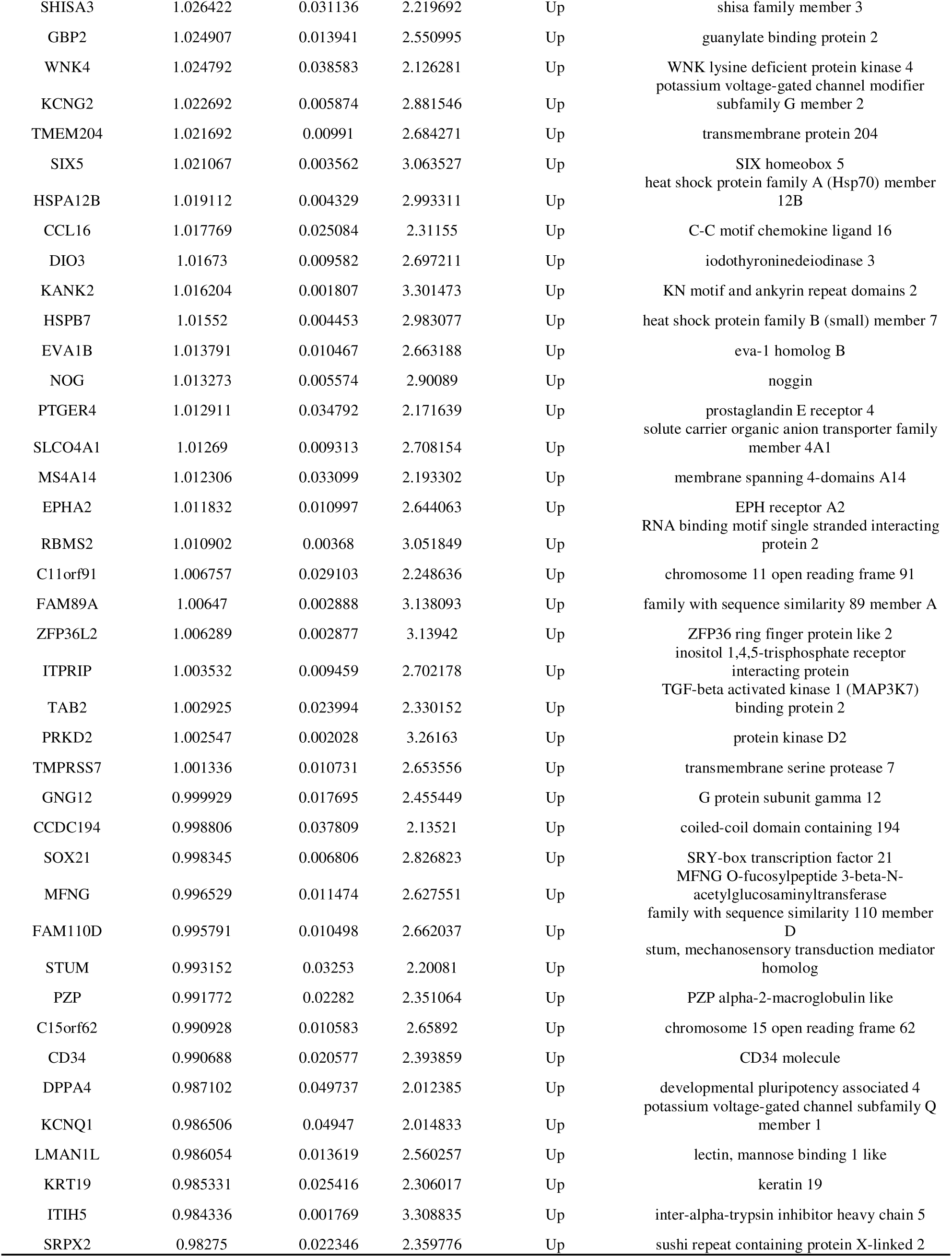
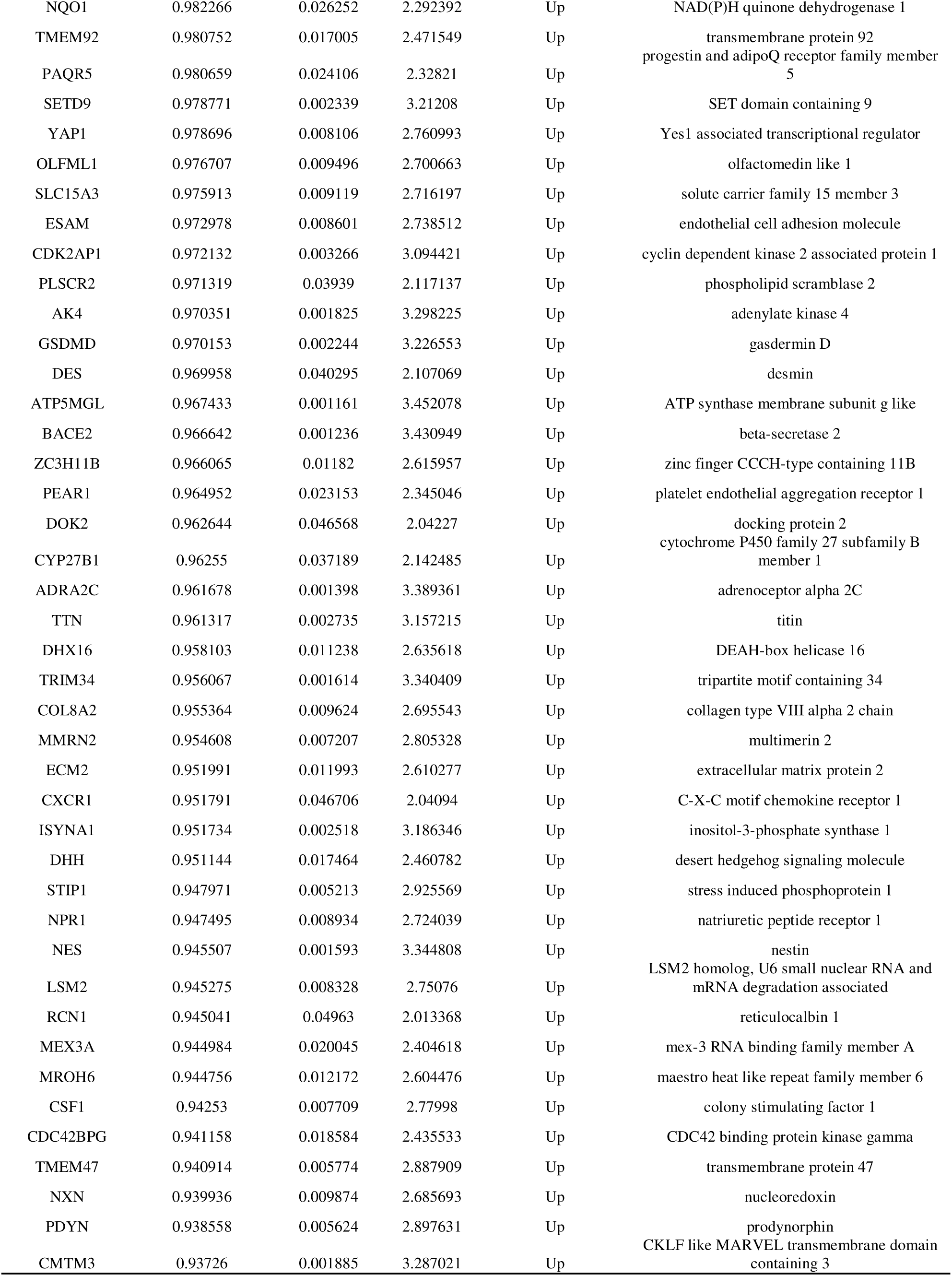
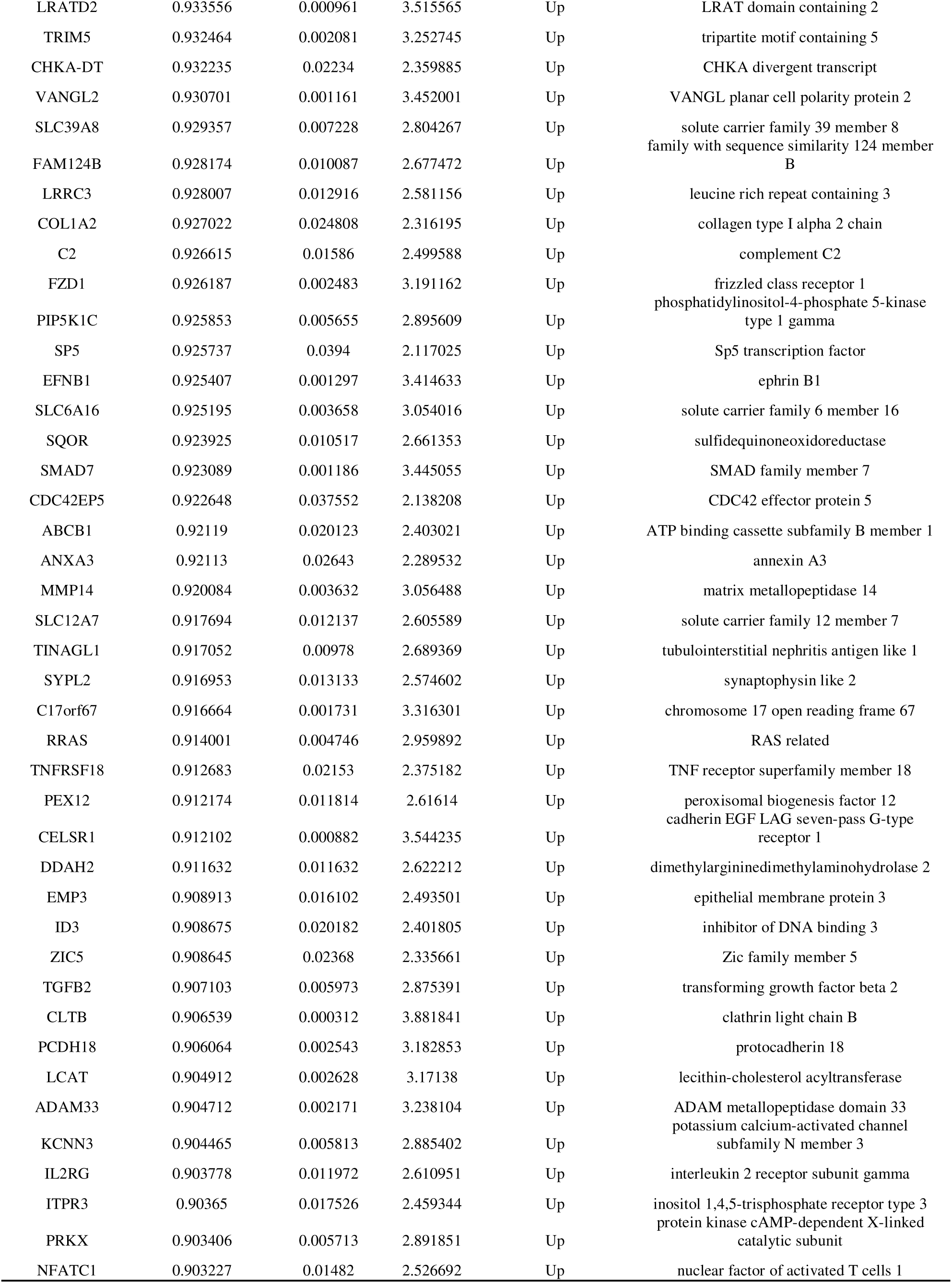
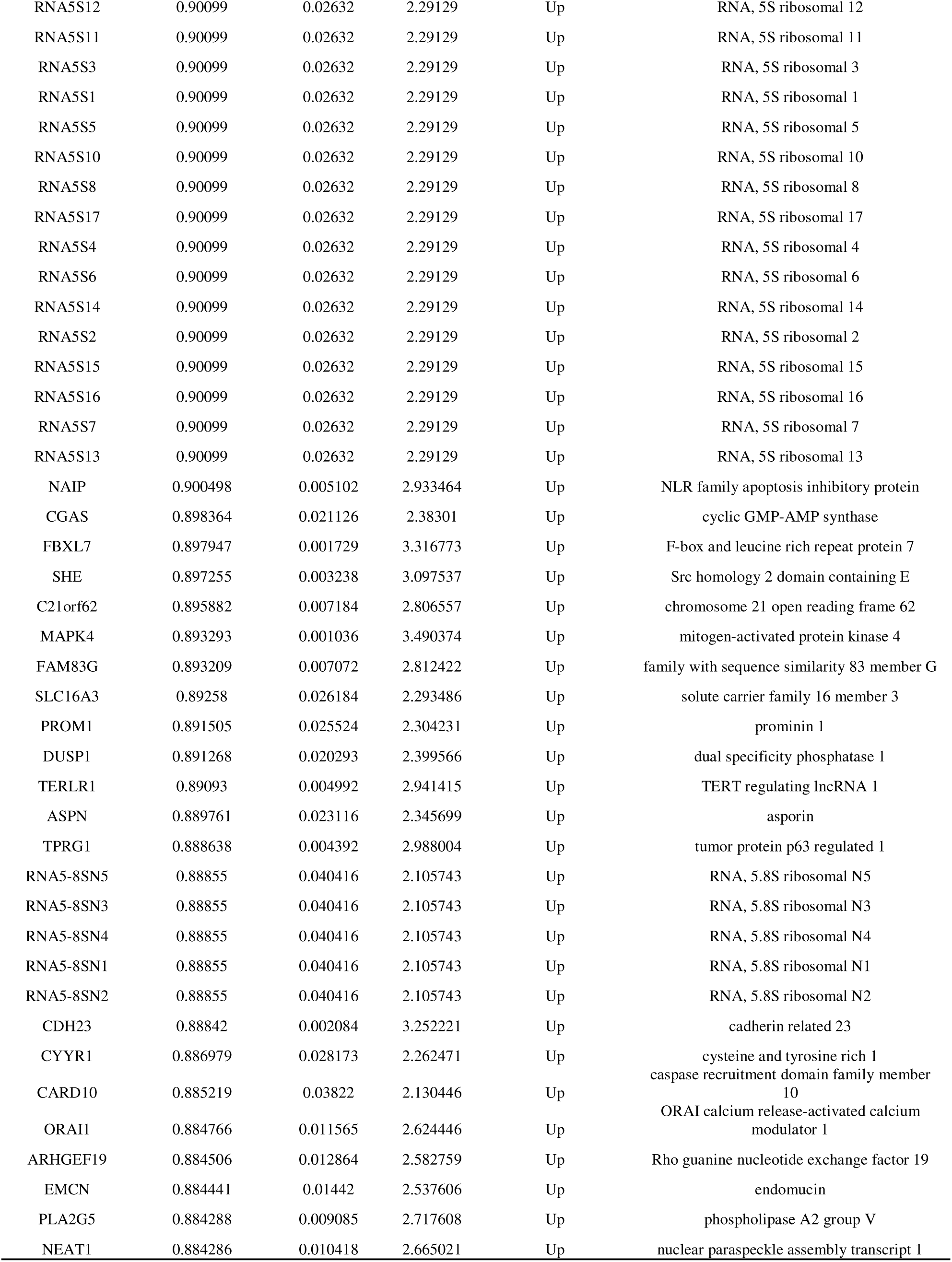
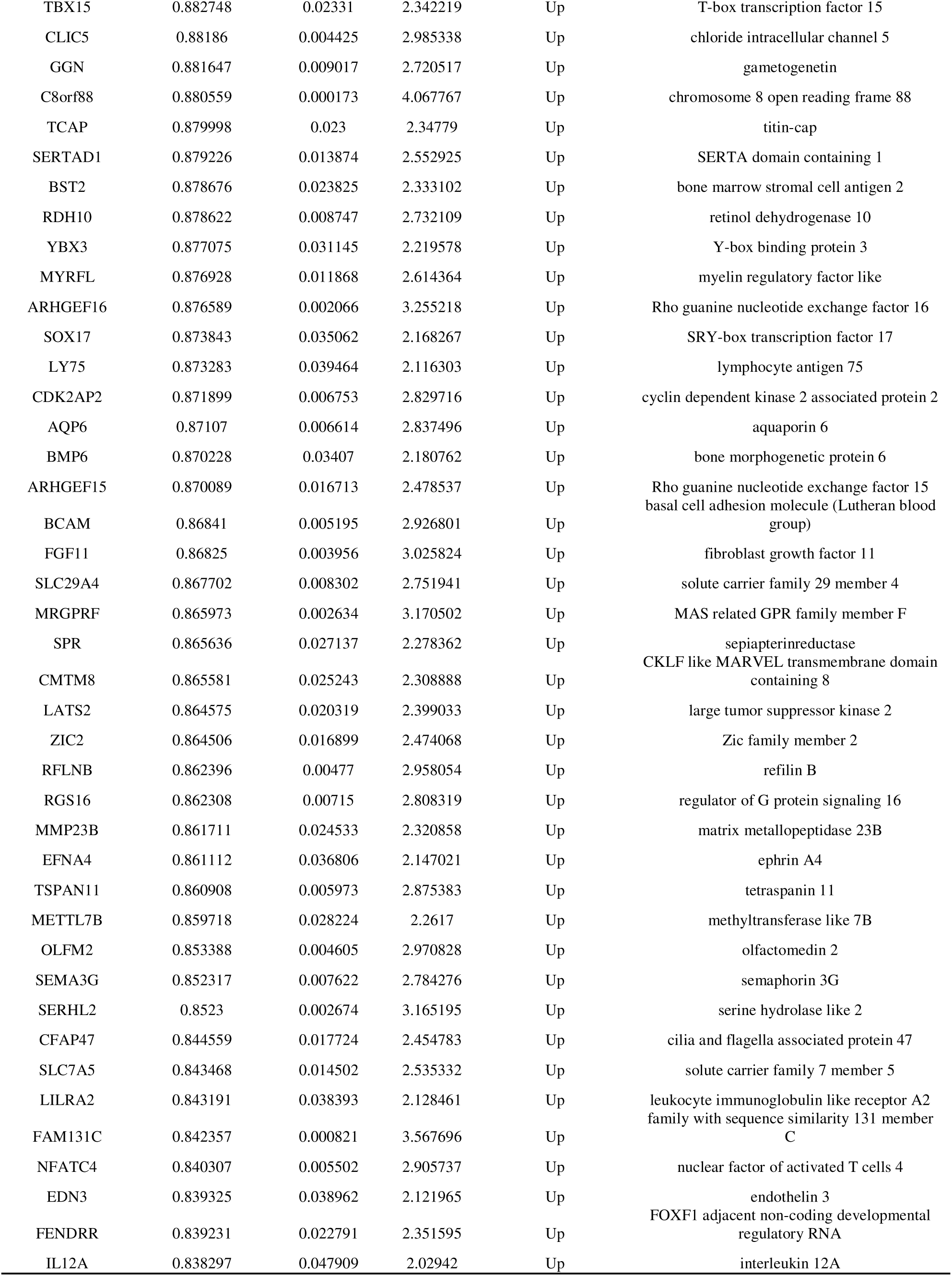
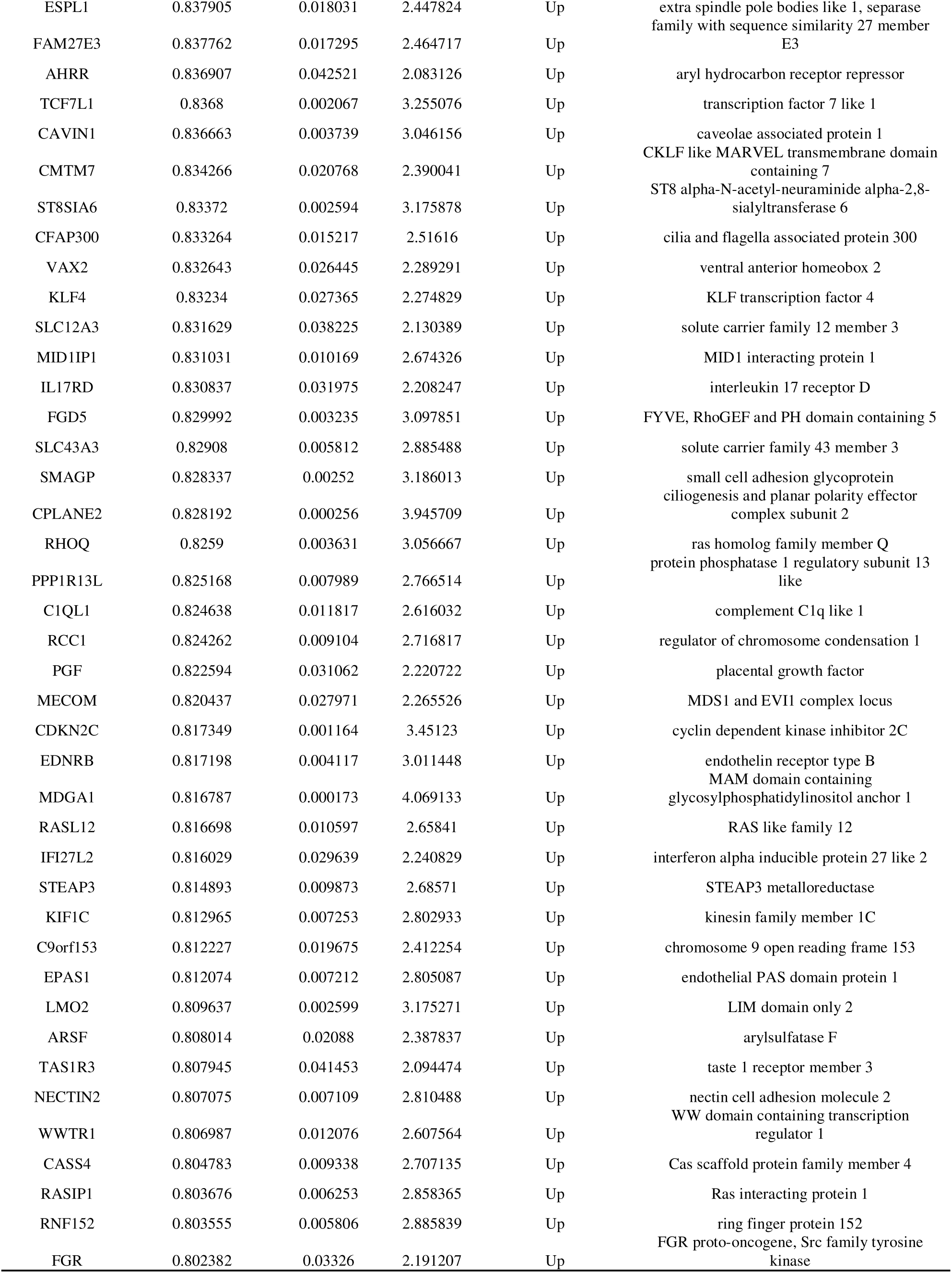
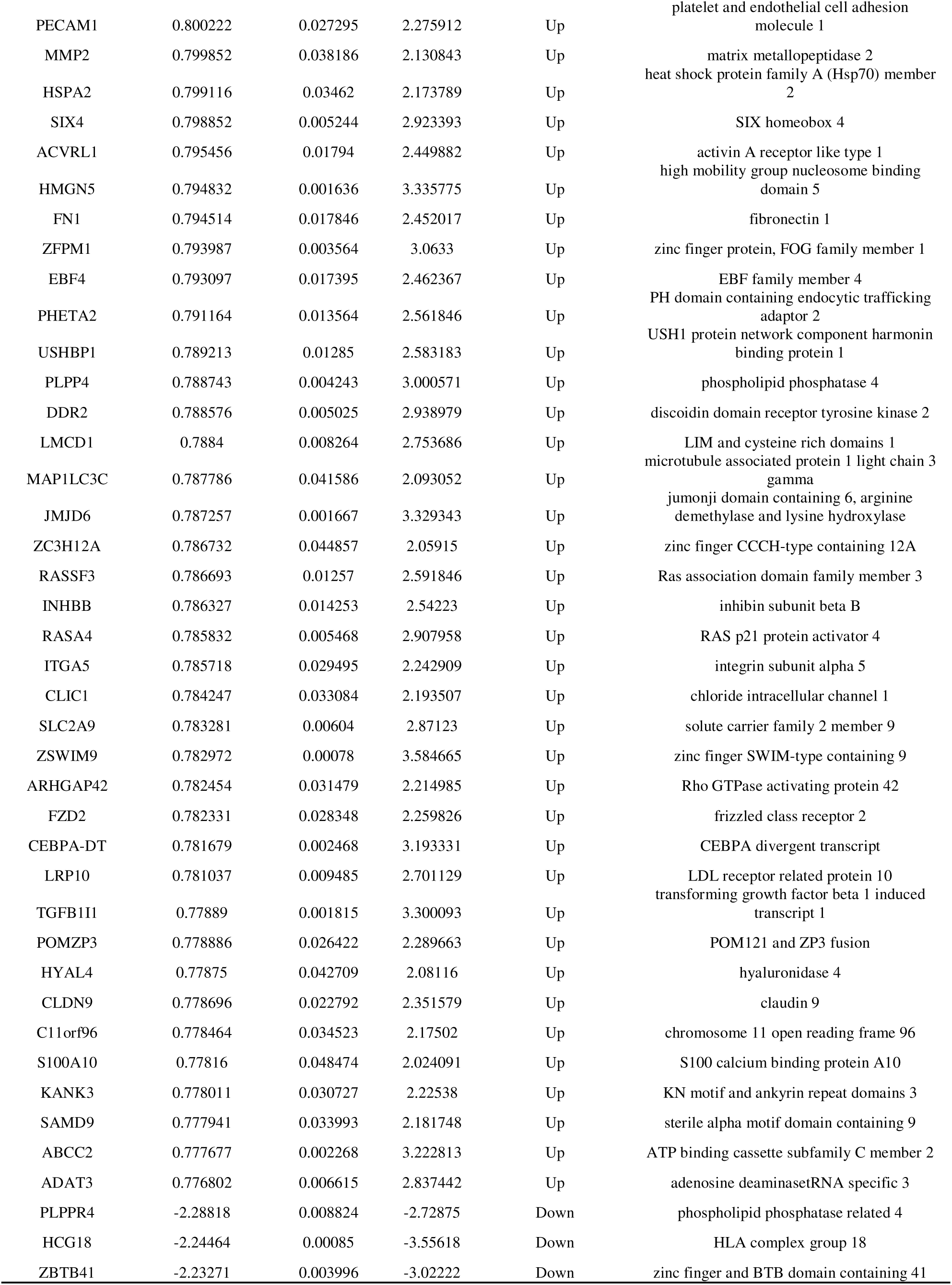
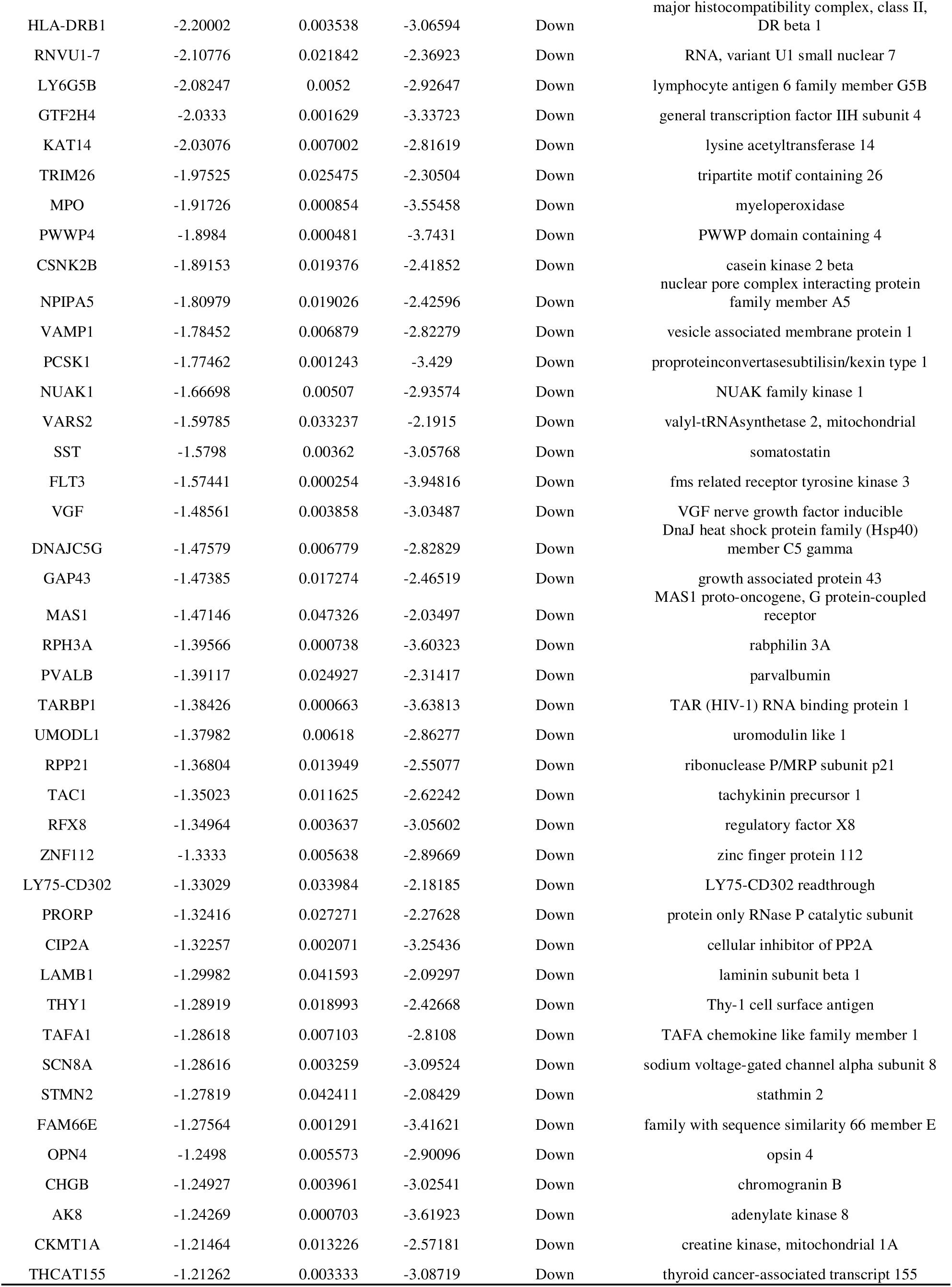
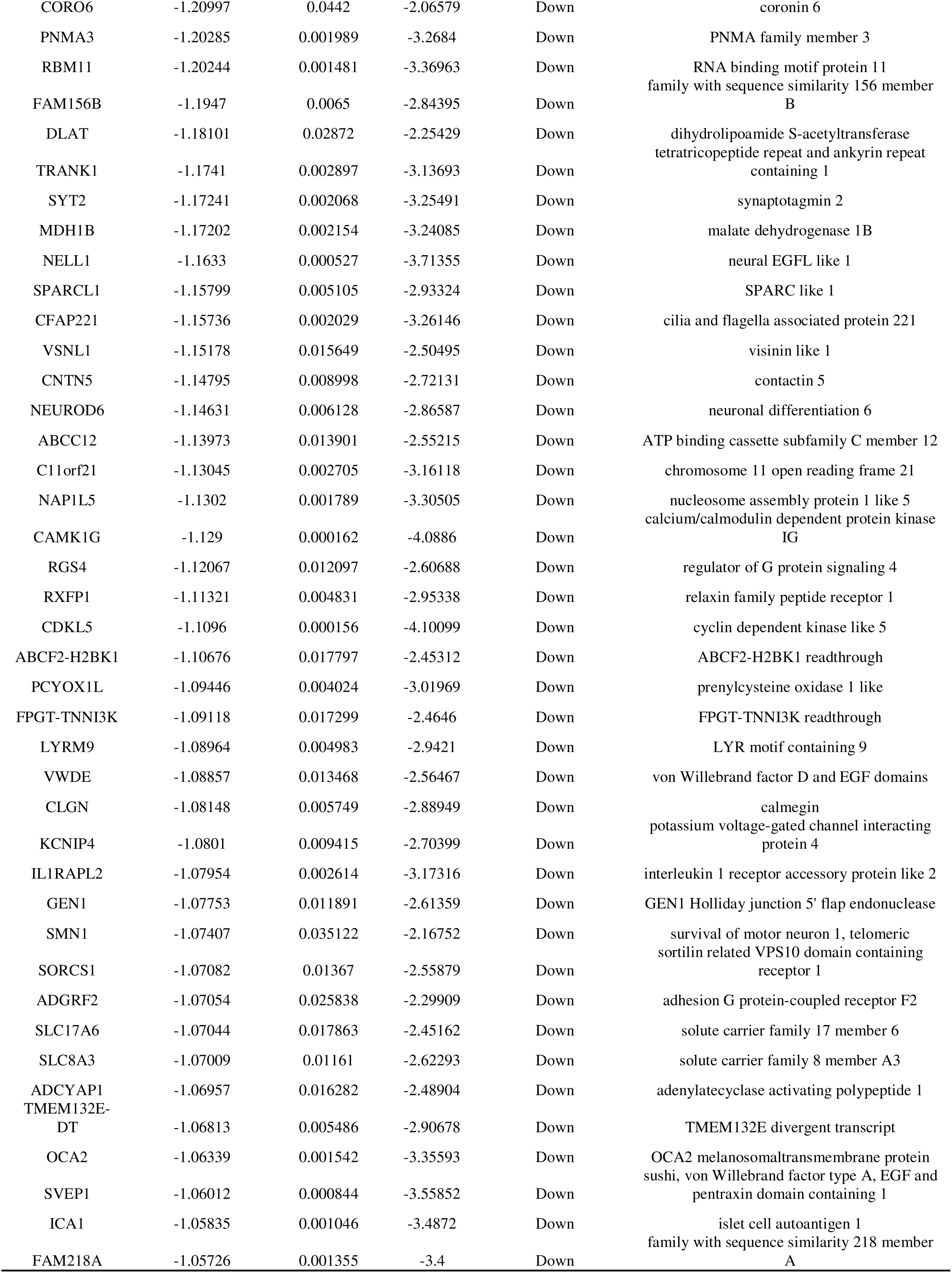
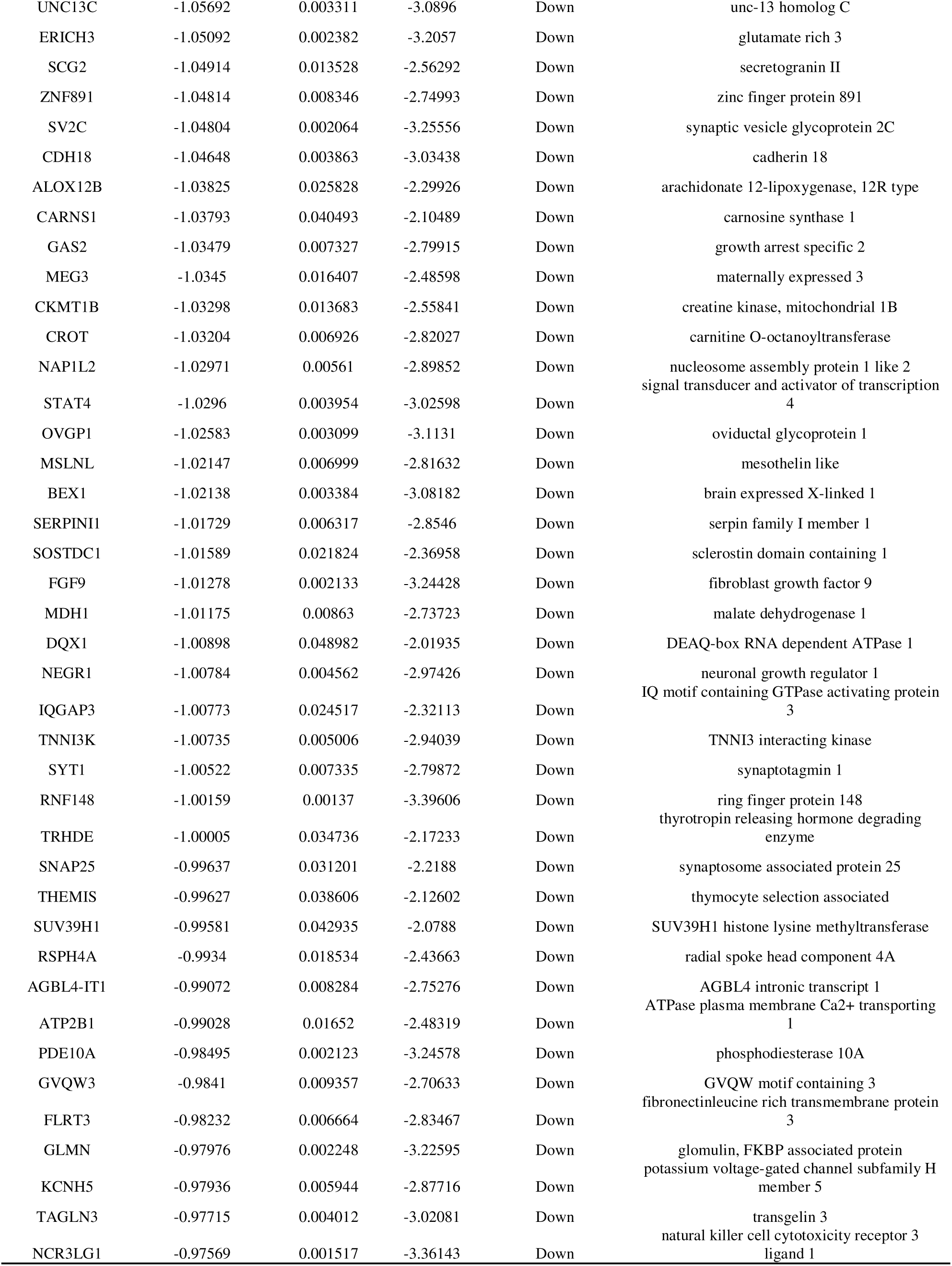
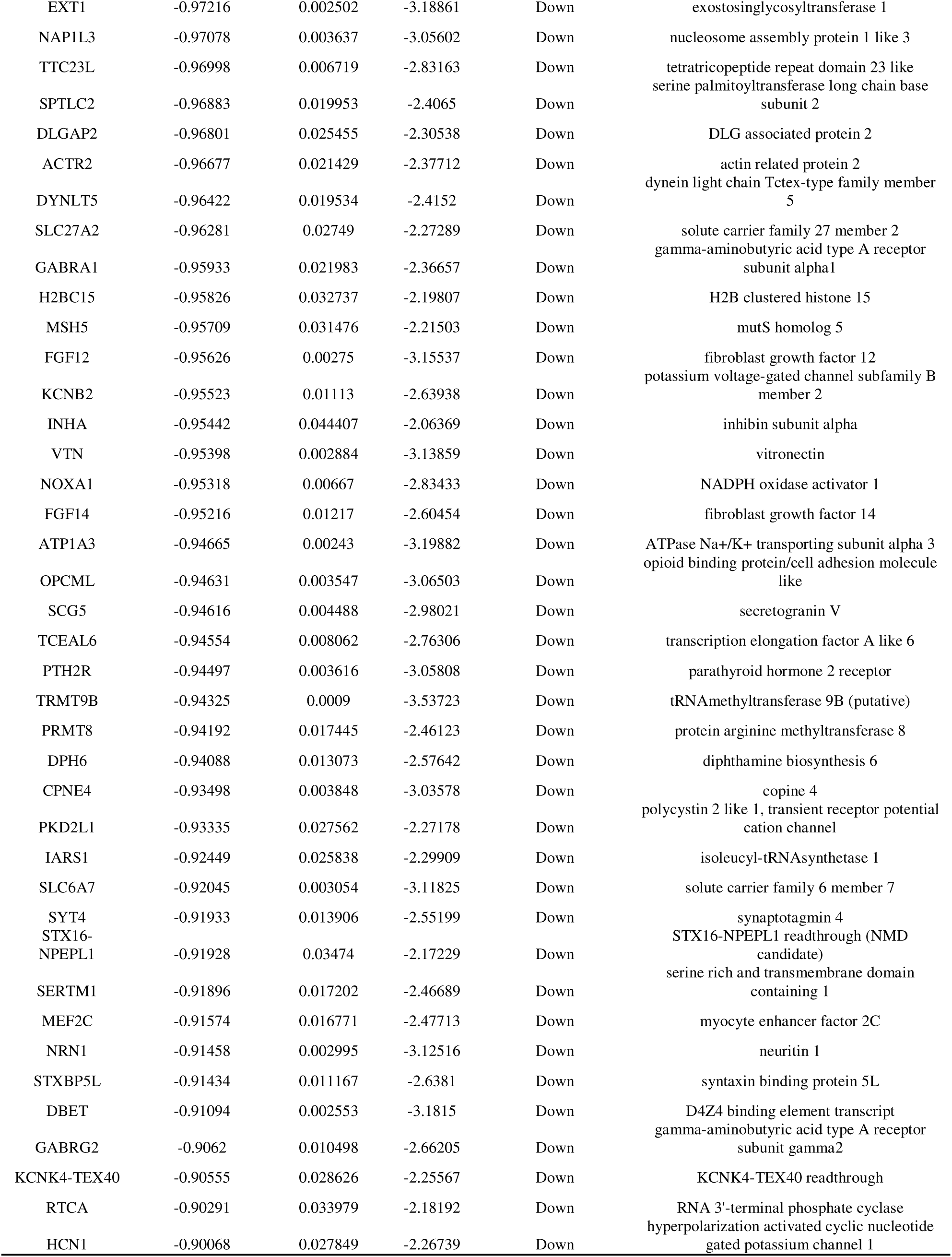
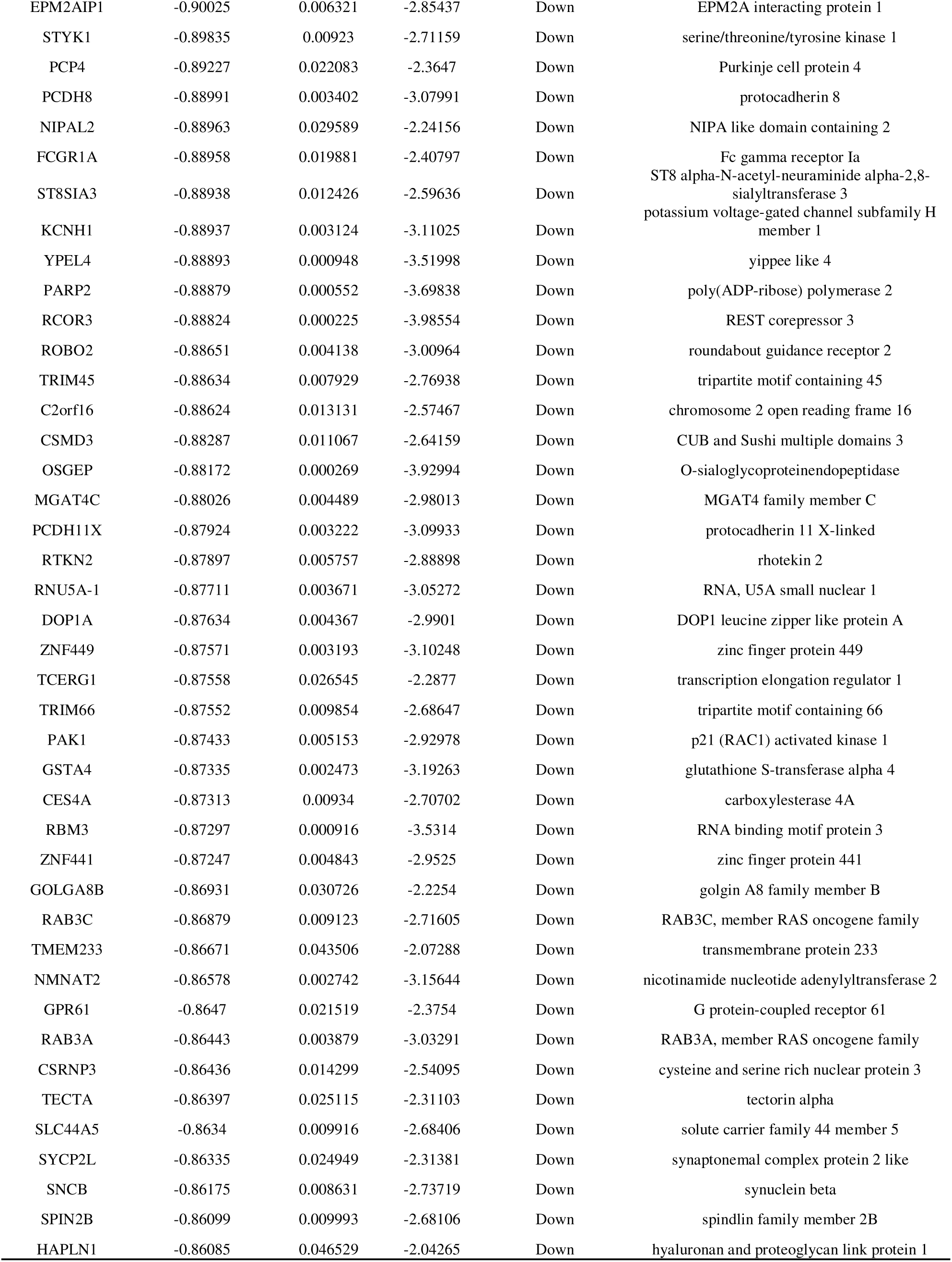
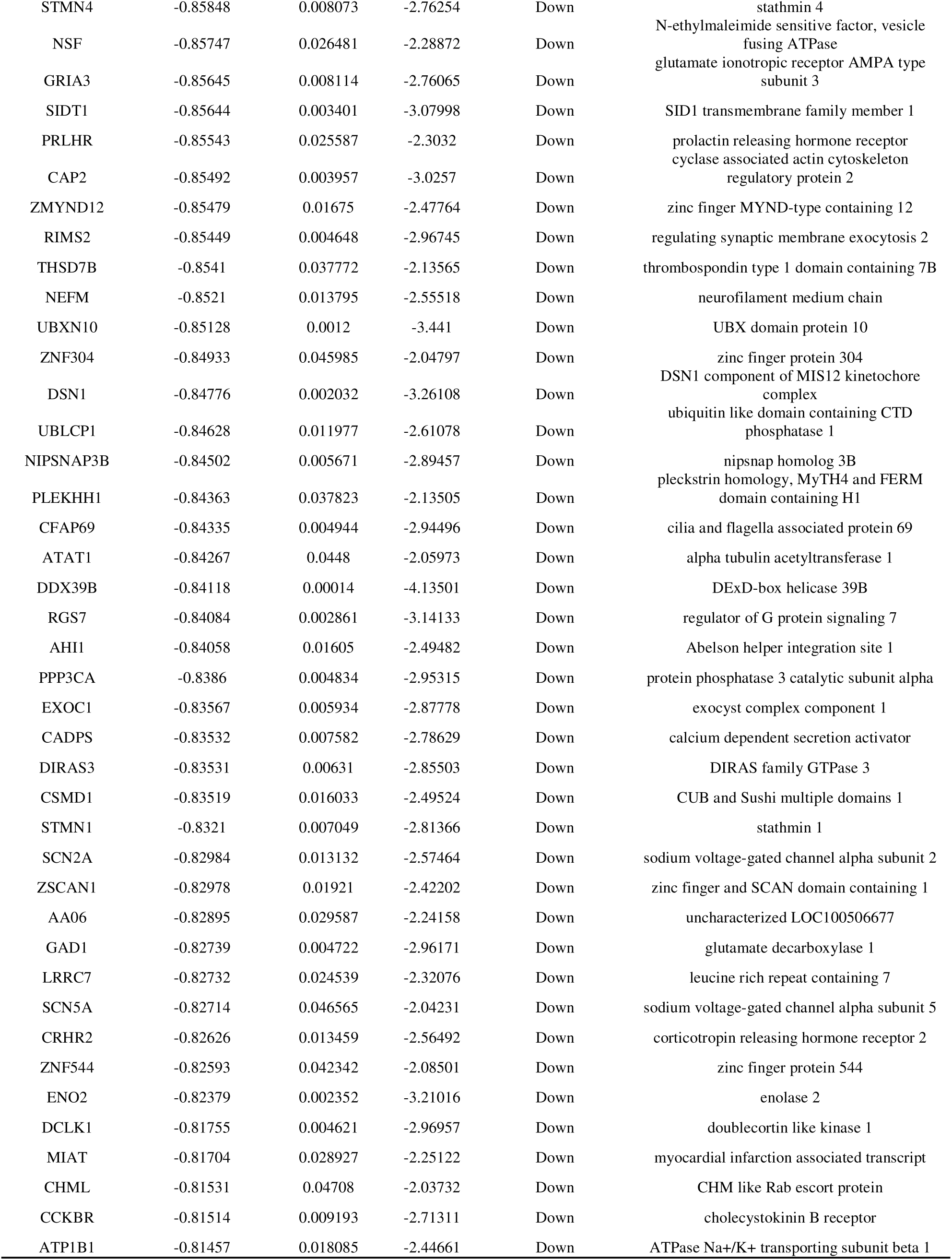
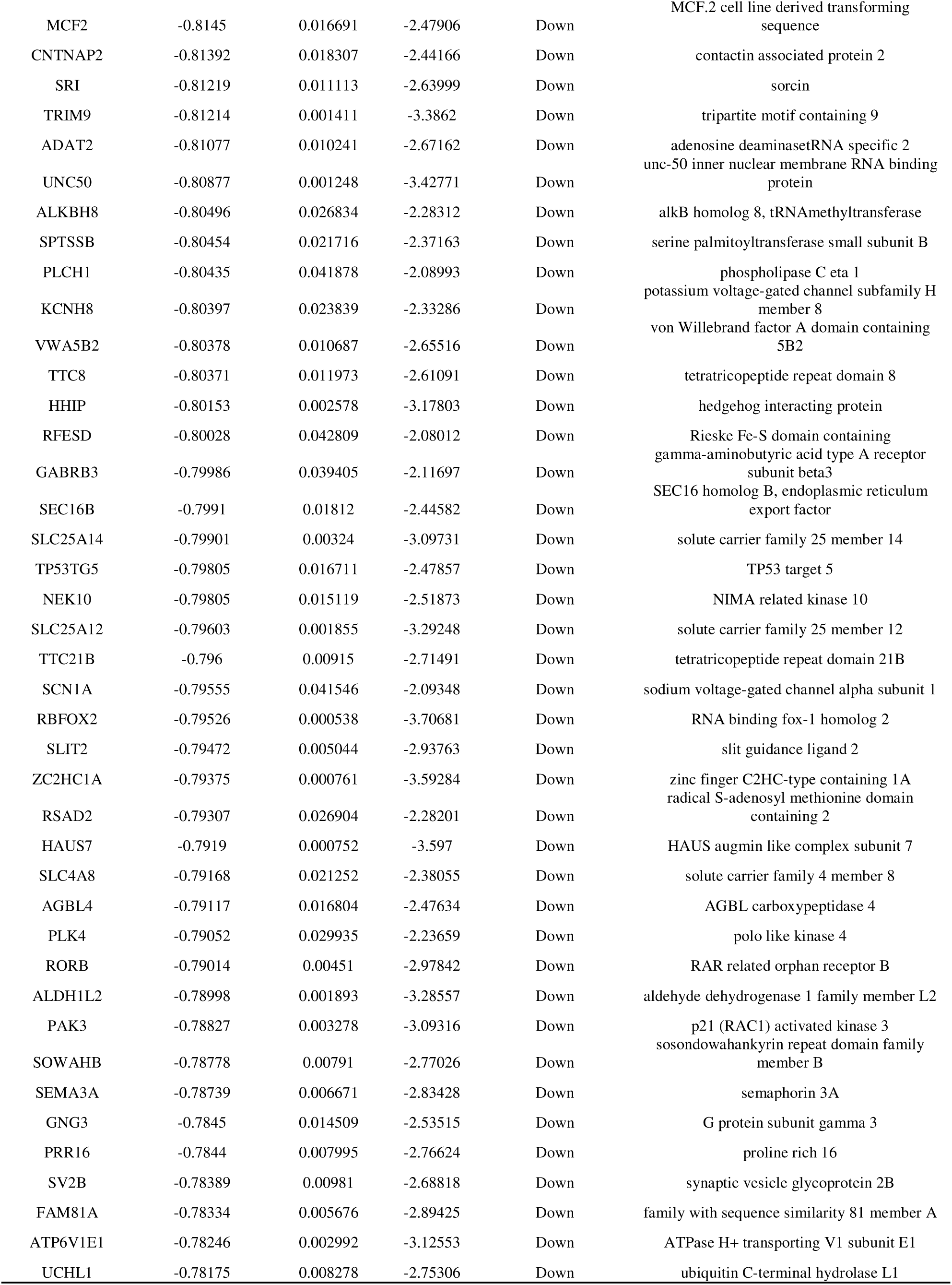
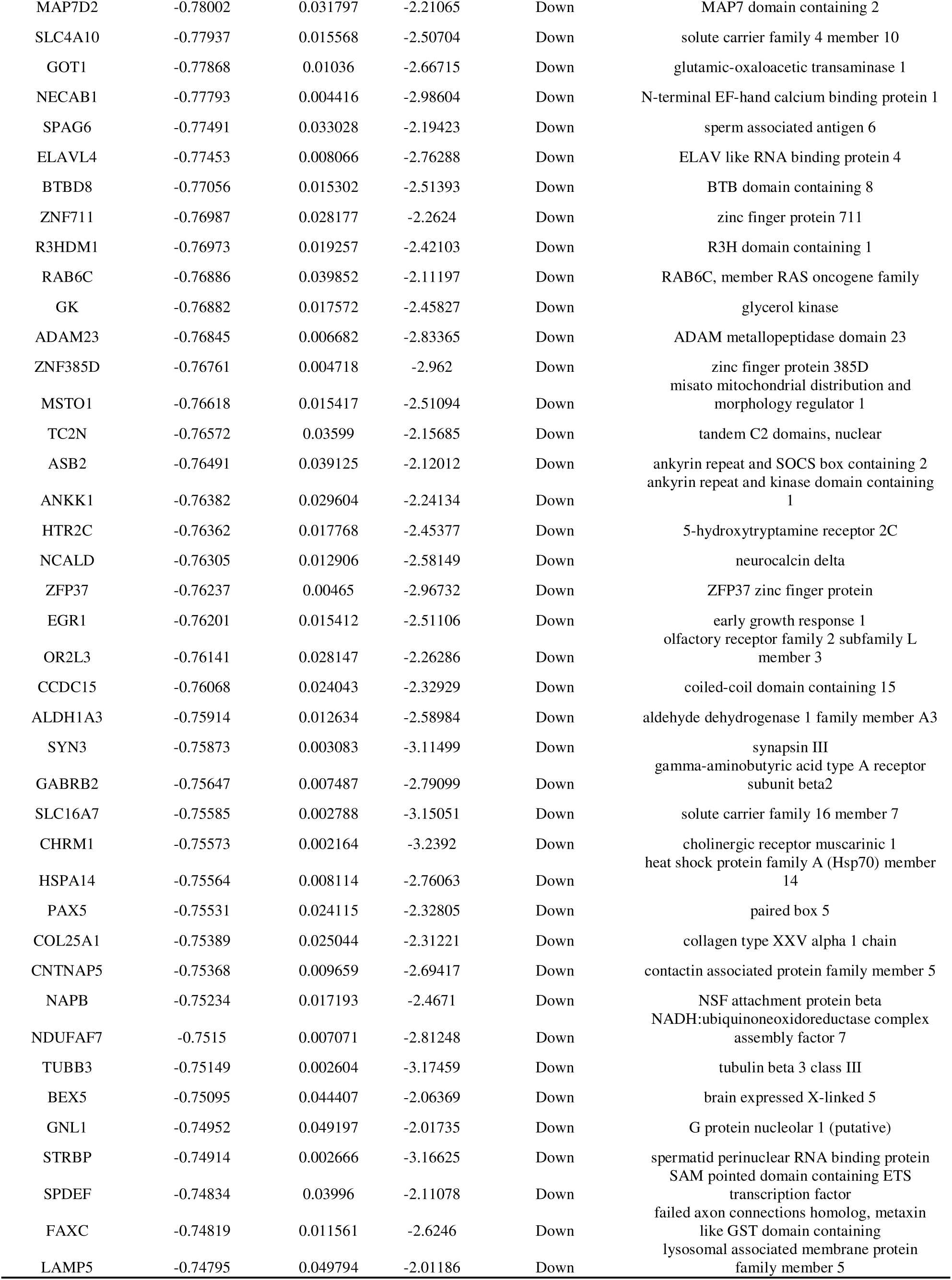
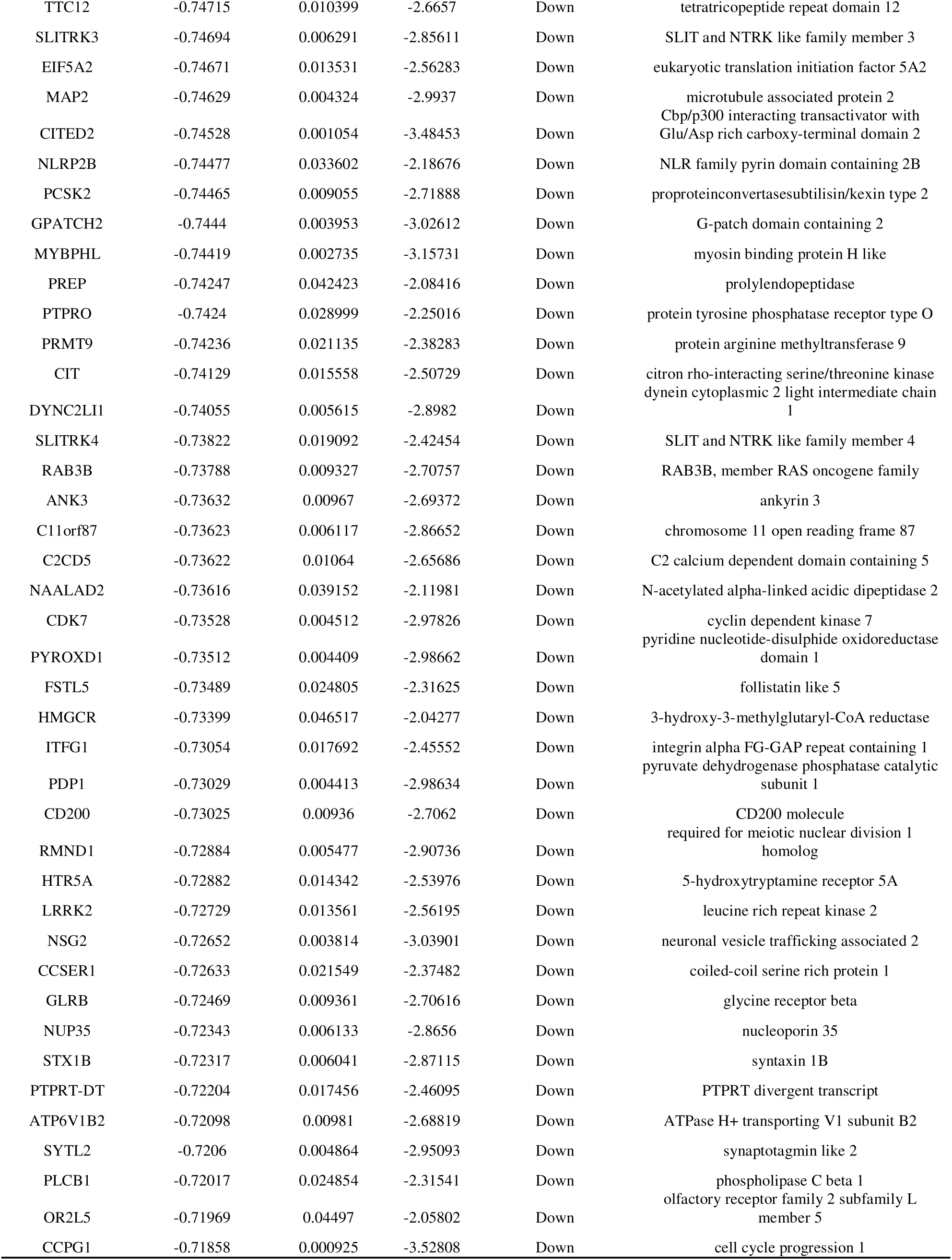
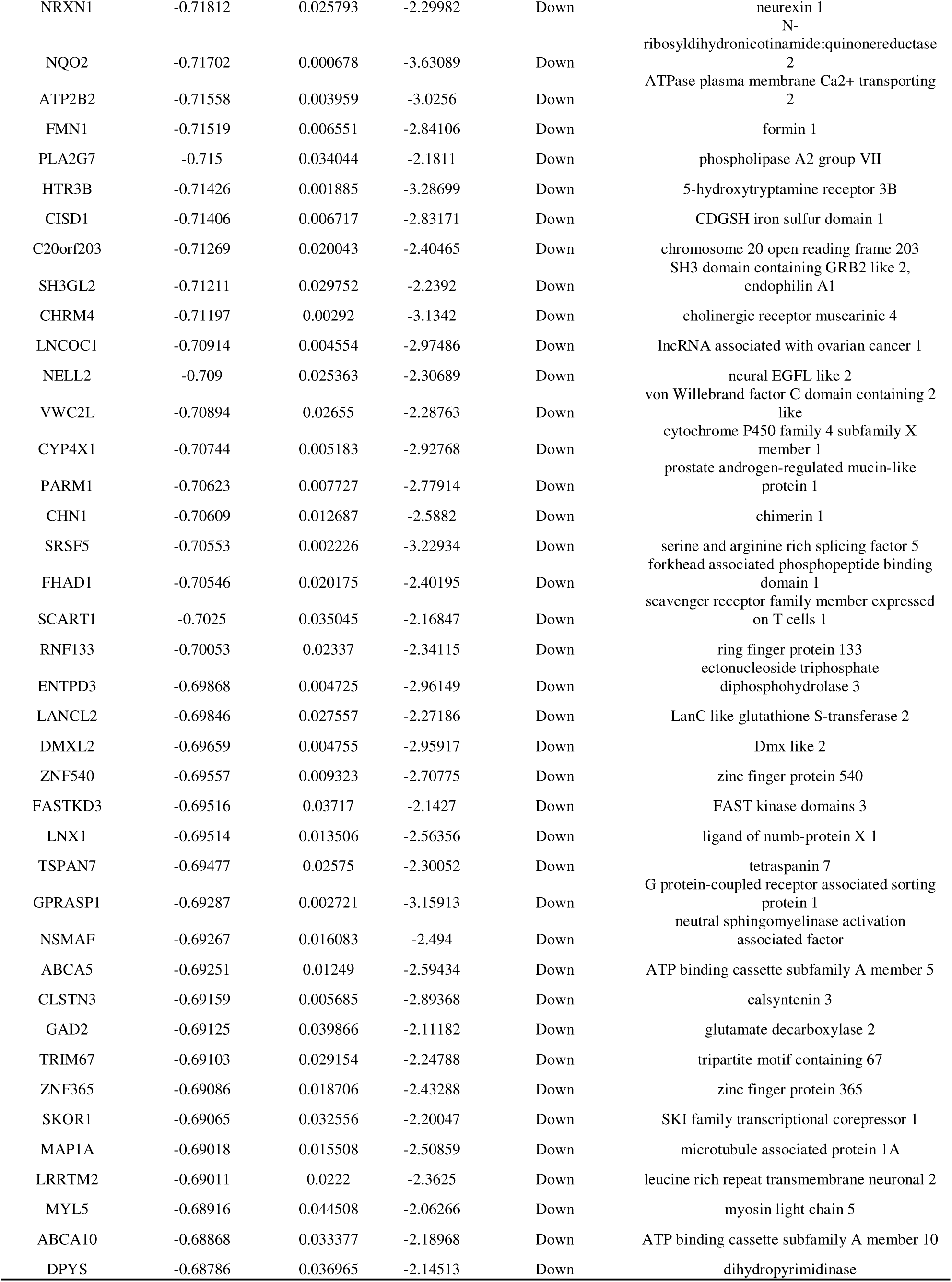
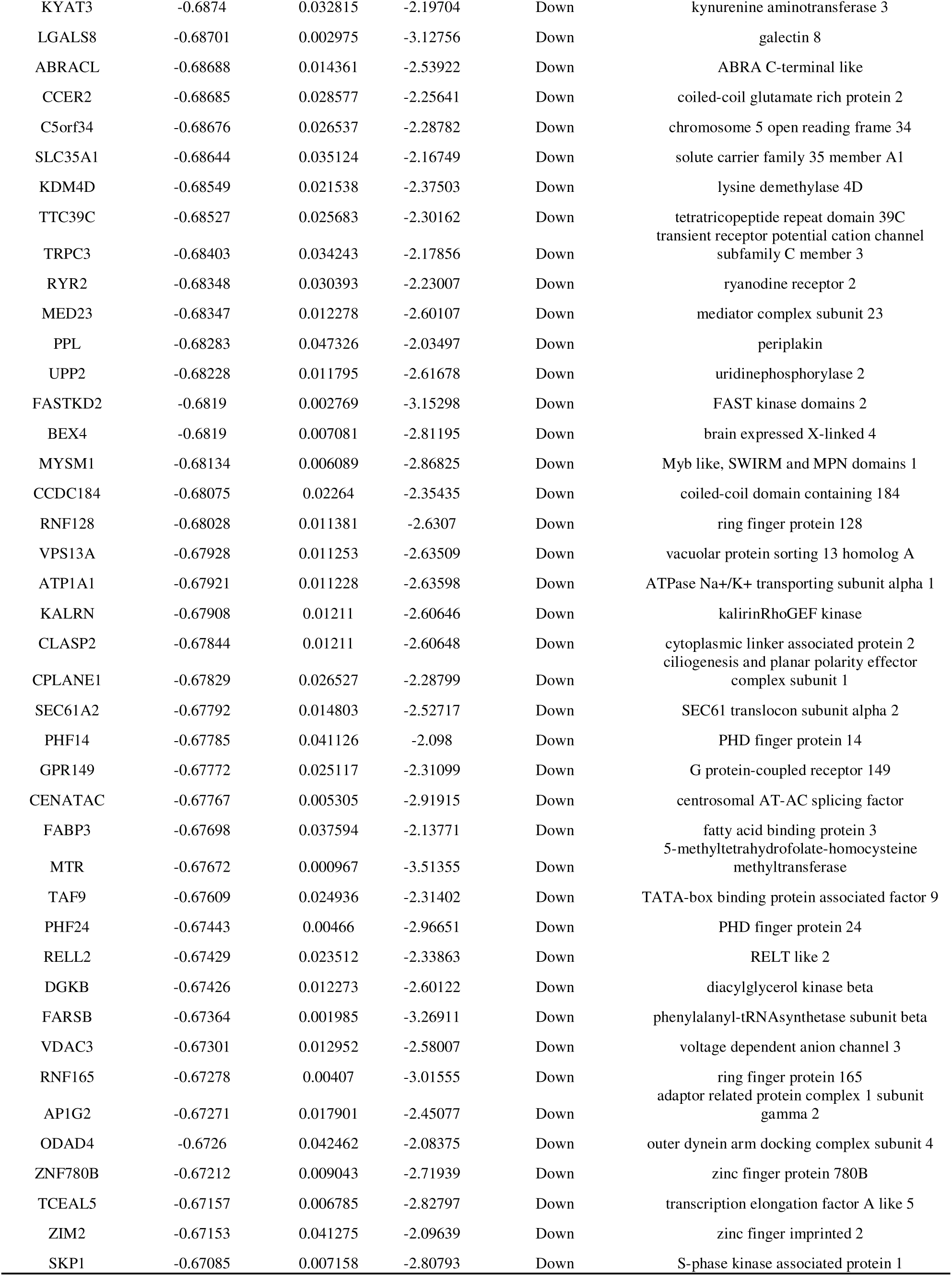
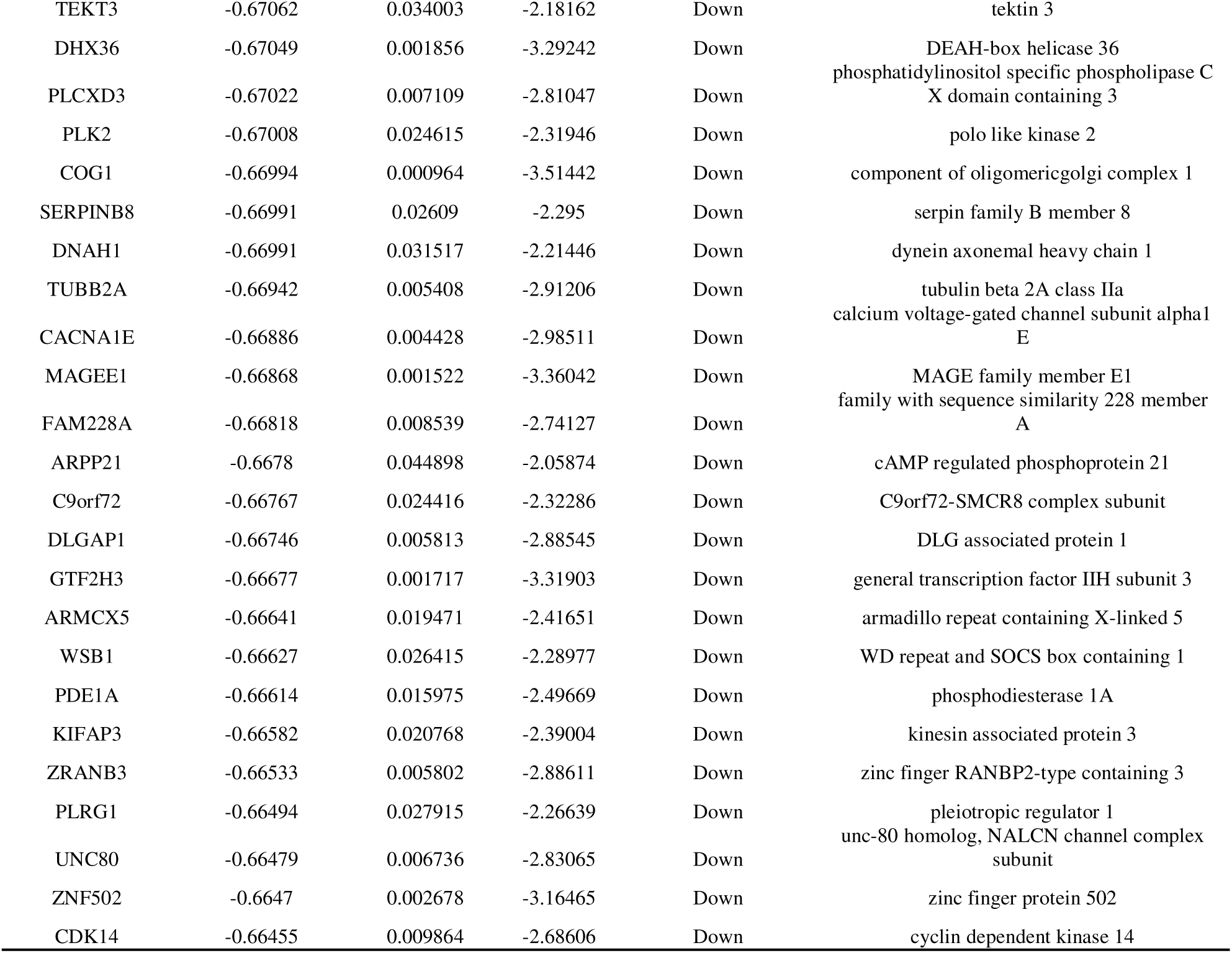
The statistical metrics for key differentially expressed genes (DEGs)

### GO and pathway enrichment analyses of DEGs **|**

With the purpose of investigating the biological activities and pathways associated with DEGs among the AD cases, GO and REACTOME pathway enrichment analyses were performed on the 958 DEGs. Besides, the significantly enriched GO terms were cellular response to response to stimulus, biological regulation, multicellular organism development, developmental process (BP); cytoplasm, membrane, cell junction, plasma membrane (CC); small molecule binding, identical protein binding, ion binding, anion binding (MF) (Table 2). As revealed by the REACTOME enrichment analysis, the 958 DEGs were mostly associated with signal transduction, muscle contraction, cardiac conduction and neuronal system (Table 3).

**Table 2.**
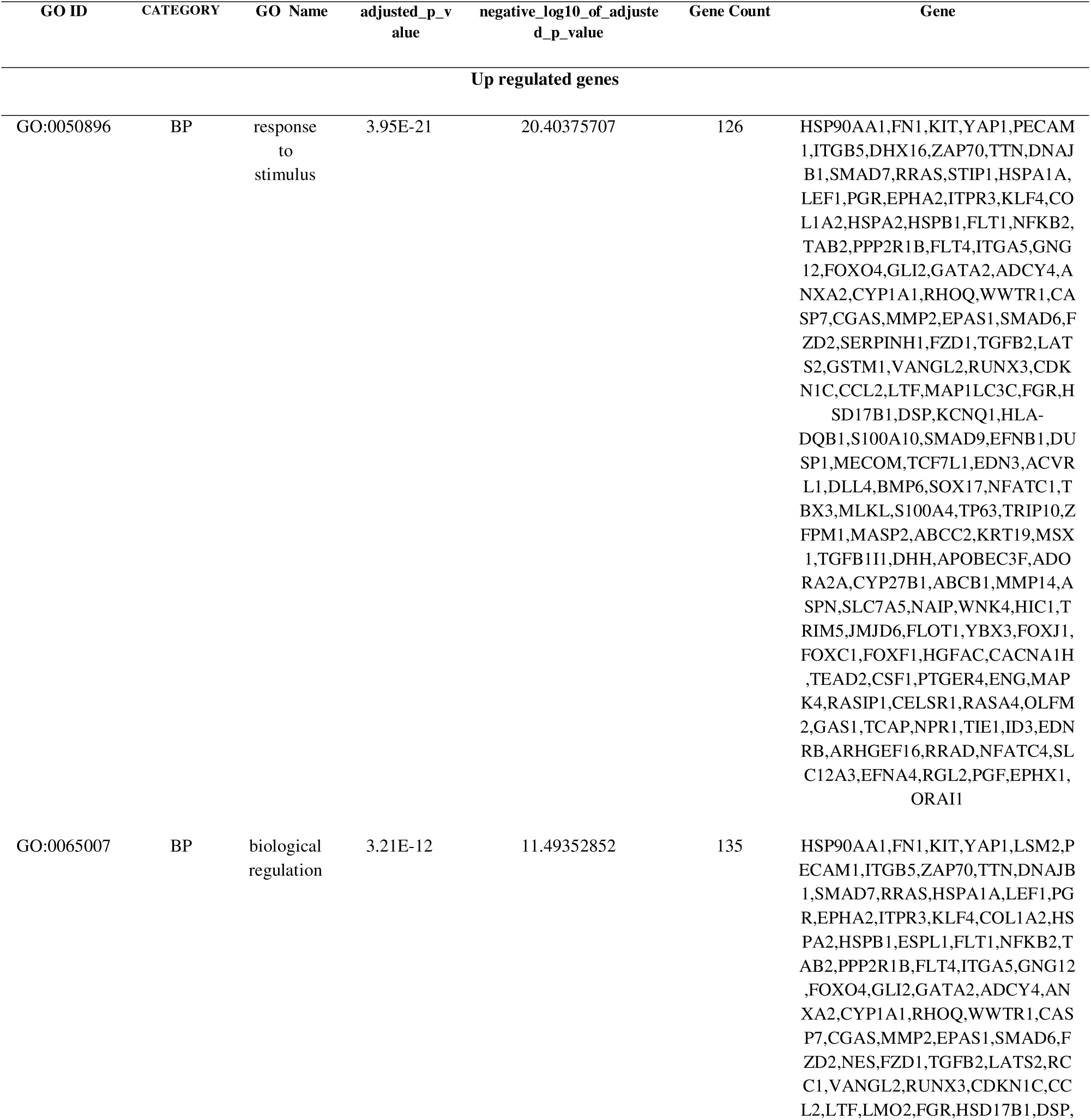
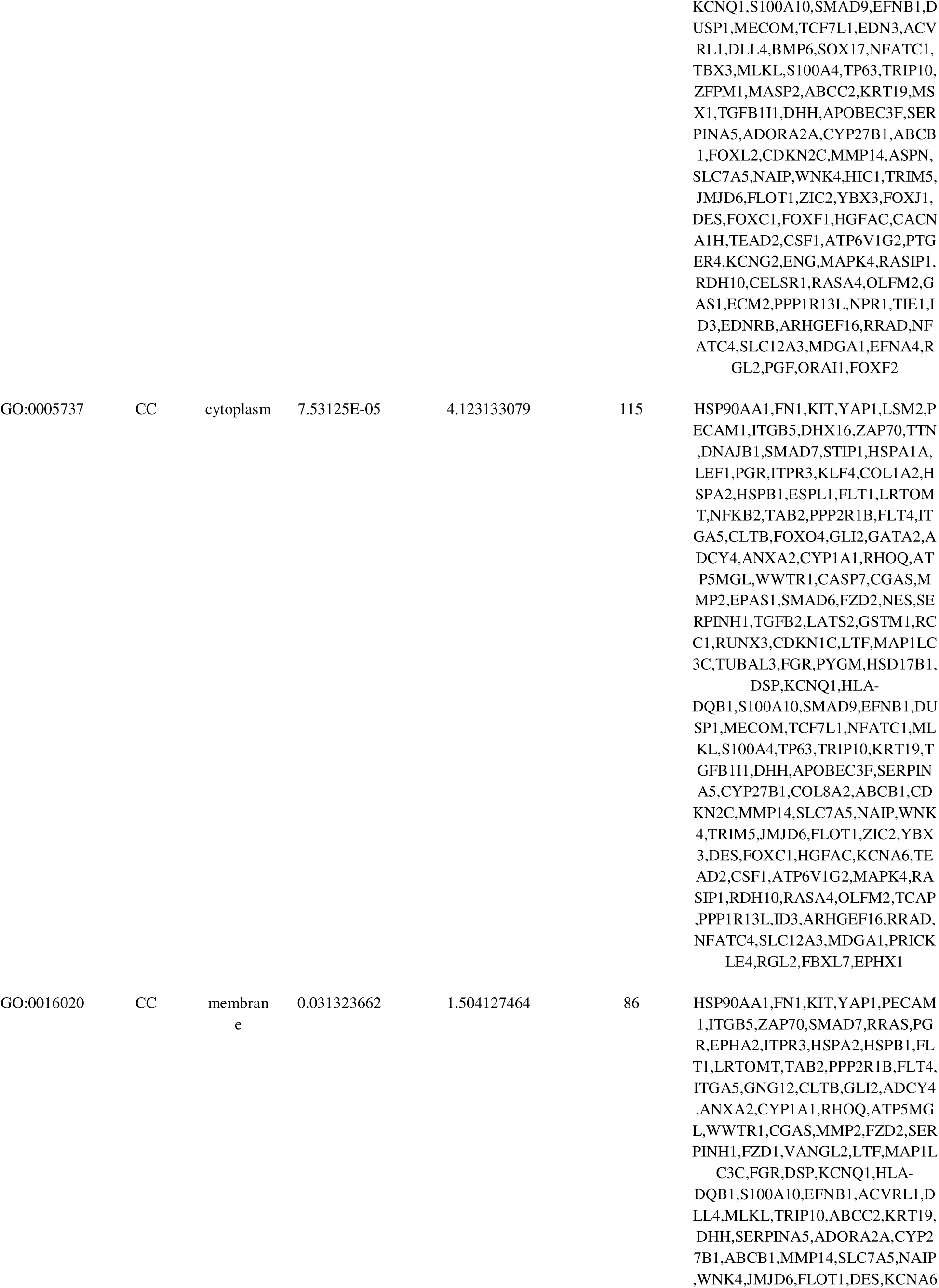
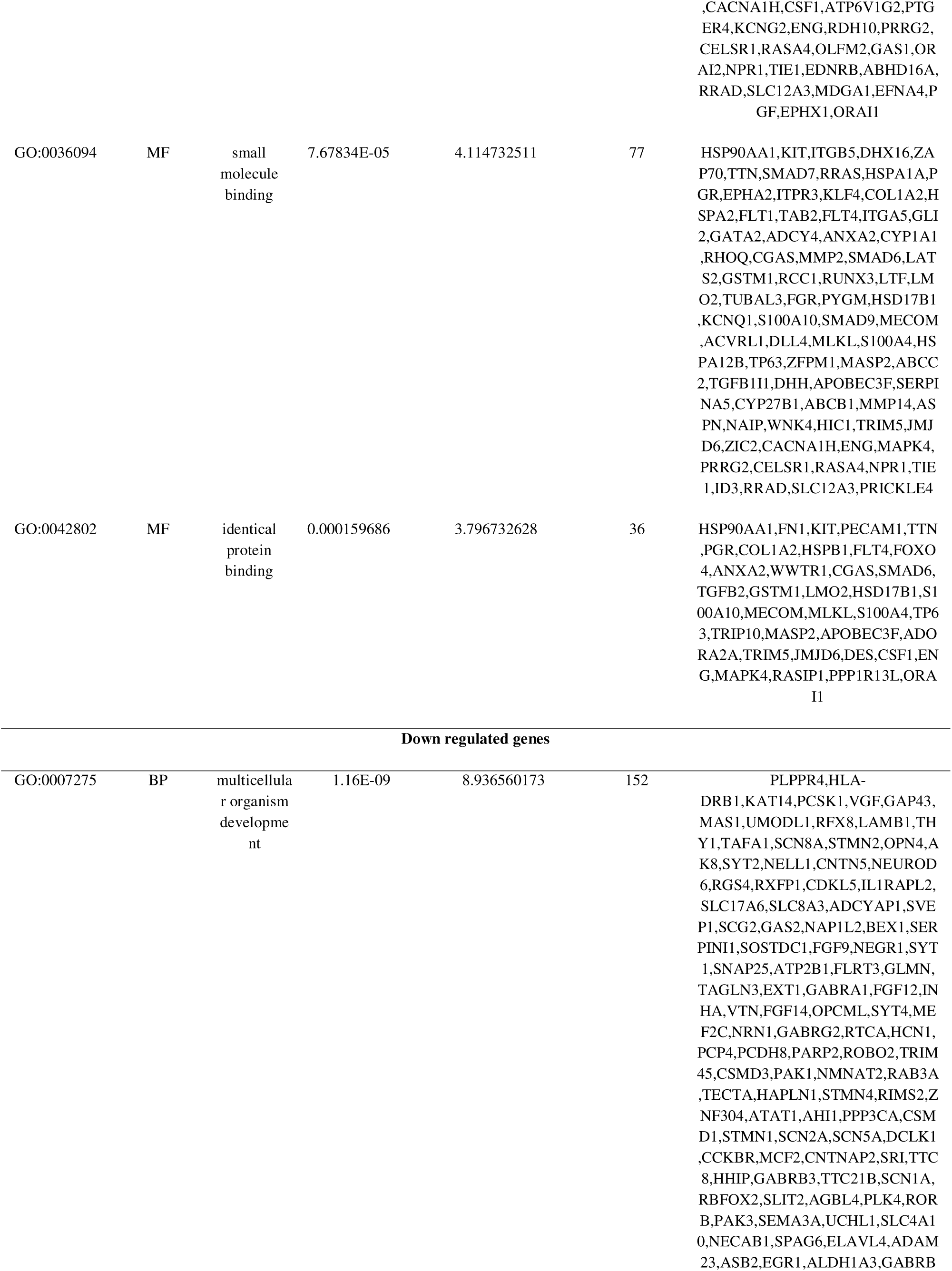
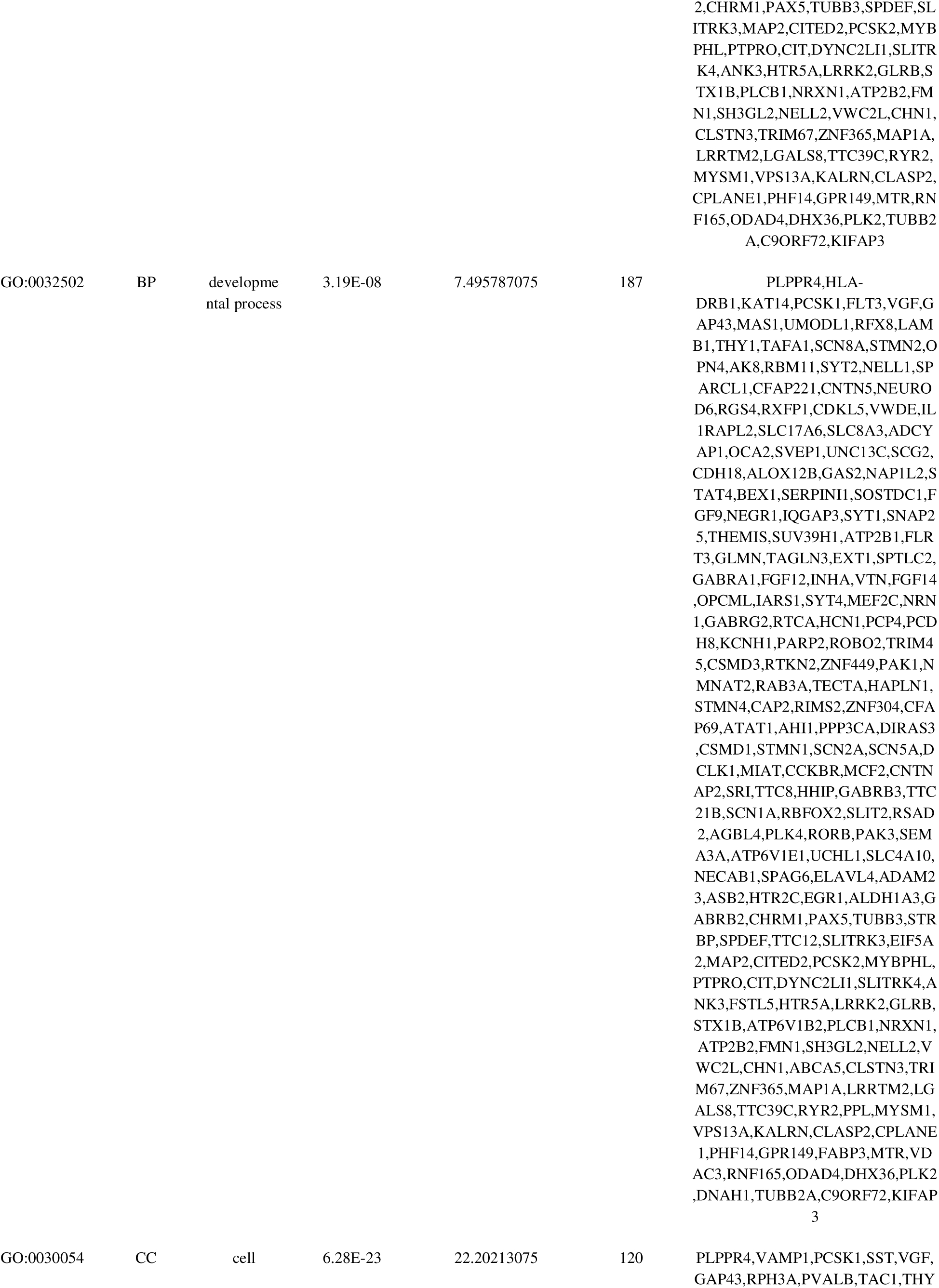
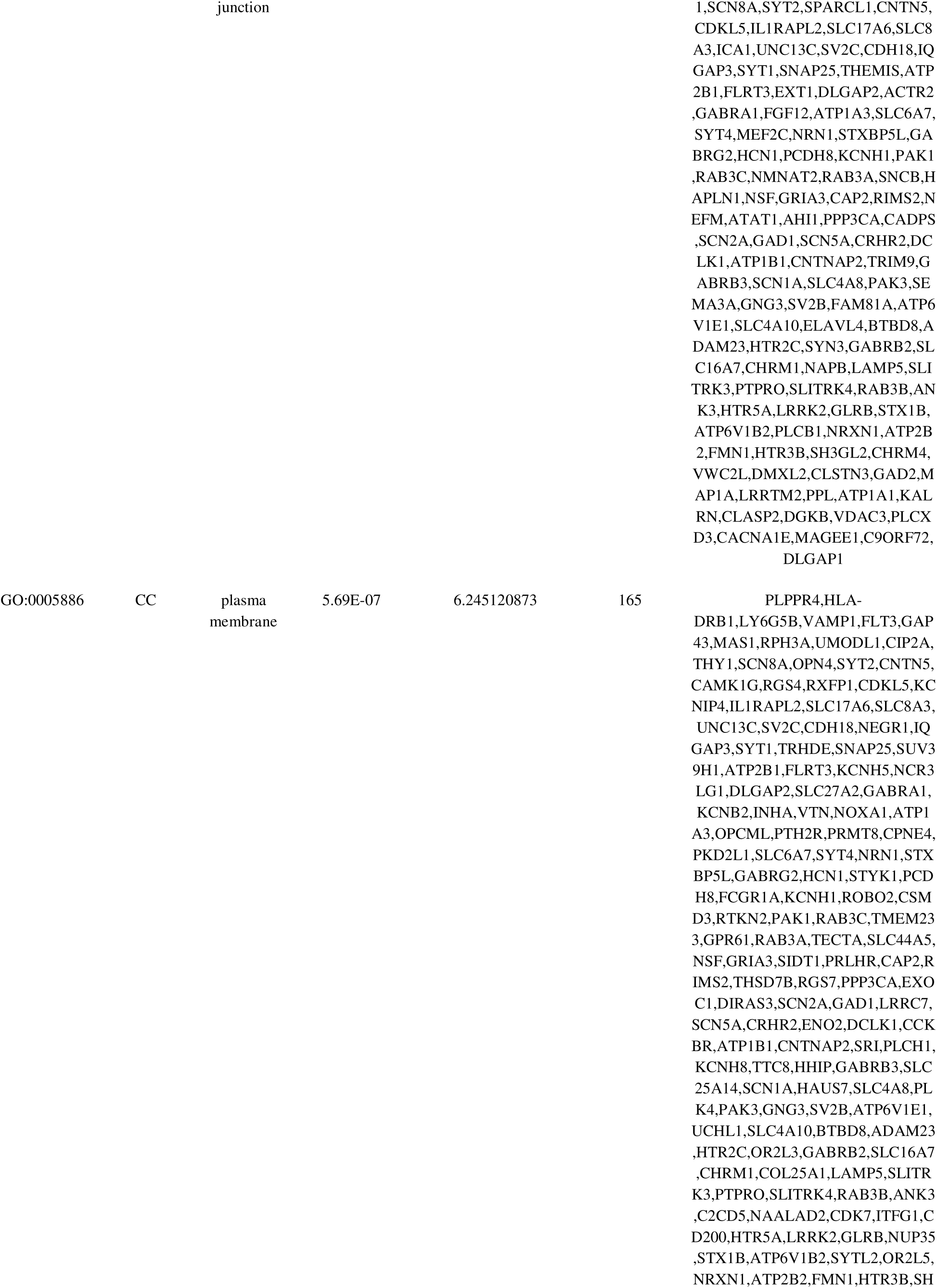
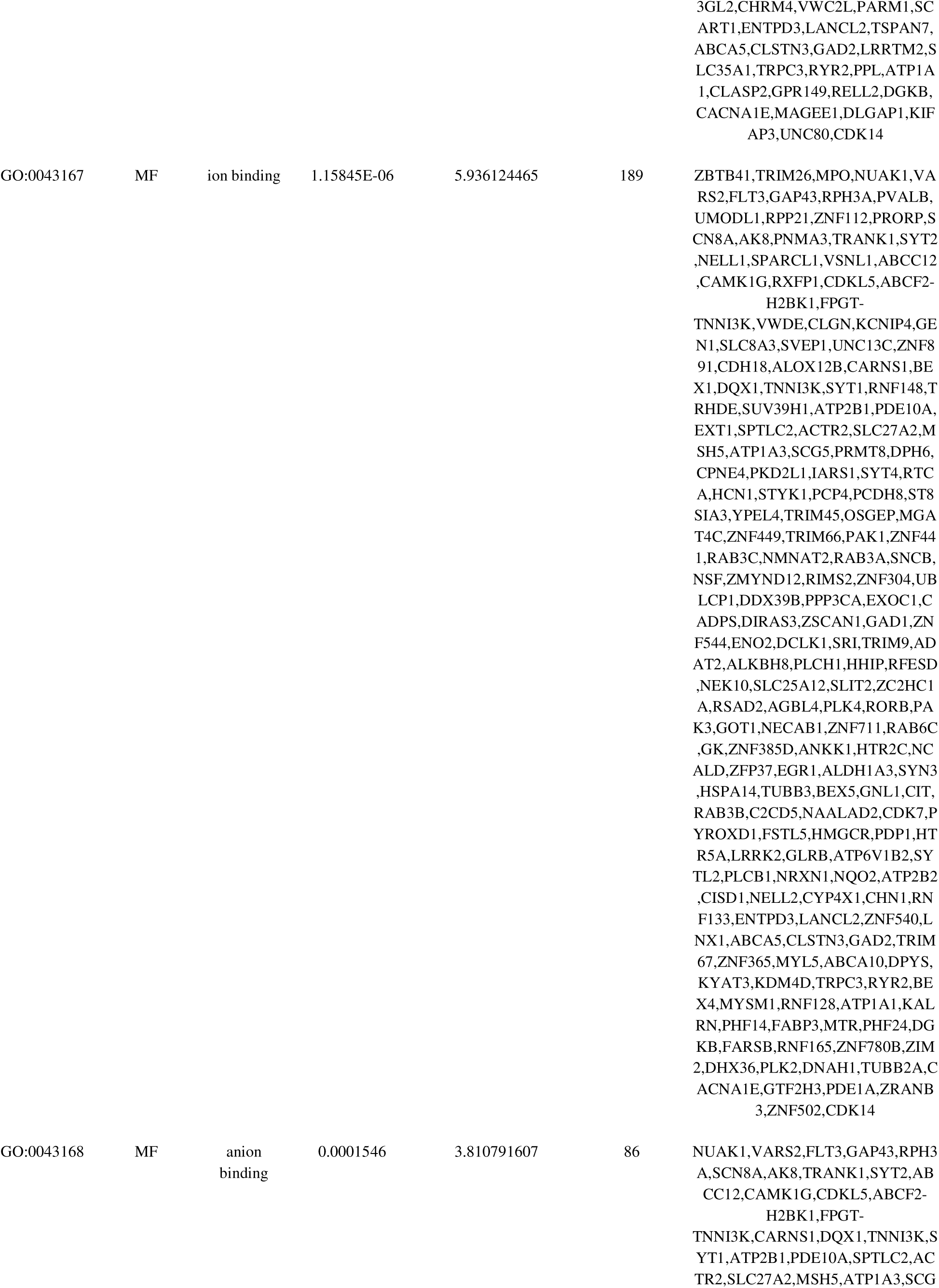
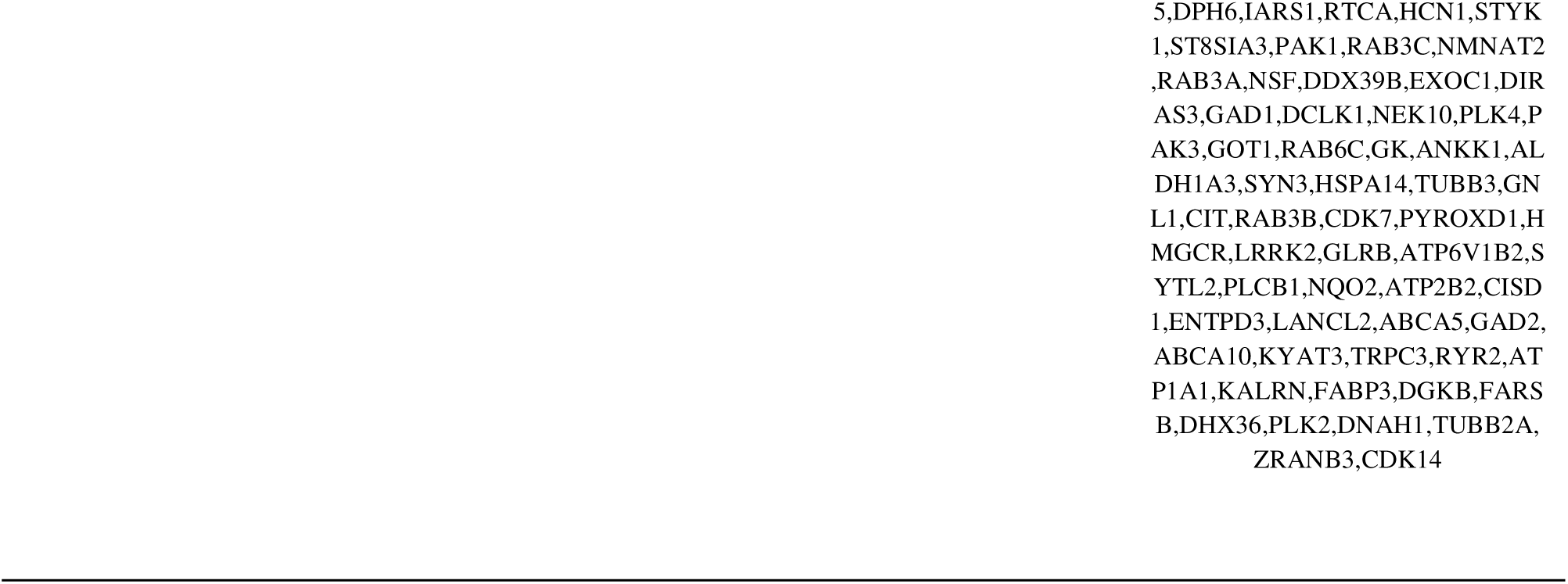
The enriched GO terms of the up and down regulated differentially expressed genes.

**Table 3.**
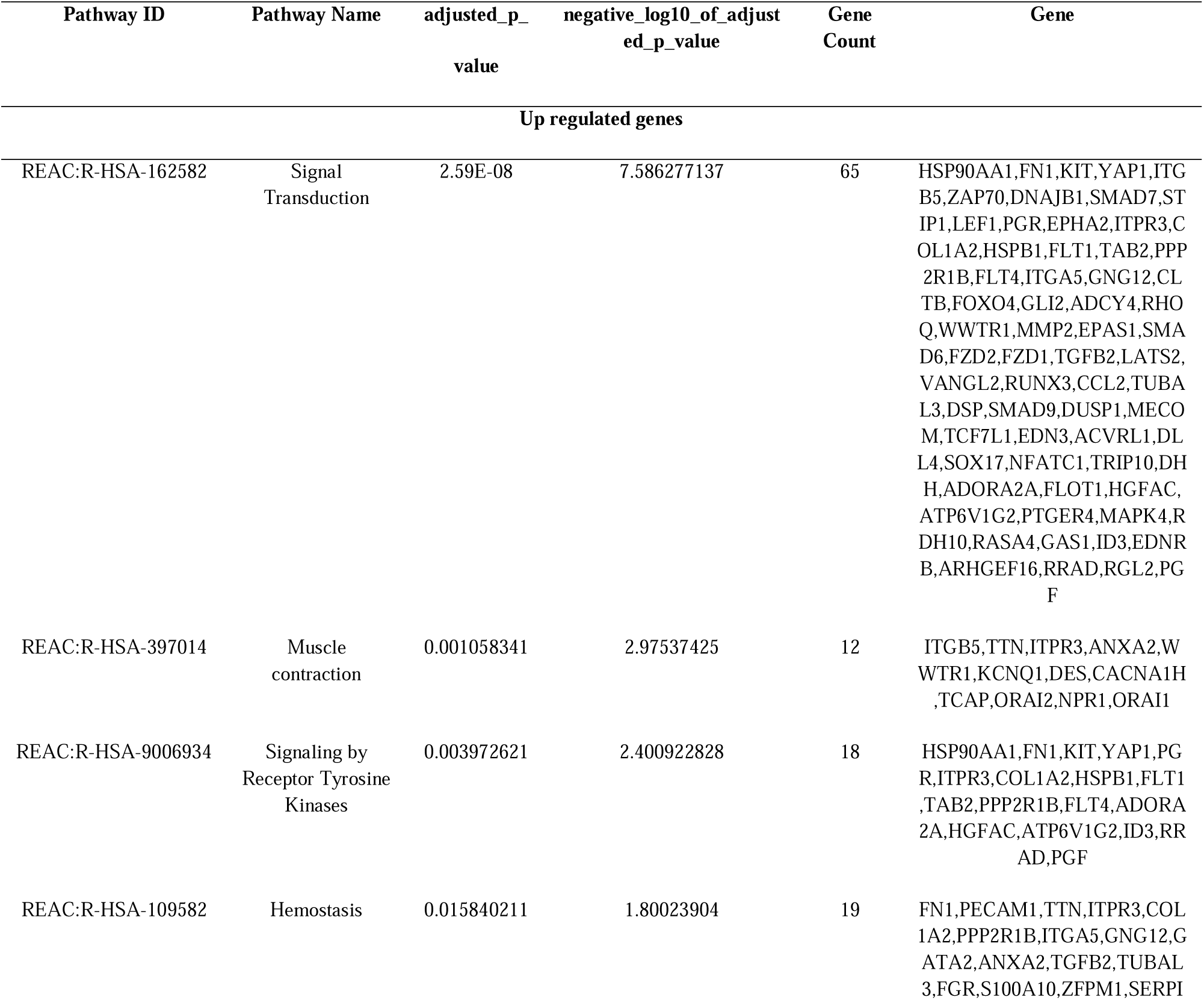
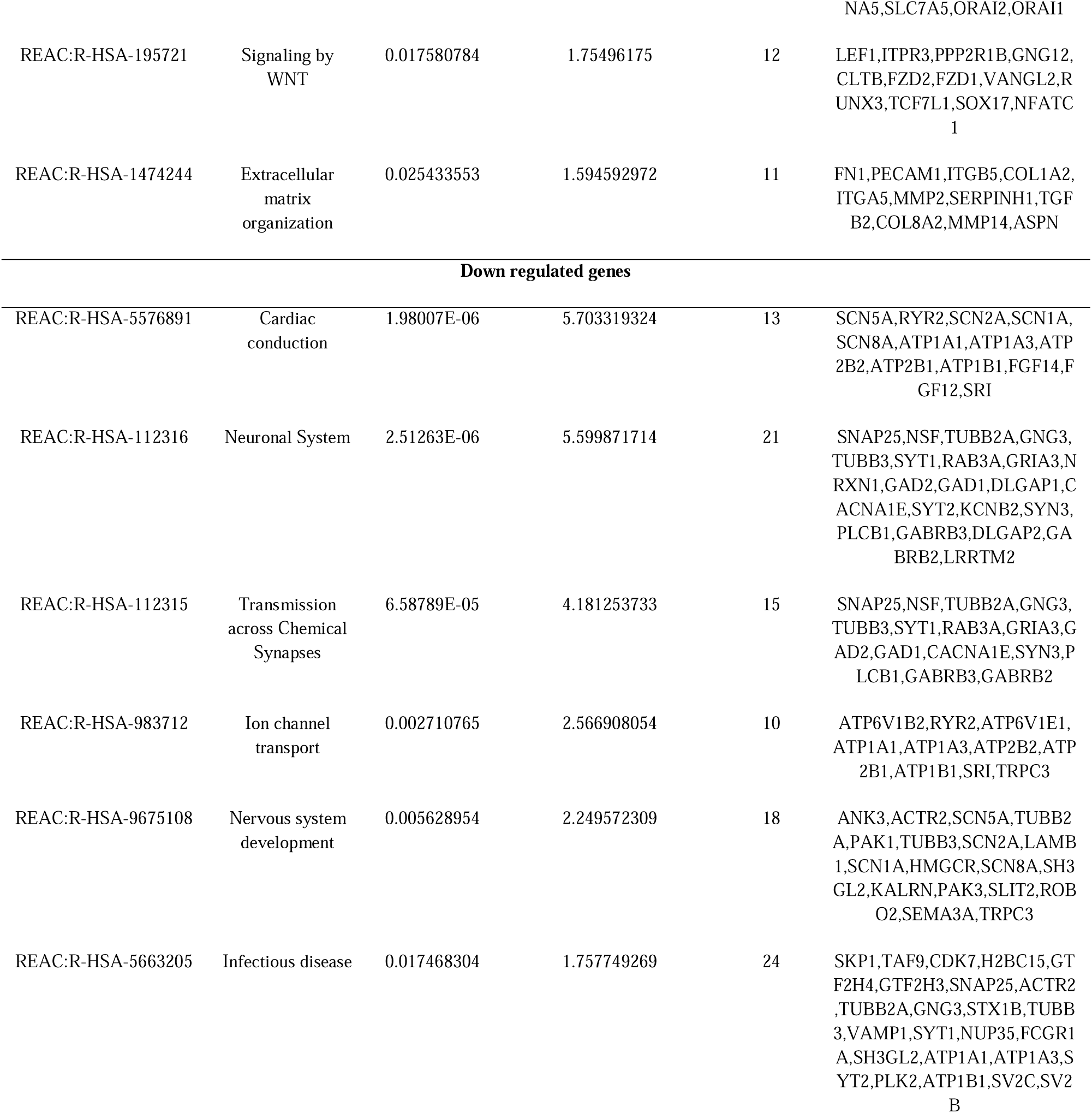
The enriched pathway terms of the up and down regulated differentially expressed genes.

### Construction of the PPI network and module analysis

To analyze the PPI network of the 958 DEGs, the database STRING was applied. Cytoscape network visualization was obtained on the basis of the STRING database (Fig. 3). Analysis of PPI networks of AD DEGs revealed 4886 nodes and 10342 edges, respectively. Using the node degree, betweenness, stress and closeness algorithms, 10 hub genes were obtained through the Network Analyzer plugin of the software Cytoscape. Subsequently, according to the rank score, seven first-level hub genes, including HSP90AA1, FN1, KIT, YAP1, LSM2, SKP1, EIF5A2, TAF9, DDX39B and CDK7 were identified (Table 4). The PEWCC in the database was used for further module analysis. Two significant modules were obtained according to the degree of interconnection among hub genes. A total of 37 nodes and 179 edges were included in Module 1 (Fig. 4A), and a total of 37 nodes and 179 edges were included in Module 2 (Fig. 4B). Module 1 was mainly enriched in signal transduction, response to stimulus, biological regulation, cytoplasm and small molecule binding. Module 2 was mainly enriched in cell junction, plasma membrane, neuronal system, infectious disease, multicellular organism development and transmission across chemical synapses.

**Fig. 3.**
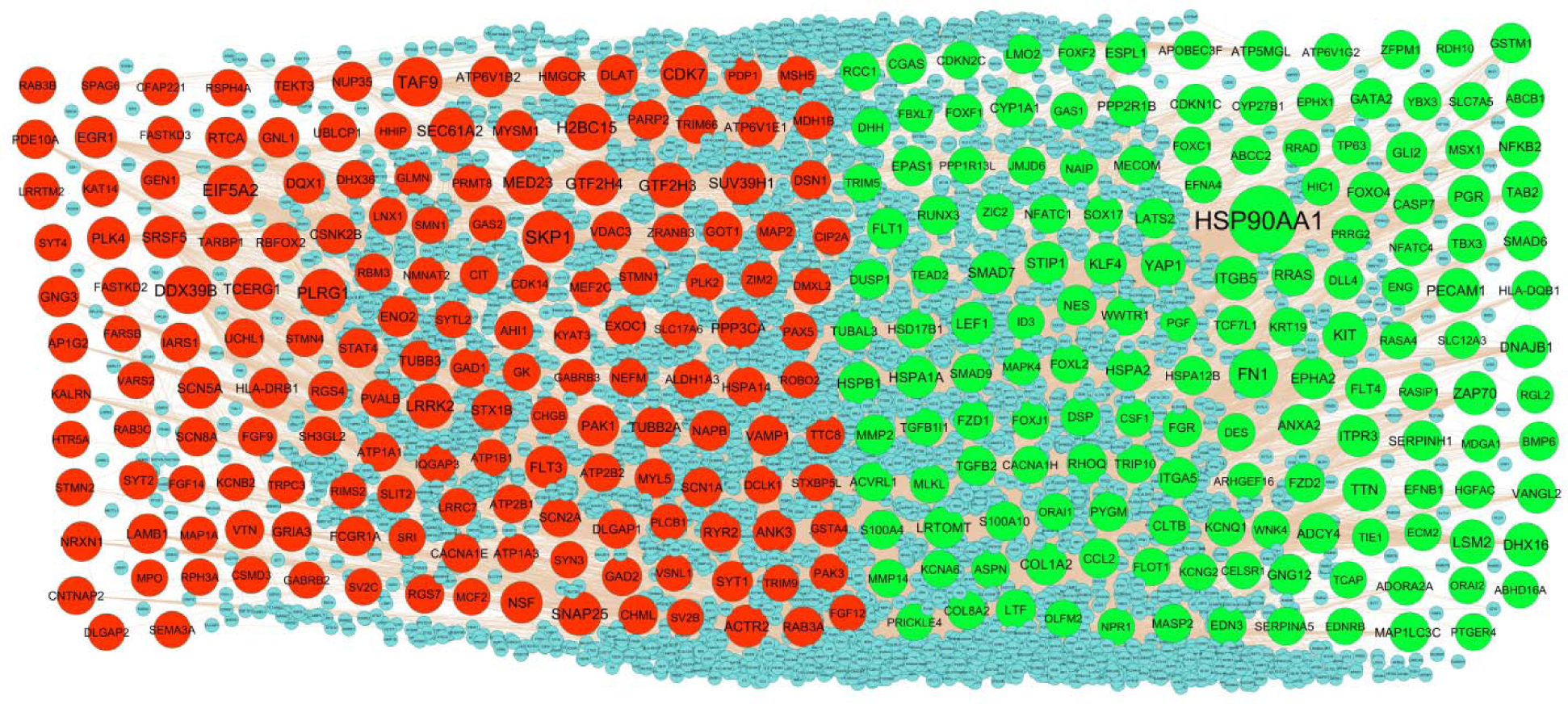
PPI network of DEGs. Up regulated genes are marked in t green; down regulated genes are marked in red.

**Fig. 4.**
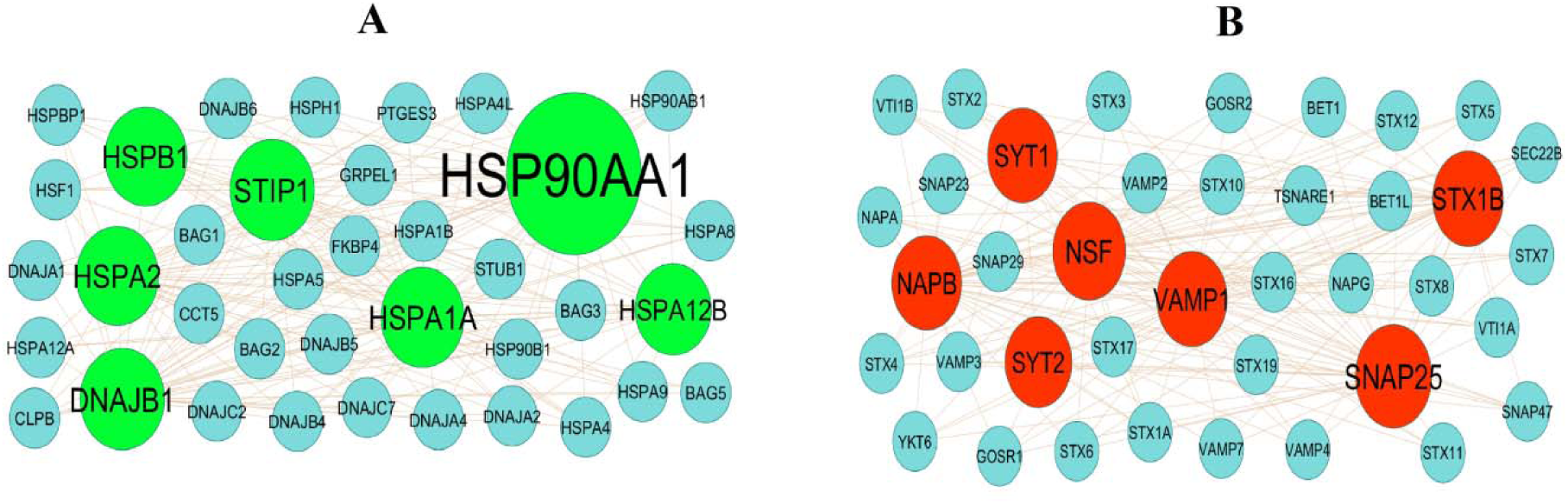
Modules selected from the PPI network. (A) The most significant module was obtained from PPI network with 37 nodes and 179 edges for up regulated genes (B) The most significant module was obtained from PPI network with 37 nodes and 179 edges for down regulated genes. Up regulated genes are marked in parrot green; down regulated genes are marked in red.

**Table 4.**
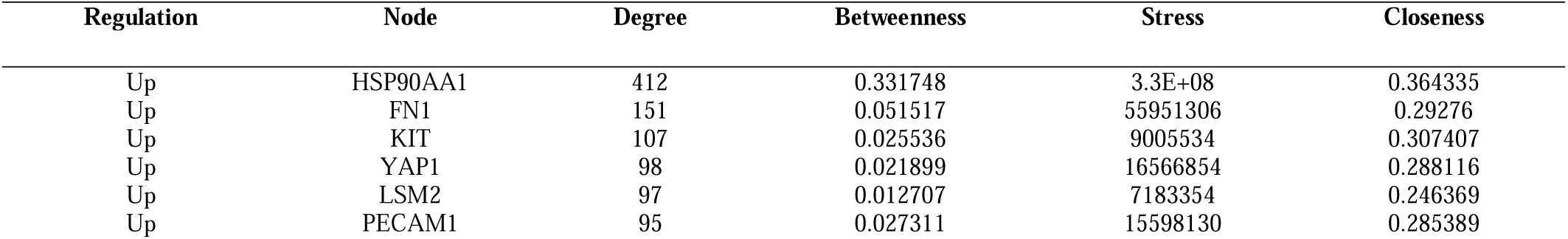
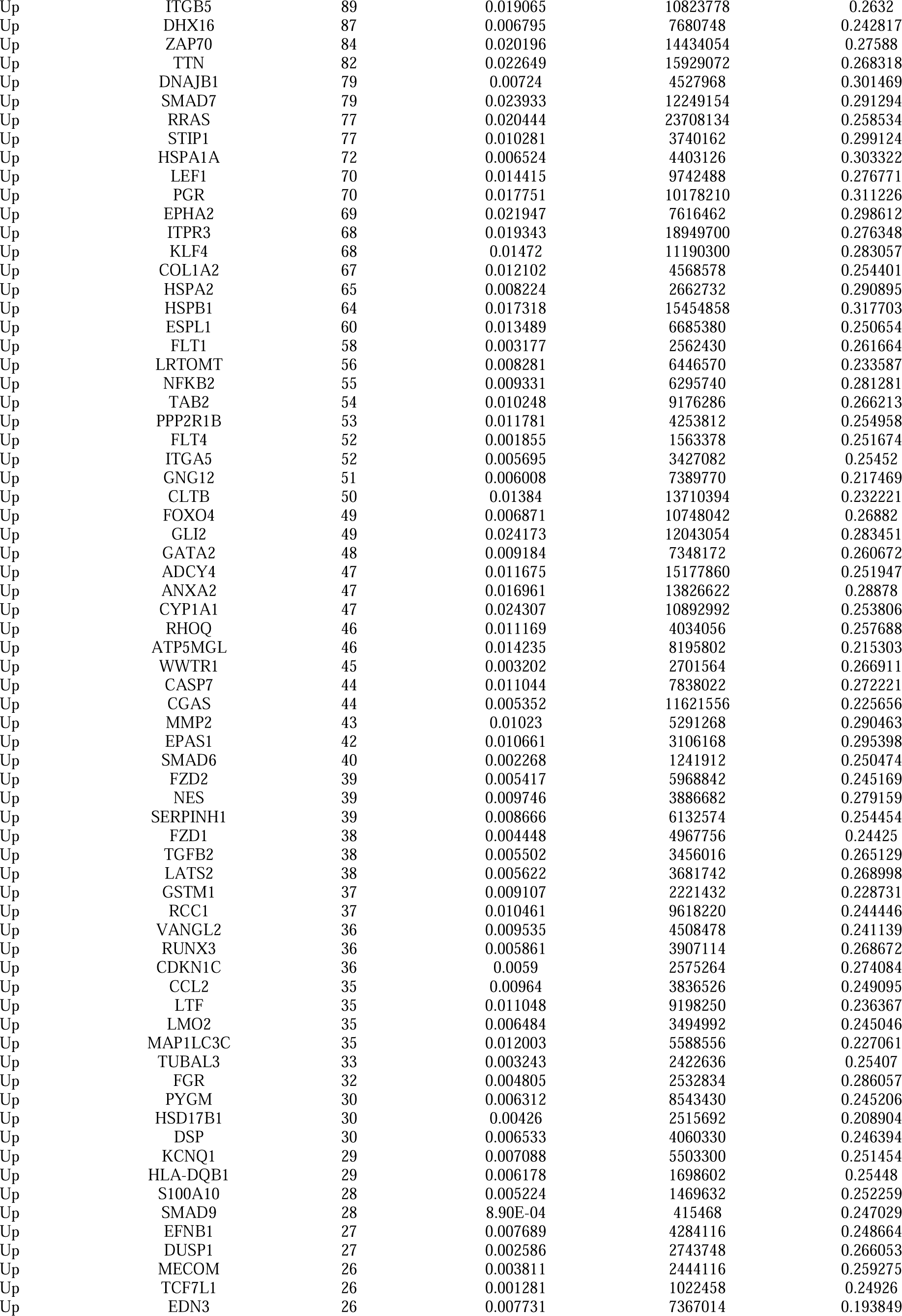
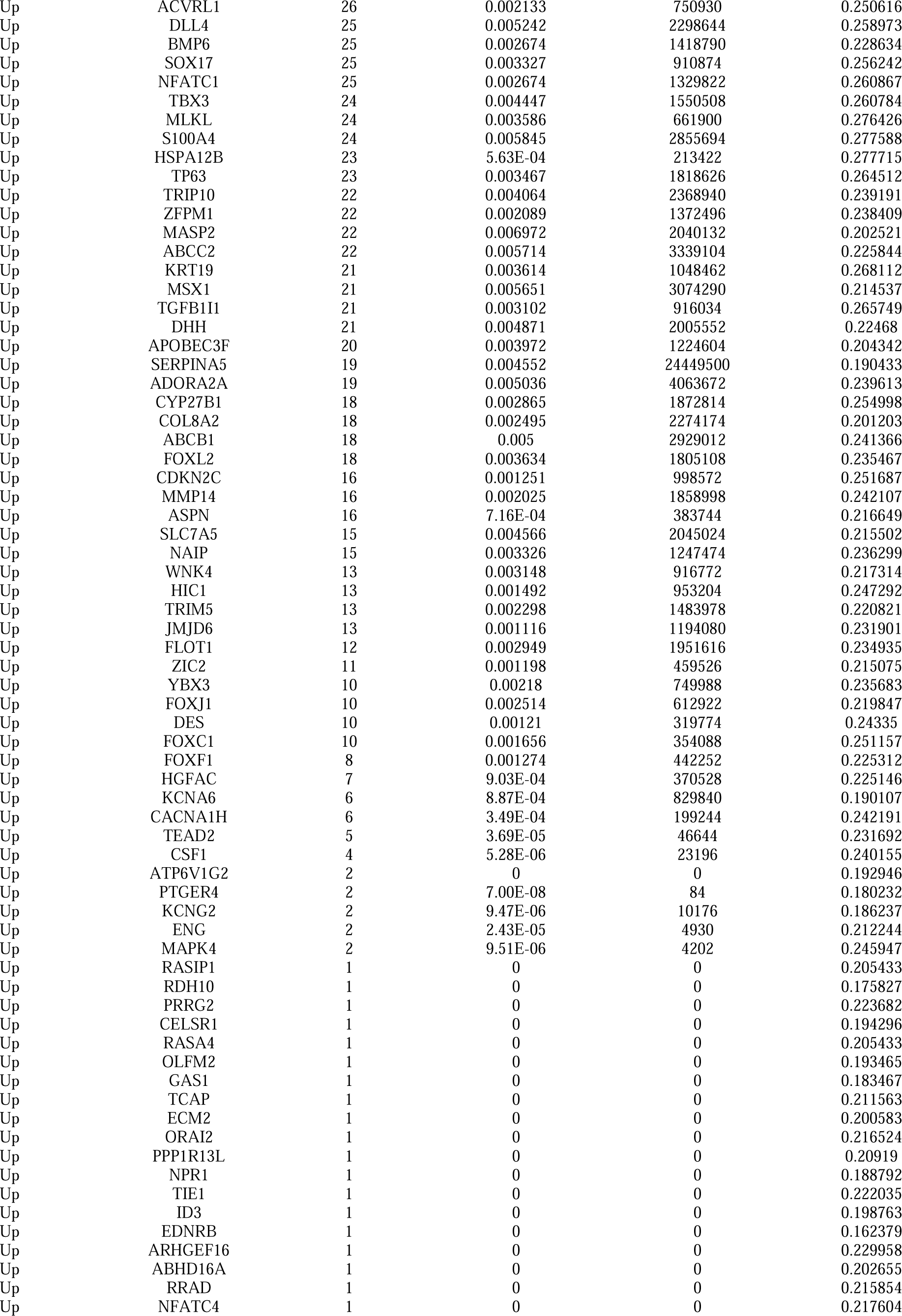
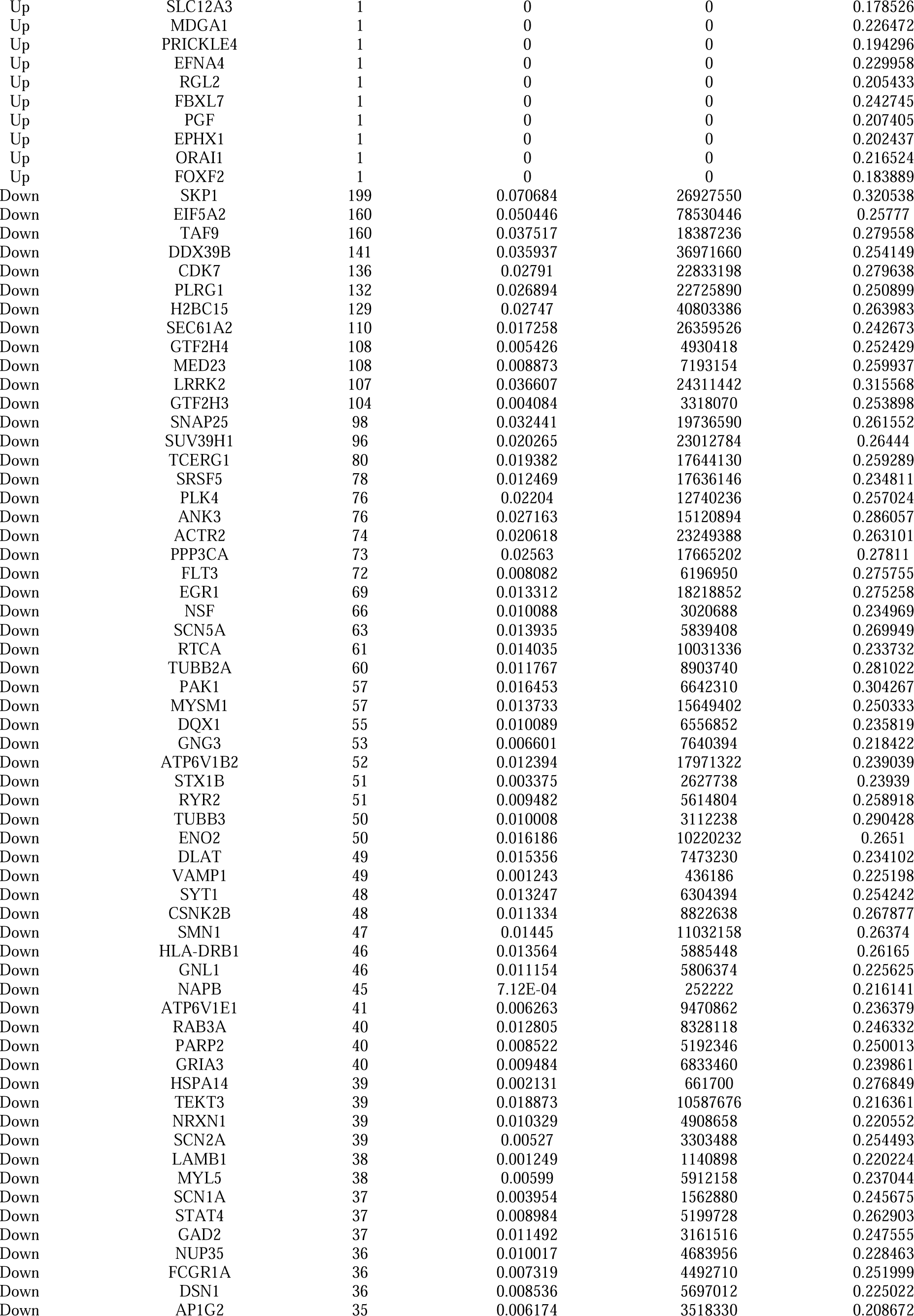
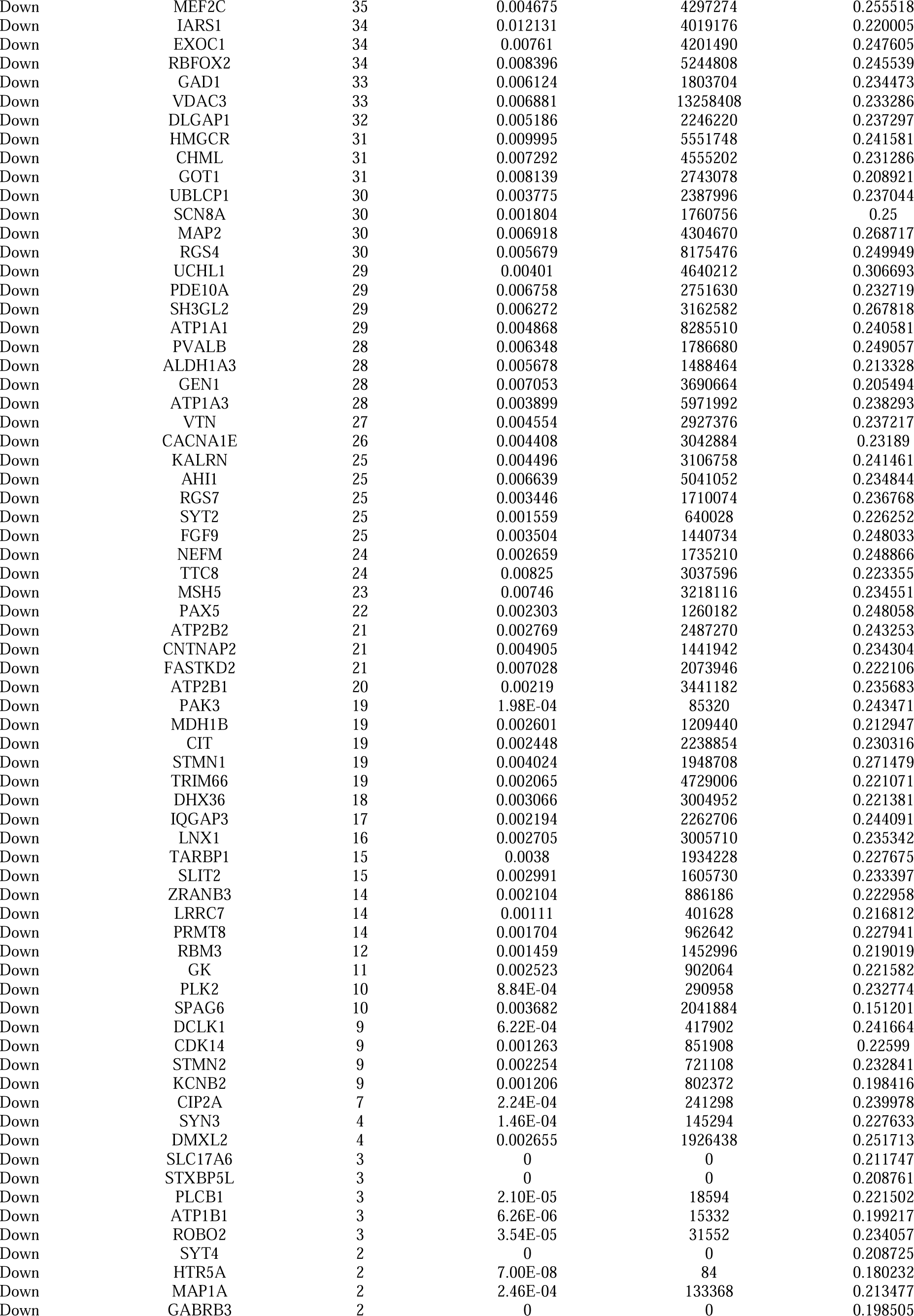
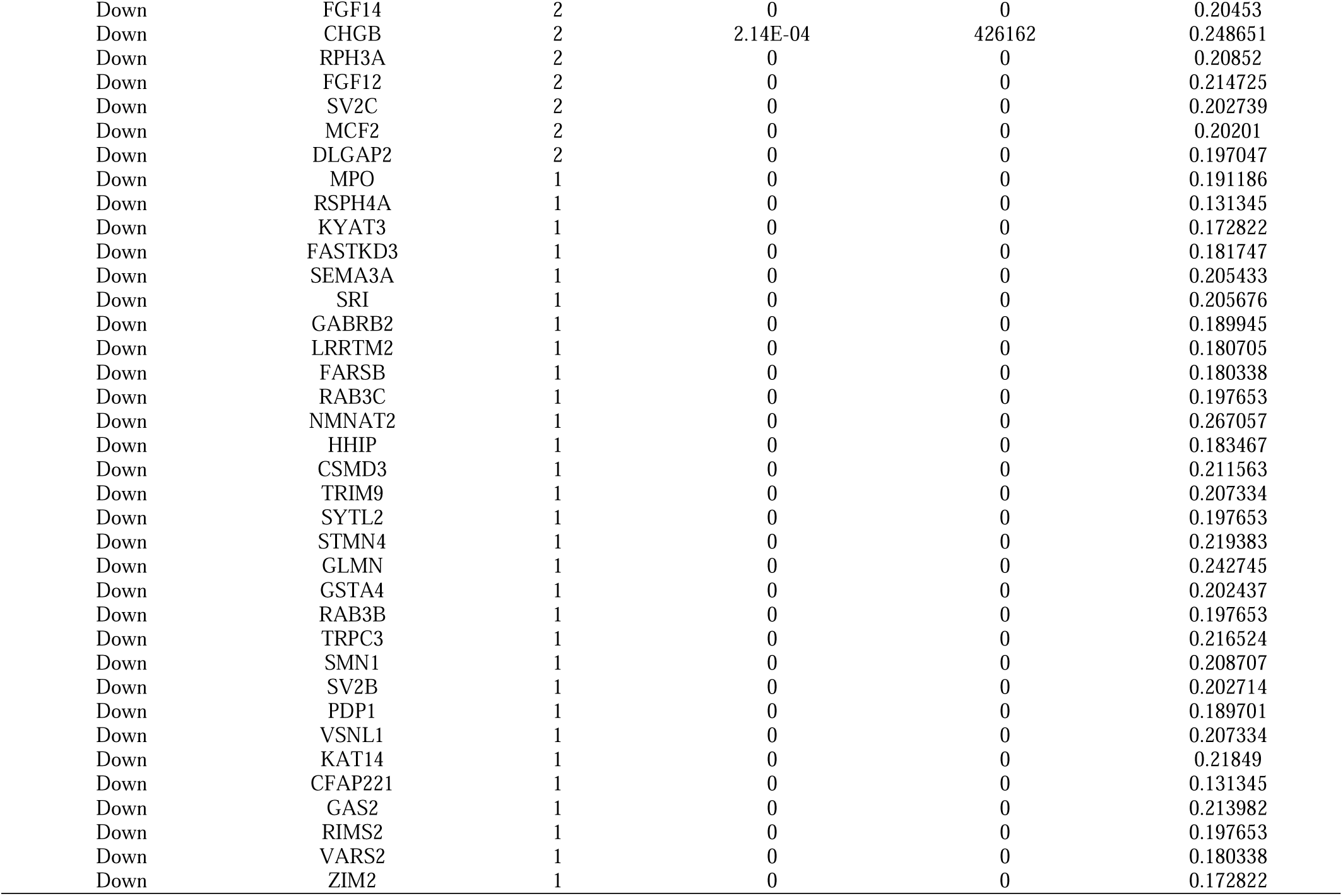
Topology table for up and down regulated genes.

### Construction of the miRNA-hub gene regulatory network

MiRNA-hub gene regulatory network was generated using the miRNet web tool. The network contained 4666 (miRNA: 4344; Hub Gene: 322) nodes and 47545 edges (Fig.5). HSP90AA1 was regulated by 538 miRNAs (ex; hsa-mir-545-3p); FN1 was regulated by 439 miRNAs (ex; hsa-mir-296-3p); YAP1 was regulated by 412 miRNAs (ex; hsa-mir-183-5p); STIP1 was regulated by 309 miRNAs (ex; hsa-miR-23a-5p); TTN was regulated by 297 miRNAs (ex; hsa-miR-3909); DDX39B was regulated by 370 miRNAs (ex; hsa-miR-548f-5p); TCERG1 was regulated by 334 miRNAs (ex; hsa-miR-429); EIF5A2 was regulated by 273 miRNAs (ex; hsa-mir-4521); SKP1 was regulated by 234 miRNAs (ex; hsa-miR-4735-5p); PLRG1 was regulated by 216 miRNAs (ex; hsa-miR-489-3p) (Table 5).

**Fig. 5.**
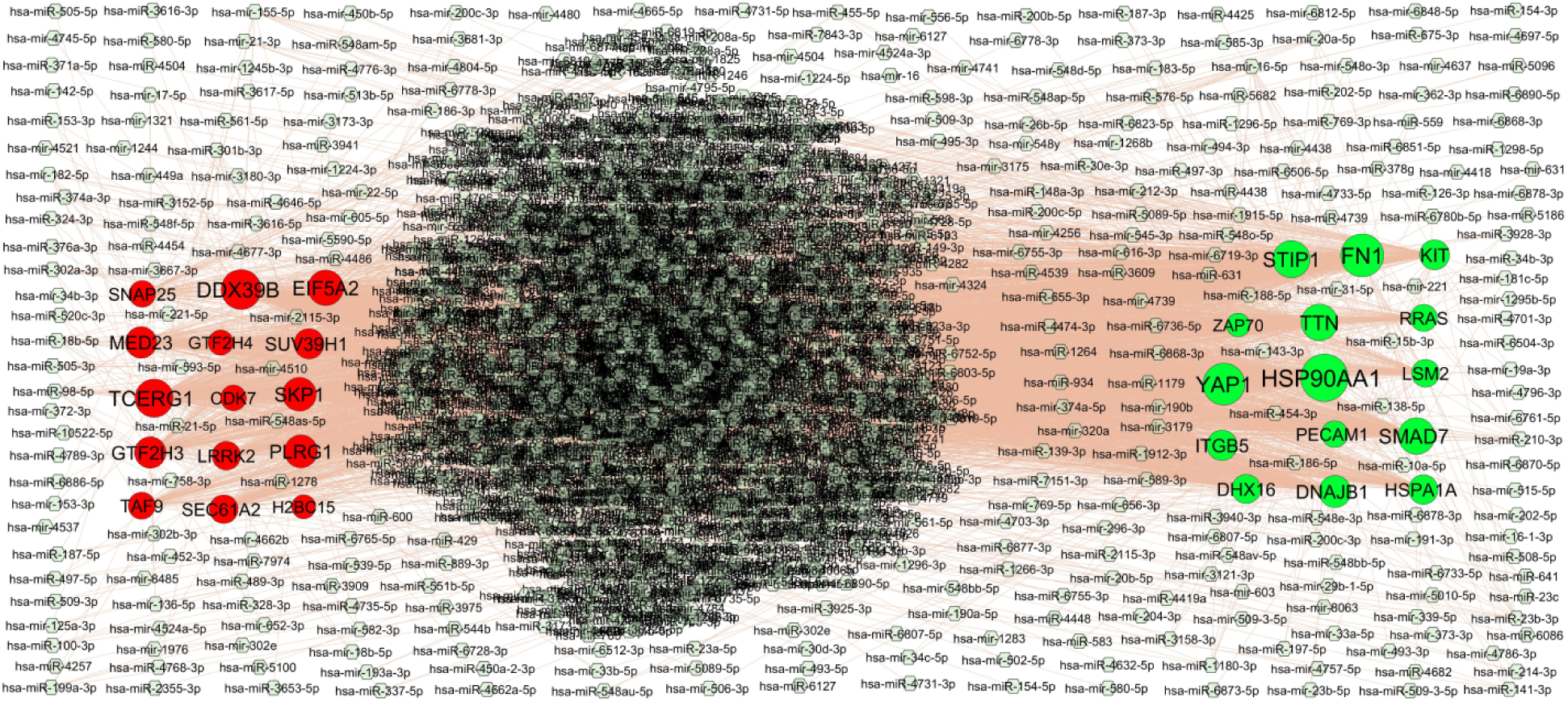
Hub gene - miRNA regulatory network. The light gray color diamond nodes represent the key miRNAs; up regulated genes are marked in green; down regulated genes are marked in red.

**Table 5.** MiRNA - hub gene and TF – hub gene topology table.

| Regulation | Hub Genes | Degree | MicroRNA | Regulation | Hub Genes | Degree | TF |
| --- | --- | --- | --- | --- | --- | --- | --- |
| Up | HSP90AA1 | 538 | hsa-mir-545-3p | Up | LSM2 | 57 | PLAG1 |
| Up | FN1 | 439 | hsa-mir-296-3p | Up | SMAD7 | 14 | NFIC |
| Up | YAP1 | 412 | hsa-mir-183-5p | Up | FN1 | 14 | CREB1 |
| Up | STIP1 | 309 | hsa-miR-23a-5p | Up | ZAP70 | 14 | USF2 |
| Up | TTN | 297 | hsa-miR-3909 | Up | HSP90AA1 | 13 | SREBF1 |
| Up | SMAD7 | 296 | hsa-miR-3616-5p | Up | RRAS | 12 | RELA |
| Up | DNAJB1 | 196 | hsa-mir-1976 | Up | DHX16 | 11 | SRY |
| Up | ITGB5 | 159 | hsa-miR-598-3p | Up | TTN | 11 | PDX1 |
| Up | KIT | 150 | hsa-miR-6736-5p | Up | HSPA1A | 10 | YY1 |
| Up | HSPA1A | 140 | hsa-mir-22-5p | Up | YAP1 | 9 | STAT3 |
| Up | DHX16 | 130 | hsa-miR-769-3p | Up | KIT | 8 | TFAP2A |
| Up | LSM2 | 102 | hsa-mir-4739 | Up | STIP1 | 7 | ARID3A |
| Up | RRAS | 86 | hsa-mir-1245b-3p | Up | DNAJB1 | 7 | HINFP |
| Up | PECAM1 | 77 | hsa-miR-148a-3p | Up | ITGB5 | 3 | ELK1 |
| Up | ZAP70 | 9 | hsa-miR-631 | Up | PECAM1 | 2 | FOXC1 |
| Down | DDX39B | 370 | hsa-miR-548f-5p | Down | DDX39B | 56 | MEF2A |
| Down | TCERG1 | 334 | hsa-miR-429 | Down | EIF5A2 | 10 | MAX |
| Down | EIF5A2 | 273 | hsa-mir-4521 | Down | CDK7 | 9 | CEBPB |
| Down | SKP1 | 234 | hsa-miR-4735-5p | Down | GTF2H4 | 9 | POU2F2 |
| Down | PLRG1 | 216 | hsa-miR-489-3p | Down | SUV39H1 | 9 | NFYA |
| Down | MED23 | 185 | hsa-miR-301b-3p | Down | TCERG1 | 9 | FOXA1 |
| Down | GTF2H3 | 183 | hsa-miR-302e | Down | TAF9 | 8 | NFKB1 |
| Down | SUV39H1 | 154 | hsa-miR-4632-5p | Down | LRRK2 | 8 | TEAD1 |
| Down | LRRK2 | 122 | hsa-miR-582-3p | Down | SNAP25 | 6 | SP1 |
| Down | SEC61A2 | 121 | hsa-miR-1180-3p | Down | PLK4 | 6 | HOXA5 |
| Down | TAF9 | 88 | hsa-miR-30e-3p | Down | GTF2H3 | 6 | ESR1 |
| Down | SNAP25 | 77 | hsa-miR-153-3p | Down | MED23 | 5 | EN1 |
| Down | CDK7 | 57 | hsa-miR-1264 | Down | PLRG1 | 4 | SRF |
| Down | GTF2H4 | 43 | hsa-miR-15b-3p | Down | SKP1 | 4 | FOXL1 |
| Down | H2BC15 | 20 | hsa-miR-21-5p | Down | SEC61A2 | 4 | GATA3 |

### Construction of the TF-hub gene regulatory network

TF-hub gene regulatory network was generated using the NetworkAnalyst web tool. The network contained 411 (TF:96; Hub Gene: 315) nodes and 2812 edges (Fig.6). LSM2 was regulated by 57 TFs (ex; PLAG1); SMAD7 was regulated by 14 TFs (ex; NFIC); FN1 was regulated by 14 TFs (ex; CREB1); ZAP70 was regulated by 14 TFs (ex; USF2); HSP90AA1 was regulated by 13 TFs (ex; SREBF1); DDX39B was regulated by 56 TFs (ex; MEF2A); EIF5A2 was regulated by 10 TFs (ex; MAX); CDK7 was regulated by 9 TFs (ex; CEBPB); GTF2H4 was regulated by 9 TFs (ex; POU2F2); SUV39H1 was regulated by 9 TFs (ex; NFYA) (Table 5).

**Fig. 6.**
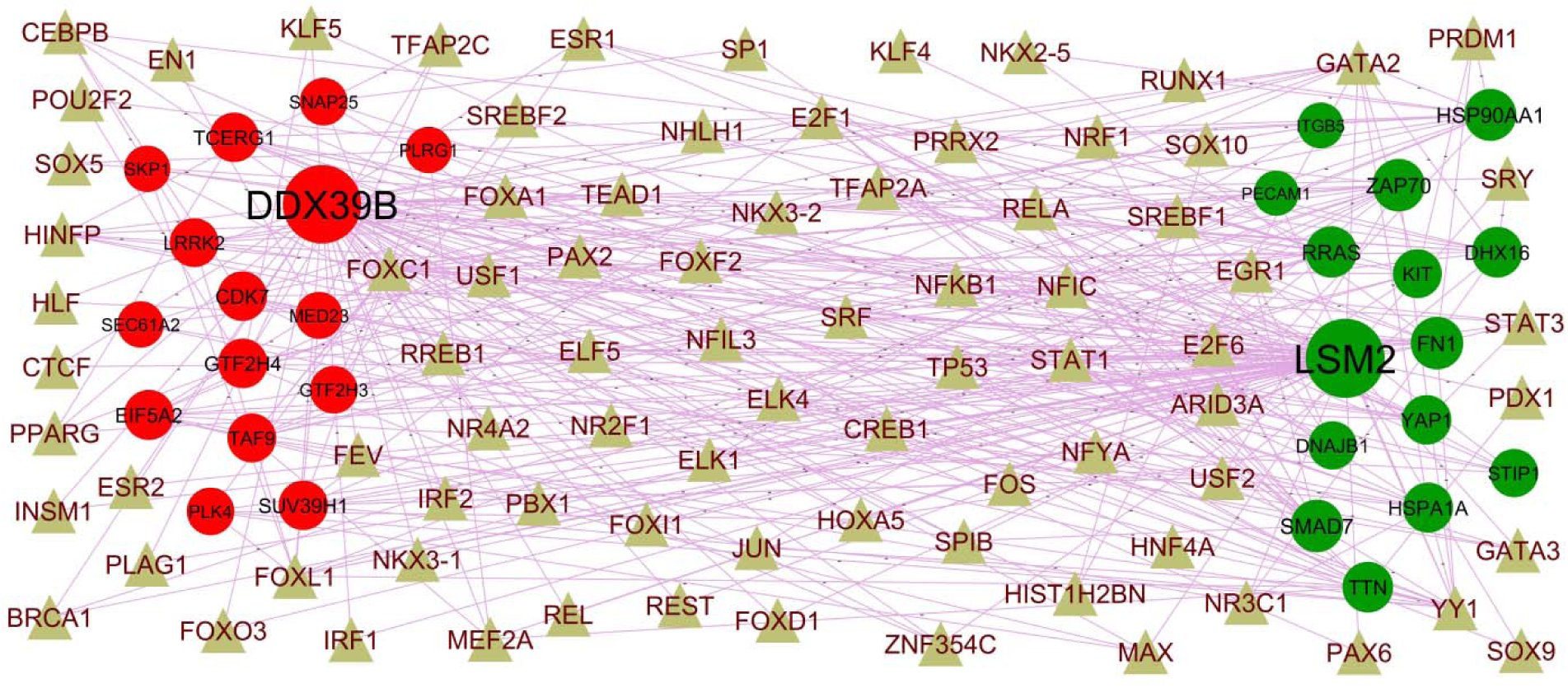
Hub gene - TF regulatory network. The brown color triangle nodes represent the key TFs; up regulated genes are marked in dark green; down regulated genes are marked in dark red.

### Receiver operating characteristic curve (ROC) analysis

Furthermore, we assessed the diagnostic efficacy of individual hub genes. In the AD, HSP90AA1 (AUC = 0.909), FN1 (AUC = 0.926), KIT (AUC = 0.922), YAP1 (AUC = 0.906), LSM2 (AUC = 0.912), SKP1 (AUC = 0.920), EIF5A2 (AUC = 0.923), TAF9 (AUC = 0.914), DDX39B (AUC =0.902) and CDK7 (AUC = 0.918) demonstrated favorable diagnostic efficiency for differentiating patients with AD from normal controls (Fig.7).

**Fig. 7.**
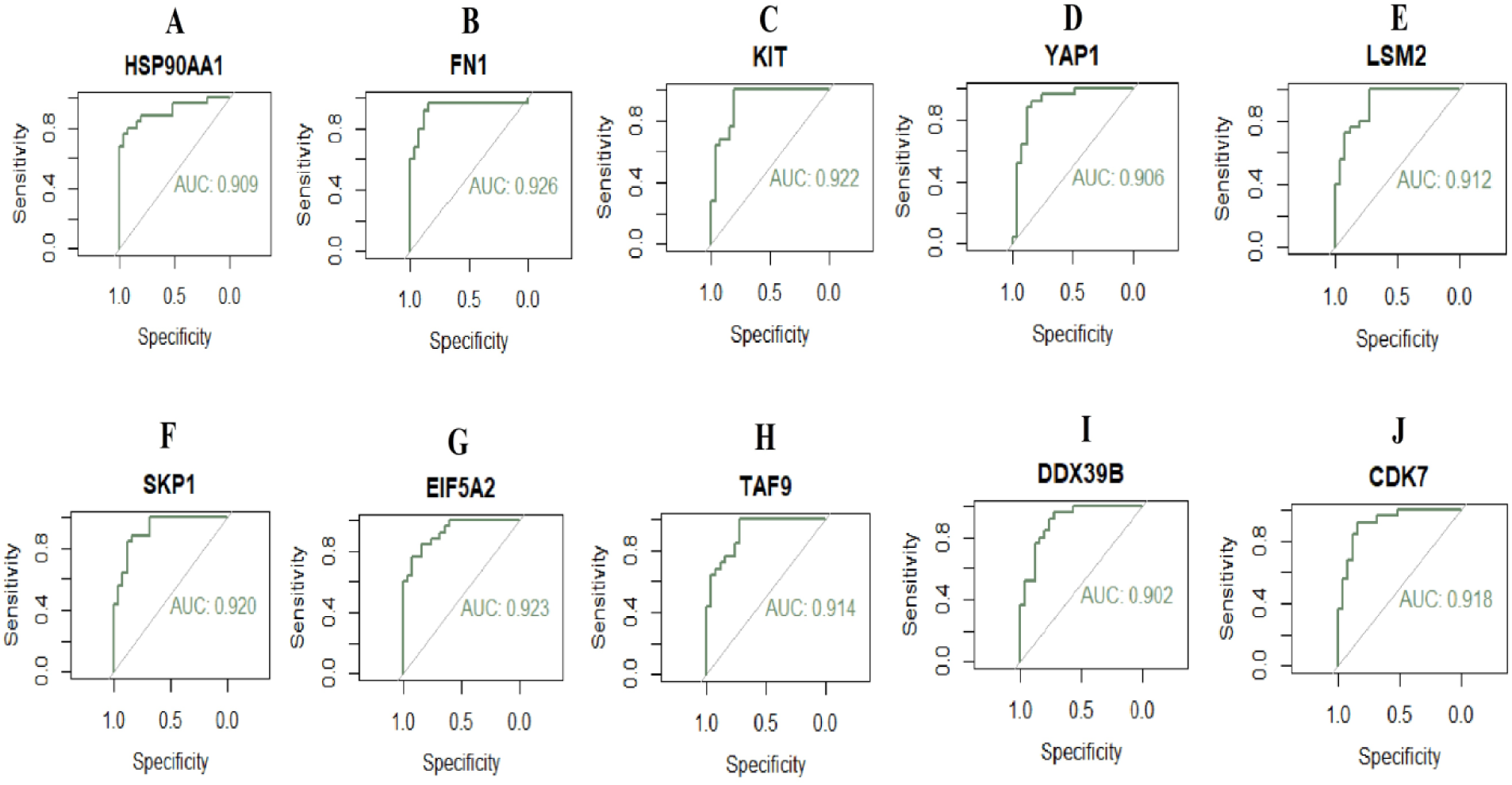
ROC curve analyses of hub genes. A) HSP90AA1 B) FN1 C) KIT D) YAP1 E) LSM2 F) SKP1 G) EIF5A2 H) TAF9 I) DDX39B J) CDK7

### Discussion

AD is a major cause of central nervous system dysfunction in the elderly population and leads to a great public health burden [60]. Thus, outstanding screening NGS techniques and accurate diagnosis remains the great test for lowering the incidence of AD. In the current investigation, integrated bioinformatics and NGS data analysis was used to identify the potential key genes related to AD. By performing DEGs analysis, 479 up regulated and 479 down regulated genes were successfully identified (|log FC| > 0.7764, |log FC| < -0.664 and adjust P-value < .05), respectively. Research has shown that HSPA1A [61], GSTM1 [62], HLA-DRB1 [63] and MPO (myeloperoxidase) [64] plays an important role in the pathogenesis of AD. Altering expressing level of HSPA1A [65], GSTM1 [66], HLA-DRB1 [67] and MPO (myeloperoxidase) [68] can facilitate vascular diseases. A studies have indicated that its altered expression of HSPA1A [69], TGFB2-OT1 [70], GSTM1 [71], G0S2 [72], HCG18 [73], HLA-DRB1 [74], TRIM26 [75] and MPO (myeloperoxidase) [76] are closely associated with the onset and progression of inflammation. HSPA1A [77], GSTM1 [78], G0S2 [79], HCG18 [80], HLA-DRB1 [81] and MPO (myeloperoxidase) [82] are significantly related to the diabetes mellitus. HSPA1A [83], GSTM1 [78], SOX7 [84], G0S2 [85], HCG18 [86], HLA-DRB1 [87] and MPO (myeloperoxidase) [88] might serve as molecular markers for cardiovascular diseases. HSPA1A [89], GSTM1 [90] and MPO (myeloperoxidase) [91] might take part in the occurrence of obesity. HSPA1A [92], GSTM1 [93] and HLA-DRB1 [94] might be involved in the occurrence and development of schizophrenia. HSPA1A [95], GSTM1 [96], HLA-DRB1 [63] and MPO (myeloperoxidase) [97] might play an important role in the pathophysiology of Parkinson’s diseases. KCNA6 [98], GSTM1 [99], SOX7 [100], HLA-DRB1 [101] and MPO (myeloperoxidase) [102] can participate in the occurrence and development of epilepsy. GSTM1 [103], SOX7 [104], HLA-DRB1[105] and MPO (myeloperoxidase) [76] might mediate the process of brain injury. Previous studies have shown that GSTM1 [106], HCG18 [107], HLA-DRB1 [108], TRIM26 [75] and MPO (myeloperoxidase) [109] might promote infections. GSTM1 [110], HLA-DRB1 [111] and MPO (myeloperoxidase) [112] expression is altered in the patients with hypertension. GSTM1 [113], HLA-DRB1 [114] and MPO (myeloperoxidase) [115] have been proposed as novel biomarkers for multiple sclerosis progression. Regulation of GSTM1 [116], HLA-DRB1 [117] and MPO (myeloperoxidase) [118] levels might be a novel treatment option against autism spectrum disorder. GSTM1 [119], HLA-DRB1 [120] and MPO (myeloperoxidase) [121] participates in the occurrence and development of stroke. Excessive activation of GSTM1 [122], HLA-DRB1 [123] and MPO (myeloperoxidase) [124] promotes the development of bipolar disorder. Research has reported that altered MPO (myeloperoxidase) [125] expression in the amyotrophic lateral sclerosis. Studies have shown that MPO (myeloperoxidase) [126] plays a certain role in depression. This analysis led to the identification of DEGs as key biomarkers that could be of mechanistic relevance for AD pathogenesis and progression.

In this investigation, the GO and REACTOME pathway enrichment analyses of the key DEGs in AD were analyzed by using g:Profiler online tool. These analyses could help to find some key factors involved in the regulation of AD. For example, signal transduction [127], muscle contraction [128], signaling by receptor tyrosine kinases [129], hemostasis [130], signaling by WNT [131], extracellular matrix organization [132], cardiac conduction [133], neuronal system [134], transmission across chemical synapses [135], ion channel transport [136], nervous system development [137] and infectious disease [138] are involved in the regulation of the AD and might serve as signaling pathways of AD. The expression of HSP90AA1 [139], FN1 [140], YAP1 [141], PECAM1 [142], EPHA2 [143], KLF4 [144], HSPA2 [145], FLT1 [146], ITGA5 [147], ANXA2 [148], CASP7 [149], CGAS (cyclic GMP-AMP synthase) [150], MMP2 [151], TGFB2 [152], GSTM1 [153], CCL2 [154], LTF (lactotransferrin) [155], HSD17B1 [156], HLA-DQB1 [157], S100A10 [158], EFNB1 [159], ACVRL1 [160], BMP6 [161], MLKL (mixed lineage kinase domain like pseudokinase) [162], ADORA2A [163], ABCB1 [164], MMP14 [165], JMJD6 [166], CSF1 [167], PGF (placental growth factor) [168], PYGM (glycogen phosphorylase, muscle associated) [169], SERPINA5 [170], MDGA1 [171], FBXL7 [172], VGF (VGF nerve growth factor inducible) [173], GAP43 [174], THY1 [175], STMN2 [176], CNTN5 [177], NEUROD6 [178], RGS4 [179], RXFP1 [180], SYT1 [181], SNAP25 [182], VTN (vitronectin) [183], FGF14 [184], MEF2C [185], NRN1 [186], GABRG2 [187], HCN1 [188], PCP4 [189], RPH3A [190], AHI1 [191], CCKBR (cholecystokinin B receptor) [192], CNTNAP2 [193], SRI (sorcin) [194], RORB (RAR related orphan receptor B) [195], PAK3 [196], SEMA3A [197], UCHL1 [198], ELAVL4 [199], EGR1 [200], CHRM1 [201], MAP2 [202], LRRK2 [203], MAP1A [204], RYR2 [205], PLK2 [206], C9ORF72 [207], VAMP1 [208], SST (somatostatin) [209], PVALB (parvalbumin) [210], TAC1 [211], SPARCL1 [212], DLGAP2 [213], CAP2 [214], SV2B [215], LAMP5 [216], RAB3A [217], VSNL1 [218], SCG5 [219], DDX39B [220], CDK7 [221], HMGCR (3-hydroxy-3-methylglutaryl-CoA reductase) [222], ABCA5 [223], TRPC3 [224] and FABP3 [225] were altered in the AD. HSP90AA1 [226], FN1 [227], YAP1 [228], PECAM1 [229], TTN (titin) [230], SMAD7 [231], LEF1 [232], PGR (progesterone receptor) [233], EPHA2 [143], KLF4 [144], HSPB1 [234], FLT1 [235], NFKB2 [236], TAB2 [237], ITGA5 [238], GNG12 [239], FOXO4 [240], GLI2 [241], GATA2 [242], ANXA2[243], CYP1A1 [244], WWTR1 [245], CASP7 [246], CGAS (cyclic GMP-AMP synthase) [247], MMP2 [248], EPAS1 [249], SMAD6 [250], TGFB2 [251], LATS2 [252], GSTM1 [253], VANGL2 [254], RUNX3 [255], CCL2 [256], LTF (lactotransferrin) [257], HSD17B1 [258], HLA-DQB1 [259], EFNB1 [260], DUSP1 [261], DLL4 [262], BMP6 [263], SOX17 [264], NFATC1 [265]. MLKL (mixed lineage kinase domain like pseudokinase) [266], S100A4 [267], TP63 [268], MASP2 [269], CYP27B1 [270], ABCB1 [271], MMP14 [272], SLC7A5 [273], NAIP (NLR family apoptosis inhibitory protein) [274], WNK4 [275], JMJD6 [276], FOXJ1 [277], FOXC1 [278], FOXF1 [279], CSF1 [280], PTGER4 [281], ENG (endoglin) [282], NPR1 [283], TIE1 [284], EDNRB (endothelin receptor type B) [285], ORAI1 [286], PPP1R13L [287], FBXL7 [288], HSPA12B [289], VGF (VGF nerve growth factor inducible) [290], GAP43 [291], MAS1 [292], THY1 [293], THY1 [294], NELL1 [295], RGS4 [296], CDKL5 [297], FGF9 [298], SNAP25 [299], ATP2B1 [300], TAGLN3 [301], VTN (vitronectin) [302], MEF2C [303], GABRG2 [304], HCN1 [305], PARP2 [306], TRIM45 [307], PAK1 [308], HAPLN1 [309], ATAT1 [310], CSMD1 [311], DCLK1 [312], HHIP (hedgehog interacting protein) [313], SLIT2 [314], AGBL4 [315], SEMA3A [316], UCHL1 [317], EGR1 [318], PAX5 [319], CITED2 [320], PTPRO (protein tyrosine phosphatase receptor type O) [321], LRRK2 [322], NRXN1 [323], TRIM67 [324], MYSM1 [325], CLASP2 [326], PLK2 [327], C9ORF72 [328], SST (somatostatin) [329], SPARCL1 [330], THEMIS (thymocyte selection associated) [331], ATP1A3 [332], KCNH1 [333], CRHR2 [334], TRIM9 [335], BTBD8 [336], LAMP5 [337], NUAK1 [338], FLT3 [339], SUV39H1 [340], PDE10A [341], PRMT8 [342], NCALD (neurocalcin delta) [343], CDK7 [344], HMGCR (3-hydroxy-3-methylglutaryl-CoA reductase) [345], PDP1 [346], NQO2 [347], CISD1 [348], LANCL2 [349], TRPC3 [350] and FABP3 [351] are found to be associated with inflammation. Genes include HSP90AA1 [352], FN1 [353], YAP1 [354], PECAM1 [355], TTN (titin) [230], SMAD7 [356], LEF1 [357], EPHA2 [358], KLF4 [359], COL1A2 [360], HSPB1 [361], FLT1 [362], TAB2 [363], FLT4 [364], GATA2 [365], ANXA2 [366], CYP1A1 [367], CGAS (cyclic GMP-AMP synthase) [368], MMP2 [369], EPAS1 [370], SMAD6 [371], SERPINH1 [372], LATS2 [252], GSTM1 [373], RUNX3 [374], CDKN1C [375], CCL2 [376], LTF (lactotransferrin) [377], DSP (desmoplakin) [378], KCNQ1 [379], HLA-DQB1 [259], DUSP1 [261], DLL4 [380], BMP6 [381], SOX17 [382], MLKL (mixed lineage kinase domain like pseudokinase) [383], TP63 [384], MASP2 [385], MSX1 [386]; ADORA2A [387], CYP27B1 [388], ABCB1 [389], MMP14 [390], ASPN (asporin) [391], JMJD6 [392], FLOT1 [393], FOXJ1 [394], FOXC1 [395], FOXF1 [396], CACNA1H [397], CSF1 [398], ENG (endoglin) [399], CELSR1 [400], TCAP (titin-cap) [401], NPR1 [402], TIE1 [403], RGL2 [404], PGF (placental growth factor) [405], EPHX1 [406], ORAI1 [407], NES (nestin) [408], DES (desmin) [409], RDH10 [410], PPP1R13L [411], HSPA12B [412], PCSK1 [413], GAP43 [414], MAS1 [415], THY1 [416], SCN8A [417], RGS4 [418], RXFP1 [419], CDKL5 [420], SVEP1 [421], NAP1L2 [422], BEX1 [423], FGF9 [424], ATP2B1 [425], FLRT3 [426], FGF12 [427], MEF2C [428], PARP2 [429], PAK1 [430], HAPLN1 [431], SCN5A [432], DCLK1 [433], RBFOX2 [434], SLIT2 [435], PAK3 [436], SEMA3A [316], UCHL1 [437], ADAM23 [438], EGR1 [439], PAX5 [440], CITED2 [441], MYBPHL (myosin binding protein H like) [442], LRRK2 [443], PLCB1 [444], RYR2 [445], PLK2 [446], SPARCL1 [447], ATP1A3 [448], KCNH1 [449], NSF (N-ethylmaleimide sensitive factor, vesicle fusing ATPase) [450], CAP2 [451], HTR2C [452], VDAC3 [453], VARS2 [454], FLT3 [455], CARNS1 [456], SUV39H1 [457], PDE10A [458], PKD2L1 [459], DIRAS3 [460], RSAD2 [461], ANKK1 [462], HMGCR (3-hydroxy-3-methylglutaryl-CoA reductase) [463], CISD1 [348], ABCA5 [464], TRPC3 [465], FABP3 [466] and PDE1A [467] are associated with the risk of cardiovascular diseases. Studies have shown that HSP90AA1 [468], YAP1 [469], DNAJB1 [470], PGR (progesterone receptor) [471], EPHA2 [472], KLF4 [473], HSPB1 [474], FLT1 [475], TAB2 [476], GLI2 [477], GATA2 [478], ANXA2 [479], CASP7 [480], CGAS (cyclic GMP-AMP synthase) [481], MMP2 [482], SMAD6 [483], FZD2 [484], GSTM1 [485], CCL2 [486], LTF (lactotransferrin) [487], DUSP1 [488], BMP6 [489], SOX17 [490], NFATC1 [491], MLKL (mixed lineage kinase domain like pseudokinase) [492], S100A4 [493], MASP2 [494], ADORA2A [495], ABCB1 [496], FOXJ1 [497], FOXC1 [498], CSF1 [499], CELSR1 [500], EDNRB (endothelin receptor type B) [501], NES (nestin) [502], MDGA1 [503], HSPA12B [504], VGF (VGF nerve growth factor inducible) [290], GAP43 [505], THY1 [506], SCG2 [507], NEGR1 [508], SYT1 [509], SNAP25 [510], ATP2B1 [511], MEF2C [512], PARP2 [513], TRIM45 [307], PAK1 [514], NMNAT2 [515], SLIT2 [516], SEMA3A [517], UCHL1 [518], ELAVL4 [519], EGR1 [520], CHRM1 [521], MAP2 [522], LRRK2 [523], CLSTN3 [524], TRIM67 [525], C9ORF72 [526], VAMP1 [527], SST (somatostatin) [528], PVALB (parvalbumin) [529], SPARCL1 [530], NSF (N-ethylmaleimide sensitive factor, vesicle fusing ATPase) [531], SV2B [532], HTR2C [533], RPH3A [534], SUV39H1 [535], PDE10A [536], PRMT8 [342], ENO2 [537], GOT1 [538], ANKK1 [539], TRPC3 [540] and FABP3 [541] participate in regulating brain injury. HSP90AA1 [468], PECAM1 [542], TTN (titin) [543], STIP1 [544], KLF4 [545], COL1A2 [546], TAB2 [547], GATA2 [548], ANXA2 [549], CYP1A1 [550], CASP7 [551], CGAS (cyclic GMP-AMP synthase) [552], MMP2 [553], GSTM1 [554], CCL2 [555], LTF (lactotransferrin) [556], KCNQ1 [557], HLA-DQB1 [558], DUSP1 [559], DLL4 [560], ABCB1 [561], MMP14 [562], FOXJ1 [563], ENG (endoglin) [564], CELSR1 [565], EDNRB (endothelin receptor type B) [566], EPHX1 [567], ORAI1 [568], NES (nestin) [569], HSPA12B [570], VGF (VGF nerve growth factor inducible) [571], GAP43 [572], THY1 [573], FGF9 [574], NAP25 [510], VTN (vitronectin) [575], GABRG2 [576], PAK1 [577], NMNAT2 [578], SEMA3A [579], UCHL1 [580], SPAG6 [581], EGR1 [520], MAP2 [582], CITED2 [583], LRRK2 [584], SNCB (synuclein beta) [585], TRIM9 [335], GAD2 [586], SUV39H1 [587], PDE10A [588] and ENO2 [537] expression level is significantly altered in the stroke. HSP90AA1 [589], YAP1 [590], PECAM1 [591], SMAD7 [592], PGR (progesterone receptor) [593], EPHA2 [594], KLF4 [595], COL1A2 [596], FLT1 [597], NFKB2 [598], GATA2 [599], ANXA2 [600], CYP1A1 [601], CGAS (cyclic GMP-AMP synthase) [602], MMP2 [603], EPAS1 [604], VANGL2 [605], CCL2 [606], LTF (lactotransferrin) [607], KCNQ1 [608], HLA-DQB1 [609], SMAD9 [610], EFNB1 [611], DUSP1 [612], ACVRL1 [613], DLL4 [614], SOX17 [615], NFATC1 [616], S100A4 [617], CYP27B1 [388], ABCB1 [618], ASPN (asporin) [619], WNK4 [620], FOXC1 [621], FOXF1 [622], CACNA1H [623], ENG (endoglin) [624], TCAP (titin-cap) [625], NPR1 [626], SLC12A3 [627], ORAI1 [628], NES (nestin) [408], PCSK1 [629], VGF (VGF nerve growth factor inducible) [630], MAS1 [631], SNAP25 [632], ATP2B1 [633], FGF12 [634], MEF2C [635], CSMD1 [636], CCKBR (cholecystokinin B receptor) [637], GABRB3 [638], TTC21B [639], SEMA3A [640], UCHL1 [641], EGR1 [642], ALDH1A3 [643], CHN1 [644], RYR2 [645], PHF14 [646], SST (somatostatin) [647], SPARCL1 [648], NSF (N-ethylmaleimide sensitive factor, vesicle fusing ATPase) [649], CRHR2 [650], ATP1B1 [651], ATP1A1 [652], DLGAP1 [653], VARS2 [454], PDE10A [654] and TRPC3 [655] have been revealed to be regulated in hypertension. HSP90AA1 [656], YAP1 [657], COL1A2 [658], GATA2 [659], ANXA2 [660], CGAS (cyclic GMP-AMP synthase) [661], MMP2 [662], CCL2 [663], LTF (lactotransferrin) [664], S100A10 [665], DUSP1 [666], ADORA2A [667], ABCB1 [668], FLOT1 [669], CSF1 [670], NES (nestin) [671], MDGA1 [672], VGF (VGF nerve growth factor inducible) [673], GAP43 [674], THY1 [675], RGS4 [676], RXFP1 [180], CDKL5 [677], NEGR1 [678], SYT1 [679], SNAP25 [680], NRN1 [681], HCN1 [682], PAK1 [683], AHI1 [684], CNTNAP2 [685], SCN1A [686], SLIT2 [687], PAK3 [688], SEMA3A [689], UCHL1 [690], EGR1 [691], MAP2 [692], HTR5A [693], LRRK2 [694], NRXN1 [695], TRIM67 [696], MYSM1 [697], C9ORF72 [698], SST (somatostatin) [699], PVALB (parvalbumin) [700], TAC1 [701], SV2C [702], ATP1A3 [703], NSF (N-ethylmaleimide sensitive factor, vesicle fusing ATPase) [704], CRHR2 [705], HTR2C [452], RAB3B [706], ATP6V1B2 [707], HTR3B [708], DMXL2 [709], CACNA1E [710], TRANK1 [711], SUV39H1 [712], ANKK1 [713], NCALD (neurocalcin delta) [343] and TRPC3 [714] have been reported to be altered expression in depression. Studies have found that the FN1 [227], YAP1 [715], PECAM1 [716], ZAP70 [717], SMAD7 [718], STIP1 [719], EPHA2 [720], KLF4 [721], NFKB2 [722], GNG12 [723], GATA2 [724], ANXA2 [725], CYP1A1 [726], CASP7 [727], CGAS (cyclic GMP-AMP synthase) [728], MMP2 [729], SMAD6 [730], LATS2 [731], GSTM1 [732], RUNX3 [733], CCL2 [734], LTF (lactotransferrin) [735], HLA-DQB1 [736], DUSP1 [737], DLL4 [738], BMP6 [739], NFATC1 [740], S100A4 [741], ABCC2 [742], APOBEC3F [743], ADORA2A [744], CYP27B1 [745], ABCB1 [746], NAIP (NLR family apoptosis inhibitory protein) [747], WNK4 [748], TRIM5 [749], YBX3 [750], FOXJ1 [751], CSF1 [752], GAS1 [753], NPR1 [754], ID3 [755], PGF (placental growth factor) [756], ORAI1 [757], GAP43 [758], THY1 [759], OPN4 [760], NELL1 [761], FGF9 [762], SNAP25 [763], ATP2B1 [764], FLRT3 [765], MEF2C [766], PAK1 [767], SLIT2 [768], UCHL1 [769], EGR1 [318], PAX5 [770], LRRK2 [771], C9ORF72 [772], ATP1A3 [773], ATP1A1 [774], FLT3 [775], SUV39H1 [776], SPTLC2 [777], DDX39B [778], ENO2 [779], RSAD2 [780], CDK7 [781], HMGCR (3-hydroxy-3-methylglutaryl-CoA reductase) [782] and LANCL2 [783] plays a vital role in the development of infection. The expression levels of KIT (KIT proto-oncogene, receptor tyrosine kinase) [784], PECAM1 [785], GSTM1 [786], ADORA2A [787], CACNA1H [788], LAMB1 [789], NEGR1 [790], SNAP25 [791], EXT1 [792], FGF12 [793], MEF2C [794], CSMD3 [795], AHI1 [796], SCN2A [797], CNTNAP2 [798], GABRB3 [799], SCN1A [800], SEMA3A [801], UCHL1 [802], ALDH1A3 [803], PAX5 [804], MAP2 [805], ANK3 [806], LRRK2 [807], NRXN1 [808], ATP2B2 [809], RYR2 [810], SST (somatostatin) [811], PVALB (parvalbumin) [812], TAC1 [813], SPARCL1 [814], DLGAP2 [815], ATP1A3 [816], SNCB (synuclein beta) [817], NSF (N-ethylmaleimide sensitive factor, vesicle fusing ATPase) [818], GAD1 [819], CRHR2 [820], HTR2C [821], SYN3 [822], NAPB (NSF attachment protein beta) [823], DMXL2 [709], ATP1A1 [824], RPH3A [825], UBLCP1 [826]. ENO2 [827], SLC25A12 [828], ZNF711 [829], HMGCR (3-hydroxy-3-methylglutaryl-CoA reductase) [830] and FABP3 [831] have been proved to be elevated in autism spectrum disorder. Some researchers have reported that altered genes include KIT (KIT proto-oncogene, receptor tyrosine kinase) [832], TTN (titin) [833], HSPB1 [834], CYP1A1 [835], MMP2 [836], FZD2 [837], CCL2 [838], S100A4 [839], CYP27B1 [840], ABCB1 [841], CACNA1H [842], CSF1 [843], ENG (endoglin) [844], VGF (VGF nerve growth factor inducible) [845], GAP43 [846], STMN2 [847], MEF2C [848], SEMA3A [849], UCHL1 [850], ELAVL4 [851], LRRK2 [852], C9ORF72 [207], KIFAP3 [853], SPTLC2 [854] and GOT1 [855] expression in the amyotrophic lateral sclerosis. Transcription of YAP1 [856], PGR (progesterone receptor) [857], KLF4 [858], COL1A2 [859], HSPA2 [860], HSPB1 [861], FLT1 [862], TAB2 [863], PPP2R1B [864], FLT4 [865], GATA2 [866], ANXA2 [867], CYP1A1 [868], CGAS (cyclic GMP-AMP synthase) [869], MMP2 [870], SMAD6 [371], GSTM1 [871], CDKN1C [872], CCL2 [873], LTF (lactotransferrin) [874], KCNQ1 [875], DUSP1 [876], S100A4 [877], CYP27B1 [878], ABCB1 [879], MMP14 [880], WNK4 [881], EDNRB (endothelin receptor type B) [882], PGF (placental growth factor) [883], CDKN2C [884], PCSK1 [885], VGF (VGF nerve growth factor inducible) [886], GAP43 [887], THY1 [888], RGS4 [889], FGF9 [890], NEGR1 [891], SNAP25 [892], EXT1 [893], AHI1 [894], SCN5A [895], DCLK1 [433], CNTNAP2 [896], HHIP (hedgehog interacting protein) [897], SLIT2 [898], PAK3 [899], EGR1 [900], CITED2 [901], PTPRO (protein tyrosine phosphatase receptor type O) [902], LRRK2 [903], CLSTN3 [904], TRIM67 [324], SST (somatostatin) [905], GAD2 [906], FLT3 [907], SUV39H1 [908], PDE10A [909], ANKK1 [910], NCALD (neurocalcin delta) [911], HMGCR (3-hydroxy-3- methylglutaryl-CoA reductase) [830], CISD1 [912] and FABP3 [913] were significantly altered in patients with obesity. A previous study reported that the YAP1 [914], PECAM1 [915], SMAD7 [916], KLF4 [917], FLT1 [918], ANXA2 [919], MMP2 [920], EPAS1 [921], TGFB2 [922], GSTM1 [923], CCL2 [924], LTF (lactotransferrin) [925], HLA-DQB1 [926], EFNB1 [927], SOX17 [928], CYP27B1 [929], FOXC1 [930], ENG (endoglin) [931], EDNRB (endothelin receptor type B) [932], GAP43 [933], ADCYAP1 [934], FGF9 [935], NEGR1 [936], AHI1 [937], DCLK1 [938], SEMA3A [939], UCHL1 [940], MAP2 [941], LRRK2 [942], NRXN1 [323], C9ORF72 [943], SST (somatostatin) [944], PVALB (parvalbumin) [945], TAC1 [946], SPARCL1 [947], THEMIS (thymocyte selection associated) [948], GRIA3 [949], SYN3 [950] and GAD2 [951] genes were associated with multiple sclerosis. PECAM1 [952], ZAP70 [953], KLF4 [954], HSPB1 [955], FLT1 [956], TAB2 [957], FOXO4 [958], GLI2 [959], ANXA2 [867], CYP1A1 [960], CGAS (cyclic GMP-AMP synthase) [869], MMP2 [961], SERPINH1 [962], TGFB2 [963], LATS2 [964], GSTM1 [965], RUNX3 [966], CDKN1C [967], CCL2 [968]. LTF (lactotransferrin) [969], KCNQ1 [608], DLL4 [970], BMP6 [971], MLKL (mixed lineage kinase domain like pseudokinase) [972], S100A4 [973], MASP2 [974], ADORA2A [975], CYP27B1 [976]. ABCB1 [977], MMP14 [165], WNK4 [978], FLOT1 [979], CSF1 [980], ENG (endoglin) [981], EDNRB (endothelin receptor type B) [882], SLC12A3 [982], EPHX1 [983], ORAI1 [984], PCSK1 [413], GAP43 [985], MAS1 [986], THY1 [987], STMN2 [988], RGS4 [989], ADCYAP1 [990], FGF9 [991], NEGR1 [992], SNAP25 [993], VTN (vitronectin) [994], MEF2C [995], PAK1 [996], AHI1 [997], DCLK1 [998], HHIP (hedgehog interacting protein) [999], RBFOX2 [1000], SLIT2 [1001], SEMA3A [1002], UCHL1 [1003], EGR1 [1004], CITED2 [1005], PCSK2 [1006], LRRK2 [1007], RYR2 [1008], SST (somatostatin) [329], ICA1 [1009], CRHR2 [705], HTR3B [1010], GAD2 [1011], ATP1A1 [1012], CACNA1E [1013], SUV39H1 [1014], PDE10A [909], MSH5 [1015], ANKK1 [1016], HMGCR (3-hydroxy-3-methylglutaryl-CoA reductase) [1017], CISD1 [1018], LANCL2 [1019], TRPC3 [1020], FABP3 [1021], PDE1A [467] and PDE1A [1022] have been proposed as biomarkers for diabetes mellitus progression. Regulation of ZAP70 [1023], STIP1 [1024], KLF4 [1025], HSPB1 [1026], ITGA5 [1027], GATA2 [1028], CYP1A1 [1029], CGAS (cyclic GMP-AMP synthase) [1030], MMP2 [1031], TGFB2 [922], GSTM1 [1032], RUNX3 [1033], CCL2 [1034], LTF (lactotransferrin) [1035], HLA-DQB1 [1036], MSX1 [1037], ADORA2A [495], ABCB1 [1038], CSF1 [1039], RCC1 [1040], VGF (VGF nerve growth factor inducible) [1041], GAP43 [1042], THY1 [1043], SLC17A6 [1044], SYT1 [1045], SNAP25 [182], SYT4 [1046], MEF2C [1047], NMNAT2 [1048], CSMD1 [1049], CCKBR (cholecystokinin B receptor) [1050], SEMA3A [1051], UCHL1 [1052], ELAVL4 [1053], EGR1 [1054], CHRM1 [1055], MAP2 [1056], PTPRO (protein tyrosine phosphatase receptor type O) [1057], LRRK2 [203], STX1B [1058], SH3GL2 [1059], MTR (5-methyltetrahydrofolate-homocysteine methyltransferase) [1060], PLK2 [1061], C9ORF72 [1062], SST (somatostatin) [1063], PVALB (parvalbumin) [1064], SV2C [1065], SNCB (synuclein beta) [1066], CAP2 [214], TRIM9 [1067], HTR2C [1068], SYN3 [1069], PDE10A [1070], ANKK1 [1071], GNL1 [1072], NQO2 [1073], CISD1 [1074], ABCA5 [1075] and FABP3 [1076] levels might be a novel treatment option against Parkinson’s disease. Previous studies have shown that SMAD7 [1077], KLF4 [1078], FOXO4 [1079], ANXA2 [1080], CYP1A1 [1081], CGAS (cyclic GMP-AMP synthase) [1082], MMP2 [1083], GSTM1 [1084], CCL2 [1085], KCNQ1 [1086], HLA-DQB1 [1087], DUSP1 [1088], ABCC2 [1089], ADORA2A [1090], ABCB1 [1091], CACNA1H [1092], CELSR1 [1093], EPHX1 [1094], ORAI1 [1095], NES (nestin) [1096], GAP43 [1097], SCN8A [417], CDKL5 [1098], SERPINI1 [1099], FGF9 [1100], SYT1 [1101], SNAP25 [1102], GABRA1 [1103], FGF12 [1104], MEF2C [1105], GABRG2 [304], HCN1 [1106], CSMD3 [1107], PPP3CA [1108], SCN2A [1109], SCN5A [1110], CNTNAP2 [1111], GABRB3 [1112], SCN1A [1113], SLIT2 [1114], RORB (RAR related orphan receptor B) [1115], SEMA3A [1116], UCHL1 [1117], SLC4A10 [1118], ADAM23 [1119], EGR1 [1120], GABRB2 [1121], CHRM1 [1122], TUBB3 [1123], MAP2 [1124], ANK3 [1125], STX1B [1126], PLCB1 [1127], NRXN1 [1128], RYR2 [1129], VPS13A [1130], MTR (5-methyltetrahydrofolate-homocysteine methyltransferase) [1131], TUBB2A [1132], C9ORF72 [1133], SST (somatostatin) [1134], PVALB (parvalbumin) [1135], DLGAP2 [1136], KCNH1 [1137], NSF (N-ethylmaleimide sensitive factor, vesicle fusing ATPase) [1138], ATP1A3 [1139], GRIA3 [1140], GAD1 [1141], SV2B [1142], HTR2C [1143], NAPB (NSF attachment protein beta) [823], ATP6V1B2 [1144], CACNA1E [1145], VARS2 [1146], RPH3A [825], RPP21 [1147], IARS1 [1148], ANKK1 [1149] and TRPC3 [1150] might promote epilepsy. Recent studies have also suggested that RRAS (RAS related) [1151], CGAS (cyclic GMP-AMP synthase) [1152], DUSP1 [1153], ADORA2A [1154], NFATC4 [1155], VGF (VGF nerve growth factor inducible) [1156], FGF9 [1157], SNAP25 [1158], UCHL1 [1159], CHRM1 [1160], MAP2 [1161], NRXN1 [1162], VPS13A [1163], C9ORF72 [1164] , SST (somatostatin) [1165] and PDE10A [1166] might promote Huntington’s disease. LEF1 [1167], GSTM1 [1168], CCL2 [1169], HLA-DQB1 [1170], ABCB1 [1171], MDGA1 [1172], VGF (VGF nerve growth factor inducible) [1173], THY1 [675], SCN8A [1174], RGS4 [1175], CDKL5 [1176], ADCYAP1 [1177], SNAP25 [1178], NRN1 [1179], AHI1 [1180], CSMD1 [1181], RORB (RAR related orphan receptor B) [1182], GABRB2 [1183], MAP2 [692], CITED2 [1184], ANK3 [1125], HTR5A [1185], C9ORF72 [1186], SST (somatostatin) [1187], ATP1A3 [1186], NSF (N-ethylmaleimide sensitive factor, vesicle fusing ATPase) [1187], GRIA3 [1188], CAP2 [1191], CADPS (calcium dependent secretion activator) [1192], GAD1 [1193], HTR2C [1194], HTR3B [1195], TRANK1 [711], PDE10A [1196], ANKK1 [1197] and TRPC3 [1198] might play an important role in the onset, development, and treatment of bipolar disorder. Some studies have shown that HSPB1 [1199], GLI2 [1200], CGAS (cyclic GMP-AMP synthase) [1201], MMP2 [1202], FZD1 [1203], GSTM1 [1204], CCL2 [1205], HLA-DQB1 [1206], BMP6 [1207], MASP2 [1208], ADORA2A [1209], ABCB1 [1210], CELSR1 [1211], ZIC2 [1212], DES (desmin) [1213], MDGA1 [1172], VGF (VGF nerve growth factor inducible) [1214], GAP43 [1215], THY1 [675], RGS4 [1216], ADCYAP1 [1217], SCG2 [507], FGF9 [1218], SYT1 [1219], SNAP25 [680], FGF14 [1220], MEF2C [1221], NRN1 [1179], GABRG2 [1222], HCN1 [1223], PCDH8 [1224], PAK1 [1225], RAB3A [1226], AHI1 [1180], CSMD1 [1227], SCN2A [1228], SCN5A [1229], DCLK1 [1230], CCKBR (cholecystokinin B receptor) [1231], CNTNAP2 [685], GABRB3 [1232], SCN1A [800], PAK3 [1233], SEMA3A [689], UCHL1 [1234], EGR1 [1235], GABRB2 [1183], CHRM1 [1236], MAP2 [1237], ANK3 [1238], HTR5A [1239], LRRK2 [1240], PLCB1 [1241], NRXN1 [1242], MAP1A [1243], RYR2 [1244], C9ORF72 [1245], VAMP1 [1246], SST (somatostatin) [1247], PVALB (parvalbumin) [1248], DLGAP2 [1249], ATP1A3 [1250], KCNH1 [1251], SNCB (synuclein beta) [1252], NSF (N-ethylmaleimide sensitive factor, vesicle fusing ATPase) [1187], GRIA3 [1253], CAP2 [214], GAD1 [1254], HTR2C [1255], SYN3 [1256], HTR3B [1257], CHRM4 [1258], DMXL2 [709], GAD2 [1259], DLGAP1 [1260], VSNL1 [1261], PDE10A [1262], ANKK1 [1263], FSTL5 [1264], NQO2 [1265] and FABP3 [831] plays a certain role in schizophrenia. GATA2 [1266], MMP2 [1267], S100A4 [1268], EDNRB (endothelin receptor type B) [1269], PGF (placental growth factor) [1270], ORAI1 [1271], MAS1 [1272], VTN (vitronectin) [1273], SEMA3A [1274], UCHL1 [1275], EGR1 [1276], SUV39H1 [1277] and PDE10A [1278] were significantly altered in patients with vascular disease. ANXA2 [1279], MMP2 [1280], CCL2 [1281], ABCB1 [1282], PGF (placental growth factor) [1283], VGF (VGF nerve growth factor inducible) [1284], GAP43 [1285], SNAP25 [1286], CNTNAP2 [1287], UCHL1 [1288], CHRM1 [1055], MAP2 [1289], LRRK2 [1290], C9ORF72 [1133], SST (somatostatin) [1291], DLGAP2 [213], SNCB (synuclein beta) [1066], TRIM9 [1067], PDE10A [1292] and FABP3 [1293] can participate in the occurrence and development of dementia. The results of this investigation suggest that enriched genes might play a key role in the pathogenesis of AD and its associated complications.

The complexity and diversity of AD have hindered our accurate understanding of the disease. In this investigation, we used a comprehensive bioinformatics approach to find hub genes form PPI network and modules that influence the progression of AD and its associated complications. HSP90AA1 [139], FN1 [140], YAP1 [141], DDX39B [220], CDK7 [221], HSPA2 [145], HSPA1A [61], SYT1 [181], VAMP1 [208] and SNAP25 [182] might be involved in the development of AD. Regulation of HSP90AA1 [226], FN1 [227], YAP1 [228], SKP1 [1294], CDK7 [344], HSPB1 [234], HSPA1A [69], HSPA12B [289] and SNAP25 [299] levels might be a novel treatment option against inflammation. Previous studies have demonstrated that HSP90AA1 [352], FN1 [353], YAP1 [354], HSPB1 [361], HSPA1A [83], HSPA12B [412] and NSF (N-ethylmaleimide sensitive factor, vesicle fusing ATPase) [450] are linked with the development mechanisms of cardiovascular diseases. HSP90AA1 [468], YAP1 [469], HSPB1 [474], DNAJB1 [470], HSPA12B [504], SYT1 [509], NSF (N-ethylmaleimide sensitive factor, vesicle fusing ATPase) [531], VAMP1 [527] and SNAP25 [510] participate in pathogenic processes of brain injury. Previous research suggested some biomarkers, such as HSP90AA1 [468], STIP1 [544], HSPA12B [570] and SNAP25 [510] could be valuable in the diagnosis and prognosis of stroke. HSP90AA1 [589], YAP1 [590], NSF (N-ethylmaleimide sensitive factor, vesicle fusing ATPase) [649] and SNAP25 [632] expression has significant diagnosis value in hypertension patients and acts as potential targets for hypertension.- targeted therapy. Research has shown that HSP90AA1 [656], YAP1 [657], SYT1 [679], NSF (N-ethylmaleimide sensitive factor, vesicle fusing ATPase) [704] and SNAP25 [680] plays an important role in the pathogenesis of depression. Altered expression of FN1 [227], YAP1 [715], SKP1 [1295], DDX39B [778], CDK7 [781], STIP1 [719] and SNAP25 [763] are significantly associated with the infections. KIT (KIT proto-oncogene, receptor tyrosine kinase) [784], NAPB (NSF attachment protein beta) [823], NSF (N-ethylmaleimide sensitive factor, vesicle fusing ATPase) [818] and SNAP25 [791] are strongly involved in the pathogenesis of autism spectrum disorder. A recent study revealed that KIT (KIT proto-oncogene, receptor tyrosine kinase) [832] and HSPB1 [834] expression was elevated in amyotrophic lateral sclerosis. YAP1 [856], HSPB1 [861], HSPA2 [860], HSPA1A [89] and SNAP25 [892] expression has been found to be altered in patients with obesity. Altered levels of YAP1 [914] have been associated with impaired cognitive performance in patients with multiple sclerosis. Previous studies have shown that SKP1 [1296], HSPB1 [1026], STIP1 [1024], HSPA1A [95], SYT1 [1045], STX1B [1058] and SNAP25 [182] are closely associated with Parkinson’s disease. Excessive activation of HSPB1 [955], HSPA1A [77] and SNAP25 [993] have been observed in diabetes mellitus. Altered levels of HSPB1 [1199], HSPA1A [92], SYT1 [1219], NSF (N-ethylmaleimide sensitive factor, vesicle fusing ATPase) [1187], VAMP1 [1246] and SNAP25 [680] proteins exhibit schizophrenia. HSPA1A [65] plays an important role in the vascular disease. Changes in NAPB (NSF attachment protein beta) [823], SYT1 [1101], NSF (N-ethylmaleimide sensitive factor, vesicle fusing ATPase) [1138], STX1B [1126] and SNAP25 [1102] expression have been observed in epilepsy. The altered expression of NSF (N-ethylmaleimide sensitive factor, vesicle fusing ATPase) [1187] and SNAP25 [1178] are related to prognosis in bipolar disorder. SNAP25 [1158] is an emerging therapeutic target due to its regulated expression in Huntington’s disease. SNAP25 [1286] provided a clear picture of the prognosis of patients with dementia. LSM2, EIF5A2 and TAF9 might serve as a novel targets for early diagnosis and specific therapy of AD and its associated complications, and the related molecular mechanisms need to be further clarified. By intervening in AD hub genes, it might be a novel direction for individualized genomic therapy of AD and its associated complications.

To find the most important regulatory molecules among the hub genes, we analyzed the correlation between the hub genes and constructed a miRNA-hub gene regulatory network and TF-hub gene regulatory network between the hub genes and the miRNA and the TF. The miRNAs and TFs regulate the expression of many genes in cells, so abnormal expression of miRNAs and TFs might have an impact on the development of diseases. By examining the interaction between hub genes, miRNAs and TFs, we discovered that several miRNAs and TFs are implicated in AD. Previous studies focused on the role of HSP90AA1 [139], FN1 [140], YAP1 [141], DDX39B [220], CDK7 [221], hsa-mir-545-3p [1297], hsa-mir-183-5p [1298], CREB1 [1299], USF2 [1300], MEF2A [1301] and CEBPB (CCAAT enhancer binding protein beta) [1302] in AD development and growth. HSP90AA1 [226], FN1 [227], YAP1 [228], TTN (titin) [230], SMAD7 [231], SKP1 [1294], CDK7 [344], SUV39H1 [340], hsa-miR-23a-5p [1303], PLAG1 [1304], NFIC (nuclear factor I C) [1305], CREB1 [1306], MEF2A [1307] and CEBPB (CCAAT enhancer binding protein beta) [1308] are associated with prognosis in patients with inflammation. Increasing evidence has convincingly demonstrated that altered expression of HSP90AA1 [352], FN1 [353], YAP1 [354], TTN (titin) [230], SMAD7 [356], SUV39H1 [457], hsa-mir-296-3p [1309], NFIC (nuclear factor I C) [1310], CREB1 [1311], SREBF1 [1312], MEF2A [1313] and NFYA (nuclear transcription factor Y subunit alpha) [1314] are a prognosis factors in cardiovascular diseases. HSP90AA1 [468], YAP1 [469], SUV39H1 [535], USF2 [1315] and CEBPB (CCAAT enhancer binding protein beta) [1316] have been reported involved in the brain injury. HSP90AA1 [468], STIP1 [544], TTN (titin) [543], SUV39H1 [587] and CEBPB (CCAAT enhancer binding protein beta) [1317] were identified as a candidate causal biomarkers of a stroke. HSP90AA1 [589], YAP1 [590], SMAD7 [592] and NFYA (nuclear transcription factor Y subunit alpha) [1314] were positively associated with hypertension. HSP90AA1 [656], YAP1 [657], SUV39H1 [712], hsa-mir-183-5p [1318], CREB1 [1319] and MAX (MYC associated factor X) [1320] expression might be regarded as an indicator of susceptibility to depression. Altered expression of FN1 [227], YAP1 [715], STIP1 [719], SMAD7 [718], ZAP70 [717], DDX39B [778], SKP1 [1295], CDK7 [781], SUV39H1 [776] and hsa-miR-23a-5p [1321] are associated with prognosis in patients with infections. YAP1 [856], SUV39H1 [908], hsa-mir-296-3p [1322], hsa-miR-23a-5p [1323], PLAG1 [1324], SREBF1 [1325] and CEBPB (CCAAT enhancer binding protein beta) [1326] are a potential markers for the detection and prognosis of obesity at an early age. A previous study reported that YAP1 [914], SMAD7 [916] and hsa-mir-183-5p [1327] are altered expressed in multiple sclerosis. The levels of STIP1 [1024], ZAP70 [1023], SKP1 [1296], hsa-miR-23a-5p [1328], CREB1 [1329] and SREBF1 [1330] might be a predictive biomarkers for the Parkinson’s disease. Research has revealed that TTN (titin) [833], hsa-mir-183-5p [1331], hsa-miR-23a-5p [1332] and SREBF1 [1330] are expressed in amyotrophic lateral sclerosis. SMAD7 [1077], hsa-miR-23a-5p [1333], hsa-mir-4521 [1334] and MEF2A [1335] might aid in the development of personalized therapies for patients with epilepsy. ZAP70 [953], SUV39H1 [1014], hsa-mir-296-3p [1322], hsa-miR-23a-5p [1336], CREB1 [1337] and SREBF1 [1338] are essential for diabetes mellitus development. Recent studies have shown that TCERG1 [1339] and CREB1 [1340] might play an important role in regulating the Huntington’s disease. SUV39H1 [1277] expression was shown to be regulated in vascular disease and associated with prognosis. Hsa-mir-183-5p [1298] and SREBF1 [1341] were associated with dementia. CREB1 [1342] was an important target biomarker of bipolar disorder. CREB1 [1343] and SREBF1 [1344] have been reported to be expressed in schizophrenia. Hence, novel biomarkers include LSM2, EIF5A2, PLRG1, GTF2H4, hsa-miR-3909, hsa-miR-548f-5p, hsa-miR-429, hsa-miR-4735-5p, hsa-miR-489-3p and POU2F2 might play a role in the development of AD through transcriptional regulation-related mechanisms. This investigation suggests that these hub genes, miRNAs and TFs are closely related to the development of AD and its associated complications.

In conclusion, the present investigation identified novel genes and signaling pathways which might be associated in AD progression through the integrated analysis of NGS dataset. These results might contribute to a better understanding of the molecular mechanisms which underlie AD and provide a series of key biomarkers. However, further experiments are required to validate the findings of the current investigations. Additionally, the majority of included investigations focused on how a hub gene, miRNAs, TFs and signaling pathway contribute to the advancement of AD, with limited study concerning the interaction of multi-genes and multi-pathways. Therefore, further experiments with additional patient cohorts are also required to confirm the results of this investigation. In vivo and in vitro investigation of gene and pathway interaction is essential to delineate the specific roles of the identified novel genes, which might help to confirm gene functions and reveal the mechanisms underlying AD.

## Acknowledgement

I thanks very much to Caldwell AB, Anantharaman BG, Ramachandran S, Galasko DR, Desplats PA, Wagner SL, Subramaniam S, Department of Bioengineering, University of California, La Jolla, San Diego, CA, the authors who deposited their NGS dataset GSE203206, into the public GEO database.

## Ethical approval

This article does not contain any studies with human participants or animals performed by any of the authors.

## Informed consent

No informed consent because this study does not contain human or animals participants.

## Availability of data and materials

The datasets supporting the conclusions of this article are available in the GEO (Gene Expression Omnibus) (https://www.ncbi.nlm.nih.gov/geo/) repository. [(GSE203206) https://www.ncbi.nlm.nih.gov/geo/query/acc.cgi?acc= GSE203206]

## Consent for publication

Not applicable.

## Author Contributions

B. V. - Writing original draft, and review and editing

C. V. - Software and investigation

## References

1. Murphy MP, LeVine H 3rd. Alzheimer’s disease and the amyloid-beta peptide. J Alzheimers Dis. 2010;19(1):311–323. doi:10.3233/JAD-2010-1221

2. Gorantla NV, Chinnathambi S. Autophagic Pathways to Clear the Tau Aggregates in Alzheimer’s Disease. Cell Mol Neurobiol. 2021;41(6):1175–1181. doi:10.1007/s10571-020-00897-0

3. Fukutani Y, Cairns NJ, Shiozawa M, Sasaki K, Sudo S, Isaki K, Lantos PL. Neuronal loss and neurofibrillary degeneration in the hippocampal cortex in late-onset sporadic Alzheimer’s disease. Psychiatry Clin Neurosci. 2000;54(5):523–529. doi:10.1046/j.1440-1819.2000.00747.x

4. Zvěřová M. Clinical aspects of Alzheimer’s disease. Clin Biochem. 2019;72:3–6. doi:10.1016/j.clinbiochem.2019.04.015

5. Reitz C. Genetic diagnosis and prognosis of Alzheimer’s disease: challenges and opportunities. Expert Rev Mol Diagn. 2015;15(3):339–348. doi:10.1586/14737159.2015.1002469

6. Stern Y. Cognitive reserve in ageing and Alzheimer’s disease. Lancet Neurol. 2012;11(11):1006–1012. doi:10.1016/S1474-4422(12)70191-6

7. Serrano-Pozo A, Das S, Hyman BT. APOE and Alzheimer’s disease: advances in genetics, pathophysiology, and therapeutic approaches. Lancet Neurol. 2021;20(1):68–80. doi:10.1016/S1474-4422(20)30412-9

8. Tolppanen AM, Taipale H, Hartikainen S. Head or brain injuries and Alzheimer’s disease: A nested case-control register study. Alzheimers Dement. 2017;13(12):1371–1379. doi:10.1016/j.jalz.2017.04.010

9. Guo T, Landau SM, Jagust WJ; Alzheimer’s Disease Neuroimaging Initiative. Age, vascular disease, and Alzheimer’s disease pathologies in amyloid negative elderly adults. Alzheimers Res Ther. 2021;13(1):174. doi:10.1186/s13195-021-00913-5

10. Vigasova D, Nemergut M, Liskova B, Damborsky J. Multi-pathogen infections and Alzheimer’s disease. Microb Cell Fact. 2021;20(1):25. doi:10.1186/s12934-021-01520-7

11. Akiyama H, Barger S, Barnum S, Bradt B, Bauer J, Cole GM, Cooper NR, Eikelenboom P, Emmerling M, Fiebich BL, et al. Inflammation and Alzheimer’s disease. Neurobiol Aging. 2000;21(3):383–421. doi:10.1016/s0197-4580(00)00124-x

12. Wainaina MN, Chen Z, Zhong C. Environmental factors in the development and progression of late-onset Alzheimer’s disease. Neurosci Bull. 2014;30(2):253–270. doi:10.1007/s12264-013-1425-9

13. Burillo J, Marqués P, Jiménez B, González-Blanco C, Benito M, Guillén C. Insulin Resistance and Diabetes Mellitus in Alzheimer’s Disease. Cells. 2021;10(5):1236. doi:10.3390/cells10051236

14. Picone P, Di Carlo M, Nuzzo D. Obesity and Alzheimer’s disease: Molecular bases. Eur J Neurosci. 2020;52(8):3944–3950. doi:10.1111/ejn.14758

15. Malone JE, Elkasaby MI, Lerner AJ. Effects of Hypertension on Alzheimer’s Disease and Related Disorders. Curr Hypertens Rep. 2022;24(12):615–625. doi:10.1007/s11906-022-01221-5

16. Leszek J, Mikhaylenko EV, Belousov DM, Koutsouraki E, Szczechowiak K, Kobusiak-Prokopowicz M, Mysiak A, Diniz BS, Somasundaram SG, Kirkland CE, et al. The Links between Cardiovascular Diseases and Alzheimer’s Disease. Curr Neuropharmacol. 2021;19(2):152–169. doi:10.2174/1570159X18666200729093724

17. Yuan D, Huang B, Gu M, Qin BE, Su Z, Dai K, Peng FH, Jiang Y. Exploring Shared Genetic Signatures of Alzheimer’s Disease and Multiple Sclerosis: A Bioinformatic Analysis Study. Eur Neurol. 2023;86(6):363–376. doi:10.1159/000533397

18. Graves LV, Holden HM, Delano-Wood L, Bondi MW, Woods SP, Corey-Bloom J, Salmon DP, Delis DC, Gilbert PE. Total recognition discriminability in Huntington’s and Alzheimer’s disease. J Clin Exp Neuropsychol. 2017;39(2):120–130. doi:10.1080/13803395.2016.1204993

19. Zhuang Z, Yang R, Wang W, Qi L, Huang T. Associations between gut microbiota and Alzheimer’s disease, major depressive disorder, and schizophrenia. J Neuroinflammation. 2020;17(1):288. doi:10.1186/s12974-020-01961-8

20. Alexiou A, Soursou G, Yarla NS, Md Ashraf G. Proteins Commonly Linked to Autism Spectrum Disorder and Alzheimer’s Disease. Curr Protein Pept Sci. 2018;19(9):876–880. doi:10.2174/1389203718666170911145321

21. Verde F, Aiello EN, Adobbati L, Poletti B, Solca F, Tiloca C, Sangalli D, Maranzano A, Muscio C, Ratti A, Zago S, et al. Coexistence of Amyotrophic Lateral Sclerosis and Alzheimer’s Disease: Case Report and Review of the Literature. J Alzheimers Dis. 2023;95(4):1383–1399. doi:10.3233/JAD-230562

22. Zhou J, Yu JT, Wang HF, Meng XF, Tan CC, Wang J, Wang C, Tan L. Association between stroke and Alzheimer’s disease: systematic review and meta-analysis. J Alzheimers Dis. 2015;43(2):479–489. doi:10.3233/JAD-140666

23. Fang Y, Si X, Wang J, Wang Z, Chen Y, Liu Y, Yan Y, Tian J, Zhang B, Pu J. Alzheimer Disease and Epilepsy: A Mendelian Randomization Study. Neurology. 2023;101(4):e399–e409. doi:10.1212/WNL.0000000000207423

24. Porsteinsson AP, Rangaraju S, Spires-Jones TL, O’Banion MK. Alzheimer’s disease and related dementias: From risk factors to disease pathogenesis. Eur J Neurosci. 2022;56(9):5337–5341. doi:10.1111/ejn.15857

25. Mayo S, Benito-León J, Peña-Bautista C, Baquero M, Cháfer-Pericás C. Recent Evidence in Epigenomics and Proteomics Biomarkers for Early and Minimally Invasive Diagnosis of Alzheimer’s and Parkinson’s Diseases. Curr Neuropharmacol. 2021;19(8):1273–1303. doi:10.2174/1570159X19666201223154009

26. Besga A, Gonzalez I, Echeburua E, Savio A, Ayerdi B, Chyzhyk D, Madrigal JL, Leza JC, Graña M, Gonzalez-Pinto AM. Discrimination between Alzheimer’s Disease and Late Onset Bipolar Disorder Using Multivariate Analysis. Front Aging Neurosci. 2015;7:231. doi:10.3389/fnagi.2015.00231

27. Huang YY, Gan YH, Yang L, Cheng W, Yu JT. Depression in Alzheimer’s Disease: Epidemiology, Mechanisms, and Treatment. Biol Psychiatry. 2024;95(11):992–1005. doi:10.1016/j.biopsych.2023.10.008

28. Sharma K. Cholinesterase inhibitors as Alzheimer’s therapeutics (Review). Mol Med Rep. 2019;20(2):1479–1487. doi:10.3892/mmr.2019.10374

29. Müller WE, Mutschler E, Riederer P. Noncompetitive NMDA receptor antagonists with fast open-channel blocking kinetics and strong voltage-dependency as potential therapeutic agents for Alzheimer’s dementia. Pharmacopsychiatry. 1995;28(4):113–124. doi:10.1055/s-2007-979603

30. Peng Y, Jin H, Xue YH, Chen Q, Yao SY, Du MQ, Liu S. Current and future therapeutic strategies for Alzheimer’s disease: an overview of drug development bottlenecks. Front Aging Neurosci. 2023;15:1206572. doi:10.3389/fnagi.2023.1206572

31. Pujar M, Vastrad B, Kavatagimath S, Vastrad C, Kotturshetti S. Identification of candidate biomarkers and pathways associated with type 1 diabetes mellitus using bioinformatics analysis. Sci Rep. 2022;12(1):9157. doi:10.1038/s41598-022-13291-1

32. Ganekal P, Vastrad B, Vastrad C, Kotrashetti S. Identification of biomarkers, pathways, and potential therapeutic targets for heart failure using next-generation sequencing data and bioinformatics analysis. Ther Adv Cardiovasc Dis. 2023;17:17539447231168471. doi:10.1177/17539447231168471

33. Quiroz YT, Aguillon D, Aguirre-Acevedo DC, Vasquez D, Zuluaga Y, Baena AY, Madrigal L, Hincapié L, Sanchez JS, Langella S, et al. APOE3 Christchurch Heterozygosity and Autosomal Dominant Alzheimer’s Disease. N Engl J Med. 2024;390(23):2156–2164. doi:10.1056/NEJMoa2308583

34. Xu C, Liu G, Ji H, Chen W, Dai D, Chen Z, Zhou D, Xu L, Hu H, Cui W, et al. Elevated methylation of OPRM1 and OPRL1 genes in Alzheimer’s disease. Mol Med Rep. 2018;18(5):4297–4302. doi:10.3892/mmr.2018.9424

35. Davies DA, Adlimoghaddam A, Albensi BC. Role of Nrf2 in Synaptic Plasticity and Memory in Alzheimer’s Disease. Cells. 2021;10(8):1884. doi:10.3390/cells10081884

36. Zajac DJ, Simpson J, Zhang E, Parikh I, Estus S. Expression of INPP5D Isoforms in Human Brain: Impact of Alzheimer’s Disease Neuropathology and Genetics. Genes (Basel). 2023;14(3):763. doi:10.3390/genes14030763

37. Parikh I, Fardo DW, Estus S. Genetics of PICALM expression and Alzheimer’s disease. PLoS One. 2014;9(3):e91242. doi:10.1371/journal.pone.0091242

38. Akhtar A, Sah SP. Insulin signaling pathway and related molecules: Role in neurodegeneration and Alzheimer’s disease. Neurochem Int. 2020;135:104707. doi:10.1016/j.neuint.2020.104707

39. Deczkowska A, Weiner A, Amit I. The Physiology, Pathology, and Potential Therapeutic Applications of the TREM2 Signaling Pathway. Cell. 2020;181(6):1207–1217. doi:10.1016/j.cell.2020.05.003

40. Wang J, Zhang J, Yu ZL, Chung SK, Xu B. The roles of dietary polyphenols at crosstalk between type 2 diabetes and Alzheimer’s disease in ameliorating oxidative stress and mitochondrial dysfunction via PI3K/Akt signaling pathways. Ageing Res Rev. 2024;99:102416. doi:10.1016/j.arr.2024.102416

41. Kapoor A, Nation DA. Role of Notch signaling in neurovascular aging and Alzheimer’s disease. Semin Cell Dev Biol. 2021;116:90–97. doi:10.1016/j.semcdb.2020.12.011

42. Saura CA, Valero J. The role of CREB signaling in Alzheimer’s disease and other cognitive disorders. Rev Neurosci. 2011;22(2):153–169. doi:10.1515/RNS.2011.018

43. Caldwell AB, Anantharaman BG, Ramachandran S, Nguyen P, Liu Q, Trinh I, Galasko DR, Desplats PA, Wagner SL, Subramaniam S. Transcriptomic profiling of sporadic Alzheimer’s disease patients. Mol Brain. 2022;15(1):83. doi:10.1186/s13041-022-00963-2

44. Clough E, Barrett T. The Gene Expression Omnibus Database. Methods Mol Biol. 2016;1418:93–110. doi:10.1007/978-1-4939-3578-9_5

45. Ritchie ME, Phipson B, Wu D, Hu Y, Law CW, Shi W, Smyth GK. limma powers differential expression analyses for RNA-sequencing and microarray studies. Nucleic Acids Res. 2015;43(7):e47. doi:10.1093/nar/gkv007

46. Green GH, Diggle PJ. On the operational characteristics of the Benjamini and Hochberg False Discovery Rate procedure. Stat Appl Genet Mol Biol. 2007;6:Article27. doi:10.2202/1544-6115.1302

47. Reimand J, Kull M, Peterson H, Hansen J, Vilo J. g:Profiler--a web-based toolset for functional profiling of gene lists from large-scale experiments. Nucleic Acids Res. 2007;35(Web Server issue):W193–W200. doi:10.1093/nar/gkm226

48. Thomas PD. The Gene Ontology and the Meaning of Biological Function. Methods Mol Biol. 2017;1446:15 24. doi:10.1007/978-1-4939-3743-1_2

49. Fabregat A, Jupe S, Matthews L, Sidiropoulos K, Gillespie M, Garapati P, Haw R, Jassal B, Korninger F, May B et al The Reactome Pathway Knowledgebase. Nucleic Acids Res. 2018;46(D1):D649–D655. doi:10.1093/nar/gkx1132

50. Szklarczyk D, Gable AL, Lyon D, Junge A, Wyder S, Huerta-Cepas J, Simonovic M, Doncheva NT, >Morris JH, Bork P, et al. STRING v11: protein-protein association networks with increased coverage, supporting functional discovery in genome-wide experimental datasets. Nucleic Acids Res. 2019;47(d1):D607-D613. doi:10.1093/nar/gky113

51. Shannon P, Markiel A, Ozier O, Baliga NS, Wang JT, Ramage D, Amin N, Schwikowski B, Ideker T Cytoscape: a software environment for integrated models of biomolecular interaction networks. Genome Res 2003;13(11):2498–2504. doi:10.1101/gr.1239303

52. Luo X, Guo L, Dai XJ, Wang Q, Zhu W, Miao X, Gong H. Abnormal intrinsic functional hubs in alcohol dependence: evidence from a voxelwise degree centrality analysis. Neuropsychiatr Dis Treat. 2017;13:2011–2020. doi:10.2147/NDT.S142742

53. Li Y, Li W, Tan Y, Liu F, Cao Y, Lee KY. Hierarchical Decomposition for Betweenness Centrality Measure of Complex Networks. Sci Rep. 2017;7:46491.. doi:10.1038/srep46491

54. Gilbert M, Li Z, Wu XN, Rohr L, Gombos S, Harter K, Schulze WX. Comparison of path-based centrality measures in protein-protein interaction networks revealed proteins with phenotypic relevance during adaptation to changing nitrogen environments. J Proteomics. 2021;235:104114. doi:10.1016/j.jprot.2021.104114

55. Li G, Li M, Wang J, Li Y, Pan Y. United Neighborhood Closeness Centrality and Orthology for Predicting Essential Proteins. IEEE/ACM Trans Comput Biol Bioinform. 2020;17(4):1451–1458. doi:10.1109/TCBB.2018.2889978

56. Zaki N, Efimov D, Berengueres J. Protein complex detection using interaction reliability assessment and weighted clustering coefficient. BMC Bioinformatics. 2013;14:163. doi:10.1186/1471-2105-14

57. Fan Y, Xia J (2018) miRNet-Functional Analysis and Visual Exploration of miRNA-Target Interactions in a Network Context. Methods Mol Biol 1819:215–233. doi:10.1007/978-1-4939-8618-7_10

58. Zhou G, Soufan O, Ewald J, Hancock REW, Basu N, Xia J (2019) NetworkAnalyst 3.0: a visual analytics platform for comprehensive gene expression profiling and meta-analysis. Nucleic Acids Res 47:W234–W241. doi:10.1093/nar/gkz240

59. Robin X, Turck N, Hainard A, Tiberti N, Lisacek F, Sanchez JC, Müller M. pROC: an open-source package for R and S+ to analyze and compare ROC curves. BMC Bioinformatics 2011;12:77. doi:10.1186/1471-2105-12-77

60. Rostagno AA. Pathogenesis of Alzheimer’s Disease. Int J Mol Sci. 2022;24(1):107. doi:10.3390/ijms24010107

61. Wang Y, Yang Y, Liang C, Zhang H. Exploring the Roles of Key Mediators IKBKE and HSPA1A in Alzheimer’s Disease and Hepatocellular Carcinoma through Bioinformatics Analysis. Int J Mol Sci. 2024;25(13):6934. doi:10.3390/ijms25136934

62. Jafarian Z, Saliminejad K, Kamali K, Ohadi M, Kowsari A, Nasehi L, Khorram Khorshid HR. Association of glutathione S-transferases M1, P1 and T1 variations and risk of late-onset Alzheimer’s disease. Neurol Res. 2018;40(1):41–44. doi:10.1080/01616412.2017.1390902

63. Le Guen Y, Luo G, Ambati A, Damotte V, Jansen I, Yu E, Nicolas A, de Rojas I, Peixoto Leal T, Miyashita A, et al. Multiancestry analysis of the HLA locus in Alzheimer’s and Parkinson’s diseases uncovers a shared adaptive immune response mediated by HLA-DRB1*04 subtypes. Proc Natl Acad Sci U S A. 2023;120(36):e2302720120. doi:10.1073/pnas.2302720120

64. Smyth LCD, Murray HC, Hill M, van Leeuwen E, Highet B, Magon NJ, Osanlouy M, Mathiesen SN, Mockett B, Singh-Bains MK, et al. Neutrophil-vascular interactions drive myeloperoxidase accumulation in the brain in Alzheimer’s disease. Acta Neuropathol Commun. 2022;10(1):38. doi:10.1186/s40478-022-01347-2

65. Dulin E, García-Barreno P, Guisasola MC. Extracellular heat shock protein 70 (HSPA1A) and classical vascular risk factors in a general population. Cell Stress Chaperones. 2010;15(6):929–937. doi:10.1007/s12192-010-0201-2

66. Nørskov MS, Frikke-Schmidt R, Loft S, Sillesen H, Grande P, Nordestgaard BG, Tybjaerg-Hansen A. Copy number variation in glutathione S-transferases M1 and T1 and ischemic vascular disease: four studies and meta-analyses. Circ Cardiovasc Genet. 2011;4(4):418-428. doi:10.1161/CIRCGENETICS.111.959809

67. Lundström E, Gustafsson JT, Jönsen A, Leonard D, Zickert A, Elvin K, Sturfelt G, Nordmark G, Bengtsson AA, Sundin U, et al. HLA-DRB1*04/*13 alleles are associated with vascular disease and antiphospholipid antibodies in systemic lupus erythematosus. Ann Rheum Dis. 2013;72(6):1018–1025. doi:10.1136/annrheumdis-2012-201760

68. Lau D, Baldus S. Myeloperoxidase and its contributory role in inflammatory vascular disease. Pharmacol Ther. 2006;111(1):16–26. doi:10.1016/j.pharmthera.2005.06.023

69. Kotowska J, Jówko E, Cieśliński I, Gromisz W, Sadowski J. IL-6 and HSPA1A Gene Polymorphisms May Influence the Levels of the Inflammatory and Oxidative Stress Parameters and Their Response to a Chronic Swimming Training. Int J Environ Res Public Health. 2022;19(13):8127. doi:10.3390/ijerph19138127

70. Huang S, Lu W, Ge D, Meng N, Li Y, Su L, Zhang S, Zhang Y, Zhao B, Miao J. A new microRNA signal pathway regulated by long noncoding RNA TGFB2-OT1 in autophagy and inflammation of vascular endothelial cells. Autophagy. 2015;11(12):2172–2183. doi:10.1080/15548627.2015.1106663

71. Zeng X, Tian G, Zhu J, Yang F, Zhang R, Li H, An Z, Li J, Song J, Jiang J, et al. Air pollution associated acute respiratory inflammation and modification by GSTM1 and GSTT1 gene polymorphisms: a panel study of healthy undergraduates. Environ Health. 2023;22(1):14. doi:10.1186/s12940-022-00954-9

72. Matsunaga N, Ikeda E, Kakimoto K, Watanabe M, Shindo N, Tsuruta A, Ikeyama H, Hamamura K, Higashi K, Yamashita T, et al. Inhibition of G0/G1 Switch 2 Ameliorates Renal Inflammation in Chronic Kidney Disease. EBioMedicine. 2016;13:262–273. doi:10.1016/j.ebiom.2016.10.008

73. Luo Y, He Y, Wang Y, Xu Y, Yang L. LncRNA HCG18 promotes inflammation and apoptosis in intervertebral disc degeneration via the miR-495-3p/FSTL1 axis. Mol Cell Biochem. 2024;479(1):171–181. doi:10.1007/s11010-023-04716-0

74. Klimenta B, Nefic H, Prodanovic N, Jadric R, Hukic F. Association of biomarkers of inflammation and HLA-DRB1 gene locus with risk of developing rheumatoid arthritis in females. Rheumatol Int. 2019;39(12):2147–2157. doi:10.1007/s00296-019-04429-y

75. Zhao G, Li Y, Chen T, Liu F, Zheng Y, Liu B, Zhao W, Qi X, Sun W, Gao C. TRIM26 alleviates fatal immunopathology by regulating inflammatory neutrophil infiltration during Candida infection. PLoS Pathog. 2024;20(1):e1011902. doi:10.1371/journal.ppat.1011902

76. Chen S, Chen H, Du Q, Shen J. Targeting Myeloperoxidase (MPO) Mediated Oxidative Stress and Inflammation for Reducing Brain Ischemia Injury: Potential Application of Natural Compounds. Front Physiol. 2020;11:433. doi:10.3389/fphys.2020.00433

77. Rosas PC, Nagaraja GM, Kaur P, Panossian A, Wickman G, Garcia LR, Al-Khamis FA, Asea AA. Hsp72 (HSPA1A) Prevents Human Islet Amyloid Polypeptide Aggregation and Toxicity: A New Approach for Type 2 Diabetes Treatment. PLoS One. 2016;11(3):e0149409. doi:10.1371/journal.pone.0149409

78. Sobha SP, Kesavarao KE. Progonostic effect of GSTM1/GSTT1 polymorphism in determining cardiovascular diseases risk among type 2 diabetes patients in South Indian population. Mol Biol Rep. 2023;50(8):6415–6423. doi:10.1007/s11033-023-08514-1

79. El-Assaad W, El-Kouhen K, Mohammad AH, Yang J, Morita M, Gamache I, Mamer O, Avizonis D, Hermance N, Kersten S, et al. Deletion of the gene encoding G0/G 1 switch protein 2 (G0s2) alleviates high-fat-diet-induced weight gain and insulin resistance, and promotes browning of white adipose tissue in mice. Diabetologia. 2015;58(1):149–157. doi:10.1007/s00125-014-3429-z

80. Xia Y, Zhang Y, Wang H. Upregulated lncRNA HCG18 in Patients with Non-Alcoholic Fatty Liver Disease and Its Regulatory Effect on Insulin Resistance. Diabetes Metab Syndr Obes. 2021;14:4747–4756. doi:10.2147/DMSO.S333431

81. Sayad A, Akbari MT, Pajouhi M, Mostafavi F, Kazemnejad A, Zamani M. Investigation The Role of Gender on The HLA-DRB1 and -DQB1 Association with Type 1 Diabetes Mellitus in Iranian Patients. Cell J. 2013;15(2):108–115.

82. Gómez García A, Rivera Rodríguez M, Gómez Alonso C, Rodríguez Ochoa DY, Alvarez Aguilar C. Myeloperoxidase is associated with insulin resistance and inflammation in overweight subjects with first-degree relatives with type 2 diabetes mellitus. Diabetes Metab J. 2015;39(1):59–65. doi:10.4093/dmj.2015.39.1.59

83. Jenei ZM, Széplaki G, Merkely B, Karádi I, Zima E, Prohászka Z. Persistently elevated extracellular HSP70 (HSPA1A) level as an independent prognostic marker in post-cardiac-arrest patients. Cell Stress Chaperones. 2013;18(4):447–454. doi:10.1007/s12192-012-0399-2

84. Huang RT, Guo YH, Yang CX, Gu JN, Qiu XB, Shi HY, Xu YJ, Xue S, Yang YQ. SOX7 loss-of-function variation as a cause of familial congenital heart disease. Am J Transl Res. 2022;14(3):1672–1684.

85. Xiong T, Lv XS, Wu GJ, Guo YX, Liu C, Hou FX, Wang JK, Fu YF, Liu FQ. Single-Cell Sequencing Analysis and Multiple Machine Learning Methods Identified G0S2 and HPSE as Novel Biomarkers for Abdominal Aortic Aneurysm. Front Immunol. 2022;13:907309. doi:10.3389/fimmu.2022.907309

86. Luo Y, Jiang Y, Zhong T, Li Z, He J, Li X, Cui K. LncRNA HCG18 affects diabetic cardiomyopathy and its association with miR-9-5p/IGF2R axis. Heliyon. 2024;10(3):e24604. doi:10.1016/j.heliyon.2024.e24604

87. Sreekanth MS, Esdan Basha SK, Arun Kumar G, Govindaraju S, Pradeep Nayar N, Pitchappan RM. Association of IL-1β +3953 C and HLA-DRB1*15 with Coronary Artery and Rheumatic Heart Diseases in South India. Hum Immunol. 2016;77(12):1275–1279. doi:10.1016/j.humimm.2016.08.003

88. Ramachandra CJA, Ja KPMM, Chua J, Cong S, Shim W, Hausenloy DJ. Myeloperoxidase As a Multifaceted Target for Cardiovascular Protection. Antioxid Redox Signal. 2020;32(15):1135–1149. doi:10.1089/ars.2019.7971

89. Pilch W, Wyrostek J, Major P, Zuziak R, Piotrowska A, Czerwińska-Ledwig O, Grzybkowska A, Zasada M, Ziemann E, Żychowska M. The effect of whole-body cryostimulation on body composition and leukocyte expression of HSPA1A, HSPB1, and CRP in obese men. Cryobiology. 2020;94:100–106. doi:10.1016/j.cryobiol.2020.04.002

90. Cui Z, Liu Y, Wan W, Xu Y, Hu Y, Ding M, Dou X, Wang R, Li H, Meng Y, et al. Ethacrynic acid targets GSTM1 to ameliorate obesity by promoting browning of white adipocytes. Protein Cell. 2021;12(6):493–501. doi:10.1007/s13238-020-00717-7

91. Qaddoumi MG, Alanbaei M, Hammad MM, Al Khairi I, Cherian P, Channanath A, Thanaraj TA, Al-Mulla F, Abu-Farha M, Abubaker J. Investigating the Role of Myeloperoxidase and Angiopoietin-like Protein 6 in Obesity and Diabetes. Sci Rep. 2020;10(1):6170. doi:10.1038/s41598-020-63149-7

92. Kowalczyk M, Kucia K, Owczarek A, Suchanek-Raif R, Merk W, Paul-Samojedny M, Kowalski J. Association Studies of HSPA1A and HSPA1L Gene Polymorphisms With Schizophrenia. Arch Med Res. 2018;49(5):342–349. doi:10.1016/j.arcmed.2018.10.002

93. Liu H, Xu Y, Peng J. Glutathione S-Transferase M1/T1 Polymorphisms and Schizophrenia Risk: A New Method for Quality Assessment and a Systematic Review. Neuropsychiatr Dis Treat. 2023;19:97–107. doi:10.2147/NDT.S376942

94. Seshasubramanian V, Raghavan V, SathishKannan AD, Naganathan C, Ramachandran A, Arasu P, Rajendren P, John S, Mowry B, Rangaswamy T, et al. Association of HLA-A, -B, -C, -DRB1 and -DQB1 alleles at amino acid level in individuals with schizophrenia: A study from South India. Int J Immunogenet. 2020;47(6):501–511. doi:10.1111/iji.12507

95. Asad Samani L, Ghaedi K, Majd A, Peymani M, Etemadifar M. Coordinated modification in expression levels of HSPA1A/B, DGKH, and NOTCH2 in Parkinson’s patients’ blood and substantia nigra as a diagnostic sign: the transcriptomes’ relationship. Neurol Sci. 2023;44(8):2753–2761. doi:10.1007/s10072-023-06738-4

96. Pinhel MA, Sado CL, Longo Gdos S, Gregório ML, Amorim GS, Florim GM, Mazeti CM, Martins DP, Oliveira Fde N, Nakazone MA, et al. ullity of GSTT1/GSTM1 related to pesticides is associated with Parkinson’s disease. Arq Neuropsiquiatr. 2013;71(8):527–532. doi:10.1590/0004-282X20130076

97. Boonpraman N, Yoon S, Kim CY, Moon JS, Yi SS. NOX4 as a critical effector mediating neuroinflammatory cytokines, myeloperoxidase and osteopontin, specifically in astrocytes in the hippocampus in Parkinson’s disease. Redox Biol. 2023;62:102698. doi:10.1016/j.redox.2023.102698

98. Salpietro V, Galassi Deforie V, Efthymiou S, O’Connor E, Marcé-Grau A, Maroofian R, Striano P, Zara F, Morrow MM; SYNAPS Study Group; et al. De novo KCNA6 variants with attenuated KV 1.6 channel deactivation in patients with epilepsy. Epilepsia. 2023;64(2):443–455. doi:10.1111/epi.17455

99. Prabha TS, Kumaraswami K, Kutala VK. Association of GSTT1 and GSTM1 polymorphisms in South Indian Epilepsy Patients. Indian J Exp Biol. 2016;54(11):783–787.

100. Zhu Z, Wang S, Cao Q, Li G. CircUBQLN1 Promotes Proliferation but Inhibits Apoptosis and Oxidative Stress of Hippocampal Neurons in Epilepsy via the miR-155-Mediated SOX7 Upregulation. J Mol Neurosci. 2021;71(9):1933–1943. doi:10.1007/s12031-021-01838-2

101. Bui TP, Nguyen LTT, Le PL, Le NTT, Nguyen TD, Van Nguyen L, Van Nguyen AT, Trinh TH. Next-generation sequencing-based HLA typing reveals the association of HLA-B*46:01:01 and HLA-DRB1*09:01:02 alleles with carbamazepine-induced hypersensitivity reactions in Vietnamese patients with epilepsy. Hum Immunol. 2023;84(3):186–195. doi:10.1016/j.humimm.2023.01.005

102. Shao C, Yuan J, Liu Y, Qin Y, Wang X, Gu J, Chen G, Zhang B, Liu HK, Zhao J, et al. Epileptic brain fluorescent imaging reveals apigenin can relieve the myeloperoxidase-mediated oxidative stress and inhibit ferroptosis. Proc Natl Acad Sci U S A. 2020;117(19):10155–10164. doi:10.1073/pnas.1917946117

103. Chernyak YI, Itskovich VB, D’yakovich OA, Kolesnikov SI. Role of cytochrome P450-dependent monooxygenases and polymorphic variants of GSTT1 and GSTM1 genes in the formation of brain lesions in individuals chronically exposed to mercury. Bull Exp Biol Med. 2013;156(1):15–18. doi:10.1007/s10517-013-2266-2

104. Huang S, Wang HL. Salvianolic acid A improves nerve regeneration and repairs nerve defects in rats with brain injury by downregulating miR-212-3p-mediated SOX7. Kaohsiung J Med Sci. 2023;39(12):1222–1232. doi:10.1002/kjm2.12779

105. Tur C, Ramagopalan S, Altmann DR, Bodini B, Cercignani M, Khaleeli Z, Miller DH, Thompson AJ, Ciccarelli O. HLA-DRB1*15 influences the development of brain tissue damage in early PPMS. Neurology. 2014;83(19):1712–1718. doi:10.1212/WNL.0000000000000959

106. Bortolli APR, Vieira VK, Treco IC, Pascotto CR, Wendt GW, Lucio LC. GSTT1 and GSTM1 polymorphisms with human papillomavirus infection in women from southern Brazil: a case-control study. Mol Biol Rep. 2022;49(7):6467–6474. doi:10.1007/s11033-022-07475-1

107. Greco S, Made’ A, Mutoli M, Zhang L, Piella SN, Vausort M, Lumley AI, Beltrami AP, Srivastava PK, Milani V, et al. HCG18, LEF1AS1 and lncCEACAM21 as biomarkers of disease severity in the peripheral blood mononuclear cells of COVID-19 patients. J Transl Med. 2023;21(1):758. doi:10.1186/s12967-023-04497-6

108. Del Río-Ospina L, Camargo M, Soto-De León SC, Sánchez R, Moreno-Pérez DA, Patarroyo ME, Patarroyo MA. dentifying the HLA DRB1-DQB1 molecules and predicting epitopes associated with high-risk HPV infection clearance and redetection. Sci Rep. 2020;10(1):7306. doi:10.1038/s41598-020-64268-x

109. Hasmann A, Wehrschuetz-Sigl E, Marold A, Wiesbauer H, Schoeftner R, Gewessler U, Kandelbauer A, Schiffer D, Schneider KP, Binder B, et al. Analysis of myeloperoxidase activity in wound fluids as a marker of infection. Ann Clin Biochem. 2013;50(Pt 3):245–254. doi:10.1258/acb.2011.010249

110. Levy R, Le TH. Role of GSTM1 in Hypertension, CKD, and Related Diseases across the Life Span. Kidney360. 2022;3(12):2153–2163. doi:10.34067/KID.0004552022

111. Li QM, Li B, Sun YF, Zhang H, Yang XL, Cai LW, Chen H, Ding YH, Jiang J. Association of HLA-DRB1 single nucleotide polymorphisms with left ventricular remodelling in elderly patients with essential hypertension. J Int Med Res. 2012;40(6):2152–2159. doi:10.1177/030006051204000613

112. Klinke A, Berghausen E, Friedrichs K, Molz S, Lau D, Remane L, Berlin M, Kaltwasser C, Adam M, Mehrkens D, et al. Myeloperoxidase aggravates pulmonary arterial hypertension by activation of vascular Rho-kinase. JCI Insight. 2018;3(11):e97530. doi:10.1172/jci.insight.97530

113. Aliomrani M, Sahraian MA, Shirkhanloo H, Sharifzadeh M, Khoshayand MR, Ghahremani MH. Correlation between heavy metal exposure and GSTM1 polymorphism in Iranian multiple sclerosis patients. Neurol Sci. 2017;38(7):1271–1278. doi:10.1007/s10072-017-2934-5

114. Alcina A, Abad-Grau Mdel M, Fedetz M, Izquierdo G, Lucas M, Fernández O, Ndagire D, Catalá-Rabasa A, Ruiz A, Gayán J, et al. Multiple sclerosis risk variant HLA-DRB1*1501 associates with high expression of DRB1 gene in different human populations. PLoS One. 2012;7(1):e29819. doi:10.1371/journal.pone.0029819

115. Pulli B, Bure L, Wojtkiewicz GR, Iwamoto Y, Ali M, Li D, Schob S, Hsieh KL, Jacobs AH, Chen JW. Multiple sclerosis: myeloperoxidase immunoradiology improves detection of acute and chronic disease in experimental model. Radiology. 2015;275(2):480–489. doi:10.1148/radiol.14141495

116. Rahbar MH, Samms-Vaughan M, Saroukhani S, Bressler J, Hessabi M, Grove ML, Shakspeare-Pellington S, Loveland KA, Beecher C, McLaughlin W. Associations of Metabolic Genes (GSTT1, GSTP1, GSTM1) and Blood Mercury Concentrations Differ in Jamaican Children with and without Autism Spectrum Disorder. Int J Environ Res Public Health. 2021;18(4):1377. doi:10.3390/ijerph18041377

117. Guerini FR, Bolognesi E, Mensi MM, Zanette M, Agliardi C, Zanzottera M, Chiappedi M, Annunziata S, García-García F, Cavallini A, et al. HLA-A, -B, -C and -DRB1 Association with Autism Spectrum Disorder Risk: A Sex-Related Analysis in Italian ASD Children and Their Siblings. Int J Mol Sci. 2024;25(18):9879. doi:10.3390/ijms25189879

118. Ceylan MF, Tural Hesapcioglu S, Yavas CP, Senat A, Erel O. Serum Ischemia-Modified Albumin Levels, Myeloperoxidase Activity and Peripheral Blood Mononuclear cells in Autism Spectrum Disorder (ASD). J Autism Dev Disord. 2021;51(7):2511–2517. doi:10.1007/s10803-020-04740-9

119. Wang R, Wang Y, Wang J, Yang K. Association of glutathione S-transferase T1 and M1 gene polymorphisms with ischemic stroke risk in the Chinese Han population. Neural Regen Res. 2012;7(18):1420–1427. doi:10.3969/j.issn.1673-5374.2012.18.009

120. Murali V, Rathika C, Ramgopal S, Padma Malini R, Arun Kumar MJ, Neethi Arasu V, Jeyaram Illiayaraja K, Balakrishnan K. Susceptible and protective associations of HLA DRB1*/DQB1* alleles and haplotypes with ischaemic stroke. Int J Immunogenet. 2016;43(3):159–165. doi:10.1111/iji.12266

121. Wang Y, Jia Y, Xu Q, Wang R, Sun L, Guo D, Shi M, Yang P, Wang Y, Liu F, et al. Association between myeloperoxidase and the risks of ischemic stroke, heart failure, and atrial fibrillation: A Mendelian randomization study. Nutr Metab Cardiovasc Dis. 2023;33(1):210–218. doi:10.1016/j.numecd.2022.09.027

122. Mohammadynejad P, Saadat I, Ghanizadeh A, Saadat M. Bipolar disorder and polymorphisms of glutathione S-transferases M1 (GSTM1) and T1 (GSTT1). Psychiatry Res. 2011;186(1):144–146. doi:10.1016/j.psychres.2010.06.017

123. Le Clerc S, Lombardi L, Baune BT, Amare AT, Schubert KO, Hou L, Clark SR, Papiol S, Cearns M, Heilbronner U, et al. HLA-DRB1 and HLA-DQB1 genetic diversity modulates response to lithium in bipolar affective disorders. Sci Rep. 2021;11(1):17823. doi:10.1038/s41598-021-97140-7

124. Selek S, Altindag A, Saracoglu G, Aksoy N. Oxidative markers of Myeloperoxidase and Catalase and their diagnostic performance in bipolar disorder. J Affect Disord. 2015;181:92–95. doi:10.1016/j.jad.2015.03.058

125. Peng J, Pan J, Mo J, Peng Y. MPO/HOCl Facilitates Apoptosis and Ferroptosis in the SOD1G93A Motor Neuron of Amyotrophic Lateral Sclerosis. Oxid Med Cell Longev. 2022;2022:8217663. doi:10.1155/2022/8217663

126. Daher J. A Potential Link between Myeloperoxidase Modified LDL, Atherosclerosis and Depression. Int J Mol Sci. 2024;25(16):8805. doi:10.3390/ijms25168805

127. Petersen RB, Nunomura A, Lee HG, Casadesus G, Perry G, Smith MA, Zhu X. Signal transduction cascades associated with oxidative stress in Alzheimer’s disease. J Alzheimers Dis. 2007;11(2):143–152. doi:10.3233/jad-2007-11202

128. Mukhamedyarov MA, Grishin SN, Yusupova ER, Zefirov AL, Palotás A. Alzheimer’s beta-amyloid-induced depolarization of skeletal muscle fibers: implications for motor dysfunctions in dementia. Cell Physiol Biochem. 2009;23(1-3):109–114. doi:10.1159/000204099

129. Majumder P, Roy K, Bagh S, Mukhopadhyay D. Receptor tyrosine kinases (RTKs) consociate in regulatory clusters in Alzheimer’s disease and type 2 diabetes. Mol Cell Biochem. 2019;459(1-2):171–182. doi:10.1007/s11010-019-03560-5

130. Ziliotto N, Bernardi F, Piazza F. Hemostasis components in cerebral amyloid angiopathy and Alzheimer’s disease. Neurol Sci. 2021;42(8):3177–3188. doi:10.1007/s10072-021-05327-7

131. Jia L, Piña-Crespo J, Li Y. Restoring Wnt/β-catenin signaling is a promising therapeutic strategy for Alzheimer’s disease. Mol Brain. 2019;12(1):104. doi:10.1186/s13041-019-0525-5

132. Anwar MM, Özkan E, Gürsoy-Özdemir Y. The role of extracellular matrix alterations in mediating astrocyte damage and pericyte dysfunction in Alzheimer’s disease: A comprehensive review. Eur J Neurosci. 2022;56(9):5453–5475. doi:10.1111/ejn.15372

133. Bordier P, Garrigue S, Lanusse S, Margaine J, Robert F, Gencel L, Lafitte A. Cardiovascular effects and risk of syncope related to donepezil in patients with Alzheimer’s disease. CNS Drugs. 2006;20(5):411–417. doi:10.2165/00023210-200620050-00005

134. Van Beek AH, Claassen JA. The cerebrovascular role of the cholinergic neural system in Alzheimer’s disease. Behav Brain Res. 2011;221(2):537–542. doi:10.1016/j.bbr.2009.12.047

135. Guo L, Tian J, Du H. Mitochondrial Dysfunction and Synaptic Transmission Failure in Alzheimer’s Disease. J Alzheimers Dis. 2017;57(4):1071–1086. doi:10.3233/JAD-160702

136. Liu R, Collier JM, Abdul-Rahman NH, Capuk O, Zhang Z, Begum G. Dysregulation of Ion Channels and Transporters and Blood-Brain Barrier Dysfunction in Alzheimer’s Disease and Vascular Dementia. Aging Dis. 2024;15(4):1748–1770.. doi:10.14336/AD.2023.1201

137. Cui H, Freeman C, Jacobson GA, Small DH. Proteoglycans in the central nervous system: role in development, neural repair, and Alzheimer’s disease. IUBMB Life. 2013;65(2):108–120. doi:10.1002/iub.1118

138. Douros A, Santella C, Dell’Aniello S, Azoulay L, Renoux C, Suissa S, Brassard P. Infectious Disease Burden and the Risk of Alzheimer’s Disease: A Population-Based Study. J Alzheimers Dis. 2021;81(1):329–338. doi:10.3233/JAD-201534

139. Astillero-Lopez V, Villar-Conde S, Gonzalez-Rodriguez M, Flores-Cuadrado A, Ubeda-Banon I, Saiz-Sanchez D, Martinez-Marcos A. Proteomic analysis identifies HSP90AA1, PTK2B, and ANXA2 in the human entorhinal cortex in Alzheimer’s disease: Potential role in synaptic homeostasis and Aβ pathology through microglial and astroglial cells. Brain Pathol. 2024;34(4):e13235. doi:10.1111/bpa.13235

140. Bhattarai P, Gunasekaran TI, Belloy ME, Reyes-Dumeyer D, Jülich D, Tayran H, Yilmaz E, Flaherty D, Turgutalp B, Sukumar G, Alba C, et al. Rare genetic variation in fibronectin 1 (FN1) protects against APOEε4 in Alzheimer’s disease. Acta Neuropathol. 2024;147(1):70. doi:10.1007/s00401-024-02721-1

141. Xu M, Zhang DF, Luo R, Wu Y, Zhou H, Kong LL, Bi R, Yao YG. A systematic integrated analysis of brain expression profiles reveals YAP1 and other prioritized hub genes as important upstream regulators in Alzheimer’s disease. Alzheimers Dement. 2018;14(2):215–229. doi:10.1016/j.jalz.2017.08.012

142. Sanfilippo C, Castrogiovanni P, Imbesi R, Di Rosa M. CHI3L2 Expression Levels Are Correlated with AIF1, PECAM1, and CALB1 in the Brains of Alzheimer’s Disease Patients. J Mol Neurosci. 2020;70(10):1598–1610. doi:10.1007/s12031-020-01667-9

143. Ma X, Zhang Y, Gou D, Ma J, Du J, Wang C, Li S, Cui H. Metabolic Reprogramming of Microglia Enhances Proinflammatory Cytokine Release through EphA2/p38 MAPK Pathway in Alzheimer’s Disease. J Alzheimers Dis. 2022;88(2):771–785. doi:10.3233/JAD-220227

144. Zhao K, Liu J, Sun T, Zeng L, Cai Z, Li Z, Liu R. The miR-25802/KLF4/NF-κB signaling axis regulates microglia-mediated neuroinflammation in Alzheimer’s disease. Brain Behav Immun. 2024;118:31–48. doi:10.1016/j.bbi.2024.02.016

145. Dong Y, Li T, Ma Z, Zhou C, Wang X, Li J. HSPA1A, HSPA2, and HSPA8 Are Potential Molecular Biomarkers for Prognosis among HSP70 Family in Alzheimer’s Disease. Dis Markers. 2022;2022:9480398. doi:10.1155/2022/9480398

146. He R, Cheng J, Qiu Y, Hu Y, Liu J, Wang TH, Cao X. IGF1R and FLT1 in female endothelial cells and CHD2 in male microglia play important roles in Alzheimer’s disease based on gender difference analysis. Exp Gerontol. 2024;194:112512. doi:10.1016/j.exger.2024.112512

147. Mahzarnia A, Lutz MW, Badea A. A Continuous Extension of Gene Set Enrichment Analysis Using the Likelihood Ratio Test Statistics Identifies Vascular Endothelial Growth Factor as a Candidate Pathway for Alzheimer’s Disease via ITGA5. J Alzheimers Dis. 2024;97(2):635–648. doi:10.3233/JAD-230934

148. Ye L, Zhao J, Xiao Z, Gu W, Liu X, Ajuyo NMC, Min Y, Pei Y, Wang D. Integrative Human Genetic and Cellular Analysis of the Pathophysiological Roles of AnxA2 in Alzheimer’s Disease. Antioxidants (Basel). 2024;13(10):1274. doi:10.3390/antiox13101274

149. Zhang X, Zhu C, Beecham G, Vardarajan BN, Ma Y, Lancour D, Farrell JJ, Chung J; Alzheimer’s Disease Sequencing Project; Mayeux R, et al. A rare missense variant of CASP7 is associated with familial late-onset Alzheimer’s disease. Alzheimers Dement. 2019;15(3):441–452. doi:10.1016/j.jalz.2018.10.005

150. Govindarajulu M, Ramesh S, Beasley M, Lynn G, Wallace C, Labeau S, Pathak S, Nadar R, Moore T, Dhanasekaran M. Role of cGAS-Sting Signaling in Alzheimer’s Disease. Int J Mol Sci. 2023;24(9):8151. doi:10.3390/ijms24098151

151. Ocenasova A, Shawkatova I, Javor J, Parnicka Z, Minarik G, Kralova M, Kiralyova I, Mikolaskova I, Durmanova V. MMP2 rs243866 and rs2285053 Polymorphisms and Alzheimer’s Disease Risk in Slovak Caucasian Population. Life (Basel). 2023;13(4):882. doi:10.3390/life13040882

152. Peress NS, Perillo E. Differential expression of TGF-beta 1, 2 and 3 isotypes in Alzheimer’s disease: a comparative immunohistochemical study with cerebral infarction, aged human and mouse control brains. J Neuropathol Exp Neurol. 1995;54(6):802–811. doi:10.1097/00005072-199511000-00007

153. Jafarian Z, Saliminejad K, Kamali K, Ohadi M, Kowsari A, Nasehi L, Khorram Khorshid HR. Association of glutathione S-transferases M1, P1 and T1 variations and risk of late-onset Alzheimer’s disease. Neurol Res. 2018;40(1):41–44. doi:10.1080/01616412.2017.1390902

154. Arfaei R, Mikaeili N, Daj F, Boroumand A, Kheyri A, Yaraghi P, Shirzad Z, Keshavarz M, Hassanshahi G, Jafarzadeh A, et al. Decoding the role of the CCL2/CCR2 axis in Alzheimer’s disease and innovating therapeutic approaches: Keeping All options open. Int Immunopharmacol. 2024;135:112328. doi:10.1016/j.intimp.2024.112328

155. Wang L, Zhou BQ, Li YH, Jiang QQ, Cong WH, Chen KJ, Wen XM, Wu ZZ. Lactoferrin modification of berberine nanoliposomes enhances the neuroprotective effects in a mouse model of Alzheimer’s disease. Neural Regen Res. 2023;18(1):226–232. doi:10.4103/1673-5374.344841

156. Lee JH, Gurney S, Pang D, Temkin A, Park N, Janicki SC, Zigman WB, Silverman W, Tycko B, Schupf N. Polymorphisms in HSD17B1: Early Onset and Increased Risk of Alzheimer’s Disease in Women with Down Syndrome. Curr Gerontol Geriatr Res. 2012;2012:361218. doi:10.1155/2012/361218

157. Shigemizu D, Fukunaga K, Yamakawa A, Suganuma M, Fujita K, Kimura T, Watanabe K, Mushiroda T, Sakurai T, Niida S, et al. The HLA-DRB1*09:01-DQB1*03:03 haplotype is associated with the risk for late-onset Alzheimer’s disease in APOE [Formula: see text]4-negative Japanese adults. NPJ Aging. 2024;10(1):3. doi:10.1038/s41514-023-00131-3

158. King A, Szekely B, Calapkulu E, Ali H, Rios F, Jones S, Troakes C. The Increased Densities, But Different Distributions, of Both C3 and S100A10 Immunopositive Astrocyte-Like Cells in Alzheimer’s Disease Brains Suggest Possible Roles for Both A1 and A2 Astrocytes in the Disease Pathogenesis. Brain Sci. 2020;10(8):503. doi:10.3390/brainsci10080503

159. Al Rahim M, Yoon Y, Dimovasili C, Shao Z, Huang Q, Zhang E, Kezunovic N, Chen L, Schaffner A, Huntley GW, et al. Presenilin1 familial Alzheimer disease mutants inactivate EFNB1- and BDNF-dependent neuroprotection against excitotoxicity by affecting neuroprotective complexes of N-methyl-d-aspartate receptor. Brain Commun. 2020;2(2):fcaa100. doi:10.1093/braincomms/fcaa100

160. Adams SL, Benayoun L, Tilton K, Mellott TJ, Seshadri S, Blusztajn JK, Delalle I.Immunohistochemical Analysis of Activin Receptor-Like Kinase 1 (ACVRL1/ALK1) Expression in the Rat and Human Hippocampus: Decline in CA3 During Progression of Alzheimer’s Disease. J Alzheimers Dis. 2018;63(4):1433–1443. doi:10.3233/JAD-171065

161. Sun L, Guo C, Song Y, Sheng J, Xiao S; Alzheimer’s Disease Neuroimaging Initiative. Blood BMP6 Associated with Cognitive Performance and Alzheimer’s Disease Diagnosis: A Longitudinal Study of Elders. J Alzheimers Dis. 2022;88(2):641–651. doi:10.3233/JAD-220279

162. Eftekharzadeh M, Simorgh S, Doshmanziari M, Hassanzadeh L, Shariatpanahi M. Human adipose-derived stem cells reduce receptor-interacting protein 1, receptor-interacting protein 3, and mixed lineage kinase domain-like pseudokinase as necroptotic markers in rat model of Alzheimer’s disease. Indian J Pharmacol. 2020;52(5):392–401. doi:10.4103/ijp.IJP_545_19

163. Siokas V, Mouliou DS, Liampas I, Aloizou AM, Folia V, Zoupa E, Papadimitriou A, Lavdas E, Bogdanos DP, Dardiotis E. Analysis of ADORA2A rs5760423 and CYP1A2 rs762551 Genetic Variants in Patients with Alzheimer’s Disease. Int J Mol Sci. 2022;23(22):14400. doi:10.3390/ijms232214400

164. Sita G, Hrelia P, Tarozzi A, Morroni F. P-glycoprotein (ABCB1) and Oxidative Stress: Focus on Alzheimer’s Disease. Oxid Med Cell Longev. 2017;2017:7905486. doi:10.1155/2017/7905486

165. Cheng J, Liu HP, Lee CC, Chen MY, Lin WY, Tsai FJ. Matrix metalloproteinase 14 modulates diabetes and Alzheimer’s disease cross-talk: a meta-analysis. Neurol Sci. 2018;39(2):267–274. doi:10.1007/s10072-017-3166-4

166. Merchant JP, Zhu K, Henrion MYR, Zaidi SSA, Lau B, Moein S, Alamprese ML, Pearse RV 2nd Bennett DA, Ertekin-Taner N, et al. Predictive network analysis identifies JMJD6 and other potential key drivers in Alzheimer’s disease. Commun Biol. 2023;6(1):503. doi:10.1038/s42003-023-04791-5

167. Pons V, Lévesque P, Plante MM, Rivest S. Conditional genetic deletion of CSF1 receptor in microglia ameliorates the physiopathology of Alzheimer’s disease. Alzheimers Res Ther. 2021;13(1):8. doi:10.1186/s13195-020-00747-7

168. Yang HS, Yau WW, Carlyle BC, Trombetta BA, Zhang C, Shirzadi Z, Schultz AP, Pruzin JJ, Fitzpatrick CD, Kirn DR, et al. Plasma VEGFA and PGF impact longitudinal tau and cognition in preclinical Alzheimer’s disease. Brain. 2024;147(6):2158–2168. doi:10.1093/brain/awae034

169. Wang T, Zhou YQ, Wang Y, Zhang L, Zhu X, Wang XY, Wang JH, Han LK, Meng J, Zhang X, et al. Long-term potentiation-based screening identifies neuronal PYGM as a synaptic plasticity regulator participating in Alzheimer’s disease. Zool Res. 2023;44(5):867–881. doi:10.24272/j.issn.2095-8137.2023.123

170. Matchett BJ, Lincoln SJ, Baker M, Tamvaka N, Labuzan SA, Hicks Sirmans TN, Moloney CM, Helminger J, Hinkle KM, Cabrera-Rodriguez J, et al. The SERPINA5 coding variant E228Q does not contribute to clinicopathologic characteristics in Alzheimer’s disease: A cross-sectional study. Medicine (Baltimore). 2023;102(24):e34017. doi:10.1097/MD.0000000000034017

171. Rudan Njavro J, Klotz J, Dislich B, Wanngren J, Shmueli MD, Herber J, Kuhn PH, Kumar R, Koeglsperger T, Conrad M, et al. Mouse brain proteomics establishes MDGA1 and CACHD1 as in vivo substrates of the Alzheimer protease BACE1. FASEB J. 2020;34(2):2465–2482. doi:10.1096/fj.201902347R

172. Tosto G, Fu H, Vardarajan BN, Lee JH, Cheng R, Reyes-Dumeyer D, Lantigua R, Medrano M, Jimenez-Velazquez IZ, Elkind MS, et al. F-box/LRR-repeat protein 7 is genetically associated with Alzheimer’s disease. Ann Clin Transl Neurol. 2015;2(8):810–820. doi:10.1002/acn3.223

173. Beckmann ND, Lin WJ, Wang M, Cohain AT, Charney AW, Wang P, Ma W, Wang YC, Jiang C, Audrain M, et al. Multiscale causal networks identify VGF as a key regulator of Alzheimer’s disease. Nat Commun. 2020;11(1):3942. doi:10.1038/s41467-020-17405-z

174. Franzmeier N, Dehsarvi A, Steward A, Biel D, Dewenter A, Roemer SN, Wagner F, Groß M, Brendel M, Moscoso A, et al. Elevated CSF GAP-43 is associated with accelerated tau accumulation and spread in Alzheimer’s disease. Nat Commun. 2024;15(1):202. doi:10.1038/s41467-023-44374-w

175. Doert A, Pilatus U, Zanella F, Müller WE, Eckert GP. ¹H- and ¹³C-NMR spectroscopy of Thy-1-APPSL mice brain extracts indicates metabolic changes in Alzheimer’s disease. J Neural Transm (Vienna). 2015;122(4):541–550. doi:10.1007/s00702-015-1387-3

176. Agra Almeida Quadros AR, Li Z, Wang X, Ndayambaje IS, Aryal S, Ramesh N, Nolan M, Jayakumar R, Han Y, Stillman H, et al. Cryptic splicing of stathmin-2 and UNC13A mRNAs is a pathological hallmark of TDP-43-associated Alzheimer’s disease. Acta Neuropathol. 2024;147(1):9. doi:10.1007/s00401-023-02655-0

177. Dauar MT, Picard C, Labonté A, Breitner J, Rosa-Neto P, Villeneuve S, Poirier J; PREVENT-AD Research Group. Contactin 5 and Apolipoproteins Interplay in Alzheimer’s Disease. J Alzheimers Dis. 2024;98(4):1361–1375. doi:10.3233/JAD-231003

178. Satoh J, Yamamoto Y, Asahina N, Kitano S, Kino Y. RNA-Seq data mining: downregulation of NeuroD6 serves as a possible biomarker for alzheimer’s disease brains. Dis Markers. 2014;2014:123165. doi:10.1155/2014/123165

179. Emilsson L, Saetre P, Jazin E. Low mRNA levels of RGS4 splice variants in Alzheimer’s disease: association between a rare haplotype and decreased mRNA expression. Synapse. 2006;59(3):173–176. doi:10.1002/syn.20226

180. Lee JH, Koh SQ, Guadagna S, Francis PT, Esiri MM, Chen CP, Wong PT, Dawe GS, Lai MK. Altered relaxin family receptors RXFP1 and RXFP3 in the neocortex of depressed Alzheimer’s disease patients. Psychopharmacology (Berl). 2016;233(4):591–598. doi:10.1007/s00213-015-4131-7

181. Shi Z, Zhang K, Zhou H, Jiang L, Xie B, Wang R, Xia W, Yin Y, Gao Z, Cui D, et al. Increased miR-34c mediates synaptic deficits by targeting synaptotagmin 1 through ROS-JNK-p53 pathway in Alzheimer’s Disease. Aging Cell. 2020;19(3):e13125. doi:10.1111/acel.13125

182. Wang Q, Tao S, Xing L, Liu J, Xu C, Xu X, Ding H, Shen Q, Yu X, Zheng Y. SNAP25 is a potential target for early stage Alzheimer’s disease and Parkinson’s disease. Eur J Med Res. 2023;28(1):570. doi:10.1186/s40001-023-01360-8

183. Ruzha Y, Ni J, Quan Z, Li H, Qing H. Role of Vitronectin and Its Receptors in Neuronal Function and Neurodegenerative Diseases. Int J Mol Sci. 2022;23(20):12387. doi:10.3390/ijms232012387

184. Wang L, Jing R, Wang X, Wang B, Guo K, Zhao J, Gao S, Xu N, Xuan X. A method for the expression of fibroblast growth factor 14 and assessment of its neuroprotective effect in an Alzheimer’s disease model. Ann Transl Med. 2021;9(12):994. doi:10.21037/atm-21-2492

185. Ren J, Zhang S, Wang X, Deng Y, Zhao Y, Xiao Y, Liu J, Chu L, Qi X. MEF2C ameliorates learning, memory, and molecular pathological changes in Alzheimer’s disease in vivo and in vitro. Acta Biochim Biophys Sin (Shanghai). 2022;54(1):77–90. doi:10.3724/abbs.2021012

186. Hurst C, Pugh DA, Abreha MH, Duong DM, Dammer EB, Bennett DA, Herskowitz JH, Seyfried NT. Integrated Proteomics to Understand the Role of Neuritin (NRN1) as a Mediator of Cognitive Resilience to Alzheimer’s Disease. Mol Cell Proteomics. 2023;22(5):100542. doi:10.1016/j.mcpro.2023.100542

187. Wang J, Zhou C, Huang Z, et al. Repetitive Transcranial Magnetic Stimulation-Mediated Neuroprotection in the 5xFAD Mouse Model of Alzheimer’s Disease Through GABRG2 and SNAP25 Modulation. Mol Neurobiol. 2024. doi:10.1007/s12035-024-04354-7

188. Datta D, Perone I, Morozov YM, Arellano J, Duque A, Rakic P, van Dyck CH, Arnsten AFT. Localization of PDE4D, HCN1 channels, and mGluR3 in rhesus macaque entorhinal cortex may confer vulnerability in Alzheimer’s disease. Cereb Cortex. 2023;33(24):11501–11516. doi:10.1093/cercor/bhad382

189. Hu D, Dong X, Wang Q, Liu M, Luo S, Meng Z, Feng Z, Zhou W, Song W. PCP4 Promotes Alzheimer’s Disease Pathogenesis by Affecting Amyloid-β Protein Precursor Processing. J Alzheimers Dis. 2023;94(2):737–750. doi:10.3233/JAD-230192

190. Nguyen DPQ, Pham S, Jallow AW, Ho NT, Le B, Quang HT, Lin YF, Lin YF. Multiple Transcriptomic Analyses Explore Potential Synaptic Biomarker Rabphilin-3A for Alzheimer’s Disease. Sci Rep. 2024;14(1):18717. doi:10.1038/s41598-024-66693-8

191. Sheu JJ, Yang LY, Sanotra MR, Wang ST, Lu HT, Kam RSY, Hsu IU, Kao SH, Lee CK, Shieh JC, et al. Reduction of AHI1 in the serum of Taiwanese with probable Alzheimer’s disease. Clin Biochem. 2020;76:24–30. doi:10.1016/j.clinbiochem.2019.11.011

192. Zhang N, Sui Y, Jendrichovsky P, Feng H, Shi H, Zhang X, Xu S, Sun W, Zhang H, Chen X, et al. Cholecystokinin B receptor agonists alleviates anterograde amnesia in cholecystokinin-deficient and aged Alzheimer’s disease mice. Alzheimers Res Ther. 2024;16(1):109. doi:10.1186/s13195-024-01472-1

193. van Abel D, Michel O, Veerhuis R, Jacobs M, van Dijk M, Oudejans CB. Direct downregulation of CNTNAP2 by STOX1A is associated with Alzheimer’s disease. J Alzheimers Dis. 2012;31(4):793–800. doi:10.3233/JAD-2012-120472

194. Berrocal M, Saez L, Mata AM. Sorcin Activates the Brain PMCA and Blocks the Inhibitory Effects of Molecular Markers of Alzheimer’s Disease on the Pump Activity. Int J Mol Sci. 2021;22(11):6055. doi:10.3390/ijms22116055

195. Lehrer S, Rheinstein PH. RORB, an Alzheimer’s disease susceptibility gene, is associated with viral encephalitis, an Alzheimer’s disease risk factor. Clin Neurol Neurosurg. 2023;233:107984. doi:10.1016/j.clineuro.2023.107984

196. McPhie DL, Coopersmith R, Hines-Peralta A, Chen Y, Ivins KJ, Manly SP, Kozlowski MR, Neve KA, Neve RL. DNA synthesis and neuronal apoptosis caused by familial Alzheimer disease mutants of the amyloid precursor protein are mediated by the p21 activated kinase PAK3. J Neurosci. 2003;23(17):6914–6927. doi:10.1523/JNEUROSCI.23-17-06914.2003

197. Good PF, Alapat D, Hsu A, Chu C, Perl D, Wen X, Burstein DE, Kohtz DS. A role for semaphorin 3A signaling in the degeneration of hippocampal neurons during Alzheimer’s disease. J Neurochem. 2004;91(3):716–736. doi:10.1111/j.1471-4159.2004.02766.x

198. Tramutola A, Di Domenico F, Barone E, Perluigi M, Butterfield DA. It Is All about (U)biquitin: Role of Altered Ubiquitin-Proteasome System and UCHL1 in Alzheimer Disease. Oxid Med Cell Longev. 2016;2016:2756068. doi:10.1155/2016/2756068

199. van der Linden RJ, Gerritsen JS, Liao M, Widomska J, Pearse RV 2nd, White FM, Franke B, Young-Pearse TL, Poelmans G. RNA-binding protein ELAVL4/HuD ameliorates Alzheimer’s disease-related molecular changes in human iPSC-derived neurons. Prog Neurobiol. 2022;217:102316. doi:10.1016/j.pneurobio.2022.102316

200. Hu YT, Chen XL, Huang SH, Zhu QB, Yu SY, Shen Y, Sluiter A, Verhaagen J, Zhao J, Swaab D, et al. Early growth response-1 regulates acetylcholinesterase and its relation with the course of Alzheimer’s disease. Brain Pathol. 2019;29(4):502–512. doi:10.1111/bpa.12688

201. Sanfilippo C, Giuliano L, Castrogiovanni P, Imbesi R, Ulivieri M, Fazio F, Blennow K, Zetterberg H, Di Rosa M. Sex, Age, and Regional Differences in CHRM1 and CHRM3 Genes Expression Levels in the Human Brain Biopsies: Potential Targets for Alzheimer’s Disease-related Sleep Disturbances. Curr Neuropharmacol. 2023;21(3):740–760. doi:10.2174/1570159X21666221207091209

202. D’Andrea MR, Nagele RG. MAP-2 immunolabeling can distinguish diffuse from dense-core amyloid plaques in brains with Alzheimer’s disease. Biotech Histochem. 2002;77(2):95–103.

203. Henderson MX, Sengupta M, Trojanowski JQ, Lee VMY. Alzheimer’s disease tau is a prominent pathology in LRRK2 Parkinson’s disease. Acta Neuropathol Commun. 2019;7(1):183. doi:10.1186/s40478-019-0836-x

204. Cai B, Shao N, Ye T, Zhou P, Si W, Song H, Wang G, Kou J. Phosphorylation of MAP 1A regulates hyperphosphorylation of Tau in Alzheimer’s disease model. Neuropathol Appl Neurobiol. 2023;49(5):e12934. doi:10.1111/nan.12934

205. Yao J, Chen SRW. RyR2-dependent modulation of neuronal hyperactivity: A potential therapeutic target for treating Alzheimer’s disease. J Physiol. 2024;602(8):1509–1518. doi:10.1113/JP283824

206. Martínez-Drudis L, Bérard M, Musiol D, Rivest S, Oueslati A. Pharmacological inhibition of PLK2 kinase activity mitigates cognitive decline but aggravates APP pathology in a sex-dependent manner in APP/PS1 mouse model of Alzheimer’s disease. Heliyon. 2024;10(20):e39571. doi:10.1016/j.heliyon.2024.e39571

207. Shu L, Sun Q, Zhang Y, Xu Q, Guo J, Yan X, Tang B. The Association between C9orf72 Repeats and Risk of Alzheimer’s Disease and Amyotrophic Lateral Sclerosis: A Meta-Analysis. Parkinsons Dis. 2016;2016:5731734. doi:10.1155/2016/5731734

208. Sevlever D, Zou F, Ma L, Carrasquillo S, Crump MG, Culley OJ, Hunter TA, Bisceglio GD, Younkin L, Allen M, et al. Genetically-controlled Vesicle-Associated Membrane Protein 1 expression may contribute to Alzheimer’s pathophysiology and susceptibility. Mol Neurodegener. 2015;10:18. doi:10.1186/s13024-015-0015-x

209. Almeida VN. Somatostatin and the pathophysiology of Alzheimer’s disease. Ageing Res Rev. 2024;96:102270. doi:10.1016/j.arr.2024.102270

210. Kumar P, Goettemoeller AM, Espinosa-Garcia C, Tobin BR, Tfaily A, Nelson RS, Natu A, Dammer EB, Santiago JV, Malepati S, et al. Native-state proteomics of Parvalbumin interneurons identifies unique molecular signatures and vulnerabilities to early Alzheimer’s pathology. Nat Commun. 2024;15(1):2823. doi:10.1038/s41467-024-47028-7

211. Zhu M, Tang M, Du Y. Identification of TAC1 Associated with Alzheimer’s Disease Using a Robust Rank Aggregation Approach. J Alzheimers Dis. 2023;91(4):1339–1349. doi:10.3233/JAD-220950

212. Seddighi S, Varma VR, An Y, Varma S, Beason-Held LL, Tanaka T, Kitner-Triolo MH, Kraut MA, Davatzikos C, Thambisetty M. SPARCL1 Accelerates Symptom Onset in Alzheimer’s Disease and Influences Brain Structure and Function During Aging. J Alzheimers Dis. 2018;61(1):401–414. doi:10.3233/JAD-170557

213. Ouellette AR, Neuner SM, Dumitrescu L, Anderson LC, Gatti DM, Mahoney ER, Bubier JA, Churchill G, Peters L, Huentelman MJ, et al. Cross-Species Analyses Identify Dlgap2 as a Regulator of Age-Related Cognitive Decline and Alzheimer’s Dementia. Cell Rep. 2020;32(9):108091. doi:10.1016/j.celrep.2020.108091

214. Di Maio A, De Rosa A, Pelucchi S, Garofalo M, Marciano B, Nuzzo T, Gardoni F, Isidori AM, Di Luca M, Errico F, et al. Analysis of mRNA and Protein Levels of CAP2, DLG1 and ADAM10 Genes in Post-Mortem Brain of Schizophrenia, Parkinson’s and Alzheimer’s Disease Patients. Int J Mol Sci. 2022;23(3):1539. doi:10.3390/ijms23031539

215. Detrait E, Maurice T, Hanon E, Leclercq K, Lamberty Y. Lack of synaptic vesicle protein SV2B protects against amyloid-β -induced oxidative stress, cholinergic deficit and cognitive impairment in mice. Behav Brain Res. 2014;271:277–285. doi:10.1016/j.bbr.2014.06.013

216. Deng Y, Bi M, Delerue F, Forrest SL, Chan G, van der Hoven J, van Hummel A, Feiten AF, Lee S, Martinez-Valbuena I, et al. Loss of LAMP5 interneurons drives neuronal network dysfunction in Alzheimer’s disease. Acta Neuropathol. 2022;144(4):637–650. doi:10.1007/s00401-022-02457-w

217. Davidsson P, Bogdanovic N, Lannfelt L, Blennow K. Reduced expression of amyloid precursor protein, presenilin-1 and rab3a in cortical brain regions in Alzheimer’s disease. Dement Geriatr Cogn Disord. 2001;12(4):243–250. doi:10.1159/000051266

218. Lin CW, Chang LC, Tseng GC, Kirkwood CM, Sibille EL, Sweet RA. VSNL1 Co-Expression Networks in Aging Include Calcium Signaling, Synaptic Plasticity, and Alzheimer’s Disease Pathways. Front Psychiatry. 2015;6:30. doi:10.3389/fpsyt.2015.00030

219. Zhuang X, Xia Y, Liu Y, Guo T, Xia Z, Wang Z, Zhang G. SCG5 and MITF may be novel markers of copper metabolism immunorelevance in Alzheimer’s disease. Sci Rep. 2024;14(1):13619. doi:10.1038/s41598-024-64599-z

220. Soosanabadi M, Bayat H, Kamali K, Saliminejad K, Banan M, Khorram Khorshid HR. Association Study of IL-4 -590 C/T and DDX39B - 22 G/C Polymorphisms with the Risk of Late-Onset Alzheimer’s Disease in Iranian Population. Curr Aging Sci. 2015;8(3):276–281. doi:10.2174/187460980803151027125919

221. Zhu X, Rottkamp CA, Raina AK, Brewer GJ, Ghanbari HA, Boux H, Smith MA. Neuronal CDK7 in hippocampus is related to aging and Alzheimer disease. Neurobiol Aging. 2000;21(6):807–813. doi:10.1016/s0197-4580(00)00217-7

222. Zhou X, Wu X, Wang R, Han L, Li H, Zhao W. Mechanisms of 3-Hydroxyl 3-Methylglutaryl CoA Reductase in Alzheimer’s Disease. Int J Mol Sci. 2023;25(1):170. doi:10.3390/ijms25010170

223. Fu Y, Hsiao JH, Paxinos G, Halliday GM, Kim WS. ABCA5 regulates amyloid-β peptide production and is associated with Alzheimer’s disease neuropathology. J Alzheimers Dis. 2015;43(3):857–869. doi:10.3233/JAD-141320

224. Wang J, Chen L, Wang Z, Zhang S, Ding D, Lin G, Zhang H, Boda VK, Kong D, Ortyl TC, et al. TRPC3 suppression ameliorates synaptic dysfunctions and memory deficits in Alzheimer’s disease. Preprint. bioRxiv. 2024;2024.09.16.611061. doi:10.1101/2024.09.16.611061

225. Dulewicz M, Kulczyńska-Przybik A, Słowik A, Borawska R, Mroczko B. Fatty Acid Binding Protein 3 (FABP3) and Apolipoprotein E4 (ApoE4) as Lipid Metabolism-Related Biomarkers of Alzheimer’s Disease. J Clin Med. 2021;10(14):3009. doi:10.3390/jcm10143009

226. Zuo Y, Dang R, Peng H, Hu P, Yang Y. LL37-mtDNA regulates viability, apoptosis, inflammation, and autophagy in lipopolysaccharide-treated RLE-6TN cells by targeting Hsp90aa1. Open Life Sci. 2024;19(1):20220943. doi:10.1515/biol-2022-0943

227. Yu W, Ye H, Li Y, Bao X, Ni Y, Chen X, Sun Y, Chen A, Zhou W, Li J. Pneumocystis carinii infection drives upregulation of Fn1 expression that causes pulmonary fibrosis with an inflammatory response. Rev Iberoam Micol. 2024;41(1):17–26. doi:10.1016/j.riam.2024.04.002

228. Xu M, Li XY, Song L, Tao C, Fang J, Tao L. miR-484 targeting of Yap1-induced LPS-inhibited proliferation, and promoted apoptosis and inflammation in cardiomyocyte. Biosci Biotechnol Biochem. 2021;85(2):378–385. doi:10.1093/bbb/zbaa009

229. Qin WD, Mi SH, Li C, Wang GX, Zhang JN, Wang H, Zhang F, Ma Y, Wu DW, Zhang M. Low shear stress induced HMGB1 translocation and release via PECAM-1/PARP-1 pathway to induce inflammation response. PLoS One. 2015;10(3):e0120586. doi:10.1371/journal.pone.0120586

230. Mueller M, Zwinger L, Klaassen S, Poller W, Monserrat Iglesias L, Pablo Ochoa J, Klingel K, Landmesser U, Heidecker B. Severe heart failure in the setting of inflammatory cardiomyopathy with likely pathogenic titin variant. Int J Cardiol Heart Vasc. 2022;39:100969. doi:10.1016/j.ijcha.2022.100969

231. Stolfi C, Troncone E, Marafini I, Monteleone G. Role of TGF-Beta and Smad7 in Gut Inflammation, Fibrosis and Cancer. Biomolecules. 2020;11(1):17. doi:10.3390/biom11010017

232. Zhao P, Sun L, Zhao C. TCF1/LEF1 triggers Wnt-dependent chemokine/cytokine-induced inflammation and cadherin pathways to drive T-ALL cell migration. Biochem Biophys Rep. 2023;34:101457. doi:10.1016/j.bbrep.2023.101457

233. Park CJ, Lin PC, Zhou S, Barakat R, Bashir ST, Choi JM, Cacioppo JA, Oakley OR, Duffy DM, Lydon JP, et al. Progesterone Receptor Serves the Ovary as a Trigger of Ovulation and a Terminator of Inflammation. Cell Rep. 2020;31(2):107496. doi:10.1016/j.celrep.2020.03.060

234. Overstreet AC, Burge M, Bellar A, McMullen M, Czarnecki D, Huang E, Pathak V, Finney C, Vij R, Dasarathy S, et al. Evidence that extracellular HSPB1 contributes to inflammation in alcohol-associated hepatitis. Preprint. medRxiv. 2024;2024.09.06.24313193. doi:10.1101/2024.09.06.24313193

235. Yoo SA, Yoon HJ, Kim HS, Chae CB, De Falco S, Cho CS, Kim WU. Role of placenta growth factor and its receptor flt-1 in rheumatoid inflammation: a link between angiogenesis and inflammation. Arthritis Rheum. 2009;60(2):345–354. doi:10.1002/art.24289

236. Chawla M, Mukherjee T, Deka A, Chatterjee B, Sarkar UA, Singh AK, Kedia S, Lum J, Dhillon MK, Banoth B, et al. An epithelial Nfkb2 pathway exacerbates intestinal inflammation by supplementing latent RelA dimers to the canonical NF-κB module. Proc Natl Acad Sci U S A. 2021;118(25):e2024828118. doi:10.1073/pnas.2024828118

237. Liu KL, Yang YC, Yao HT, Chia TW, Lu CY, Li CC, Tsai HJ, Lii CK, Chen HW. Docosahexaenoic acid inhibits inflammation via free fatty acid receptor FFA4, disruption of TAB2 interaction with TAK1/TAB1 and downregulation of ERK-dependent Egr-1 expression in EA.hy926 cells. Mol Nutr Food Res. 2016;60(2):430–443. doi:10.1002/mnfr.201500178

238. Knyazev E, Maltseva D, Raygorodskaya M, Shkurnikov M. HIF-Dependent NFATC1 Activation Upregulates ITGA5 and PLAUR in Intestinal Epithelium in Inflammatory Bowel Disease. Front Genet. 2021;12:791640. doi:10.3389/fgene.2021.791640

239. Larson KC, Lipko M, Dabrowski M, Draper MP. Gng12 is a novel negative regulator of LPS-induced inflammation in the microglial cell line BV-2. Inflamm Res. 2010;59(1):15–22. doi:10.1007/s00011-009-0062-2

240. Fu W, Hu W, Shi L, Mundra JJ, Xiao G, Dustin ML, Liu CJ. Foxo4- and Stat3-dependent IL-10 production by progranulin in regulatory T cells restrains inflammatory arthritis. FASEB J. 2017;31(4):1354–1367. doi:10.1096/fj.201601134R

241. Han W, Allam SA, Elsawa SF. GLI2-Mediated Inflammation in the Tumor Microenvironment. Adv Exp Med Biol. 2020;1263:55–65. doi:10.1007/978-3-030-44518-8_5

242. Baba H, Kimura N, Kanegane H, Miya F, Kosaki K, Morio T, Koike R. GATA2 deficiency of a novel missense variant with multiorgan inflammation. Rheumatology (Oxford). 2024;63(8):e226–e228. doi:10.1093/rheumatology/keae062

243. Chen J, Liu Y, Xia S, Ye X, Chen L. Annexin A2 (ANXA2) regulates the transcription and alternative splicing of inflammatory genes in renal tubular epithelial cells. BMC Genomics. 2022;23(1):544. doi:10.1186/s12864-022-08748-6

244. Kyoreva M, Li Y, Hoosenally M, Hardman-Smart J, Morrison K, Tosi I, Tolaini M, Barinaga G, Stockinger B, Mrowietz U, Nestle FO, et al. CYP1A1 Enzymatic Activity Influences Skin Inflammation Via Regulation of the AHR Pathway. J Invest Dermatol. 2021;141(6):1553–1563.e3. doi:10.1016/j.jid.2020.11.024

245. Wang X, Zheng Z, Caviglia JM, Corey KE, Herfel TM, Cai B, Masia R, Chung RT, Lefkowitch JH, Schwabe RF, et al. Hepatocyte TAZ/WWTR1 Promotes Inflammation and Fibrosis in Nonalcoholic Steatohepatitis. Cell Metab. 2016;24(6):848–862. doi:10.1016/j.cmet.2016.09.01

246. Lamkanfi M, Kanneganti TD. Caspase-7: a protease involved in apoptosis and inflammation. Int J Biochem Cell Biol. 2010;42(1):21–24. doi:10.1016/j.biocel.2009.09.013

247. Khan S, Mentrup HL, Novak EA, Siow VS, Wang Q, Crawford EC, Schneider C, Comerford TE 4th, Firek B, Rogers MB, Loughran P, et al. Cyclic GMP-AMP synthase contributes to epithelial homeostasis in intestinal inflammation via Beclin-1-mediated autophagy. FASEB J. 2022;36(5):e22282. doi:10.1096/fj.202200138R

248. Hazell GG, Peachey AM, Teasdale JE, Sala-Newby GB, Angelini GD, Newby AC, White SJ. PI16 is a shear stress and inflammation-regulated inhibitor of MMP2. Sci Rep. 2016;6:39553. doi:10.1038/srep39553

249. Yamamura K, Uruno T, Shiraishi A, Tanaka Y, Ushijima M, Nakahara T, Watanabe M, Kido-Nakahara M, Tsuge I, Furue M, et al. The transcription factor EPAS1 links DOCK8 deficiency to atopic skin inflammation via IL-31 induction. Nat Commun. 2017;8:13946. doi:10.1038/ncomms13946

250. Zhang T, Wu J, Ungvijanpunya N, Jackson-Weaver O, Gou Y, Feng J, Ho TV, Shen Y, Liu J, Richard S, et al. Smad6 Methylation Represses NFκB Activation and Periodontal Inflammation. J Dent Res. 2018;97(7):810–819. doi:10.1177/0022034518755688

251. Chen Z, Liang Y, Lu Q, Nazar M, Mao Y, Aboragah A, Yang Z, Loor JJ. Cadmium promotes apoptosis and inflammation via the circ08409/miR-133a/TGFB2 axis in bovine mammary epithelial cells and mouse mammary gland. Ecotoxicol Environ Saf. 2021;222:112477. doi:10.1016/j.ecoenv.2021.112477

252. Liu L, Huang S, Du Y, Zhou H, Zhang K, He J. Lats2 deficiency protects the heart against myocardial infarction by reducing inflammation and inhibiting mitochondrial fission and STING/p65 signaling. Int J Biol Sci. 2023;19(11):3428–3440. doi:10.7150/ijbs.84426

253. Zeng X, Tian G, Zhu J, Yang F, Zhang R, Li H, An Z, Li J, Song J, Jiang J, et al. Air pollution associated acute respiratory inflammation and modification by GSTM1 and GSTT1 gene polymorphisms: a panel study of healthy undergraduates. Environ Health. 2023;22(1):14. doi:10.1186/s12940-022-00954-9

254. Zhang K, Li Z, Lu Y, Xiang L, Sun J, Zhang H. Silencing of Vangl2 attenuates the inflammation promoted by Wnt5a via MAPK and NF-κB pathway in chondrocytes. J Orthop Surg Res. 2021;16(1):136. doi:10.1186/s13018-021-02268-x

255. Lotem J, Levanon D, Negreanu V, Bauer O, Hantisteanu S, Dicken J, Groner Y. Runx3 in Immunity, Inflammation and Cancer. Adv Exp Med Biol. 2017;962:369–393. doi:10.1007/978-981-10-3233-2_23

256. Krause K, Sabat R, Witte-Händel E, Schulze A, Puhl V, Maurer M, Wolk K. Association of CCL2 with systemic inflammation in Schnitzler syndrome. Br J Dermatol. 2019;180(4):859–868. doi:10.1111/bjd.17334

257. Artym J, Zimecki M, Kruzel ML. Lactoferrin for Prevention and Treatment of Anemia and Inflammation in Pregnant Women: A Comprehensive Review. Biomedicines. 2021;9(8):898. doi:10.3390/biomedicines9080898

258. Yildirim S, Sengul E, Aksu EH, Cinar İ, Gelen V, Tekin S, Dag Y. Selenium reduces acrylamide-induced testicular toxicity in rats by regulating HSD17B1, StAR, and CYP17A1 expression, oxidative stress, inflammation, apoptosis, autophagy, and DNA damage. Environ Toxicol. 2024;39(3):1402–1414. doi:10.1002/tox.23996

259. Portig I, Sandmoeller A, Kreilinger S, Maisch B. HLA-DQB1* polymorphism and associations with dilated cardiomyopathy, inflammatory dilated cardiomyopathy and myocarditis. Autoimmunity. 2009;42(1):33–40. doi:10.1080/08916930802258651

260. Mori T, Maeda N, Inoue K, Sekimoto R, Tsushima Y, Matsuda K, Yamaoka M, Suganami T, Nishizawa H, Ogawa Y, et al. A novel role for adipose ephrin-B1 in inflammatory response. PLoS One. 2013;8(10):e76199.doi:10.1371/journal.pone.0076199

261. Khadir A, Kavalakatt S, Dehbi M, Alarouj M, Bennakhi A, Tiss A, Elkum N. DUSP1 Is a Potential Marker of Chronic Inflammation in Arabs with Cardiovascular Diseases. Dis Markers. 2018;2018:9529621. doi:10.1155/2018/9529621

262. Huang MT, Chen YL, Lien CI, Liu WL, Hsu LC, Yagita H, Chiang BL. Notch Ligand DLL4 Alleviates Allergic Airway Inflammation via Induction of a Homeostatic Regulatory Pathway. Sci Rep. 2017;7:43535. doi:10.1038/srep43535

263. Verhamme FM, De Smet EG, Van Hooste W, Delanghe J, Verleden SE, Joos GF, Brusselle GG, Bracke KR. Bone morphogenetic protein 6 (BMP-6) modulates lung function, pulmonary iron levels and cigarette smoke-induced inflammation. Mucosal Immunol. 2019;12(2):340–351. doi:10.1038/s41385-018-0116-2

264. Liu M, Zhang L, Marsboom G, Jambusaria A, Xiong S, Toth PT, Benevolenskaya EV, Rehman J, Malik AB. Sox17 is required for endothelial regeneration following inflammation-induced vascular injury. Nat Commun. 2019;10(1):2126. doi:10.1038/s41467-019-10134-y

265. Baumgart S, Chen NM, Siveke JT, König A, Zhang JS, Singh SK, Wolf E, Bartkuhn M, Esposito I, Heßmann E, et al. Inflammation-induced NFATc1-STAT3 transcription complex promotes pancreatic cancer initiation by KrasG12D. Cancer Discov. 2014;4(6):688–701. doi:10.1158/2159-8290.CD-13-0593

266. Xu H, Du X, Liu G, Huang S, Du W, Zou S, Tang D, Fan C, Xie Y, Wei Y, et al. The pseudokinase MLKL regulates hepatic insulin sensitivity independently of inflammation. Mol Metab. 2019;23:14–23. doi:10.1016/j.molmet.2019.02.003

267. Jingjing H, Tongqian W, Shirong Y, Lan M, Jing L, Shihui M, Haijian Y, Fang Y. S100A4 promotes experimental autoimmune encephalomyelitis by impacting microglial inflammation through TLR4/NF-κB signaling pathway. Int Immunopharmacol. 2024;142(Pt A):112849. doi:10.1016/j.intimp.2024.112849

268. Kubo T, Sato S, Hida T, Minowa T, Hirohashi Y, Tsukahara T, Kanaseki T, Murata K, Uhara H, Torigoe T. IL-13 modulates ΔNp63 levels causing altered expression of barrier- and inflammation-related molecules in human keratinocytes: A possible explanation for chronicity of atopic dermatitis. Immun Inflamm Dis. 2021;9(3):734–745. doi:10.1002/iid3.427

269. Gao T, Zhu L, Liu H, Zhang X, Wang T, Fu Y, Li H, Dong Q, Hu Y, Zhang Z, et al. Highly pathogenic coronavirus N protein aggravates inflammation by MASP-2-mediated lectin complement pathway overactivation. Signal Transduct Target Ther. 2022;7(1):318. doi:10.1038/s41392-022-01133-5

270. Du J, Wei X, Ge X, Chen Y, Li YC. Microbiota-Dependent Induction of Colonic Cyp27b1 Is Associated With Colonic Inflammation: Implications of Locally Produced 1,25-Dihydroxyvitamin D3 in Inflammatory Regulation in the Colon. Endocrinology. 2017;158(11):4064–4075. doi:10.1210/en.2017-00578

271. Stoeltje L, Luc JK, Haddad T, Schrankel CS. The roles of ABCB1/P-glycoprotein drug transporters in regulating gut microbes and inflammation: insights from animal models, old and new. Philos Trans R Soc Lond B Biol Sci. 2024;379(1901):20230074. doi:10.1098/rstb.2023.0074

272. Brodzikowska A, Gondek A, Rak B, Paskal W, Pełka K, Cudnoch-Jędrzejewska A, Włodarski P. Metalloproteinase 14 (MMP-14) and hsa-miR-410-3p expression in human inflamed dental pulp and odontoblasts. Histochem Cell Biol. 2019;152(5):345–353. doi:10.1007/s00418-019-01811-6

273. Xu J, Jiang C, Cai Y, Guo Y, Wang X, Zhang J, Xu J, Xu K, Zhu W, Wang S, et al. Intervening upregulated SLC7A5 could mitigate inflammatory mediator by mTOR-P70S6K signal in rheumatoid arthritis synoviocytes. Arthritis Res Ther. 2020;22(1):200. doi:10.1186/s13075-020-02296-8

274. Mazrouei S, Ziaei A, Tanhaee AP, Keyhanian K, Esmaeili M, Baradaran A, Salehi M. Apoptosis inhibition or inflammation: the role of NAIP protein expression in Hodgkin and non-Hodgkin lymphomas compared to non-neoplastic lymph node. J Inflamm (Lond). 2012;9(1):4. doi:10.1186/1476-9255-9-4

275. Hsieh PC, Huang KL, Peng CK, Wu YK, Liu GT, Kuo CY, Wang MC, Lan CC. Aqueous extract of Descuraniae Semen attenuates lipopolysaccharide-induced inflammation by inhibiting ER stress and WNK4-SPAK-NKCC1 pathway. J Cell Mol Med. 2024;28(15):e18589. doi:10.1111/jcmm.18589

276. Zhang BS, Zhang XM, Ito M, Yajima S, Yoshida K, Ohno M, Nishi E, Wang H, Li SY, Kubota M, et al. JMJD6 Autoantibodies as a Potential Biomarker for Inflammation-Related Diseases. Int J Mol Sci. 2024;25(9):4935. doi:10.3390/ijms25094935

277. Lin L, Spoor MS, Gerth AJ, Brody SL, Peng SL. Modulation of Th1 activation and inflammation by the NF-kappaB repressor Foxj1. Science. 2004;303(5660):1017-1020. doi:10.1126/science.1093889

278. Jia Y, Pan J. CKLF1, transcriptionally activated by FOXC1, promotes hypoxia/reoxygenation induced oxidative stress and inflammation in H9c2 cells by NLRP3 inflammasome activation. Exp Ther Med. 2023;27(2):59. doi:10.3892/etm.2023.12347

279. Kalin TV, Meliton L, Meliton AY, Zhu X, Whitsett JA, Kalinichenko VV. Pulmonary mastocytosis and enhanced lung inflammation in mice heterozygous null for the Foxf1 gene. Am J Respir Cell Mol Biol. 2008;39(4):390–399. doi:10.1165/rcmb.2008-0044OC

280. Lin W, Xu D, Austin CD, Caplazi P, Senger K, Sun Y, Jeet S, Young J, Delarosa D, Suto E, et al. Function of CSF1 and IL34 in Macrophage Homeostasis, Inflammation, and Cancer. Front Immunol. 2019;10:2019. doi:10.3389/fimmu.2019.02019

281. Na YR, Jung D, Stakenborg M, Jang H, Gu GJ, Jeong MR, Suh SY, Kim HJ, Kwon YH, Sung TS, et al. Prostaglandin E2 receptor PTGER4-expressing macrophages promote intestinal epithelial barrier regeneration upon inflammation. Gut. 2021;70(12):2249–2260. doi:10.1136/gutjnl-2020-322146

282. Meurer SK, Weiskirchen R. Endoglin: An ’Accessory’ Receptor Regulating Blood Cell Development and Inflammation. Int J Mol Sci. 2020;21(23):9247. doi:10.3390/ijms21239247

283. Long C, Liu H, Zhan W, Chen L, Wu A, Yang L, Chen S.Null Function of Npr1 Disturbs Immune Response in Colonic Inflammation During Early Postnatal Stage. Inflammation. 2022;45(6):2419–2432. doi:10.1007/s10753-022-01702-4

284. Korhonen EA, Lampinen A, Giri H, Anisimov A, Kim M, Allen B, Fang S, D’Amico G, Sipilä TJ, Lohela M, et al. Tie1 controls angiopoietin function in vascular remodeling and inflammation. J Clin Invest. 2016;126(9):3495–3510. doi:10.1172/JCI84923

285. Xu G, Gong Y, Lu F, Wang B, Yang Z, Chen L, Min J, Cheng C, Jiang T. Endothelin receptor B enhances liver injury and pro-inflammatory responses by increasing G-protein-coupled receptor kinase-2 expression in primary biliary cholangitis. Sci Rep. 2022;12(1):19772. doi:10.1038/s41598-022-21816-x

286. Ahmad S, Wrennall JA, Goriounova AS, Sekhri M, Iskarpatyoti JA, Ghosh A, Abdelwahab SH, Voeller A, Rai M, Mahida RY, et al. pecific Inhibition of Orai1-mediated Calcium Signalling Resolves Inflammation and Clears Bacteria in an Acute Respiratory Distress Syndrome Model. Am J Respir Crit Care Med. 2024;209(6):703–715. doi:10.1164/rccm.202308-1393OC

287. Yin J, Wang C, Vogel U, Ma Y, Zhang Y, Wang H, Sun Z, Du S. Common variants of pro-inflammatory gene IL1B and interactions with PPP1R13L and POLR1G in relation to lung cancer among Northeast Chinese. Sci Rep. 2023;13(1):7352. doi:10.1038/s41598-023-34069-z

288. Lian X, Zhang X, Chen W, Xue F, Wang G. Dexmedetomidine mitigates neuroinflammation in an Alzheimer’s disease mouse model via the miR-204-3p/FBXL7 signaling axis. Brain Res. 2024;1822:148612. doi:10.1016/j.brainres.2023.148612

289. Tu F, Wang X, Zhang X, Ha T, Wang Y, Fan M, Yang K, Gill PS, Ozment TR, Dai Y, et al. Novel Role of Endothelial Derived Exosomal HSPA12B in Regulating Macrophage Inflammatory Responses in Polymicrobial Sepsis. Front Immunol. 2020;11:825. doi:10.3389/fimmu.2020.00825

290. Fairbanks CA, Peterson CD, Speltz RH, Riedl MS, Kitto KF, Dykstra JA, Braun PD, Sadahiro M, Salton SR, Vulchanova L. The VGF-derived peptide TLQP-21 contributes to inflammatory and nerve injury-induced hypersensitivity. Pain. 2014;155(7):1229–1237. doi:10.1016/j.pain.2014.03.012

291. Kato N, Nemoto K, Arino H, Fujikawa K. Influence of peripheral inflammation on growth-associated phosphoprotein (GAP-43) expression in dorsal root ganglia and on nerve recovery after crush injury. Neurosci Res. 2003;45(3):297–303. doi:10.1016/s0168-0102(02)00234-1

292. Yan S, Ju X, Lao J, et al. Overexpression of the Mas1 gene mitigated LPS-induced inflammatory injury in mammary epithelial cells by inhibiting the NF-κB/MAPKs signaling pathways. Front Vet Sci. 2024;11:1446366. doi:10.3389/fvets.2024.1446366

293. Jósvay K, Winter Z, Katona RL, Pecze L, Marton A, Buhala A, Szakonyi G, Oláh Z, Vizler C. Besides neuro-imaging, the Thy1-YFP mouse could serve for visualizing experimental tumours, inflammation and wound-healing. Sci Rep. 2014;4:6776. doi:10.1038/srep06776

294. Alrashdi B, Dawod B, Tacke S, Kuerten S, Côté PD, Marshall JS. Mice Heterozygous for the Sodium Channel Scn8a (Nav1.6) Have Reduced Inflammatory Responses During EAE and Following LPS Challenge. Front Immunol. 2021;12:533423. doi:10.3389/fimmu.2021.533423

295. Wang L, Li X, Song Y, Zhang L, Ye L, Zhou X, Song D, Huang D. NELL1 augments osteogenesis and inhibits inflammation of human periodontal ligament stem cells induced by BMP9. J Periodontol. 2022;93(7):977–987. doi:10.1002/JPER.20-0517

296. Xie Z, Chan EC, Druey KM. R4 Regulator of G Protein Signaling (RGS) Proteins in Inflammation and Immunity. AAPS J. 2016;18(2):294–304. doi:10.1208/s12248-015-9847-0

297. Leoncini S, De Felice C, Signorini C, Zollo G, Cortelazzo A, Durand T, Galano JM, Guerranti R, Rossi M, Ciccoli L, et al. Cytokine Dysregulation in MECP2-and CDKL5-Related Rett Syndrome: Relationships with Aberrant Redox Homeostasis, Inflammation, and ω-3 PUFAs. Oxid Med Cell Longev. 2015;2015:421624. doi:10.1155/2015/421624

298. Mulder DJ, Pacheco I, Hurlbut DJ, Mak N, Furuta GT, MacLeod RJ, Justinich CJ. FGF9-induced proliferative response to eosinophilic inflammation in oesophagitis. Gut. 2009;58(2):166–173. doi:10.1136/gut.2008.157628

299. Duan XL, Guo Z, He YT, Li YX, Liu YN, Bai HH, Li HL, Hu XD, Suo ZW. SNAP25/syntaxin4/VAMP2/Munc18-1 Complexes in Spinal Dorsal Horn Contributed to Inflammatory Pain. Neuroscience. 2020;429:203–212. doi:10.1016/j.neuroscience.2020.01.003

300. Yang X, Lu L, Wu C, Zhang F. ATP2B1-AS1 exacerbates sepsis-induced cell apoptosis and inflammation by regulating miR-23a-3p/TLR4 axis. Allergol Immunopathol (Madr). 2023;51(2):17–26. doi:10.15586/aei.v51i2.782

301. Arnaud L, Benech P, Greetham L, Stephan D, Jimenez A, Jullien N, García-González L, Tsvetkov PO, Devred F, Sancho-Martinez I, et al. APOE4 drives inflammation in human astrocytes via TAGLN3 repression and NF-κB activation. Cell Rep. 2022;40(7):111200. doi:10.1016/j.celrep.2022.111200

302. Ho TC, Yeh SI, Chen SL, Tsao YP. Integrin αv and Vitronectin Prime Macrophage-Related Inflammation and Contribute the Development of Dry Eye Disease. Int J Mol Sci. 2021;22(16):8410. doi:10.3390/ijms22168410

303. Wang D, Qian W, Wu D, Wu Y, Lu K, Zou G. METTL3 promotes microglial inflammation via MEF2C in spinal cord injury. Cell Tissue Res. 2024;395(2):189–197. doi:10.1007/s00441-023-03855-6

304. Sui J, Zhan L, Ji S, Wu W, Chen Y, Yun F, Liang W, Wang J, Cao M, Shen D, et al. Differential inflammation responses determine the variable phenotypes of epilepsy induced by GABRG2 mutations. CNS Neurosci Ther. 2024;30(2):e14583. doi:10.1111/cns.14583

305. Acosta C, McMullan S, Djouhri L, Gao L, Watkins R, Berry C, Dempsey K, Lawson SN. HCN1 and HCN2 in Rat DRG neurons: levels in nociceptors and non-nociceptors, NT3-dependence and influence of CFA-induced skin inflammation on HCN2 and NT3 expression. PLoS One. 2012;7(12):e50442. doi:10.1371/journal.pone.0050442

306. Bencsics M, Bányai B, Ke H, Csépányi-Kömi R, Sasvári P, Dantzer F, Hanini N, Benkő R, Horváth EM. PARP2 downregulation in T cells ameliorates lipopolysaccharide-induced inflammation of the large intestine. Front Immunol. 2023;14:1135410. doi:10.3389/fimmu.2023.1135410

307. Xia Q, Zhan G, Mao M, Zhao Y, Li X. TRIM45 causes neuronal damage by aggravating microglia-mediated neuroinflammation upon cerebral ischemia and reperfusion injury. Exp Mol Med. 2022;54(2):180–193. doi:10.1038/s12276-022-00734-y

308. Chen M, Pan L, Chen D, Wu Y, Ye J, Li K, Zhang N, Xu J. PAK1 Promotes Inflammation Induced by Sepsis through the Snail/CXCL2 Signaling Pathway. ACS Infect Dis. 2024;10(4):1370–1378. doi:10.1021/acsinfecdis.4c00052

309. Chen Y, Wang B, Chen Y, Wu Q, Lai WF, Wei L, Nandakumar KS, Liu D. HAPLN1 Affects Cell Viability and Promotes the Pro-Inflammatory Phenotype of Fibroblast-Like Synoviocytes. Front Immunol. 2022;13:888612. doi:10.3389/fimmu.2022.888612

310. Zhang Q, Zhou L, Xie H, Zhang H, Gao X. HAGLR aggravates neuropathic pain and promotes inflammatory response and apoptosis of lipopolysaccharide-treated SH-SY5Y cells by sequestering miR-182-5p from ATAT1 and activating NLRP3 inflammasome. Neurochem Int. 2021;145:105001. doi:10.1016/j.neuint.2021.105001

311. Tuysuz EC, Mourati E, Rosberg R, Moskal A, Gialeli C, Johansson E, Governa V, Belting M, Pietras A, Blom AM. Tumor suppressor role of the complement inhibitor CSMD1 and its role in TNF-induced neuroinflammation in gliomas. J Exp Clin Cancer Res. 2024;43(1):98. doi:10.1186/s13046-024-03019-6

312. Yi J, Bergstrom K, Fu J, Shan X, McDaniel JM, McGee S, Qu D, Houchen CW, Liu X, Xia L. Dclk1 in tuft cells promotes inflammation-driven epithelial restitution and mitigates chronic colitis. Cell Death Differ. 2019;26(9):1656–1669. doi:10.1038/s41418-018-0237-x

313. Yun JH, Lee C, Liu T, Liu S, Kim EY, Xu S, Curtis JL, Pinello L, Bowler RP, Silverman EK, et al. Hedgehog interacting protein-expressing lung fibroblasts suppress lymphocytic inflammation in mice. JCI Insight. 2021;6(17):e144575. doi:10.1172/jci.insight.144575

314. Sun H, Li Z, Fan C, Liu S, Yan K, Huang G, Li S. Slit guidance ligand 2 promotes the inflammatory response of periodontitis through activation of the NF-κB signaling pathway. J Periodontal Res. 2022;57(3):578–586. doi:10.1111/jre.12987

315. Zhang S, Cheng L, Su Y, Qian Z, Wang Z, Chen C, Li R, Zhang A, He J, Mao J, et al. AGBL4 promotes malignant progression of glioblastoma via modulation of MMP-1 and inflammatory pathways. Front Immunol. 2024;15:1420182. Published 2024 Jun 28. doi:10.3389/fimmu.2024.1420182

316. Rienks M, Carai P, Bitsch N, Schellings M, Vanhaverbeke M, Verjans J, Cuijpers I, Heymans S, Papageorgiou A. Sema3A promotes the resolution of cardiac inflammation after myocardial infarction. Basic Res Cardiol. 2017;112(4):42. doi:10.1007/s00395-017-0630-5

317. Zhang Z, Liu N, Chen X, Zhang F, Kong T, Tang X, Yang Q, Chen W, Xiong X, Chen X. UCHL1 regulates inflammation via MAPK and NF-κB pathways in LPS-activated macrophages. Cell Biol Int. 2021;45(10):2107–2117. doi:10.1002/cbin.11662

318. Lehman CW, Smith A, Kelly J, Jacobs JL, Dinman JD, Kehn-Hall K. EGR1 Upregulation during Encephalitic Viral Infections Contributes to Inflammation and Cell Death. Viruses. 2022;14(6):1210. doi:10.3390/v14061210

319. Isidro-Hernández M, Mayado A, Casado-García A, Martínez-Cano J, Palmi C, Fazio G, Orfao A, Ribera J, Ribera JM, Zamora L, et al. Inhibition of inflammatory signaling in Pax5 mutant cells mitigates B-cell leukemogenesis. Sci Rep. 2020;10(1):19189. doi:10.1038/s41598-020-76206-y

320. Zhang X, Chen W, Liu W, Li D, Shen W. CITED2 alleviates lipopolysaccharide-induced inflammation and pyroptosis in - human lung fibroblast by inhibition of NF-κB pathway. Allergol Immunopathol (Madr). 2022;50(4):64–70. doi:10.15586/aei.v50i4.628

321. Huan Z, Tang Y, Xu C, Cai J, Yao H, Wang Y, Bu F, Ge X. PTPRO knockdown protects against inflammation in hemorrhage shock-induced lung injury involving the NF-κB signaling pathway. Respir Res. 2022;23(1):195. doi:10.1186/s12931-022-02118-2

322. Cabezudo D, Baekelandt V, Lobbestael E. Multiple-Hit Hypothesis in Parkinson’s Disease: LRRK2 and Inflammation. Front Neurosci. 2020;14:376. doi:10.3389/fnins.2020.00376

323. Marchese E, Valentini M, Di Sante G, Cesari E, Adinolfi A, Corvino V, Ria F, Sette C, Geloso MC. Alternative splicing of neurexins 1-3 is modulated by neuroinflammation in the prefrontal cortex of a murine model of multiple sclerosis. Exp Neurol. 2021;335:113497. doi:10.1016/j.expneurol.2020.113497

324. Luo Q, Jahangir A, He J, Huang C, Xia Y, Jia L, Wei X, Pan T, Du Y, Mu B, et al. Ameliorating Effects of TRIM67 against Intestinal Inflammation and Barrier Dysfunction Induced by High Fat Diet in Obese Mice. Int J Mol Sci. 2022;23(14):7650. doi:10.3390/ijms23147650

325. Panda S, Gekara NO. The deubiquitinase MYSM1 dampens NOD2-mediated inflammation and tissue damage by inactivating the RIP2 complex. Nat Commun. 2018;9(1):4654. doi:10.1038/s41467-018-07016-0

326. Karki P, Ke Y, Zhang CO, Li Y, Tian Y, Son S, Yoshimura A, Kaibuchi K, Birukov KG, Birukova AA. SOCS3-microtubule interaction via CLIP-170 and CLASP2 is critical for modulation of endothelial inflammation and lung injury. J Biol Chem. 2021;296:100239. doi:10.1074/jbc.RA120.014232

327. Kim DE, Byeon HE, Kim DH, Kim SG, Yim H. Plk2-mediated phosphorylation and translocalization of Nrf2 activates anti-inflammation through p53/Plk2/p21cip1 signaling in acute kidney injury. Cell Biol Toxicol. 2023;39(4):1509–1529. doi:10.1007/s10565-022-09741-1

328. Pang W, Hu F. C9ORF72 suppresses JAK-STAT mediated inflammation. iScience. 2023;26(5):106579. doi:10.1016/j.isci.2023.106579

329. Hernández C, Arroba AI, Bogdanov P, Ramos H, Simó-Servat O, Simó R, Valverde AM. Effect of Topical Administration of Somatostatin on Retinal Inflammation and Neurodegeneration in an Experimental Model of Diabetes. J Clin Med. 2020;9(8):2579. doi:10.3390/jcm9082579

330. Miao Y, Wu S, Gong Z, Chen Y, Xue F, Liu K, Zou J, Feng Y, Li G. SPARCL1 promotes chondrocytes extracellular matrix degradation and inflammation in osteoarthritis via TNF/NF-κB pathway. J Orthop Translat. 2024;46:116–128. doi:10.1016/j.jot.2024.02.009

331. Yang C, Blaize G, Marrocco R, Rouquié N, Bories C, Gador M, Mélique S, Joulia E, Benamar M, Dejean AS, et al. THEMIS enhances the magnitude of normal and neuroinflammatory type 1 immune responses by promoting TCR-independent signals. Sci Signal. 2022;15(742):eabl5343. doi:10.1126/scisignal.abl5343

332. Tang W, Dong M, Teng F, Cui J, Zhu X, Wang W, Wuniqiemu T, Qin J, Yi L, Wang S, et al. TMT-based quantitative proteomics reveals suppression of SLC3A2 and ATP1A3 expression contributes to the inhibitory role of acupuncture on airway inflammation in an OVA-induced mouse asthma model. Biomed Pharmacother. 2021;134:111001. doi:10.1016/j.biopha.2020.111001

333. Shen C, Kuang Y, Xu S, Li R, Wang J, Zou Y, Wang C, Xu S, Liang L, Lin C, et al. Nitidine chloride inhibits fibroblast like synoviocytes- mediated rheumatoid synovial inflammation and joint destruction by targeting KCNH1. Int Immunopharmacol. 2021;101(Pt A):108273. doi:10.1016/j.intimp.2021.108273

334. Deng S, Guo A, Huang Z, Guan K, Zhu Y, Chan C, Gui J, Song C, Li X. The exploration of neuroinflammatory mechanism by which CRHR2 deficiency induced anxiety disorder. Prog Neuropsychopharmacol Biol Psychiatry. 2024;128:110844. doi:10.1016/j.pnpbp.2023.110844

335. Zeng J, Wang Y, Luo Z, Chang LC, Yoo JS, Yan H, Choi Y, Xie X, Deverman BE, Gradinaru V, et al. TRIM9-Mediated Resolution of Neuroinflammation Confers Neuroprotection upon Ischemic Stroke in Mice. Cell Rep. 2023;42(8):113050. doi:10.1016/j.celrep.2023.113050

336. Yang X, He Z, Dong Q, Nai S, Duan X, Yu J, Zhao N, Du X, Chen L. Btbd8 deficiency reduces susceptibility to colitis by enhancing intestinal barrier function and suppressing inflammation. Front Immunol. 2024;15:1382661. doi:10.3389/fimmu.2024.1382661

337. Gracia-Maldonado G, Clark J, Burwinkel M, Greenslade B, Wunderlich M, Salomonis N, Leone D, Gatti E, Pierre P, Kumar AR, et al. LAMP-5 is an essential inflammatory-signaling regulator and novel immunotherapy target for mixed lineage leukemia-rearranged acute leukemia. Haematologica. 2022;107(4):803–815. doi:10.3324/haematol.2020.257451

338. Sheng M, Huo S, Jia L, Weng Y, Liu W, Lin Y, Yu W. NUAK1 promotes metabolic dysfunction-associated steatohepatitis progression by activating Caspase 6-driven pyroptosis and inflammation. Hepatol Commun. 2024;8(7):e0479. doi:10.1097/HC9.0000000000000479

339. Xu Y, Zhan Y, Lew AM, Naik SH, Kershaw MH. Differential development of murine dendritic cells by GM-CSF versus Flt3 ligand has implications for inflammation and trafficking. J Immunol. 2007;179(11):7577–7584. doi:10.4049/jimmunol.179.11.7577

340. Chen TT, Wu SM, Ho SC, Chuang HC, Liu CY, Chan YF, Kuo LW, Feng PH, Liu WT, Chen KY, et al. Reduction Is Implicated in Abnormal Inflammation in COPD. Sci Rep. 2017;7:46667. doi:10.1038/srep46667

341. Huang J, Tang D, Cao Y, Wang Y, Long J, Wei L, Song H. Inhibition of PDE10A-Rescued TBI-Induced Neuroinflammation and Apoptosis through the cAMP/PKA/NLRP3 Pathway. Evid Based Complement Alternat Med. 2022;2022:3311250. doi:10.1155/2022/3311250

342. Zheng K, Zhang Y, Zhang C, Ye W, Ye C, Tan X, Xiong Y. PRMT8 Attenuates Cerebral Ischemia/Reperfusion Injury via Modulating Microglia Activation and Polarization to Suppress Neuroinflammation by Upregulating Lin28a. ACS Chem Neurosci. 2022;13(7):1096–1104. doi:10.1021/acschemneuro.2c00096

343. Zhang YX, Zhang XT, Li HJ, Zhou TF, Zhou AC, Zhong ZL, Liu YH, Yuan LL, Zhu HY, Luan D, et al. Antidepressant-like effects of helicid on a chronic unpredictable mild stress-induced depression rat model: Inhibiting the IKK/IκBα/NF-κB pathway through NCALD to reduce inflammation. Int Immunopharmacol. 2021;93:107165. doi:10.1016/j.intimp.2020.107165

344. Hong H, Zeng Y, Jian W, Li L, Lin L, Mo Y, Liu M, Fang S, Xia Y. CDK7 inhibition suppresses rheumatoid arthritis inflammation via blockage of NF-κB activation and IL-1β/IL-6 secretion. J Cell Mol Med. 2018;22(2):1292–1301. doi:10.1111/jcmm.13414

345. Chen Y, Ku H, Zhao L, Wheeler DC, Li LC, Li Q, Varghese Z, Moorhead JF, Powis SH, Huang A, et al. Inflammatory stress induces statin resistance by disrupting 3-hydroxy-3-methylglutaryl-CoA reductase feedback regulation. Arterioscler Thromb Vasc Biol. 2014;34(2):365–376. doi:10.1161/ATVBAHA.113.301301

346. Feng X, Lu J, Wu Y, Xu H. MiR-18a-3p improves cartilage matrix remodeling and inhibits inflammation in osteoarthritis by suppressing PDP1. J Physiol Sci. 2022;72(1):3. doi:10.1186/s12576-022-00827-3

347. Belgath AA, Emam AM, Taujanskas J, Bryce RA, Freeman S, Stratford IJ. Discovery of Potent Benzothiazole Inhibitors of Oxidoreductase NQO2, a Target for Inflammation and Cancer. Int J Mol Sci. 2024;25(22):12025. doi:10.3390/ijms252212025

348. Hua J, Gao Z, Zhong S, Wei B, Zhu J, Ying R. CISD1 protects against atherosclerosis by suppressing lipid accumulation and inflammation via mediating Drp1. Biochem Biophys Res Commun. 2021;577:80–88. doi:10.1016/j.bbrc.2021.08.023

349. Tubau-Juni N, Hontecillas R, Leber A, Maturavongsadit P, Chauhan J, Bassaganya-Riera J. First-in-class topical therapeutic omilancor ameliorates disease severity and inflammation through activation of LANCL2 pathway in psoriasis. Sci Rep. 2021;11(1):19827. doi:10.1038/s41598-021-99349-y

350. Jing C, Dongming Z, Hong C, Quan N, Sishi L, Caixia L. TRPC3 Overexpression Promotes the Progression of Inflammation-Induced Preterm Labor and Inhibits T Cell Activation. Cell Physiol Biochem. 2018;45(1):378–388. doi:10.1159/000486912

351. Nguyen HC, Bu S, Nikfarjam S, Rasheed B, Michels DCR, Singh A, Singh S, Marszal C, McGuire JJ, Feng Q, et al. Loss of fatty acid binding protein 3 ameliorates lipopolysaccharide-induced inflammation and endothelial dysfunction. J Biol Chem. 2023;299(3):102921. doi:10.1016/j.jbc.2023.102921

352. Zhu WS, Guo W, Zhu JN, Tang CM, Fu YH, Lin QX, Tan N, Shan ZX. Hsp90aa1: a novel target gene of miR-1 in cardiac ischemia/reperfusion injury. Sci Rep. 2016;6:24498. doi:10.1038/srep24498

353. M.M. Page, K.L. Ellis, D.C. Chan, J. Pang, A.J. Hooper, D.A. Bell, J.R. Burnett, E.K. Moses, G.F. Watts. A variant in the fibronectin (FN1) gene, rs1250229-T, is associated with decreased risk of coronary artery disease in familial hypercholesterolaemia. J Clin Lipidol. 2022;16(4):525–529. doi:10.1016/j.jacl.2022.05.065

354. Li TY, Su W, Li LL, Zhao XG, Yang N, Gai JX, Lv X, Zhang J, Huang MQ, Zhang Q, et al. Critical role of PAFR/YAP1 positive feedback loop in cardiac fibrosis. Acta Pharmacol Sin. 2022;43(11):2862–2872. doi:10.1038/s41401-022-00903-9

355. Elrayess MA, Talmud PJ. Platelet endothelial cell adhesion molecule-1 (PECAM-1) & coronary heart disease. Indian J Med Res. 2005;121(2):77–79.

356. Yuan J, Chen H, Ge D, Xu Y, Xu H, Yang Y, Gu M, Zhou Y, Zhu J, Ge T, et al. Mir-21 Promotes Cardiac Fibrosis After Myocardial Infarction Via Targeting Smad7. Cell Physiol Biochem. 2017;42(6):2207–2219. doi:10.1159/000479995

357. Cho HM, Lee KH, Shen YM, Shin TJ, Ryu PD, Choi MC, Kang KS, Cho JY. Transplantation of hMSCs Genome Edited with LEF1 Improves Cardio-Protective Effects in Myocardial Infarction. Mol Ther Nucleic Acids. 2020;19:1186–1197. doi:10.1016/j.omtn.2020.01.007

358. Tian D, Qin Q, Liu R, Wang Z, Li X, Xu Q, Lv Q. Diagnostic Value of Circulating Progranulin and Its Receptor EphA2 in Predicting the Atheroma Burden in Patients with Coronary Artery Disease. Dis Markers. 2021;2021:6653501. doi:10.1155/2021/6653501

359. Wang Z, Li C, Sun X, Li Z, Li J, Wang L, Sun Y. Hypermethylation of miR-181b in monocytes is associated with coronary artery disease and promotes M1 polarized phenotype via PIAS1-KLF4 axis. Cardiovasc Diagn Ther. 2020;10(4):738–751. doi:10.21037/cdt-20-407

360. Bowers SLK, Meng Q, Kuwabara Y, Huo J, Minerath R, York AJ, Sargent MA, Prasad V, Saviola AJ, Galindo DC, et al. Col1a2-Deleted Mice Have Defective Type I Collagen and Secondary Reactive Cardiac Fibrosis with Altered Hypertrophic Dynamics. Cells. 2023;12(17):2174. doi:10.3390/cells12172174

361. Kraemer BF, Mannell H, Lamkemeyer T, Franz-Wachtel M, Lindemann S. Heat-Shock Protein 27 (HSPB1) Is Upregulated and Phosphorylated in Human Platelets during ST-Elevation Myocardial Infarction. Int J Mol Sci. 2019;20(23):5968. doi:10.3390/ijms20235968

362. Mauricio R, Singh K, Sanghavi M, Ayers CR, Rohatgi A, Vongpatanasin W, de Lemos JA, Khera A. Soluble Fms-like tyrosine kinase-1 (sFlt-1) is associated with subclinical and clinical atherosclerotic cardiovascular disease: The Dallas Heart Study. Atherosclerosis. 2022;346:46–52. doi:10.1016/j.atherosclerosis.2022.02.026

363. Deng Q, Wang X, Gao J, Xia X, Wang Y, Zhang Y, Chen Y. Growth restriction and congenital heart disease caused by a novel TAB2 mutation: A case report. Exp Ther Med. 2023;25(6):258. doi:10.3892/etm.2023.11957

364. Monaghan RM, Naylor RW, Flatman D, Kasher PR, Williams SG, Keavney BD. FLT4 causes developmental disorders of the cardiovascular and lymphovascular systems via pleiotropic molecular mechanisms. Cardiovasc Res. 2024;120(10):1164–1176. doi:10.1093/cvr/cvae104

365. Muiya NP, Wakil S, Al-Najai M, Tahir AI, Baz B, Andres E, Al-Boudari O, Al-Tassan N, Al-Shahid M, Meyer BF, et al. A study of the role of GATA2 gene polymorphism in coronary artery disease risk traits. Gene. 2014;544(2):152–158. doi:10.1016/j.gene.2014.04.064

366. Ji Z, Guo J, Zhang R, Zuo W, Xu Y, Qu Y, Tao Z, Li X, Li Y, Yao Y, et al. ADAM8 deficiency in macrophages promotes cardiac repair after myocardial infarction via ANXA2-mTOR-autophagy pathway. J Adv Res. 2024. doi:10.1016/j.jare.2024.07.037

367. Eskandari M, Awsat Mellati A, Mahmoodi K, Kamali K, Soltanpour MS. Association of the CYP1A1 rs4646903 polymorphism with susceptibility and severity of coronary artery disease. Mol Biol Res Commun. 2021;10(2):22–61. doi:10.22099/mbrc.2021.39141.1574

368. Zhang W, Zhang Y, Han L, Bo T, Qi Z, Zhong H, Xu H, Hu L, Chen S, Zhang S. Double-stranded DNA enhances platelet activation, thrombosis, and myocardial injury via cyclic GMP-AMP synthase. Cardiovasc Res. 2024. doi:10.1093/cvr/cvae218

369. Wang LY, Tan CS, Lai MKP, Hilal S. Factors Associated with RANTES, EMMPIRIN, MMP2 and MMP9, and the Association of These Biomarkers with Cardiovascular Disease in a Multi-Ethnic Population. J Clin Med. 2022;11(24):7281. doi:10.3390/jcm11247281

370. Sluimer JC. Metabolic and Shear Stress Regulate Endothelial Epas1 in Atherosclerosis. Circ Res. 2024;135(8):838–840. doi:10.1161/CIRCRESAHA.124.325131

371. Niu HM, Liu CL. The aberrant expression of Smad6 and TGF-β in obesity linked cardiac disease. Eur Rev Med Pharmacol Sci. 2017;21(1):138–142.

372. Rusu-Nastase EG, Lupan AM, Marinescu CI, Neculachi CA, Preda MB, Burlacu A. MiR-29a Increase in Aging May Function as a Compensatory Mechanism Against Cardiac Fibrosis Through SERPINH1 Downregulation. Front Cardiovasc Med. 2022;8:810241. doi:10.3389/fcvm.2021.810241

373. Sobha SP, Kesavarao KE. Progonostic effect of GSTM1/GSTT1 polymorphism in determining cardiovascular diseases risk among type 2 diabetes patients in South Indian population. Mol Biol Rep. 2023;50(8):6415–6423. doi:10.1007/s11033-023-08514-1

374. Su Z, Lu H, Jiang H, Zhu H, Li Z, Zhang P, Ni P, Shen H, Xu W, Xu H. IFN-γ-producing Th17 cells bias by HMGB1-T-bet/RUNX3 axis might contribute to progression of coronary artery atherosclerosis. Atherosclerosis. 2015;243(2):421–428. doi:10.1016/j.atherosclerosis.2015.09.037

375. Rodríguez I, Coto E, Reguero JR, González P, Andrés V, Lozano I, Martín M, Alvarez V, Morís C. Role of the CDKN1A/p21, CDKN1C/p57, and CDKN2A/p16 genes in the risk of atherosclerosis and myocardial infarction. Cell Cycle. 2007;6(5):620–625. doi:10.4161/cc.6.5.3927

376. Gholamalizadeh H, Ensan B, Sukhorukov VN, Sahebkar A. Targeting the CCL2-CCR2 signaling pathway: potential implications of statins beyond cardiovascular diseases. J Pharm Pharmacol. 2024;76(2):138–153. doi:10.1093/jpp/rgad112

377. Videm V, Dahl H, Wålberg LE, Wiseth R. Functional polymorphisms in the LTF gene and risk of coronary artery stenosis. Hum Immunol. 2012;73(5):554–559. doi:10.1016/j.humimm.2012.02.014

378. Wang W, Murray B, Tichnell C, Gilotra NA, Zimmerman SL, Gasperetti A, Scheel P, Tandri H, Calkins H, James CA. Clinical characteristics and risk stratification of desmoplakin cardiomyopathy. Europace. 2022;24(2):268–277. doi:10.1093/europace/euab183

379. Tulay P, Temel SG, Ergoren MC. Investigation of KCNQ1 polymorphisms as biomarkers for cardiovascular diseases in the Turkish Cypriots for establishing preventative medical measures. Int J Biol Macromol. 2019;124:537–540. doi:10.1016/j.ijbiomac.2018.11.227

380. Cong L, Zhao L, Shi Y, Bai Y, Guo Z. Circ_0006476 modulates macrophage apoptosis through the miR-3074-5p/DLL4 axis: implications for Notch signalling pathway regulation in cardiovascular disease. Aging (Albany NY). 2024;16(16):11857–11876. doi:10.18632/aging.206049

381. Banach J, Gilewski W, Słomka A, Buszko K, Błażejewski J, Karasek D, Rogowicz D, Żekanowska E, Sinkiewicz W. Bone morphogenetic protein 6-a possible new player in pathophysiology of heart failure. Clin Exp Pharmacol Physiol. 2016;43(12):1247–1250. doi:10.1111/1440-1681.12665

382. Lee JW, Lee CS, Son H, Lee J, Kang M, Chai J, Cho HJ, Kim HS. SOX17-mediated LPAR4 expression plays a pivotal role in cardiac development and regeneration after myocardial infarction. Exp Mol Med. 2023;55(7):1424–1436. doi:10.1038/s12276-023-01025-w

383. Rasheed A, Robichaud S, Nguyen MA, Geoffrion M, Wyatt H, Cottee ML, Dennison T, Pietrangelo A, Lee R, Lagace TA, et al. Loss of MLKL (Mixed Lineage Kinase Domain-Like Protein) Decreases Necrotic Core but Increases Macrophage Lipid Accumulation in Atherosclerosis. Arterioscler Thromb Vasc Biol. 2020;40(5):1155–1167. doi:10.1161/ATVBAHA.119.313640

384. Martínez-Campelo L, Blanco-Verea A, López-Fernández T, Martínez-Monzonís A, Buño A, Mazón P, Zamora P, Norton N, Reddy JS, Velasco-Ruiz A, et al. Meta-analysis of genome-wide association studies for cancer therapy-related cardiovascular dysfunction and functional mapping highlight an intergenic region close to TP63. Sci Rep. 2024;14(1):18413. doi:10.1038/s41598-024-69064-5

385. Catarino SJ, Boldt AB, Beltrame MH, Nisihara RM, Schafranski MD, de Messias-Reason IJ. Association of MASP2 polymorphisms and protein levels with rheumatic fever and rheumatic heart disease. Hum Immunol. 2014;75(12):1197–1202. doi:10.1016/j.humimm.2014.10.003

386. Li FF, Han Y, Shi S, Li X, Zhu XD, Zhou J, Shao QL, Li XQ, Liu SL. Characterization of Transcriptional Repressor Gene MSX1 Variations for Possible Associations with Congenital Heart Diseases. PLoS One. 2015;10(11):e0142666. doi:10.1371/journal.pone.0142666

387. Zhai YJ, Liu P, He HR, Zheng XW, Wang Y, Yang QT, Dong YL, Lu J.The association of ADORA2A and ADORA2B polymorphisms with the risk and severity of chronic heart failure: a case-control study of a northern Chinese population. Int J Mol Sci. 2015;16(2):2732–2746. doi:10.3390/ijms16022732

388. Wilke RA, Simpson RU, Mukesh BN, Bhupathi SV, Dart RA, Ghebranious NR, McCarty CA. Genetic variation in CYP27B1 is associated with congestive heart failure in patients with hypertension. Pharmacogenomics. 2009;10(11):1789–1797. doi:10.2217/pgs.09.101

389. Bharath G, Vishnuprabu DP, Preethi L, Nagappan AS, Dhianeshwaran Isravanya RT, Bhaskar LV, Swaminathan N, Munirajan AK. SLCO1B1 and ABCB1 variants synergistically influence the atorvastatin treatment response in South Indian coronary artery disease patients. Pharmacogenomics. 2022;23(12):683–694. doi:10.2217/pgs-2022-0044

390. Ray BK, Shakya A, Turk JR, Apte SS, Ray A. Induction of the MMP-14 gene in macrophages of the atherosclerotic plaque: role of SAF-1 in the induction process. Circ Res. 2004;95(11):1082–1090. doi:10.1161/01.RES.0000150046.48115.80

391. Zhang K, Wu M, Qin X, Wen P, Wu Y, Zhuang J. Asporin is a Potential Promising Biomarker for Common Heart Failure. DNA Cell Biol. 2021;40(2):303–315. doi:10.1089/dna.2020.5995

392. Guo Z, Hu YH, Feng GS, Valenzuela Ripoll C, Li ZZ, Cai SD, Wang QQ, Luo WW, Li Q, Liang LY, et al. JMJD6 protects against isoproterenol-induced cardiac hypertrophy via inhibition of NF-κB activation by demethylating R149 of the p65 subunit. Acta Pharmacol Sin. 2023;44(9):1777–1789. doi:10.1038/s41401-023-01086-7

393. Chaptal MC, Maraninchi M, Musto G, Mancini J, Chtioui H, Dupont-Roussel J, Marlinge M, Fromonot J, Lalevee N, Mourre F, et al. Low Density Lipoprotein Cholesterol Decreases the Expression of Adenosine A2A Receptor and Lipid Rafts-Protein Flotillin-1: Insights on Cardiovascular Risk of Hypercholesterolemia. Cells. 2024;13(6):488. doi:10.3390/cells13060488

394. Padua MB, Helm BM, Wells JR, Smith AM, Bellchambers HM, Sridhar A, Ware SM. Congenital heart defects caused by FOXJ1. Hum Mol Genet. 2023;32(14):2335–2346. doi:10.1093/hmg/ddad065

395. Wei W, Li B, Li F, Sun K, Jiang X, Xu R. Variants in FOXC1 and FOXC2 identified in patients with conotruncal heart defects. Genomics. 2024;116(3):110840. doi:10.1016/j.ygeno.2024.110840

396. Jin D, Han F. FOXF1 ameliorates angiotensin II-induced cardiac fibrosis in cardiac fibroblasts through inhibiting the TGF-β1/Smad3 signaling pathway. J Recept Signal Transduct Res. 2020;40(6):493–500. doi:10.1080/10799893.2020.1772299

397. Wang MX, Liu X, Li JM, Liu L, Lu W, Chen GC. Inhibition of CACNA1H can alleviate endoplasmic reticulum stress and reduce myocardial cell apoptosis caused by myocardial infarction. Eur Rev Med Pharmacol Sci. 2020;24(24):12887–12895. doi:10.26355/eurrev_202012_24192

398. Sjaarda J, Gerstein H, Chong M, Yusuf S, Meyre D, Anand SS, Hess S, Paré G. Blood CSF1 and CXCL12 as Causal Mediators of Coronary Artery Disease. J Am Coll Cardiol. 2018;72(3):300–310. doi:10.1016/j.jacc.2018.04.067

399. Vicen M, Igreja Sá IC, Tripská K, Vitverová B, Najmanová I, Eissazadeh S, Micuda S, Nachtigal P. Membrane and soluble endoglin role in cardiovascular and metabolic disorders related to metabolic syndrome. Cell Mol Life Sci. 2021;78(6):2405–2418. doi:10.1007/s00018-020-03701-w

400. Theis JL, Niaz T, Sundsbak RS, Fogarty ZC, Bamlet WR, Hagler DJ, Olson TM. CELSR1 Risk Alleles in Familial Bicuspid Aortic Valve and Hypoplastic Left Heart Syndrome. Circ Genom Precis Med. 2022;15(2):e003523. doi:10.1161/CIRCGEN.121.003523

401. Alaei Z, Zamani N, Rabbani B, Mahdieh N. TCAP gene is not a common cause of cardiomyopathy in Iranian patients. Eur J Med Res. 2023;28(1):376. doi:10.1186/s40001-023-01019-4

402. Huang Y, Li T, Gao S, Li S, Zhu X, Li Y, Liu D, Li W, Yang L, Liu K, et al. Investigating the role of NPR1 in dilated cardiomyopathy and its potential as a therapeutic target for glucocorticoid therapy. Front Pharmacol. 2023;14:1290253. doi:10.3389/fphar.2023.1290253

403. Fonseca J, Tomada N, Magalhães A, Rodrigues AR, Gouveia AM, Neves D. Effect of aging and cardiovascular risk factors on receptor Tie1 expression in human erectile tissue. J Sex Med. 2015;12(4):876–886. doi:10.1111/jsm.12794

404. Scotland RL, Allen L, Hennings LJ, Post GR, Post SR. The ral exchange factor rgl2 promotes cardiomyocyte survival and inhibits cardiac fibrosis. PLoS One. 2013;8(9):e73599. doi:10.1371/journal.pone.0073599

405. Accornero F, van Berlo JH, Benard MJ, Lorenz JN, Carmeliet P, Molkentin JD. Placental growth factor regulates cardiac adaptation and hypertrophy through a paracrine mechanism. Circ Res. 2011;109(3):272–280. doi:10.1161/CIRCRESAHA.111.240820

406. Edin ML, Hamedani BG, Gruzdev A, Graves JP, Lih FB, Arbes SJ 3rd Singh R, Orjuela Leon AC, Bradbury JA, DeGraff LM, et al. Epoxide hydrolase 1 (EPHX1) hydrolyzes epoxyeicosanoids and impairs cardiac recovery after ischemia. J Biol Chem. 2018;293(9):3281–3292. doi:10.1074/jbc.RA117.000298

407. Bartoli F, Sabourin J. Cardiac Remodeling and Disease: Current Understanding of STIM1/Orai1-Mediated Store-Operated Ca2+ Entry in Cardiac Function and Pathology. Adv Exp Med Biol. 2017;993:523–534. doi:10.1007/978-3-319-57732-6_26

408. Zhou JJ, Li H, Qian YL, Quan RL, Chen XX, Li L, Li Y, Wang PH, Meng XM, Jing XL, et al. Nestin represents a potential marker of pulmonary vascular remodeling in pulmonary arterial hypertension associated with congenital heart disease. J Mol Cell Cardiol. 2020;149:41–53. doi:10.1016/j.yjmcc.2020.09.005

409. Mavroidis M, Athanasiadis NC, Rigas P, Kostavasili I, Kloukina I, Te Rijdt WP, Kavantzas N, Chaniotis D, van Tintelen JP, Skaliora I, et al.Desmin is essential for the structure and function of the sinoatrial node: implications for increased arrhythmogenesis. Am J Physiol Heart Circ Physiol. 2020;319(3):H557–H570. doi:10.1152/ajpheart.00594.2019

410. Wu Y, Huang T, Li X, Shen C, Ren H, Wang H, Wu T, Fu X, Deng S, Feng Z, et al. Retinol dehydrogenase 10 reduction mediated retinol metabolism disorder promotes diabetic cardiomyopathy in male mice. Nat Commun. 2023;14(1):1181. doi:10.1038/s41467-023-36837-x

411. Tulbah S, Alruwaili N, Alhashem A, Aljohany A, Alhadeq F, Brotons DCA, Alwadai A, Al-Hassnan ZN. Variable phenotype of a null PPP1R13L allele in children with dilated cardiomyopathy. Am J Med Genet A. 2024;194(1):59–63. doi:10.1002/ajmg.a.63402

412. Fan M, Yang K, Wang X, Wang Y, Tu F, Ha T, Liu L, Williams DL, Li C. Endothelial cell HSPA12B and yes-associated protein cooperatively regulate angiogenesis following myocardial infarction. JCI Insight. 2020;5(18):e139640. doi:10.1172/jci.insight.139640

413. Wei X, Ma X, Lu R, Bai G, Zhang J, Deng R, Gu N, Feng N, Guo X. Genetic variants in PCSK1 gene are associated with the risk of coronary artery disease in type 2 diabetes in a Chinese Han population: a case control study. PLoS One. 2014;9(1):e87168. doi:10.1371/journal.pone.0087168

414. Bai S, Wang X, Hou Y, Cui Y, Song Q, Du J, Zhang Y, Xu J. ncRNA-056298 Regulates GAP43 and Promotes Cardiac Intrinsic Autonomic Nerve Remodelling in a Canine Model of Atrial Fibrillation Induction after Ganglionated Plexus Ablation. Curr Med Chem. 2024. doi:10.2174/0109298673289298240129103537

415. Zhou G, Fan L, Li Z, Li J, Kou X, Xiao M, Gao M, Qu X. G protein-coupled receptor MAS1 induces an inhibitory effect on myocardial infarction-induced myocardial injury. Int J Biol Macromol. 2022;207:72–80. doi:10.1016/j.ijbiomac.2022.02.163

416. Li Y, Song D, Mao L, Abraham DM, Bursac N. Lack of Thy1 defines a pathogenic fraction of cardiac fibroblasts in heart failure. Biomaterials. 2020;236:119824. doi:10.1016/j.biomaterials.2020.119824

417. Frasier CR, Wagnon JL, Bao YO, McVeigh LG, Lopez-Santiago LF, Meisler MH, Isom LL. Cardiac arrhythmia in a mouse model of sodium channel SCN8A epileptic encephalopathy. Proc Natl Acad Sci U S A. 2016;113(45):12838–12843. doi:10.1073/pnas.1612746113

418. Borges JI, Suster MS, Lymperopoulos A. Cardiac RGS Proteins in Human Heart Failure and Atrial Fibrillation: Focus on RGS4. Int J Mol Sci. 2023;24(7):6136. doi:10.3390/ijms24076136

419. Wingert J, Meinhardt E, Sasipong N, Pott M, Lederer C, de la Torre C, Sticht C, Most P, Katus HA, Frey N, et al. Cardiomyocyte-specific RXFP1 overexpression protects against pressure overload-induced cardiac dysfunction independently of relaxin. Biochem Pharmacol. 2024;225:116305. doi:10.1016/j.bcp.2024.116305

420. Loi M, Bastianini S, Candini G, Rizzardi N, Medici G, Papa V, Gennaccaro L, Mottolese N, Tassinari M, Uguagliati B, et al. Cardiac Functional and Structural Abnormalities in a Mouse Model of CDKL5 Deficiency Disorder. Int J Mol Sci. 2023;24(6):5552. doi:10.3390/ijms24065552

421. Duong T, Austin TR, Brody JA, Shojaie A, Battle A, Bader JS, Hong YS, Ballantyne CM, Coresh J, Gerszten RE, et al. Circulating Blood Plasma Profiling Reveals Proteomic Signature and a Causal Role for SVEP1 in Sudden Cardiac Death. Circ Genom Precis Med. 2024;17(5):e004494. doi:10.1161/CIRCGEN.123.004494

422. Lu QB, Fu X, Liu Y, Wang ZC, Liu SY, Li YC, Sun HJ. Disrupted cardiac fibroblast BCAA catabolism contributes to diabetic cardiomyopathy via a periostin/NAP1L2/SIRT3 axis. Cell Mol Biol Lett. 2023;28(1):93. doi:10.1186/s11658-023-00510-4

423. Accornero F, Schips TG, Petrosino JM, Gu SQ, Kanisicak O, van Berlo JH, Molkentin JD. BEX1 is an RNA-dependent mediator of cardiomyopathy. Nat Commun. 2017;8(1):1875. doi:10.1038/s41467-017-02005-1

424. Shi HJ, Wang MW, Sun JT, Wang H, Li YF, Chen BR, Fan Y, Wang SB, Wang ZM, Wang QM, et al. A novel long noncoding RNA FAF inhibits apoptosis via upregulating FGF9 through PI3K/AKT signaling pathway in ischemia-hypoxia cardiomyocytes. J Cell Physiol. 2019;234(12):21973–21987. doi:10.1002/jcp.28760

425. Kulkarni S, Lenin M, Ramesh R, Delphine SCW, Velu K. Evaluation of Single-Nucleotide Polymorphisms of Transcription Factor 7-Like 2 and ATP2B1 Genes as Cardiovascular Risk Predictors in Chronic Kidney Disease. Int J Appl Basic Med Res. 2019;9(4):221–225. doi:10.4103/ijabmr.IJABMR_92_19

426. Pang Y, Xu Y, Chen Q, Cheng K, Ling Y, Jang J, Ge J, Zhu W. FLRT3 and TGF-β/SMAD4 signalling: Impacts on apoptosis, autophagy and ion channels in supraventricular tachycardia. J Cell Mol Med. 2024;28(7):e18237. doi:10.1111/jcmm.18237

427. Li Q, Zhao Y, Wu G, Chen S, Zhou Y, Li S, Zhou M, Fan Q, Pu J, Hong K, et al. De Novo FGF12 (Fibroblast Growth Factor 12) Functional Variation Is Potentially Associated With Idiopathic Ventricular Tachycardia. J Am Heart Assoc. 2017;6(8):e006130. doi:10.1161/JAHA.117.006130

428. Pereira AHM, Cardoso AC, Consonni SR, Oliveira RR, Saito A, Vaggione MLB, Matos-Souza JR, Carazzolle MF, Gonçalves A, Fernandes JL, et al. MEF2C repressor variant deregulation leads to cell cycle re-entry and development of heart failure. EBioMedicine. 2020;51:102571. doi:10.1016/j.ebiom.2019.11.032

429. Huang C, Zhang X, Wang S, Shen A, Xu T, Hou Y, Gao S, Xie Y, Zeng Y, Chen J, et al. PARP-2 mediates cardiomyocyte aging and damage induced by doxorubicin through SIRT1 Inhibition. Apoptosis. 2024;29(5-6):816–834. doi:10.1007/s10495-023-01929-y

430. Wang Y, Wang S, Lei M, Boyett M, Tsui H, Liu W, Wang X. The p21-activated kinase 1 (Pak1) signalling pathway in cardiac disease: from mechanistic study to therapeutic exploration. Br J Pharmacol. 2018;175(8):1362–1374. doi:10.1111/bph.13872

431. Yan T, Song S, Sun W, Ge Y. HAPLN1 knockdown inhibits heart failure development via activating the PKA signaling pathway. BMC Cardiovasc Disord. 2024;24(1):197. doi:10.1186/s12872-024-03861-8

432. Hermida A, Jedraszak G, Ader F, Denjoy I, Fressart V, Maury P, Beyls C, Bloch A, Clerici G, Daire E, et al. Systematic analysis of SCN5A variants associated with inherited cardiac diseases. Heart Rhythm. 2024. doi:10.1016/j.hrthm.2024.08.018

433. Yang B, Zhao Y, Luo W, Zhu W, Jin L, Wang M, Ye L, Wang Y, Liang G. Macrophage DCLK1 promotes obesity-induced cardiomyopathy via activating RIP2/TAK1 signaling pathway. Cell Death Dis. 2023;14(7):419. doi:10.1038/s41419-023-05960-4

434. Shen J, Shentu J, Zhong C, Huang Q, Duan S. RNA splicing factor RBFOX2 is a key factor in the progression of cancer and cardiomyopathy. Clin Transl Med. 2024;14(9):e1788. doi:10.1002/ctm2.1788

435. Li X, Zheng S, Tan W, Chen H, Li X, Wu J, Luo T, Ren X, Pyle WG, Wang L, et al. Slit2 Protects Hearts Against Ischemia-Reperfusion Injury by Inhibiting Inflammatory Responses and Maintaining Myofilament Contractile Properties. Front Physiol. 2020;11:228. doi:10.3389/fphys.2020.00228

436. Ruiz-Velasco A, Raja R, Chen X, Ganenthiran H, Kaur N, Alatawi NHO, Miller JM, Abouleisa RRE, Ou Q, Zhao X, et al. Restored autophagy is protective against PAK3-induced cardiac dysfunction. iScience. 2023;26(6):106970. doi:10.1016/j.isci.2023.106970

437. Geng B, Wang X, Park KH, Lee KE, Kim J, Chen P, Zhou X, Tan T, Yang C, Zou X, et al. UCHL1 protects against ischemic heart injury via activating HIF-1α signal pathway. Redox Biol. 2022;52:102295. doi:10.1016/j.redox.2022.102295

438. Xiang M, Luo H, Wu J, Ren L, Ding X, Wu C, Chen J, Chen S, Zhang H, Yu L, et al. ADAM23 in Cardiomyocyte Inhibits Cardiac Hypertrophy by Targeting FAK - AKT Signaling. J Am Heart Assoc. 2018;7(18):e008604. doi:10.1161/JAHA.118.008604

439. Xie Y, Li Y, Chen J, Ding H, Zhang X. Early growth response-1: Key mediators of cell death and novel targets for cardiovascular disease therapy. Front Cardiovasc Med. 2023;10:1162662. doi:10.3389/fcvm.2023.1162662

440. Wang X, Guan Z, Tang W, Wang X, Xu C, Shan E, Wang W, Gao Y. PAX5/ITGAX Contributed to the Progression of Atherosclerosis by Regulation of B Differentiation via TNF-α Signaling Pathway. DNA Cell Biol. 2023;42(2):97–104. doi:10.1089/dna.2022.0461

441. Yaqoob H, Ahmad H, Ali SI, Patel N, Arif A. Missense mutations in the CITED2 gene may contribute to congenital heart disease. BMC Cardiovasc Disord. 2024;24(1):516. doi:10.1186/s12872-024-04035-2

442. Barefield DY, Yamakawa S, Tahtah I, Sell JJ, Broman M, Laforest B, Harris S, Alvarez-Arce A, Araujo KN, Puckelwartz MJ, et al. Partial and complete loss of myosin binding protein H-like cause cardiac conduction defects. J Mol Cell Cardiol. 2022;169:28–40. doi:10.1016/j.yjmcc.2022.04.012

443. Liu Y, Chen L, Gao L, Pei X, Tao Z, Xu Y, Li R. LRRK2 deficiency protects the heart against myocardial infarction injury in mice via the P53/HMGB1 pathway. Free Radic Biol Med. 2022;191:119–127. doi:10.1016/j.freeradbiomed.2022.08.035

444. Lin YJ, Chang JS, Liu X, Tsang H, Chien WK, Chen JH, Hsieh HY, Hsueh KC, Shiao YT, Li JP, et al. Genetic variants in PLCB4/PLCB1 as susceptibility loci for coronary artery aneurysm formation in Kawasaki disease in Han Chinese in Taiwan. Sci Rep. 2015;5:14762. doi:10.1038/srep14762

445. Dobrev D, Wehrens XH. Role of RyR2 phosphorylation in heart failure and arrhythmias: Controversies around ryanodine receptor phosphorylation in cardiac disease. Circ Res. 2014;114(8):1311–1319. doi:10.1161/CIRCRESAHA.114.300568

446. Künzel SR, Hoffmann M, Weber S, Künzel K, Kämmerer S, Günscht M, Klapproth E, Rausch JSE, Sadek MS, Kolanowski T, et al. Diminished PLK2 Induces Cardiac Fibrosis and Promotes Atrial Fibrillation. Circ Res. 2021;129(8):804–820. doi:10.1161/CIRCRESAHA.121.319425

447. Cheng X, Chen X, Zhang M, Wan Y, Ge S, Cheng X. Sparcl1 and Atherosclerosis. J Inflamm Res. 2023;16:2121–2127. doi:10.2147/JIR.S406907

448. Moya-Mendez ME, Ogbonna C, Ezekian JE, Rosamilia MB, Prange L, de la Uz C, Kim JJ, Howard T, Garcia J, Nussbaum R, et al. ATP1A3-Encoded Sodium-Potassium ATPase Subunit Alpha 3 D801N Variant Is Associated With Shortened QT Interval and Predisposition to Ventricular Fibrillation Preceded by Bradycardia. J Am Heart Assoc. 2021;10(17):e019887. doi:10.1161/JAHA.120.019887

449. Noori MR, Zhang B, Pan L. Is KCNH1 mutation related to coronary artery ectasia. BMC Cardiovasc Disord. 2019;19(1):296. doi:10.1186/s12872-019-01276-4

450. Calvert JW, Gundewar S, Yamakuchi M, Park PC, Baldwin WM 3rd, Lefer DJ, Lowenstein CJ. Inhibition of N-ethylmaleimide-sensitive factor protects against myocardial ischemia/reperfusion injury. Circ Res. 2007;101(12):1247–1254. doi:10.1161/CIRCRESAHA.107.162610

451. Gurunathan S, Sebastian J, Baker J, Abdel-Hamid HZ, West SC, Feingold B, Peche V, Reyes-Múgica M, Madan-Khetarpal S, Field J. A homozygous CAP2 pathogenic variant in a neonate presenting with rapidly progressive cardiomyopathy and nemaline rods. Am J Med Genet A. 2022;188(3):970–977. doi:10.1002/ajmg.a.62590

452. Golimbet VE, Volel’ BA, Dolzhikov AV, Korovaitseva GI, Isaeva MI. Association of 5-HTR2A and 5-HTR2C serotonin receptor gene polymorphisms with depression risk in patients with coronary heart disease. Bull Exp Biol Med. 2014;156(5):680–683. doi:10.1007/s10517-014-2424-1

453. Kang K, Li J, Li R, Xu X, Liu J, Qin L, Huang T, Wu J, Jiao M, Wei M, et al. Potentially Critical Roles of NDUFB5, TIMMDC1, and VDAC3 in the Progression of Septic Cardiomyopathy Through Integrated Bioinformatics Analysis. DNA Cell Biol. 2020;39(1):105–117. doi:10.1089/dna.2019.4859

454. Kušíková K, Feichtinger RG, Csillag B, Kalev OK, Weis S, Duba HC, Mayr JA, Weis D. Case Report and Review of the Literature: A New and a Recurrent Variant in the VARS2 Gene Are Associated With Isolated Lethal Hypertrophic Cardiomyopathy, Hyperlactatemia, and Pulmonary Hypertension in Early Infancy. Front Pediatr. 2021;9:660076. doi:10.3389/fped.2021.660076

455. Pfister O, Lorenz V, Oikonomopoulos A, Xu L, Häuselmann SP, Mbah C, Kaufmann BA, Liao R, Wodnar-Filipowicz A, Kuster GM. FLT3 activation improves post-myocardial infarction remodeling involving a cytoprotective effect on cardiomyocytes. J Am Coll Cardiol. 2014;63(10):1011–1019. doi:10.1016/j.jacc.2013.08.1647

456. Xiong F, Mao R, Zhao R, Zhang L, Tan K, Liu C, Wang S, Xu M, Li Y, et al. Plasma Exosomal S1PR5 and CARNS1 as Potential Non-invasive Screening Biomarkers of Coronary Heart Disease. Front Cardiovasc Med. 2022;9:845673. doi:10.3389/fcvm.2022.845673

457. Shen JF, Fan ZB, Wu CW, Qi GX, Cao QY, Xu F. Sacubitril Valsartan Enhances Cardiac Function and Alleviates Myocardial Infarction in Rats through a SUV39H1/SPP1 Axis. Oxid Med Cell Longev. 2022;2022:5009289. doi:10.1155/2022/5009289

458. Chen S, Zhang Y, Lighthouse JK, Mickelsen DM, Wu J, Yao P, Small EM, Yan C. A Novel Role of Cyclic Nucleotide Phosphodiesterase 10A in Pathological Cardiac Remodeling and Dysfunction. Circulation. 2020;141(3):217–233. doi:10.1161/CIRCULATIONAHA.119.042178

459. Lu Z, Cui Y, Wei X, Gao P, Zhang H, Wei X, Li Q, Sun F, Yan Z, Zheng H, et al. Deficiency of PKD2L1 (TRPP3) Exacerbates Pathological Cardiac Hypertrophy by Augmenting NCX1-Mediated Mitochondrial Calcium Overload. Cell Rep. 2018;24(6):1639–1652. doi:10.1016/j.celrep.2018.07.022

460. Zhuo C, Jiang R, Lin X, Shao M. LncRNA H19 inhibits autophagy by epigenetically silencing of DIRAS3 in diabetic cardiomyopathy. Oncotarget. 2017;8(1):1429–1437. doi:10.18632/oncotarget.13637

461. Hayderi A, Kumawat AK, Shavva VS, Dreifaldt M, Sigvant B, Petri MH, Kragsterman B, Olofsson PS, Sirsjö A, Ljungberg LU. RSAD2 is abundant in atherosclerotic plaques and promotes interferon-induced CXCR3-chemokines in human smooth muscle cells. Sci Rep. 2024;14(1):8196. doi:10.1038/s41598-024-58592-9

462. Mayer O Jr, Seidlerová J, Černá V, Kučerová A, Bruthans J, Vágovičová P, Vaněk J, Timoracká K, Wohlfahrt P, Filipovský J, et al. The DRD2/ANKK1 Taq1A polymorphism is associated with smoking cessation failure in patients with coronary heart disease. Per Med. 2015;12(5):463–473. doi:10.2217/pme.15.16

463. Li H, Lewis A, Brodsky S, Rieger R, Iden C, Goligorsky MS. Homocysteine induces 3-hydroxy-3-methylglutaryl coenzyme a reductase in vascular endothelial cells: a mechanism for development of atherosclerosis?. Circulation. 2002;105(9):1037–1043. doi:10.1161/hc0902.104713

464. Du Z, Kuang S, Li Y, Han P, Liu J, Wang Z, Huang Y, Guan Y, Xu X, Liu X, et al. Family-Based Whole Genome Sequencing Identified Novel Variants in ABCA5 Gene in a Patient with Idiopathic Ventricular Tachycardia. Pediatr Cardiol. 2020;41(8):1783–1794. doi:10.1007/s00246-020-02446-4

465. Ma T, Lin S, Wang B, Wang Q, Xia W, Zhang H, Cui Y, He C, Wu H, Sun F, et al. TRPC3 deficiency attenuates high salt-induced cardiac hypertrophy by alleviating cardiac mitochondrial dysfunction. Biochem Biophys Res Commun. 2019;519(4):674–681. doi:10.1016/j.bbrc.2019.09.018

466. Li B, Syed MH, Khan H, Singh KK, Qadura M. The Role of Fatty Acid Binding Protein 3 in Cardiovascular Diseases. Biomedicines. 2022;10(9):2283. doi:10.3390/biomedicines10092283

467. Tam CHT, Lim CKP, Luk AOY, Shi M, Man Cheung H, Ng ACW, Lee HM, Lau ESH, Fan B, Jiang G, et al. Identification of a Common Variant for Coronary Heart Disease at PDE1A Contributes to Individualized Treatment Goals and Risk Stratification of Cardiovascular Complications in Chinese Patients With Type 2 Diabetes. Diabetes Care. 2023;46(6):1271–1281. doi:10.2337/dc22-2331

468. Lei W, Yiming S, Qiang P, Xin C, Peng G, Baofeng Z. Unleashing the Neurotherapeutic Potential: The Crucial Role of miR-206-3p in Facilitating Hsp90aa1-Mediated Central Nervous System Injuries During Heat Stroke. Mol Neurobiol. 2024. doi:10.1007/s12035-024-04342-x

469. Li D, Ji JX, Xu YT, Ni HB, Rui Q, Liu HX, Jiang F, Gao R, Chen G. Inhibition of Lats1/p-YAP1 pathway mitigates neuronal apoptosis and neurological deficits in a rat model of traumatic brain injury. CNS Neurosci Ther. 2018;24(10):906–916. doi:10.1111/cns.12833

470. Zhang Z, Zhang X, Wu X, Zhang Y, Lu J, Li D. Sirt1 attenuates astrocyte activation via modulating Dnajb1 and chaperone-mediated autophagy after closed head injury. Cereb Cortex. 2022;32(22):5191–5205. doi:10.1093/cercor/bhac007

471. Amirkhosravi L, Khaksari M, Sheibani V, Shahrokhi N, Ebrahimi MN, Amiresmaili S, Salmani N. Improved spatial memory, neurobehavioral outcomes, and neuroprotective effect after progesterone administration in ovariectomized rats with traumatic brain injury: Role of RU486 progesterone receptor antagonist. Iran J Basic Med Sci. 2021;24(3):349–359. doi:10.22038/ijbms.2021.50973.11591

472. Thundyil J, Manzanero S, Pavlovski D, Cully TR, Lok KZ, Widiapradja A, Chunduri P, Jo DG, Naruse C, Asano M, et al. Evidence that the EphA2 receptor exacerbates ischemic brain injury. PLoS One. 2013;8(1):e53528. doi:10.1371/journal.pone.0053528

473. Huang Z, Liu J, Xu J, Dai L, Wang H. Downregulation of miR-26b attenuates early brain injury induced by subarachnoid hemorrhage via mediating the KLF4/STAT3/HMGB1 axis. Exp Neurol. 2023;359:114270. doi:10.1016/j.expneurol.2022.114270

474. Dai Y, Hu L. HSPB1 overexpression improves hypoxic-ischemic brain damage by attenuating ferroptosis in rats through promoting G6PD expression. J Neurophysiol. 2022;128(6):1507–1517. doi:10.1152/jn.00306.2022

475. Krum JM, Mani N, Rosenstein JM. Roles of the endogenous VEGF receptors flt-1 and flk-1 in astroglial and vascular remodeling after brain injury. Exp Neurol. 2008;212(1):108–117. doi:10.1016/j.expneurol.2008.03.019

476. Luo J, Wu X, Liu H, Cui W, Guo W, Guo K, Guo H, Tao K, Li F, Shi Y, et al. Antagonism of Protease-Activated Receptor 4 Protects Against Traumatic Brain Injury by Suppressing Neuroinflammation via Inhibition of Tab2/NF-κB Signaling. Neurosci Bull. 2021;37(2):242–254. doi:10.1007/s12264-020-00601-8

477. Li Y, Liu C, Fan H, Du Y, Zhang R, Zhan S, Zhang G, Bu N. Gli2-induced lncRNA Peg13 alleviates cerebral ischemia-reperfusion injury by suppressing Yy1 transcription in a PRC2 complex-dependent manner. Metab Brain Dis. 2023;38(4):1389–1404. doi:10.1007/s11011-023-01159-w

478. Zhang AQ, Wang L, Wang YX, Hong SS, Zhong YS, Yu RY, Wu XL, Zhou BB, Yu QM, Fu HF, et al. Silencing miRNA-324-3p protects against cerebral ischemic injury via regulation of the GATA2/A1R axis. Neural Regen Res. 2022;17(11):2504–2511. doi:10.4103/1673-5374.339009

479. Liu N, Han J, Li Y, Jiang Y, Shi SX, Lok J, Whalen M, Dumont AS, Wang X. Recombinant annexin A2 inhibits peripheral leukocyte activation and brain infiltration after traumatic brain injury. J Neuroinflammation. 2021;18(1):173. doi:10.1186/s12974-021-02219-7

480. Li S, Qu X, Qin Z, Gao J, Li J, Liu J. lncfos/miR-212-5p/CASP7 Axis-Regulated miR-212-5p Protects the Brain Against Ischemic Damage. Mol Neurobiol. 2023;60(5):2767–2785. doi:10.1007/s12035-023-03216-y

481. Kumari D, Kaur S, Dandekar MP. Intricate Role of the Cyclic Guanosine Monophosphate Adenosine Monophosphate Synthase-Stimulator of Interferon Genes (cGAS-STING) Pathway in Traumatic Brain Injury-Generated Neuroinflammation and Neuronal Death. ACS Pharmacol Transl Sci. 2024;7(10):2936–2950. doi:10.1021/acsptsci.4c00310

482. Mao W, Yi X, Qin J, Tian M, Jin G. CXCL12/CXCR4 Axis Improves Migration of Neuroblasts Along Corpus Callosum by Stimulating MMP-2 Secretion After Traumatic Brain Injury in Rats. Neurochem Res. 2016;41(6):1315–1322. doi:10.1007/s11064-016-1831-2

483. Chen Z, Hu Y, Lu R, Ge M, Zhang L. MicroRNA-374a-5p inhibits neuroinflammation in neonatal hypoxic-ischemic encephalopathy via regulating NLRP3 inflammasome targeted Smad6. Life Sci. 2020;252:117664. doi:10.1016/j.lfs.2020.117664

484. Niu LJ, Xu RX, Zhang P, Du MX, Jiang XD. Suppression of Frizzled-2-mediated Wnt/Ca² signaling significantly attenuates intracellular calcium accumulation in vitro and in a rat model of traumatic brain injury. Neuroscience. 2012;213:19–28. doi:10.1016/j.neuroscience.2012.03.057

485. Chernyak YI, Itskovich VB, D’yakovich OA, Kolesnikov SI. Role of cytochrome P450-dependent monooxygenases and polymorphic variants of GSTT1 and GSTM1 genes in the formation of brain lesions in individuals chronically exposed to mercury. Bull Exp Biol Med. 2013;156(1):15–18. doi:10.1007/s10517-013-2266-2

486. Chen Y, Wang Y, Xu J, Hou T, Zhu J, Jiang Y, Sun L, Huang C, Sun L, Liu S. Multiplex Assessment of Serum Chemokines CCL2, CCL5, CXCL1, CXCL10, and CXCL13 Following Traumatic Brain Injury. Inflammation. 2023;46(1):244–255. doi:10.1007/s10753-022-01729-7

487. Li HY, Li P, Yang HG, Yao QQ, Huang SN, Wang JQ, Zheng N. Investigation and comparison of the protective activities of three functional proteins-lactoferrin, α-lactalbumin, and β-lactoglobulin-in cerebral ischemia reperfusion injury. J Dairy Sci. 2020;103(6):4895–4906. doi:10.3168/jds.2019-17725

488. Jiao W, Jiang L, Zhang Y. SNHG1 alleviates the oxidative stress and inflammatory response in traumatic brain injury through regulating miR-377-3p/DUSP1 axis. Neuroreport. 2023;34(1):17–29. doi:10.1097/WNR.0000000000001852

489. Zhang Z, Trautmann K, Artelt M, Burnet M, Schluesener HJ. Bone morphogenetic protein-6 is expressed early by activated astrocytes in lesions of rat traumatic brain injury. Neuroscience. 2006;138(1):47–53. doi:10.1016/j.neuroscience.2005.11.036

490. Wang Y, Fang M, Ren Q, Qi W, Bai X, Amin N, Zhang X, Li Z, Zhang L. Sox17 protects human brain microvascular endothelial cells from AngII-induced injury by regulating autophagy and apoptosis. Mol Cell Biochem. 2024;479(9):2337–2350. doi:10.1007/s11010-023-04838-5

491. Wu Q, Liu G, Xu L, Wen X, Cai Y, Fan W, Yao X, Huang H, Li Q. Repair of Neurological Function in Response to FK506 Through CaN/NFATc1 Pathway Following Traumatic Brain Injury in Rats. Neurochem Res. 2016;41(10):2810–2818. doi:10.1007/s11064-016-1997-7

492. Zhou Y, Zhou B, Tu H, Tang Y, Xu C, Chen Y, Zhao Z, Miao Z. he degradation of mixed lineage kinase domain-like protein promotes neuroprotection after ischemic brain injury. Oncotarget. 2017;8(40):68393–68401. doi:10.18632/oncotarget.19416

493. Lipponen A, Paananen J, Puhakka N, Pitkänen A. Analysis of Post-Traumatic Brain Injury Gene Expression Signature Reveals Tubulins, Nfe2l2, Nfkb, Cd44, and S100a4 as Treatment Targets. Sci Rep. 2016;6:31570. doi:10.1038/srep31570

494. Mercurio D, Oggioni M, Fumagalli S, Lynch NJ, Roscher S, Minuta D, Perego C, Ippati S, Wallis R, Schwaeble WJ, et al. Targeted deletions of complement lectin pathway genes improve outcome in traumatic brain injury, with MASP-2 playing a major role. Acta Neuropathol Commun. 2020;8(1):174. doi:10.1186/s40478-020-01041-1

495. Zhao Y, Zhou YG, Chen JF. Targeting the adenosine A2A receptor for neuroprotection and cognitive improvement in traumatic brain injury and Parkinson’s disease. Chin J Traumatol. 2024;27(3):125–133. doi:10.1016/j.cjtee.2023.08.003

496. Willyerd FA, Empey PE, Philbrick A, Ikonomovic MD, Puccio AM, Kochanek PM, Okonkwo DO, Clark RS. Expression of ATP-Binding Cassette Transporters B1 and C1 after Severe Traumatic Brain Injury in Humans. J Neurotrauma. 2016;33(2):226–231. doi:10.1089/neu.2015.3879

497. Cui G, Yu Z, Li Z, Wang W, Lu T, Qian C, Li J, Ding Y. Increased expression of Foxj1 after traumatic brain injury. J Mol Neurosci. 2011;45(2):145–153. doi:10.1007/s12031-011-9504-8

498. He T, Shang J, Gao C, Guan X, Chen Y, Zhu L, Zhang L, Zhang C, Zhang J, Pang T. A novel SIRT6 activator ameliorates neuroinflammation and ischemic brain injury via EZH2/FOXC1 axis. Acta Pharm Sin B. 2021;11(3):708–726. doi:10.1016/j.apsb.2020.11.002

499. Li L, Yerra L, Chang B, Mathur V, Nguyen A, Luo J. Acute and late administration of colony stimulating factor 1 attenuates chronic cognitive impairment following mild traumatic brain injury in mice. Brain Behav Immun. 2021;94:274–288. doi:10.1016/j.bbi.2021.01.022

500. Wang LH, Zhang GL, Liu XY, et al. CELSR1 Promotes Neuroprotection in Cerebral Ischemic Injury Mainly Through the Wnt/PKC Signaling Pathway. Int J Mol Sci. 2020;21(4):1267. doi:10.3390/ijms21041267

501. Michinaga S, Hishinuma S, Koyama Y. Roles of Astrocytic Endothelin ETB Receptor in Traumatic Brain Injury. Cells. 2023;12(5):719. doi:10.3390/cells12050719

502. Frisén J, Johansson CB, Török C, Risling M, Lendahl U. Rapid, widespread, and longlasting induction of nestin contributes to the generation of glial scar tissue after CNS injury. J Cell Biol. 1995;131(2):453–464. doi:10.1083/jcb.131.2.453

503. Li HL, Guo RJ, Ai ZR, Han S, Guan Y, Li JF, Wang Y. Upregulation of Spinal MDGA1 in Rats After Nerve Injury Alters Interactions Between Neuroligin-2 and Postsynaptic Scaffolding Proteins and Increases GluR1 Subunit Surface Delivery in the Spinal Cord Dorsal Horn. Neurochem Res. 2024;49(2):507–518. doi:10.1007/s11064-023-04049-w

504. Ma Y, Lu C, Li C, Li R, Zhang Y, Ma H, Zhang X, Ding Z, Liu L. Overexpression of HSPA12B protects against cerebral ischemia/reperfusion injury via a PI3K/Akt-dependent mechanism. Biochim Biophys Acta. 2013;1832(1):57–66. doi:10.1016/j.bbadis.2012.10.003

505. Dan QQ, Ma Z, Tan YX, Visar B, Chen L. AQP4 knockout promotes neurite outgrowth via upregulating GAP43 expression in infant rats with hypoxic-ischemic brain injury. Ibrain. 2022;8(3):324–337. doi:10.1002/ibra.12062

506. Richter M, Negro-Demontel ML, Blanco-Ocampo D, Taranto E, Lago N, Peluffo H. Thy1-YFP-H Mice and the Parallel Rod Floor Test to Evaluate Short- and Long-Term Progression of Traumatic Brain Injury. Curr Protoc Immunol. 2018;120:24.1.1-24.1.25. doi:10.1002/cpim.42

507. Lin C, Zhao P, Sun G, Liu N, Ji J. SCG2 mediates blood-brain barrier dysfunction and schizophrenia-like behaviors after traumatic brain injury. FASEB J. 2024;38(17):e70016. doi:10.1096/fj.202401117R

508. Qiu L, He J, Chen H, Xu X, Tao Y. CircDLGAP4 overexpression relieves oxygen-glucose deprivation-induced neuronal injury by elevating NEGR1 through sponging miR-503-3p. J Mol Histol. 2022;53(2):321–332. doi:10.1007/s10735-021-10036-8

509. Zhang J, Liang L, Miao X, Wu S, Cao J, Tao B, Mao Q, Mo K, Xiong M, et al. Contribution of the Suppressor of Variegation 3-9 Homolog 1 in Dorsal Root Ganglia and Spinal Cord Dorsal Horn to Nerve Injury-induced Nociceptive Hypersensitivity. Anesthesiology. 2016;125(4):765–778. doi:10.1097/ALN.0000000000001261

510. Si W, Sun B, Luo J, Li Z, Dou Y, Wang Q. Snap25 attenuates neuronal injury via reducing ferroptosis in acute ischemic stroke. Exp Neurol. 2023;367:114476. doi:10.1016/j.expneurol.2023.114476

511. Wang L, Tan Y, Zhu Z, Chen J, Sun Q, Ai Z, Ai C, Xing Y, He G, Liu Y. ATP2B1-AS1 Promotes Cerebral Ischemia/Reperfusion Injury Through Regulating the miR-330-5p/TLR4-MyD88-NF-κB Signaling Pathway. Front Cell Dev Biol. 2021;9:720468. doi:10.3389/fcell.2021.720468

512. Xu J, Huang X, Liu S, Chen D, Xie Y, Zhao Z. The protective effects of lncRNA ZFAS1/miR-421-3p/MEF2C axis on cerebral ischemia-reperfusion injury. Cell Cycle. 2022;21(18):1915–1931. doi:10.1080/15384101.2022.2060627

513. Kofler J, Otsuka T, Zhang Z, Noppens R, Grafe MR, Koh DW, Dawson VL, de Murcia JM, Hurn PD, Traystman RJ. Differential effect of PARP-2 deletion on brain injury after focal and global cerebral ischemia. J Cereb Blood Flow Metab. 2006;26(1):135–141. doi:10.1038/sj.jcbfm.9600173

514. Huang M, Zhang J, Li M, Cao H, Zhu Q, Yang D. PAK1 contributes to cerebral ischemia/reperfusion injury by regulating the blood-brain barrier integrity. iScience. 2023;26(8):107333. doi:10.1016/j.isci.2023.107333

515. Gu X, Ni H, Kan X, Chen C, Zhou Z, Ding Z, Li D, Liu B. Nicotinamide Mononucleotide Adenylyl Transferase 2 Inhibition Aggravates Neurological Damage after Traumatic Brain Injury in a Rat Model. J Korean Neurosurg Soc. 2023;66(4):400–408. doi:10.3340/jkns.2022.0115

516. Zhang N, Yang L, Meng L, Cui H. Inhibition of miR-200b-3p alleviates hypoxia-ischemic brain damage via targeting Slit2 in neonatal rats. Biochem Biophys Res Commun. 2020;523(4):931–938. doi:10.1016/j.bbrc.2020.01.029

517. Yang M, Wang X, Fan Y, Chen Y, Sun D, Xu X, Wang J, Gu G, Peng R, Shen T, et al. Semaphorin 3A Contributes to Secondary Blood-Brain Barrier Damage After Traumatic Brain Injury. Front Cell Neurosci. 2019;13:117. doi:10.3389/fncel.2019.00117

518. Mi Z, Graham SH. Role of UCHL1 in the pathogenesis of neurodegenerative diseases and brain injury. Ageing Res Rev. 2023;86:101856. doi:10.1016/j.arr.2023.101856

519. Zhu J, Chen Z, Yu B, Zhang L, Ai F. MicroRNA-375-3p Alleviates Salicylate-Induced Neuronal Injury by Targeting ELAVL4 in Tinnitus. J Neurol Surg B Skull Base. 2023;85(3):227–233. doi:10.1055/s-0043-1764379

520. Li YY, Guo JH, Liu YQ, Dong JH, Zhu CH. PPARγ Activation-Mediated Egr-1 Inhibition Benefits Against Brain Injury in an Experimental Ischaemic Stroke Model. J Stroke Cerebrovasc Dis. 2020;29(12):105255. doi:10.1016/j.jstrokecerebrovasdis.2020.105255

521. Zhu G, Fang Y, Cui X, Jia R, Kang X, Zhao R. Magnolol upregulates CHRM1 to attenuate Amyloid-β-triggered neuronal injury through regulating the cAMP/PKA/CREB pathway. J Nat Med. 2022;76(1):188–199. doi:10.1007/s11418-021-01574-2

522. Mondello S, Gabrielli A, Catani S, D’Ippolito M, Jeromin A, Ciaramella A, Bossù P, Schmid K, Tortella F, Wang KK, et al. Increased levels of serum MAP-2 at 6-months correlate with improved outcome in survivors of severe traumatic brain injury. Brain Inj. 2012;26(13-14):1629–1635. doi:10.3109/02699052.2012.700083

523. An J, Yang H, Park SM, Chwae YJ, Joe EH. The LRRK2-G2019S mutation attenuates repair of brain injury partially by reducing the release of osteopontin-containing monocytic exosome-like vesicles. Neurobiol Dis. 2024;197:106528. doi:10.1016/j.nbd.2024.106528

524. Qi J, Meng C, Mo J, Shou T, Ding L, Zhi T. CircAFF2 Promotes Neuronal Cell Injury in Intracerebral Hemorrhage by Regulating the miR-488/CLSTN3 Axis. Neuroscience. 2023;535:75–87. doi:10.1016/j.neuroscience.2023.10.014

525. Yu Y, Xia Q, Zhan G, Gao S, Han T, Mao M, Li X, Wang Y. TRIM67 alleviates cerebral ischemia reperfusion injury by protecting neurons and inhibiting neuroinflammation via targeting IκBα for K63-linked polyubiquitination. Cell Biosci. 2023;13(1):99. doi:10.1186/s13578-023-01056-w

526. Kahriman A, Bouley J, Tuncali I, Dogan EO, Pereira M, Luu T, Bosco DA, Jaber S, Peters OM, Brown RH Jr, et al. Repeated mild traumatic brain injury triggers pathology in asymptomatic C9ORF72 transgenic mice. Brain. 2023;146(12):5139–5152. doi:10.1093/brain/awad264

527. Jacobsson G, Piehl F, Meister B. VAMP-1 and VAMP-2 gene expression in rat spinal motoneurones: differential regulation after neuronal injury. Eur J Neurosci. 1998;10(1):301–316. doi:10.1046/j.1460-9568.1998.00050.x

528. Ihbe N, Le Prieult F, Wang Q, Distler U, Sielaff M, Tenzer S, Thal SC, Mittmann T. Adaptive Mechanisms of Somatostatin-Positive Interneurons after Traumatic Brain Injury through a Switch of α Subunits in L-Type Voltage-Gated Calcium Channels. Cereb Cortex. 2022;32(5):1093–1109. doi:10.1093/cercor/bhab268

529. Hiltunen J, Ndode-Ekane XE, Lipponen A, Drexel M, Sperk G, Puhakka N, Pitkänen A. Regulation of Parvalbumin Interactome in the Perilesional Cortex after Experimental Traumatic Brain Injury. Neuroscience. 2021;475:52–72. doi:10.1016/j.neuroscience.2021.08.018

530. Huang HB, Yang SB, Shen LJ, Lv QW, Guo M, Zhou J, Li Z, Yang CS, Wang LY, Zhang H. A prospective study on serum secreted protein acidic and rich in cysteine-like 1 as a prognostic marker for severe traumatic brain injury. Clin Chim Acta. 2019;491:19–23. doi:10.1016/j.cca.2019.01.005

531. Qiu M, Zhao X, Guo T, He H, Deng Y. N-ethylmaleimide-sensitive factor elicits a neuroprotection against ischemic neuronal injury by restoring autophagic/lysosomal dysfunction. Cell Death Discov. 2024;10(1):368. doi:10.1038/s41420-024-02144-7

532. Fronczak KM, Li Y, Henchir J, Dixon CE, Carlson SW. Reductions in Synaptic Vesicle Glycoprotein 2 Isoforms in the Cortex and Hippocampus in a Rat Model of Traumatic Brain Injury. Mol Neurobiol. 2021;58(11):6006–6019. doi:10.1007/s12035-021-02534-3

533. Liu Z, Gao W, Xu Y. Eleutheroside E alleviates cerebral ischemia-reperfusion injury in a 5-hydroxytryptamine receptor 2C (Htr2c)-dependent manner in rats. Bioengineered. 2022;13(5):11718–11731. doi:10.1080/21655979.2022.2071009

534. Zhu X, Li H, You W, Yu Z, Wang Z, Shen H, Li X, Yu H, Wang Z, Chen G. Role of Rph3A in brain injury induced by experimental cerebral ischemia-reperfusion model in rats. CNS Neurosci Ther. 2022;28(7):1124–1138. doi:10.1111/cns.13850

535. Zhang J, Liang L, Miao X, Wu S, Cao J, Tao B, Mao Q, Mo K, Xiong M, Lutz BM, et al. Contribution of the Suppressor of Variegation 3-9 Homolog 1 in Dorsal Root Ganglia and Spinal Cord Dorsal Horn to Nerve Injury-induced Nociceptive Hypersensitivity. Anesthesiology. 2016;125(4):765–778. doi:10.1097/ALN.0000000000001261

536. Beker MC, Altintas MO, Dogan E, Bayraktaroglu C, Balaban B, Ozpinar A, Sengun N, Altunay S, Kilic E. Inhibition of phosphodiesterase 10A mitigates neuronal injury by modulating apoptotic pathways in cold-induced traumatic brain injury. Mol Cell Neurosci. 2024. doi:10.1016/j.mcn.2024.103977

537. Jiang W, Stingelin L, Zhang P, Tian X, Kang N, Liu J, Aihemaiti Y, Zhou D, Tu H. Enolase2 and enolase1 cooperate against neuronal injury in stroke model. Neurosci Lett. 2021;747:135662. doi:10.1016/j.neulet.2021.135662

538. Snider S, Albano L, Gagliardi F, Comai S, Roncelli F, De Domenico P, Pompeo E, Panni P, Bens N, Calvi MR, et al. Substantially elevated serum glutamate and CSF GOT-1 levels associated with cerebral ischemia and poor neurological outcomes in subarachnoid hemorrhage patients. Sci Rep. 2023;13(1):5246. doi:10.1038/s41598-023-32302-3

539. Yue JK, Pronger AM, Ferguson AR, Temkin NR, Sharma S, Rosand J, Sorani MD, McAllister TW, Barber J, Winkler EA, et al. Association of a common genetic variant within ANKK1 with six-month cognitive performance after traumatic brain injury. Neurogenetics. 2015;16(3):169–180. doi:10.1007/s10048-015-0437-1

540. Chen X, Lu M, He X, Ma L, Birnbaumer L, Liao Y. TRPC3/6/7 Knockdown Protects the Brain from Cerebral Ischemia Injury via Astrocyte Apoptosis Inhibition and Effects on NF-кB Translocation. Mol Neurobiol. 2017;54(10):7555–7566. doi:10.1007/s12035-016-0227-2

541. Zhong FF, Wei B, Bao GX, Lou YP, Wei ME, Wang XY, Xiao X, Tian JJ. FABP3 Induces Mitochondrial Autophagy to Promote Neuronal Cell Apoptosis in Brain Ischemia-Reperfusion Injury. Neurotox Res. 2024;42(4):35. doi:10.1007/s12640-024-00712-4

542. Winneberger J, Schöls S, Lessmann K, Rández-Garbayo J, Bauer AT, Mohamud Yusuf A, Hermann DM, Gunzer M, Schneider SW, Fiehler J, et al. Platelet endothelial cell adhesion molecule-1 is a gatekeeper of neutrophil transendothelial migration in ischemic stroke. Brain Behav Immun. 2021;93:277–287. doi:10.1016/j.bbi.2020.12.026

543. Rankinen T, Rice T, Boudreau A, Leon AS, Skinner JS, Wilmore JH, Rao DC, Bouchard C. Titin is a candidate gene for stroke volume response to endurance training: the HERITAGE Family Study. Physiol Genomics. 2003;15(1):27–33. doi:10.1152/physiolgenomics.00147.2002

544. Stetskaya TA, Krapiva AB, Kobzeva KA, Gurtovoy DE, Komkova GV, Polonikov AV, Bushueva OY. Polymorphism in Genes Encoding Adaptor Proteins ST13 and STIP1 and the Risk of Ischemic Stroke: a Pilot Study. Bull Exp Biol Med. 2024;176(4):477–480. doi:10.1007/s10517-024-06050-x

545. Wang C, Li L. The critical role of KLF4 in regulating the activation of A1/A2 reactive astrocytes following ischemic stroke. J Neuroinflammation. 2023;20(1):44. doi:10.1186/s12974-023-02742-9

546. Lindahl K, Rubin CJ, Brändström H, Karlsson MK, Holmberg A, Ohlsson C, Mellström D, Orwoll E, Mallmin H, Kindmark A, et al. Heterozygosity for a coding SNP in COL1A2 confers a lower BMD and an increased stroke risk. Biochem Biophys Res Commun. 2009;384(4):501–505. doi:10.1016/j.bbrc.2009.05.006

547. Li Z, Liu B, Lambertsen KL, Clausen BH, Zhu Z, Du X, Xu Y, Poulsen FR, Halle B, Bonde C, et al. USP25 Inhibits Neuroinflammatory Responses After Cerebral Ischemic Stroke by Deubiquitinating TAB2. Adv Sci (Weinh). 2023;10(28):e2301641. doi:10.1002/advs.202301641

548. Bierman-Chow S, Holland SM, Hsu AP, Palmer C, Lynch J, Mina Y, Joo Sophie Cho H. Clinical, Imaging, and Laboratory Findings in Patients With GATA2 Deficiency Presenting With Early-Onset Ischemic Stroke. Neurology. 2023;100(7):338–341. doi:10.1212/WNL.0000000000201569

549. Liu B, Xing Z, Song F, Li D, Zhao B, Xu C, Xia W, Ji H. METTL3-mediated ANXA2 inhibition confers neuroprotection in ischemic stroke: Evidence from in vivo and in vitro studies. FASEB J. 2023;37(7):e22974. doi:10.1096/fj.202300246R

550. Karimian M, Karimnia F. CYP1A1 common gene polymorphisms and ischemic stroke risk: a meta-analysis and a structural examination. Per Med. 2023;20(3):271–281. doi:10.2217/pme-2022-0113

551. Zheng Z, Liu S, Wang C, Wang C, Tang D, Shi Y, Han X. Association of genetic polymorphisms in CASP7 with risk of ischaemic stroke. Sci Rep. 2019;9(1):18627. doi:10.1038/s41598-019-55201-y

552. Chauhan C, Kaundal RK. The role of cGAS-STING signaling in ischemic stroke: From immune response to therapeutic targeting. Drug Discov Today. 2023;28(11):103792. doi:10.1016/j.drudis.2023.103792

553. Li S, Yang S, Zhang X, Zhang Y, Zhang J, Zhang X, Li W, Niu X, Shi W, Zhang G, et al. Impact of MMP2 rs243849 and rs14070 genetic polymorphisms on the ischemic stroke susceptibility in Chinese Shaanxi population. Front Neurol. 2022;13:931437. doi:10.3389/fneur.2022.931437

554. Wang R, Wang Y, Wang J, Yang K. Association of glutathione S- transferase T1 and M1 gene polymorphisms with ischemic stroke risk in the Chinese Han population. Neural Regen Res. 2012;7(18):1420–1427. doi:10.3969/j.issn.1673-5374.2012.18.009

555. Li J, Zhang L, Xue S, Yu C, Li Y, Li S, Ye Q, Duan X, Peng D. Exploration of the mechanism of Taohong Siwu Decoction for the treatment of ischemic stroke based on CCL2/CCR2 axis. Front Pharmacol. 2024;15:1428572. doi:10.3389/fphar.2024.1428572

556. Xiao Z, Shen D, Lan T, Wei C, Wu W, Sun Q, Luo Z, Chen W, Zhang Y, Hu L, et al. Reduction of lactoferrin aggravates neuronal ferroptosis after intracerebral hemorrhagic stroke in hyperglycemic mice. Redox Biol. 2022;50:102256. doi:10.1016/j.redox.2022.102256

557. Janicki PK, Eyileten C, Ruiz-Velasco V, Pordzik J, Czlonkowska A, Kurkowska-Jastrzebska I, Sugino S, Imamura Kawasawa Y, Mirowska-Guzel D, Postula M. Increased burden of rare deleterious variants of the KCNQ1 gene in patients with large vessel ischemic stroke. Mol Med Rep. 2019;19(4):3263–3272. doi:10.3892/mmr.2019.9987

558. Murali V, Rathika C, Ramgopal S, Padma Malini R, Arun Kumar MJ, Neethi Arasu V, Jeyaram Illiayaraja K, Balakrishnan K. Susceptible and protective associations of HLA DRB1*/DQB1* alleles and haplotypes with ischaemic stroke. Int J Immunogenet. 2016;43(3):159–165. doi:10.1111/iji.12266

559. Fan J, Cao S, Chen M, Yao Q, Zhang X, Du S, Qu H, Cheng Y, Ma S, Zhang M, et al. Investigating the AC079305/DUSP1 Axis as Oxidative Stress-Related Signatures and Immune Infiltration Characteristics in Ischemic Stroke. Oxid Med Cell Longev. 2022;2022:8432352. doi:10.1155/2022/8432352

560. Zhou YF, Chen AQ, Wu JH, Mao L, Xia YP, Jin HJ, He QW, Miao QR, Yue ZY, Liu XL, et al. Sema3E/PlexinD1 signaling inhibits postischemic angiogenesis by regulating endothelial DLL4 and filopodia formation in a rat model of ischemic stroke. FASEB J. 2019;33(4):4947–4961. doi:10.1096/fj.201801706RR

561. Xu L, Wang Y. Combined influence of ABCB1 genetic polymorphism and DNA methylation on aspirin resistance in Chinese ischemic stroke patients. Acta Neurol Belg. 2022;122(4):1057–1064. doi:10.1007/s13760-021-01714-1

562. Yin Y, Zhang Y, Zhang X, Zhang Q, Wang J, Yang T, Liang C, Li W, Liu J, Ma X, et al. Association of MMP3, MMP14, and MMP25 gene polymorphisms with cerebral stroke risk: a case-control study. BMC Med Genomics. 2023;16(1):297. doi:10.1186/s12920-023-01734-1

563. Muthusamy N, Brumm A, Zhang X, Carmichael ST, Ghashghaei HT. Foxj1 expressing ependymal cells do not contribute new cells to sites of injury or stroke in the mouse forebrain. Sci Rep. 2018;8(1):1766. doi:10.1038/s41598-018-19913-x

564. Haarmann A, Zimmermann L, Bieber M, Silwedel C, Stoll G, Schuhmann MK. Regulation and Release of Vasoactive Endoglin by Brain Endothelium in Response to Hypoxia/Reoxygenation in Stroke. Int J Mol Sci. 2022;23(13):7085. doi:10.3390/ijms23137085

565. Zhang Z, Su G, Guo J, et al. Pooled genetic analysis reveals an association of SNPs of only a few genes with risk predisposition to ischemic stroke in a Chinese population. IUBMB Life. 2015;67(3):170–174. doi:10.1002/iub.1359

566. Zhang L, Sui R. Effect of SNP polymorphisms of EDN1, EDNRA, and EDNRB gene on ischemic stroke. Cell Biochem Biophys. 2014;70(1):233–239. doi:10.1007/s12013-014-9887-6

567. Can Demirdöğen B, Miçooğulları Y, Türkanoğlu Özçelik A, Adalı O. Missense Genetic Polymorphisms of Microsomal (EPHX1) and Soluble Epoxide Hydrolase (EPHX2) and Their Relation to the Risk of Large Artery Atherosclerotic Ischemic Stroke in a Turkish Population. Neuropsychiatr Dis Treat. 2021;16:3251–3265. doi:10.2147/NDT.S233992

568. Secondo A, Petrozziello T, Tedeschi V, Boscia F, Vinciguerra A, Ciccone R, Pannaccione A, Molinaro P, Pignataro G, Annunziato L. ORAI1/STIM1 Interaction Intervenes in Stroke and in Neuroprotection Induced by Ischemic Preconditioning Through Store-Operated Calcium Entry. Stroke. 2019;50(5):1240–1249. doi:10.1161/STROKEAHA.118.024115

569. Nishie H, Nakano-Doi A, Sawano T, Nakagomi T. Establishment of a Reproducible Ischemic Stroke Model in Nestin-GFP Mice with High Survival Rates. Int J Mol Sci. 2021;22(23):12997. doi:10.3390/ijms222312997

570. Chi W, Meng F, Li Y, Wang Q, Wang G, Han S, Wang P, Li J. Downregulation of miRNA-134 protects neural cells against ischemic injury in N2A cells and mouse brain with ischemic stroke by targeting HSPA12B. Neuroscience. 2014;277:111–122. doi:10.1016/j.neuroscience.2014.06.062

571. Gillis HL, Kalinina A, Xue Y, Yan K, Turcotte-Cardin V, Todd MAM, Young KG, Lagace D, Picketts DJ. VGF is required for recovery after focal stroke. Exp Neurol. 2023;362:114326. doi:10.1016/j.expneurol.2023.114326

572. Sandelius Å, Cullen NC, Källén Å, Rosengren L, Jensen C, Kostanjevecki V, Vandijck M, Zetterberg H, Blennow K. Transient increase in CSF GAP-43 concentration after ischemic stroke. BMC Neurol. 2018;18(1):202. doi:10.1186/s12883-018-1210-5

573. Wang PR, Masuda Y, Kitamura H, Yamanaka N. Tubulointerstitial injury of Thy-1 nephritis in uninephrectomized stroke-prone spontaneously hypertensive rats. J Nippon Med Sch. 2001;68(4):301–309. doi:10.1272/jnms.68.301

574. Gao XZ, Ma RH, Zhang ZX. miR-339 Promotes Hypoxia-Induced Neuronal Apoptosis and Impairs Cell Viability by Targeting FGF9/CACNG2 and Mediating MAPK Pathway in Ischemic Stroke. Front Neurol. 2020;11:436. doi:10.3389/fneur.2020.00436

575. Jia C, Lovins C, Malone HM, Keasey MP, Hagg T. Female-specific neuroprotection after ischemic stroke by vitronectin-focal adhesion kinase inhibition. J Cereb Blood Flow Metab. 2022;42(10):1961–1974. doi:10.1177/0271678X221107871

576. Ma M, Zhao J, Xie D, Chen J. Association between GABRG2 Gene Single Nucleotide Polymorphisms and Susceptibility to Ischemic Stroke in a Chinese Population. J Integr Neurosci. 2023;22(6):151. doi:10.31083/j.jin2206151

577. Bu F, Min JW, Munshi Y, Lai YJ, Qi L, Urayama A, McCullough LD, Li J. Activation of endothelial ras-related C3 botulinum toxin substrate 1 (Rac1) improves post-stroke recovery and angiogenesis via activating Pak1 in mice. Exp Neurol. 2019;322:113059. doi:10.1016/j.expneurol.2019.113059

578. Schulz A, Wagner F, Ungelenk M, Kurth I, Redecker C. Stroke-like onset of brain stem degeneration presents with unique MRI sign and heterozygous NMNAT2 variant: a case report. Transl Neurodegener. 2016;5:23. doi:10.1186/s40035-016-0069-x

579. Zhao L, Zhang M, Yan F, Cong Y. Knockdown of RMST Impedes Neuronal Apoptosis and Oxidative Stress in OGD/R-Induced Ischemic Stroke Via Depending on the miR-377/SEMA3A Signal Network. Neurochem Res. 2021;46(3):584–594. doi:10.1007/s11064-020-03194-w

580. Ren C, Kobeissy F, Alawieh A, Li N, Li N, Zibara K, Zoltewicz S, Guingab-Cagmat J, Larner SF, Ding Y, et al. Assessment of Serum UCH-L1 and GFAP in Acute Stroke Patients. Sci Rep. 2016;6:24588. doi:10.1038/srep24588

581. Hu Y, Cao X, Zhao Y, Jin Y, Li F, Xu B, Zhao M, Chen Y, Du B, Sun Y, et al. The Function of Spag6 to Repair Brain Edema Damage After Cerebral Ischemic Stroke-reperfusion. Neuroscience. 2023;522:132–149. doi:10.1016/j.neuroscience.2023.04.014

582. Mages B, Fuhs T, Aleithe S, Blietz A, Hobusch C, Härtig W, Schob S, Krueger M, Michalski D. The Cytoskeletal Elements MAP2 and NF-L Show Substantial Alterations in Different Stroke Models While Elevated Serum Levels Highlight Especially MAP2 as a Sensitive Biomarker in Stroke Patients. Mol Neurobiol. 2021;58(8):4051–4069. doi:10.1007/s12035-021-02372-3

583. Lu M, Liu Y, Xian Z, Yu X, Chen J, Tan S, Zhang P, Guo Y. VEGF to CITED2 ratio predicts the collateral circulation of acute ischemic stroke. Front Neurol. 2022;13:1000992. doi:10.3389/fneur.2022.1000992

584. Hwang JA, Choi SK, Kim SH, Kim DW. Pharmacological Inhibition of LRRK2 Exhibits Neuroprotective Activity in Mouse Photothrombotic Stroke Model. Exp Neurobiol. 2024;33(1):36–45. doi:10.5607/en23023

585. Fırtına ÖB, Salt Ö, Sayhan MB, Dibirdik I, Yucal A. Value of plasma alpha- and beta-synuclein levels in the diagnosis, severity, and functional outcome of acute ischemic stroke. Turk J Emerg Med. 2024;24(4):238–244. doi:10.4103/tjem.tjem_17_24

586. McCrory M, Murphy DF, Morris RC, Noad RF. Evaluating the GAD-2 to screen for post-stroke anxiety on an acute stroke unit. Neuropsychol Rehabil. 2023;33(3):480–496. doi:10.1080/09602011.2022.2030366

587. Sharifulina S, Dzreyan V, Guzenko V, Demyanenko S. Histone Methyltransferases SUV39H1 and G9a and DNA Methyltransferase DNMT1 in Penumbra Neurons and Astrocytes after Photothrombotic Stroke. Int J Mol Sci. 2021;22(22):12483. doi:10.3390/ijms222212483

588. Birjandi SZ, Abduljawad N, Nair S, Dehghani M, Suzuki K, Kimura H, Carmichael ST. Phosphodiesterase 10A Inhibition Leads to Brain Region-Specific Recovery Based on Stroke Type. Transl Stroke Res. 2021;12(2):303–315. doi:10.1007/s12975-020-00819-8

589. Deng ZH, Chen YX, Xue-Gao, Yang JY, Wei XY, Zhang GX, Qian JX. Mesenchymal stem cell-derived exosomes ameliorate hypoxic pulmonary hypertension by inhibiting the Hsp90aa1/ERK/pERK pathway. Biochem Pharmacol. 2024;226:116382. doi:10.1016/j.bcp.2024.116382

590. Chen YX, Deng ZH, Xue-Gao, Qiang-Du, Juan-Yin, Chen GH, Li JG, Zhao YM, Zhang HT, Zhang GX, et al. Exosomes derived from mesenchymal stromal cells exert a therapeutic effect on hypoxia-induced pulmonary hypertension by modulating the YAP1/SPP1 signaling pathway. Biomed Pharmacother. 2023;168:115816. doi:10.1016/j.biopha.2023.115816

591. Clapham KR, Rao Y, Sahay S, Sauler M, Lee PJ, Psotka MA, Fares WH, Ahmad T. PECAM-1 is Associated WithOutcomes and Response to Treatment in Pulmonary Arterial Hypertension. Am J Cardiol. 2020;127:198–199. doi:10.1016/j.amjcard.2020.04.031

592. Li Y, Li H, Xing W, Li J, Du R, Cao D, Wang Y, Yang X, Zhong G, Zhao Y, et al. Vascular smooth muscle cell-specific miRNA-214 knockout inhibits angiotensin II-induced hypertension through upregulation of Smad7. FASEB J. 2021;35(11):e21947. doi:10.1096/fj.202100766RR

593. Barberis MC, Veronese S, Bauer D, De Juli E, Harari S. Immunocytochemical detection of progesterone receptors. A study in a patient with primary pulmonary hypertension. Chest. 1995;107(3):869–872. doi:10.1378/chest.107.3.869

594. Murakoshi M, Iwasawa T, Koshida T, Suzuki Y, Gohda T, Kato K. Development of an In-House EphA2 ELISA for Human Serum and Measurement of Circulating Levels of EphA2 in Hypertensive Patients with Renal Dysfunction. Diagnostics (Basel). 2022;12(12):3023. doi:10.3390/diagnostics12123023

595. Han X, Li W, Chen C, Liu J, Sun J, Wang F, Wang C, Mu J, Gu X, Liu F, et al. Genetic variants and mRNA expression of KLF4 and KLF5 with hypertension: A combination of case-control study and cohort study. J Biomed Res. 2024. doi:10.7555/JBR.38.20240208

596. Tian DZ, Wei W, Dong YJ. Influence of COL1A2 gene variants on the incidence of hypertensive intracerebral hemorrhage in a Chinese population. Genet Mol Res. 2016;15(1):10.4238/gmr.15017369. doi:10.4238/gmr.15017369

597. Morris BJ, Chen R, Donlon TA, Kallianpur KJ, Masaki KH, Willcox BJ. Vascular endothelial growth factor receptor 1 gene (FLT1) longevity variant increases lifespan by reducing mortality risk posed by hypertension. Aging (Albany NY). 2023;15(10):3967–3983. doi:10.18632/aging.204722

598. Nagata T, Nakagawa K, Tsurumi F, Watanabe K, Endo T, Hata A. A case of novel NFKB2 variant with hypertensive emergency and nephrotic syndrome leading to CKD 5D. Pediatr Nephrol. 2024;39(9):2637–2640. doi:10.1007/s00467-024-06334-4

599. Jouneau S, Ballerie A, Kerjouan M, Demant X, Blanchard E, Lederlin M. Haemodynamically proven pulmonary hypertension in a patient with GATA2 deficiency-associated pulmonary alveolar proteinosis and fibrosis. Eur Respir J. 2017;49(5):1700407. doi:10.1183/13993003.00407-2017

600. Fähling M, Paliege A, Jönsson S, Becirovic-Agic M, Melville JM, Skogstrand T, Hultström M. NFAT5 regulates renal gene expression in response to angiotensin II through Annexin-A2-mediated posttranscriptional regulation in hypertensive rats. Am J Physiol Renal Physiol. 2019;316(1):F101–F112. doi:10.1152/ajprenal.00361.2018

601. Perepechaeva ML, Stefanova NA, Grishanova AY, Kolosova NG. The Expression of Genes CYP1A1, CYP1B1, and CYP2J3 in Distinct Regions of the Heart and Its Possible Contribution to the Development of Hypertension. Biomedicines. 2024;12(10):2374. doi:10.3390/biomedicines12102374

602. Yan X, Huang J, Zeng Y, Zhong X, Fu Y, Xiao H, Wang X, Lian H, Luo H, Li D, et al. CGRP attenuates pulmonary vascular remodeling by inhibiting the cGAS-STING-NFκB pathway in pulmonary arterial hypertension. Biochem Pharmacol. 2024;222:116093. doi:10.1016/j.bcp.2024.116093

603. Wetzl V, Tiede SL, Faerber L, Weissmann N, Schermuly RT, Ghofrani HA, Gall H. Plasma MMP2/TIMP4 Ratio at Follow-up Assessment Predicts Disease Progression of Idiopathic Pulmonary Arterial Hypertension. Lung. 2017;195(4):489–496. doi:10.1007/s00408-017-0014-5

604. Wang N, Hua J, Fu Y, An J, Chen X, Wang C, Zheng Y, Wang F, Ji Y, Li Q. Updated perspective of EPAS1 and the role in pulmonary hypertension. Front Cell Dev Biol. 2023;11:1125723. doi:10.3389/fcell.2023.1125723

605. Li Y, Cai H, Wei J, Zhu L, Yao Y, Xie M, Song L, Zhang C, Huang X, et al. Dihydroartemisinin Attenuates Hypoxic Pulmonary Hypertension via the Downregulation of miR-335 Targeting Vangl2. DNA Cell Biol. 2022;41(8):750–767. doi:10.1089/dna.2021.1113

606. Alsheikh AJ, Dasinger JH, Abais-Battad JM, Fehrenbach DJ, Yang C, Cowley AW Jr, Mattson DL. CCL2 mediates early renal leukocyte infiltration during salt-sensitive hypertension. Am J Physiol Renal Physiol. 2020;318(4):F982–F993. doi:10.1152/ajprenal.00521.2019

607. Singh A, Zapata RC, Pezeshki A, Knight CG, Tuor UI, Chelikani PK. Whey Protein and Its Components Lactalbumin and Lactoferrin Affect Energy Balance and Protect against Stroke Onset and Renal Damage in Salt-Loaded, High-Fat Fed Male Spontaneously Hypertensive Stroke-Prone Rats. J Nutr. 2020;150(4):763–774. doi:10.1093/jn/nxz312

608. Huang KC, Li TM, Liu X, Chen JH, Chien WK, Shiao YT, Tsang H, Lin TH, Liao CC, Huang SM, et al.KCNQ1 variants associate with hypertension in type 2 diabetes and affect smooth muscle contractility in vitro. J Cell Physiol. 2017;232(12):3309–3316. doi:10.1002/jcp.25775

609. Zhu F, Sun Y, Wang M, Ma S, Chen X, Cao A, Chen F, Qiu Y, Liao Y. Correlation between HLA-DRB1, HLA-DQB1 polymorphism and autoantibodies against angiotensin AT(1) receptors in Chinese patients with essential hypertension. Clin Cardiol. 2011;34(5):302–308. doi:10.1002/clc.20852

610. Chen Y, Ye C, Chen J, Lin D, Wang H, Wang S. Association of the gene polymorphisms of BMPR2, ACVRL1, SMAD9 and their interactions with the risk of essential hypertension in the Chinese Han population. Biosci Rep. 2019;39(1):BSR20181217. doi:10.1042/BSR20181217

611. Wu T, Zhang BQ, Raelson J, Yao YM, Wu HD, Xu ZX, Marois-Blanchet FC, et al. Analysis of the association of EPHB6, EFNB1 and EFNB3 variants with hypertension risks in males with hypogonadism. Sci Rep. 2018;8(1):14497. doi:10.1038/s41598-018-32836-x

612. Pan Z, Yao Y, Liu X, Wang Y, Zhang X, Zha S, Hu K. Nr1d1 inhibition mitigates intermittent hypoxia-induced pulmonary hypertension via Dusp1-mediated Erk1/2 deactivation and mitochondrial fission attenuation. Cell Death Discov. 2024;10(1):459. doi:10.1038/s41420-024-02219-5

613. Zhang X, Zhang C, Li Q, Piao C, Zhang H, Gu H. Clinical characteristics and prognosis analysis of idiopathic and hereditary pulmonary hypertension patients with ACVRL1 gene mutations. Pulm Circ. 2021;11(4):20458940211044577. doi:10.1177/20458940211044577

614. Tan G, Juan C, Mao Y, Xue G, Fang Z. Inhibition of DLL4/Notch Signaling Pathway Promotes M2 Polarization and Cell Proliferation in Pulmonary Arterial Hypertension. ACS Omega. 2024;9(36):37923–37933. doi:10.1021/acsomega.4c04307

615. Gallego-Zazo N, Miranda-Alcaraz L, Cruz-Utrilla A, Del Cerro Marín MJ, Álvarez-Fuente M, Del Mar Rodríguez oSeven Additional Patients with SOX17 Related Pulmonary Arterial Hypertension and Review of the Literature. Genes (Basel). 2023;14(10):1965. doi:10.3390/genes14101965

616. Du M, Kou L, Li S. Water-soluble chitosan regulates vascular remodeling in hypertension via NFATc1. Eur Heart J Suppl. 2016;18(Suppl F):F38. doi:10.1093/eurheartj/suw037

617. Laggner M, Hacker P, Oberndorfer F, Bauer J, Raunegger T, Gerges C, Szerafin T, Thanner J, Lang I, Skoro-Sajer N, et al. The Roles of S100A4 and the EGF/EGFR Signaling Axis in Pulmonary Hypertension with Right Ventricular Hypertrophy. Biology (Basel). 2022;11(1):118. doi:10.3390/biology11010118

618. Sychev D, Shikh N, Morozova T, Grishina E, Ryzhikova K, MalovaE. Effects of ABCB1 rs1045642 polymorphisms on the efficacy and safety of amlodipine therapy in Caucasian patients with stage I-II hypertension. Pharmgenomics Pers Med. 2018;11:157–165. doi:10.2147/PGPM.S158401

619. Hong J, Medzikovic L, Sun W, Wong B, Ruffenach G, Rhodes CJ, Brownstein A, Liang LL, Aryan L, Li M, et al. Integrative Multiomics in the Lung Reveals a Protective Role of Asporin in Pulmonary Arterial Hypertension. Circulation. 2024;150(16):1268–1287. doi:10.1161/CIRCULATIONAHA.124.069864

620. Guo XG, Ding J, Xu H, Xuan TM, Jin WQ, Yin X, Shang YP, Zhang FR, Zhu JH, et al. Comprehensive assessment of the association of WNK4 polymorphisms with hypertension: evidence from a meta-analysis. Sci Rep. 2014;4:6507. doi:10.1038/srep06507

621. Yang L, Liang H, Shen L, Guan Z, Meng X. LncRNA Tug1 involves in the pulmonary vascular remodeling in mice with hypoxic pulmonary hypertension via the microRNA-374c-mediated Foxc1. Life Sci. 2019;237:116769. doi:10.1016/j.lfs.2019.116769

622. Isobe S, Nair RV, Kang HY, Wang L, Moonen JR, Shinohara T, Cao A, Taylor S, Otsuki S, Marciano DP, et al. Reduced FOXF1 links unrepaired DNA damage to pulmonary arterial hypertension. Nat Commun. 2023;14(1):7578. doi:10.1038/s41467-023-43039-y

623. Seidel E, Schewe J, Zhang J, Dinh HA, Forslund SK, Markó L, Hellmig N, Peters J, Muller DN, Lifton RP, et al. Enhanced Ca2+ signaling, mild primary aldosteronism, and hypertension in a familial hyperaldosteronism mouse model (Cacna1hM1560V/+ ). Proc Natl Acad Sci U S A. 2021;118(17):e2014876118. doi:10.1073/pnas.2014876118

624. Gallardo-Vara E, Gamella-Pozuelo L, Perez-Roque L, Bartha JL, Garcia-Palmero I, Casal JI, López-Novoa JM, Pericacho M, Bernabeu C. Potential Role of Circulating Endoglin in Hypertension via the Upregulated Expression of BMP4. Cells. 2020;9(4):988. doi:10.3390/cells9040988

625. Nakano N, Hori H, Abe M, Shibata H, Arimura T, Sasaoka T, Sawabe M, Chida K, Arai T, Nakahara K, et al. Interaction of BMP10 with Tcap may modulate the course of hypertensive cardiac hypertrophy. Am J Physiol Heart Circ Physiol. 2007;293(6):H3396–H3403. doi:10.1152/ajpheart.00311.2007

626. Capri Y, Kwon T, Boyer O, Bourmance L, Testa N, Baudouin V, Bonnefoy R, Couderc A, Meziane C, Tournier-Lasserve E, et al. Biallelic NPR1 loss of function variants are responsible for neonatal systemic hypertension. J Med Genet. 2023;60(10):993–998. doi:10.1136/jmg-2023-109176

627. Huang CC, Chung CM, Yang CY, Leu HB, Huang PH, Lin LY, Wu TC, Lin SJ, Pan WH, Chen JW. SLC12A3 Variation and Renal Function in Chinese Patients With Hypertension. Front Med (Lausanne). 2022;9:863275. doi:10.3389/fmed.2022.863275

628. Masson B, Le Ribeuz H, Sabourin J, Laubry L, Woodhouse E, Foster R, Ruchon Y, Dutheil M, Boët A, Ghigna MR, et al. Orai1 Inhibitors as Potential Treatments for Pulmonary Arterial Hypertension. Circ Res. 2022;131(9):e102–e119. doi:10.1161/CIRCRESAHA.122.321041

629. Li XM, Ling Y, Lu DR, Lu ZQ, Liu Y, Chen HY, Gao X. The obesity-related polymorphism PCSK1 rs6235 is associated with essential hypertension in the Han Chinese population. Hypertens Res. 2012;35(10):994–999. doi:10.1038/hr.2012.79

630. Fargali S, Garcia AL, Sadahiro M, Jiang C, Janssen WG, Lin WJ, Cogliani V, Elste A, Mortillo S, Cero C, et al. The granin VGF promotes genesis of secretory vesicles, and regulates circulating catecholamine levels and blood pressure. FASEB J. 2014;28(5):2120–2133. doi:10.1096/fj.13-239509

631. Chen Y, Li S, Xu Z, Zhang Y, Zhang H, Shi L. Aerobic training-mediated DNA hypermethylation of Agtr1a and Mas1 genes ameliorate mesenteric arterial function in spontaneously hypertensive rats. Mol Biol Rep. 2021;48(12):8033–8044. doi:10.1007/s11033-021-06929-2

632. Li Q, Wong JH, Lu G, Antonio GE, Yeung DK, Ng TB, Forster LE, Yew DT. Gene expression of synaptosomal-associated protein 25 (SNAP-25) in the prefrontal cortex of the spontaneously hypertensive rat (SHR). Biochim Biophys Acta. 2009;1792(8):766–776. doi:10.1016/j.bbadis.2009.05.006

633. Kobayashi Y, Yatsu K, Haruna A, Kawano R, Ozawa M, Haze T, Komiya S, Suzuki S, Ohki Y, Fujiwara A, et al. ATP2B1 gene polymorphisms associated with resistant hypertension in the Japanese population. J Clin Hypertens (Greenwich). 2024;26(4):355–362. doi:10.1111/jch.14785

634. Yeo Y, Yi ES, Kim JM, Jo EK, Seo S, Kim RI, Kim KL, Sung JH, Park SG, Suh W. et al. FGF12 (Fibroblast Growth Factor 12) Inhibits Vascular Smooth Muscle Cell Remodeling in Pulmonary Arterial Hypertension. Hypertension. 2020;76(6):1778–1786. doi:10.1161/HYPERTENSIONAHA.120.15068

635. Sahoo S, Meijles DN, Al Ghouleh I, Tandon M, Cifuentes-Pagano E, Sembrat J, Rojas M, Goncharova E, Pagano PJ. MEF2C-MYOCD and Leiomodin1 Suppression by miRNA-214 Promotes Smooth Muscle Cell Phenotype Switching in Pulmonary Arterial Hypertension. PLoS One. 2016;11(5):e0153780. doi:10.1371/journal.pone.0153780

636. Hong KW, Go MJ, Jin HS, Lim JE, Lee JY, Han BG, Hwang SY, Lee SH, Park HK, Cho YS, et al. Genetic variations in ATP2B1, CSK, ARSG and CSMD1 loci are related to blood pressure and/or hypertension in two Korean cohorts. J Hum Hypertens. 2010;24(6):367–372. doi:10.1038/jhh.2009.86

637. Zhang QY, Guo Y, Jiang XL, Liu X, Zhao SG, Zhou XL, Yang ZW. Intestinal Cckbr-specific knockout mouse as a novel model of salt-sensitive hypertension via sodium over-absorption. J Geriatr Cardiol. 2023;20(7):538–547. doi:10.26599/1671-5411.2023.07.001

638. Agrud A, Subburaju S, Goel P, Ren J, Kumar AS, Caldarone BJ, Dai W, Chavez J, Fukumura D, Jain RK, et al. Gabrb3 endothelial cell-specific knockout mice display abnormal blood flow, hypertension, and behavioral dysfunction. Sci Rep. 2022;12(1):4922. doi:10.1038/s41598-022-08806-9

639. Olinger E, Phakdeekitcharoen P, Caliskan Y, Orr S, Mabillard H, Pickles C, Tse Y, Wood K; Genomics England Research Consortium; Sayer JA. Biallelic variants in TTC21B as a rare cause of early-onset arterial hypertension and tubuloglomerular kidney disease. Am J Med Genet C Semin Med Genet. 2022;190(1):109–120. doi:10.1002/ajmg.c.31964

640. Viazzi F, Ramesh G, Jayakumar C, Leoncini G, Garneri D, Pontremoli R. Increased urine semaphorin-3A is associated with renal damage in hypertensive patients with chronic kidney disease: a nested case-control study. J Nephrol. 2015;28(3):315–320. doi:10.1007/s40620-014-0097-5

641. Tang H, Gupta A, Morrisroe SA, Bao C, Schwantes-An TH, Gupta G, Liang S, Sun Y, Chu A, et al. Deficiency of the Deubiquitinase UCHL1 Attenuates Pulmonary Arterial Hypertension. Circulation. 2024;150(4):302–316. doi:10.1161/CIRCULATIONAHA.123.065304

642. Laggner M, Oberndorfer F, Golabi B, Bauer J, Zuckermann A, Hacker P, Lang I, Skoro-Sajer N, Gerges C, Taghavi S, et al. EGR1 Is Implicated in Right Ventricular Cardiac Remodeling Associated with Pulmonary Hypertension. Biology (Basel). 2022;11(5):677. doi:10.3390/biology11050677

643. Li D, Shao NY, Moonen JR, Zhao Z, Shi M, Otsuki S, Wang L, Nguyen T, Yan E, Marciano DP, et al. ALDH1A3 Coordinates Metabolism With Gene Regulation in Pulmonary Arterial Hypertension. Circulation. 2021;143(21):2074–2090. doi:10.1161/CIRCULATIONAHA.120.048845

644. Liang J, Zhang X, Xia W, Tong X, Qiu Y, Qiu Y, He J, Yu B, Huang H, Tao J. Promotion of Aerobic Exercise Induced Angiogenesis Is Associated With Decline in Blood Pressure in Hypertension: Result of EXCAVATION-CHN1. Hypertension. 2021;77(4):1141–1153. doi:10.1161/HYPERTENSIONAHA.120.16107

645. Tanaka S, Yamamoto T, Mikawa M, Nawata J, Fujii S, Nakamura Y, Kato T, Fukuda M, Suetomi T, Uchinoumi H, et al. Stabilization of RyR2 maintains right ventricular function, reduces the development of ventricular arrhythmias, and improves prognosis in pulmonary hypertension. Heart Rhythm. 2022;19(6):986–997. doi:10.1016/j.hrthm.2022.02.003

646. Schinagl C, Melum GR, Rødningen OK, Bjørgo K, Andresen JH. Severe persistent pulmonary hypertension of the newborn and dysmorphic features in neonate with a deletion involving TWIST1 and PHF14: a case report. J Med Case Rep. 2017;11(1):226. doi:10.1186/s13256-017-1402-4

647. Zhu H, Zhu L, Fang Z, Yang S, Chen Y, Jin Y, Zhao X, Shen C, Yao Y. Common variants at somatostatin are significantly associated with hypertension incidence in smoking and drinking populations. J Am Soc Hypertens. 2018;12(3):230–237.e12. doi:10.1016/j.jash.2017.12.009

648. Keranov S, Dörr O, Jafari L, Liebetrau C, Keller T, Troidl C, Kriechbaum S, Voss S, Richter M, Tello K, et al. SPARCL1 as a biomarker of maladaptive right ventricular remodelling in pulmonary hypertension. Biomarkers. 2020;25(3):290–295. doi:10.1080/1354750X.2020.1745889

649. Zhang HL, Liu ZH, Luo Q, Wang Y, Zhao ZH. Abnormal expression of NSF, α-SNAP and SNAP23 in pulmonary arterial hypertension in rats treated with monocrotaline. Int J Clin Exp Med. 2015;8(2):1834–1843.

650. Wang LA, Nguyen DH, Mifflin SW. CRHR2 (Corticotropin-Releasing Hormone Receptor 2) in the Nucleus of the Solitary Tract Contributes to Intermittent Hypoxia-Induced Hypertension. Hypertension. 2018;72(4):994–1001. doi:10.1161/HYPERTENSIONAHA.118.11497

651. Faruque MU, Chen G, Doumatey A, Huang H, Zhou J, Dunston GM, Rotimi CN, Adeyemo AA. Association of ATP1B1, RGS5 and SELE polymorphisms with hypertension and blood pressure in African-Americans. J Hypertens. 2011;29(10):1906–1912. doi:10.1097/HJH.0b013e32834b000d

652. Herrera VL, Pasion KA, Moran AM, Zaninello R, Ortu MF, Fresu G, Piras DA, Argiolas G, Troffa C, Glorioso V, et al. A functional 12T-insertion polymorphism in the ATP1A1 promoter confers decreased susceptibility to hypertension in a male Sardinian population. PLoS One. 2015;10(1):e0116724. doi:10.1371/journal.pone.0116724

653. Takahashi Y, Yamazaki K, Kamatani Y, Kubo M, Matsuda K, Asai S. A genome-wide association study identifies a novel candidate locus at the DLGAP1 gene with susceptibility to resistant hypertension in the Japanese population. Sci Rep. 2021;11(1):19497. doi:10.1038/s41598-021-98144-z

654. Huang YY, Yu YF, Zhang C, Chen Y, Zhou Q, Li Z, Zhou S, Li Z, Guo L, Wu D, et al. Validation of Phosphodiesterase-10 as a Novel Target for Pulmonary Arterial Hypertension via Highly Selective and Subnanomolar Inhibitors. J Med Chem. 2019;62(7):3707–3721. doi:10.1021/acs.jmedchem.9b00224

655. Hu Y, Xia W, Li Y, Wang Q, Lin S, Wang B, Zhou C, Cui Y, Jiang Y, Pu X, et al. High-salt intake increases TRPC3 expression and enhances TRPC3-mediated calcium influx and systolic blood pressure in hypertensive patients. Hypertens Res. 2020;43(7):679–687. doi:10.1038/s41440-020-0409-1

656. Xiang X, You XM, Li LQ. Expression of HSP90AA1/HSPA8 in hepatocellular carcinoma patients with depression. Onco Targets Ther. 2018;11:3013–3023. doi:10.2147/OTT.S159432

657. Ma F, Bian H, Jiao W, Zhang N. Single-cell RNA-seq reveals the role of YAP1 in prefrontal cortex microglia in depression. BMC Neurol. 2024;24(1):191. doi:10.1186/s12883-024-03685-1

658. Huang K, Zhang X, Duan J, Wang R, Wu Z, Yang C, Yang L. STAT4 and COL1A2 are potential diagnostic biomarkers and therapeutic targets for heart failure comorbided with depression. Brain Res Bull. 2022;184:68–75. doi:10.1016/j.brainresbull.2022.03.014

659. Choi M, Wang SE, Ko SY, Kang HJ, Chae SY, Lee SH, Kim YS, Duman RS, Son H. Overexpression of human GATA-1 and GATA-2 interferes with spine formation and produces depressive behavior in rats. PLoS One. 2014;9(10):e109253. doi:10.1371/journal.pone.0109253

660. Dai Y, Wei T, Huang Y, Bei Y, Lin H, Shen Z, Yu L, Yang M, Xu H, He W, et al. Upregulation of HDAC9 in hippocampal neurons mediates depression-like behaviours by inhibiting ANXA2 degradation. Cell Mol Life Sci. 2023;80(10):289. doi:10.1007/s00018-023-04945-y

661. Chen S, Li J, Yan L, Zhang X, Huang J, Zhou P. Electroacupuncture alleviates the symptom of depression in mice by regulating the cGAS-STING-NLRP3 signaling. Aging (Albany NY). 2024;16(8):6731–6744. doi:10.18632/aging.205596

662. Bobińska K, Szemraj J, Czarny P, Gałecki P. Role of MMP-2, MMP-7, MMP-9 and TIMP-2 in the development of recurrent depressive disorder. J Affect Disord. 2016;205:119–129. doi:10.1016/j.jad.2016.03.068

663. Curzytek K, Leśkiewicz M. Targeting the CCL2-CCR2 axis in depressive disorders. Pharmacol Rep. 2021;73(4):1052–1062. doi:10.1007/s43440-021-00280-w

664. Zhang J, Xin H, Wang W, Li Y, Wu R, Wei L, Su S, Wang X, Wang X, Wang X, et al. Investigating the modulatory effects of lactoferrin on depressed rats through 16S rDNA gene sequencing and LC-MS metabolomics analysis. Sci Rep. 2024;14(1):22111. doi:10.1038/s41598-024-72793-2

665. Chen MX, Oh YS, Kim Y. S100A10 and its binding partners in depression and antidepressant actions. Front Mol Neurosci. 2022;15:953066. doi:10.3389/fnmol.2022.953066

666. Zhao Y, Wang S, Chu Z, Dang Y, Zhu J, Su X. MicroRNA-101 in the ventrolateral orbital cortex (VLO) modulates depressive-like behaviors in rats and targets dual-specificity phosphatase 1 (DUSP1). Brain Res. 2017;1669:55–62. doi:10.1016/j.brainres.2017.05.020

667. Oliveira S, Ardais AP, Bastos CR, Gazal M, Jansen K, de Mattos Souza L, da Silva RA, Kaster MP, Lara DR, Ghisleni G. Impact of genetic variations in ADORA2A gene on depression and symptoms: a cross-sectional population-based study. Purinergic Signal. 2019;l(1):37-44. doi:10.1007/s11302-018-9635-2

668. Brückl TM, Uhr M. ABCB1 genotyping in the treatment of depression. Pharmacogenomics. 2016;17(18):2039–2069. doi:10.2217/pgs.16.18

669. Zhong J, Li S, Zeng W, Li X, Gu C, Liu J, Luo XJ. Integration of GWAS and brain eQTL identifies FLOT1 as a risk gene for major depressive disorder. Neuropsychopharmacology. 2019;44(9):1542–1551. doi:10.1038/s41386-019-0345-4

670. Wohleb ES, Terwilliger R, Duman CH, Duman RS. Stress-Induced Neuronal Colony Stimulating Factor 1 Provokes Microglia-Mediated Neuronal Remodeling and Depressive-like Behavior. Biol Psychiatry. 2018;83(1):38–49. doi:10.1016/j.biopsych.2017.05.026

671. Yun S, Donovan MH, Ross MN, Richardson DR, Reister R, Farnbauch LA, Fischer SJ, Riethmacher D, Gershenfeld HK, Lagace DC, et al. Stress-Induced Anxiety-and Depressive-Like Phenotype Associated with Transient Reduction in Neurogenesis in Adult Nestin-CreERT2/Diphtheria Toxin Fragment A Transgenic Mice. PLoS One. 2016;11(1):e0147256. doi:10.1371/journal.pone.0147256

672. Li YJ, Kresock E, Kuplicki R, Savitz J, McKinney BA. Differential expression of MDGA1 in major depressive disorder. Brain Behav Immun Health. 2022;26:100534. doi:10.1016/j.bbih.2022.100534

673. Ye Q, Zhang Y, Zhang Y, Chen Z, Yu C, Zheng C, Yu H, Zhou D, Li X. Low VGF is associated with executive dysfunction in patients with major depressive disorder. J Psychiatr Res. 2022;152:182–186. doi:10.1016/j.jpsychires.2022.06.030

674. Han MH, Jiao S, Jia JM, Chen Y, Chen CY, Gucek M, Markey SP, Li Z. The novel caspase-3 substrate Gap43 is involved in AMPA receptor endocytosis and long-term depression. Mol Cell Proteomics. 2013;12(12):3719–3731. doi:10.1074/mcp.M113.030676

675. Webster MJ, Vawter MP, Freed WJ. Immunohistochemical localization of the cell adhesion molecules Thy-1 and L1 in the human prefrontal cortex patients with schizophrenia, bipolar disorder, and depression. Mol Psychiatry. 1999;4(1):46–52. doi:10.1038/sj.mp.4000450

676. Zeng D, He S, Yu S, Li G, Ma C, Wen Y, Shen Y, Yu Y, Li H. Analysis of the association of MIR124-1 and its target gene RGS4 polymorphisms with major depressive disorder and antidepressant response. Neuropsychiatr Dis Treat. 2018;14:715–723. doi:10.2147/NDT.S155076

677. Barbiero I, Bianchi M, Kilstrup-Nielsen C. Therapeutic potential of pregnenolone and pregnenolone methyl ether on depressive and CDKL5 deficiency disorders: Focus on microtubule targeting. J Neuroendocrinol. 2022;34(2):e13033. doi:10.1111/jne.13033

678. Wang X, Cheng W, Zhu J, Yin H, Chang S, Yue W, Yu H. Integrating genome-wide association study and expression quantitative trait loci data identifies NEGR1 as a causal risk gene of major depression disorder. J Affect Disord. 2020;265:679–686. doi:10.1016/j.jad.2019.11.116

679. Yu S, Zhao Y, Luo Q, Gu B, Wang X, Cheng J, Wang Z, Liu D, Ho RCM, Ho CSH. Early life stress enhances the susceptibility to depression and interferes with neuroplasticity in the hippocampus of adolescent mice via regulating miR-34c-5p/SYT1 axis. J Psychiatr Res. 2024;170:262–276. doi:10.1016/j.jpsychires.2023.12.030

680. Wang Q, Wang Y, Ji W, Zhou G, He K, Li Z, Chen J, Li W, Wen Z, Shen J, et al. SNAP25 is associated with schizophrenia and major depressive disorder in the Han Chinese population. J Clin Psychiatry. 2015;76(1):e76–e82. doi:10.4088/JCP.13m08962

681. Prats C, Arias B, Ortet G, Ibáñez MI, Moya J, Pomarol-Clotet E, Fañanás L, Fatjó-Vilas M. Neurotrophins role in depressive symptoms and executive function performance: Association analysis of NRN1 gene and its interaction with BDNF gene in a non-clinical sample. J Affect Disord. 2017;211:92–98. doi:10.1016/j.jad.2016.11.017

682. Shah MM. HCN1 channels: a new therapeutic target for depressive disorders?. Sci Signal. 2012;5(244):pe44. doi:10.1126/scisignal.2003593

683. Fuchsova B, Alvarez Juliá A, Rizavi HS, Frasch AC, Pandey GN. Expression of p21-activated kinases 1 and 3 is altered in the brain of subjects with depression. Neuroscience. 2016;333:331–344. doi:10.1016/j.neuroscience.2016.07.037

684. Yang W, Li S, Li XJ. AHI1: linking depression and impaired antiviral immune response. Cell Res. 2022;32(10):869–870. doi:10.1038/s41422-022-00702-1

685. Ji W, Li T, Pan Y, Tao H, Ju K, Wen Z, Fu Y, An Z, Zhao Q, Wang T, et al. CNTNAP2 is significantly associated with schizophrenia and major depression in the Han Chinese population. Psychiatry Res. 2013;207(3):225–228. doi:10.1016/j.psychres.2012.09.024

686. Auffenberg E, Hedrich UB, Barbieri R, Miely D, Groschup B, Wuttke TV, Vogel N, Lührs P, Zanardi I, Bertelli S, et al. Hyperexcitable interneurons trigger cortical spreading depression in an Scn1a migraine model. J Clin Invest. 2021;131(21):e142202. doi:10.1172/JCI142202

687. Huang G, Wang S, Yan J, Li C, Feng J, Chen Q, Zheng X, Li H, He Y, Young AJ, et al. Depression-/Anxiety-Like Behavior Alterations in Adult Slit2 Transgenic Mice. Front Behav Neurosci. 2021;14:622257. doi:10.3389/fnbeh.2020.622257

688. Fuchsova B, Alvarez Juliá A, Rizavi HS, Frasch AC, Pandey GN. Expression of p21-activated kinases 1 and 3 is altered in the brain of subjects with depression. Neuroscience. 2016;333:331–344. doi:10.1016/j.neuroscience.2016.07.037

689. Yang Y, Guan W, Sheng XM, Gu HJ. Role of Semaphorin 3A in common psychiatric illnesses such as schizophrenia, depression, and anxiety. Biochem Pharmacol. 2024;226:116358. doi:10.1016/j.bcp.2024.116358

690. Choi JE, Lee JJ, Kang W, Kim HJ, Cho JH, Han PL, Lee KJ. Proteomic Analysis of Hippocampus in a Mouse Model of Depression Reveals Neuroprotective Function of Ubiquitin C-terminal Hydrolase L1 (UCH-L1) via Stress-induced Cysteine Oxidative Modifications. Mol Cell Proteomics. 2018;17(9):1803–1823. doi:10.1074/mcp.RA118.000835

691. Lee WH, Bondy C. Induction of EGR1/NGFI-A Gene Expression by Spreading Depression and Focal Cerebral Ischemia. Mol Cell Neurosci. 1993;4(3):225–230. doi:10.1006/mcne.1993.1028

692. Daftary S, Yon JM, Choi EK, Kim YB, Bice C, Kulikova A, Park J, Sherwood Brown E. Microtubule associated protein 2 in bipolar depression: Impact of pregnenolone. J Affect Disord. 2017;218:49–52. doi:10.1016/j.jad.2017.04.024

693. Kim YG, Chang HS, Won ES, Ham BJ, Lee MS. Serotonin-related polymorphisms in TPH1 and HTR5A genes are not associated with escitalopram treatment response in Korean patients with major depression. Neuropsychobiology. 2014;69(4):210–219. doi:10.1159/000362241

694. Filippone A, Cucinotta L, Bova V, Lanza M, Casili G, Paterniti I, Campolo M, Cuzzocrea S, Esposito E. Inhibition of LRRK2 Attenuates Depression-Related Symptoms in Mice with Moderate Traumatic Brain Injury. Cells. 2023;12(7):1040. doi:10.3390/cells12071040

695. Skiba A, Talarowska M, Szemraj J, Gałecki P. Is NRXN1 Gene Expression an Important Marker of Treatment of Depressive Disorders? A Pilot Study. J Pers Med. 2021;11(7):637. doi:10.3390/jpm11070637

696. Wang Z, Cheng X, Shuang R, Gao T, Zhao T, Hou D, Zhang Y, Yang J, Tao W. Dandouchi Polypeptide Alleviates Depressive-like Behavior and Promotes Hippocampal Neurogenesis by Activating the TRIM67/NF-κB Pathway in CUMS-Induced Mice. J Agric Food Chem. 2024;72(30):16726–16738. doi:10.1021/acs.jafc.4c02183

697. Zhang H, Liu S, Qin Q, Xu Z, Qu Y, Wang Y, Wang J, Du Z, Yuan S, Hong S, et al. Genetic and Pharmacological Inhibition of Astrocytic Mysm1 Alleviates Depressive-Like Disorders by Promoting ATP Production. Adv Sci (Weinh). 2022. doi:10.1002/advs.202204463

698. Figueroa KP, Gan SR, Perlman S, Wilmot G, Gomez CM, Schmahmann J, Paulson H, Shakkottai VG, Ying SH, Zesiewicz T, et al. C9orf72 repeat expansions as genetic modifiers for depression in spinocerebellar ataxias. Mov Disord. 2018;33(3):497–498. doi:10.1002/mds.27258

699. Fee C, Banasr M, Sibille E. Somatostatin-Positive Gamma-Aminobutyric Acid Interneuron Deficits in Depression: Cortical Microcircuit and Therapeutic Perspectives. Biol Psychiatry. 2017;82(8):549–559. doi:10.1016/j.biopsych.2017.05.024

700. Thaweethee-Sukjai B, Suttajit S, Thanoi S, Dalton CF, Reynolds GP, Nudmamud-Thanoi S. Parvalbumin Promoter Methylation Altered in Major Depressive Disorder. Int J Med Sci. 2019;16(9):1207–1214. doi:10.7150/ijms.36131

701. Bilkei-Gorzo A, Racz I, Michel K, Zimmer A. Diminished anxiety- and depression-related behaviors in mice with selective deletion of the Tac1 gene. J Neurosci. 2002;22(22):10046–10052. doi:10.1523/JNEUROSCI.22-22-10046.2002

702. Sun L, Bai D, Lin M, Eerdenidalai, Zhang L, Wang F, Jin S. miR-96 Inhibits SV2C to Promote Depression-Like Behavior and Memory Disorders in Mice. Front Behav Neurosci. 2021;14:575345. doi:10.3389/fnbeh.2020.575345

703. Kirshenbaum GS, Saltzman K, Rose B, Petersen J, Vilsen B, Roder JC. Decreased neuronal Na+, K+ -ATPase activity in Atp1a3 heterozygous mice increases susceptibility to depression-like endophenotypes by chronic variable stress. Genes Brain Behav. 2011;10(5):542–550. doi:10.1111/j.1601-183X.2011.00691.x

704. Li T, Tian Y, Li Q, Chen H, Lv H, Xie W, Han J. The Neurexin/N-Ethylmaleimide-sensitive Factor (NSF) Interaction Regulates Short Term Synaptic Depression. J Biol Chem. 2015;290(29):17656–17667. doi:10.1074/jbc.M115.644583

705. Amin M, Ott J, Gordon D, Wu R, Postolache TT, Vergare M, Gragnoli C. Comorbidity of Novel CRHR2 Gene Variants in Type 2 Diabetes and Depression. Int J Mol Sci. 2022;23(17):9819. doi:10.3390/ijms23179819

706. Tsetsenis T, Younts TJ, Chiu CQ, Kaeser PS, Castillo PE, Südhof TC. Rab3B protein is required for long-term depression of hippocampal inhibitory synapses and for normal reversal learning. Proc Natl Acad Sci U S A 2011;108(34):14300–14305. doi:10.1073/pnas.1112237108

707. Gonda X, Eszlari N, Anderson IM, Deakin JF, Bagdy G, Juhasz G. Association of ATP6V1B2 rs1106634 with lifetime risk of depression and hippocampal neurocognitive deficits: possible novel mechanisms in the etiopathology of depression. Transl Psychiatry. 2016;6(11):e945. doi:10.1038/tp.2016.221

708. Wang L, Wang M, Zhao C, Jian J, Qiao D. Association of HTR3B gene polymorphisms with depression and its executive dysfunction: a case-control study. BMC Psychiatry. 2023;23(1):128. doi:10.1186/s12888-023-04625-y

709. Costain G, Walker S, Argiropoulos B, Baribeau DA, Bassett AS, Boot E, Devriendt K, Kellam B, Marshall CR, Prasad A, et al. Rare copy number variations affecting the synaptic gene DMXL2 in neurodevelopmental disorders. J Neurodev Disord. 2019;11(1):3. doi:10.1186/s11689-019-9263-3

710. Huang C, Yang X, Zeng B, Zeng L, Gong X, Zhou C, Xia J, Lian B, Qin Y, Yang L, et al. Proteomic analysis of olfactory bulb suggests CACNA1E as a promoter of CREB signaling in microbiota-induced depression. J Proteomics. 2019;194:132–147. doi:10.1016/j.jprot.2018.11.023

711. Lai J, Zhang P, Jiang J, Mou T, Li Y, Xi C, Wu L, Gao X, Zhang D, Chen Y, et al. New Evidence of Gut Microbiota Involvement in the Neuropathogenesis of Bipolar Depression by TRANK1 Modulation: Joint Clinical and Animal Data. Front Immunol. 2021;12:789647. doi:10.3389/fimmu.2021.789647

712. Lee JE, Park SY, Han PL. Aging-Dependent Downregulation of SUV39H1 Histone Methyltransferase Increases Susceptibility to Stress-Induced Depressive Behavior. Mol Neurobiol. 2021;58(12):6427–6442. doi:10.1007/s12035-021-02529-0

713. Myrga JM, Juengst SB, Failla MD, Conley YP, Arenth PM, Grace AA, Wagner AK. OMT and ANKK1 Genetics Interact With Depression to Influence Behavior Following Severe TBI: An Initial Assessment. Neurorehabil Neural Repair. 2016;30(10):920–930. doi:10.1177/1545968316648409

714. Kim SJ. TRPC3 channel underlies cerebellar long-term depression. Cerebellum. 2013;12(3):334–337. doi:10.1007/s12311-013-0455-1

715. Lu X, Wang Y, Ma Y, Huang D, Lu Y, Liu X, Zhou R, Yu P, Zhang L, Chen J, et al. YAP1 induces marrow derived suppressor cell recruitment in Chlamydia trachomatis infection. Immunol Lett. 2022;242:8–16. doi:10.1016/j.imlet.2021.12.003

716. Fuchs M, Trampuz A, Kirschbaum S, Winkler T, Sass FA. Soluble Pecam-1 as a Biomarker in Periprosthetic Joint Infection. J Clin Med. 2021;10(4):612. doi:10.3390/jcm10040612

717. Wu Y, Wu M, Ming S, Zhan X, Hu S, Li X, Yin H, Cao C, Liu J, Li J, et al. TREM-2 promotes Th1 responses by interacting with the CD3ζ-ZAP70 complex following Mycobacterium tuberculosis infection. J Clin Invest. 2021;131(17):e137407. doi:10.1172/JCI137407

718. Argentou N, Germanidis G, Hytiroglou P, Apostolou E, Vassiliadis T, Patsiaoura K, Sideras P, Germenis AE, Speletas M. TGF-β signaling is activated in patients with chronic HBV infection and repressed by SMAD7 overexpression after successful antiviral treatment. Inflamm Res. 2016;65(5):355–365. doi:10.1007/s00011-016-0921-6

719. Cao Y, Chen F, Zhu S, Zhu D, Qi H. Staphylococcus aureus infection initiates hypoxia-mediated STIP1 homology and U-box containing protein 1 upregulation to trigger osteomyelitis. Toxicon. 2024;248:108049. doi:10.1016/j.toxicon.2024.108049

720. Vincenzi M, Mercurio FA, Leone M. EPHA2 Receptor as a Possible Therapeutic Target in Viral Infections. Curr Med Chem. 2024;31(35):5670–5701. doi:10.2174/0109298673256638231003111234

721. Nawandar DM, Wang A, Makielski K, Lee D, Ma S, Barlow E, Reusch J, Jiang R, Wille CK, Greenspan D, et al. Differentiation-Dependent KLF4 Expression Promotes Lytic Epstein-Barr Virus Infection in Epithelial Cells. PLoS Pathog. 2015;11(10):e1005195. doi:10.1371/journal.ppat.1005195

722. Aird A, Lagos M, Vargas-Hernández A, Posey JE, Coban-Akdemir Z, Jhangiani S, Mace EM, Reyes A, King A, Cavagnaro F, et al. Novel Heterozygous Mutation in NFKB2 Is Associated With Early Onset CVID and a Functional Defect in NK Cells Complicated by Disseminated CMV Infection and Severe Nephrotic Syndrome. Front Pediatr. 2019;7:303. doi:10.3389/fped.2019.00303

723. Chen YC, Hsiao CC, Chen TW, Wu CC, Chao TY, Leung SY, Eng HL, Lee CP, Wang TY, Lin MC. Whole Genome DNA Methylation Analysis of Active Pulmonary Tuberculosis Disease Identifies Novel Epigenotypes: PARP9/miR-505/RASGRP4/GNG12 Gene Methylation and Clinical Phenotypes. Int J Mol Sci. 2020;21(9):3180. doi:10.3390/ijms21093180

724. Suzuki T, Takaya S, Kunimatsu J, Kutsuna S, Hayakawa K, Shibata H, Yasumi T, Ohmagari N. GATA2 mutation underlies hemophagocytic lymphohistiocytosis in an adult with primary cytomegalovirus infection. J Infect Chemother. 2020;26(2):252–256. doi:10.1016/j.jiac.2019.07.002

725. Wang T, Zhao D, Zhang Y, Yu D, Liu G, Zhang K. Annexin A2: A Double-Edged Sword in Pathogen Infection. Pathogens. 2024;13(7):564. doi:10.3390/pathogens13070564

726. Qiao RB, Dai WH, Li W, Yang X, He DM, Gao R, Cui YQ, Wang RX, Ma XY, Wang FJ, et al. The cytochrome P4501A1 (CYP1A1) inhibitor bergamottin enhances host tolerance to multidrug-resistant Vibrio vulnificus infection. Chin J Traumatol. 2024;27(5):295–304. doi:10.1016/j.cjtee.2024.07.003

727. Gonçalves AV, Margolis SR, Quirino GFS, Mascarenhas DPA, Rauch I, Nichols RD, Ansaldo E, Fontana MF, Vance RE, Zamboni DS. Gasdermin-D and Caspase-7 are the key Caspase-1/8 substrates downstream of the NAIP5/NLRC4 inflammasome required for restriction of Legionella pneumophila. PLoS Pathog. 2019;15(6):e1007886. doi:10.1371/journal.ppat.1007886

728. Siddiqui MA, Yamashita M. Toll-Like Receptor (TLR) Signaling Enables Cyclic GMP-AMP Synthase (cGAS) Sensing of HIV-1 Infection in Macrophages. mBio. 2021;12(6):e0281721. doi:10.1128/mBio.02817-21

729. Gutierrez FR, Lalu MM, Mariano FS, Milanezi CM, Cena J, Gerlach RF, Santos JE, Torres-Dueñas D, Cunha FQ, Schulz R, et al. Increased activities of cardiac matrix metalloproteinases matrix metalloproteinase (MMP)-2 and MMP-9 are associated with mortality during the acute phase of experimental Trypanosoma cruzi infection. J Infect Dis. 2008;197(10):1468–1476. doi:10.1086/587487

730. Sultana H, Neelakanta G, Foellmer HG, Montgomery RR, Anderson JF, Koski RA, Medzhitov RM, Fikrig E. Semaphorin 7A contributes to West Nile virus pathogenesis through TGF-β1/Smad6 signaling. J Immunol. 2012;189(6):3150–3158. doi:10.4049/jimmunol.1201140

731. He TS, Dang L, Zhang J, Zhang J, Wang G, Wang E, Xia H, Zhou W, Wu S, Liu X. The Hippo signaling component LATS2 enhances innate immunity to inhibit HIV-1 infection through PQBP1-cGAS pathway. Cell Death Differ. 2022;29(1):192–205. doi:10.1038/s41418-021-00849-1

732. Abbas M, Verma S, Verma S, Siddiqui S, Khan FH, Raza ST, Siddiqi Z, Eba A, Mahdi F. Association of GSTM1 and GSTT1 gene polymorphisms with COVID-19 susceptibility and its outcome. J Med Virol. 2021;93(9):5446–5451. doi:10.1002/jmv.27076

733. Yin S, Yu J, Hu B, Lu C, Liu X, Gao X, Li W, Zhou L, Wang J, Wang D, et al. Runx3 Mediates Resistance to Intracellular Bacterial Infection by Promoting IL12 Signaling in Group 1 ILC and NCR+ILC3. Front Immunol. 2018;9:2101. doi:10.3389/fimmu.2018.02101

734. Ranjbar M, Rahimi A, Baghernejadan Z, Ghorbani A, Khorramdelazad H. Role of CCL2/CCR2 axis in the pathogenesis of COVID-19 and possible Treatments: All options on the Table. Int Immunopharmacol. 2022;113(Pt A):109325. doi:10.1016/j.intimp.2022.109325

735. Wakabayashi H, Oda H, Yamauchi K, Abe F. Lactoferrin for prevention of common viral infections. J Infect Chemother. 2014;20(11):666–671. doi:10.1016/j.jiac.2014.08.003

736. Ou G, Xu H, Yu H, Liu X, Yang L, Ji X, Wang J, Liu Z. The roles of HLA-DQB1 gene polymorphisms in hepatitis B virus infection. J Transl Med. 2018;16(1):362. doi:10.1186/s12967-018-1716-z

737. Feng J, Meng W, Chen L, Zhang X, Markazi A, Yuan W, Huang Y, Gao SJ. N6-Methyladenosine and Reader Protein YTHDF2 Enhance the Innate Immune Response by Mediating DUSP1 mRNA Degradation and Activating Mitogen-Activated Protein Kinases during Bacterial and Viral Infections. mBio. 2023;14(1):e0334922. doi:10.1128/mbio.03349-22

738. Zhan C, Sun Y, Pan J, Chen L, Yuan T. Effect of the Notch4/Dll4 signaling pathway in early gestational intrauterine infection on lung development. Exp Ther Med. 2021;22(3):972. doi:10.3892/etm.2021.10404

739. Jiyarom B, Giannakopoulos S, Strange DP, Panova N, Gale M Jr, Verma S. RIG-I and MDA5 are modulated by bone morphogenetic protein (BMP6) and are essential for restricting Zika virus infection in human Sertoli cells. Front Microbiol. 2023;13:1062499. doi:10.3389/fmicb.2022.1062499

740. Kayama H, Koga R, Atarashi K, Okuyama M, Kimura T, Mak TW, Uematsu S, Akira S, Takayanagi H, Honda K, et al. NFATc1 mediates Toll-like receptor-independent innate immune responses during Trypanosoma cruzi infection. PLoS Pathog. 2009;5(7):e1000514. doi:10.1371/journal.ppat.1000514

741. Zhang J, Jiao Y, Hou S, Tian T, Yuan Q, Hao H, Wu Z, Bao X. S100A4 contributes to colitis development by increasing the adherence of Citrobacter rodentium in intestinal epithelial cells. Sci Rep. 2017;7(1):12099. doi:10.1038/s41598-017-12256-z

742. Nishijima T, Komatsu H, Higasa K, Takano M, Tsuchiya K, Hayashida T, Oka S, Gatanaga H. Single nucleotide polymorphisms in ABCC2 associate with tenofovir-induced kidney tubular dysfunction in Japanese patients with HIV-1 infection: a pharmacogenetic study. Clin Infect Dis. 2012;55(11):1558–1567. doi:10.1093/cid/cis772

743. An P, Penugonda S, Thorball CW, Bartha I, Goedert JJ, Donfield S, Buchbinder S, Binns-Roemer E, Kirk GD, Zhang W, et al. Role of APOBEC3F Gene Variation in HIV-1 Disease Progression and Pneumocystis Pneumonia. PLoS Genet. 2016;12(3):e1005921. doi:10.1371/journal.pgen.1005921

744. Lindo J, Nogueira C, Soares R, Cunha N, Almeida MR, Rodrigues L, Coelho P, Rodrigues F, Cunha RA, Gonçalves T. Genetic Polymorphisms of P2RX7 but Not of ADORA2A Are Associated with the Severity of SARS-CoV-2 Infection. Int J Mol Sci. 2024;25(11):6135. doi:10.3390/ijms25116135

745. Zacharioudaki M, Messaritakis I, Galanakis E. Vitamin D receptor, vitamin D binding protein and CYP27B1 single nucleotide polymorphisms and susceptibility to viral infections in infants. Sci Rep. 2021;11(1):13835. doi:10.1038/s41598-021-93243-3

746. Scherrmann JM. Intracellular ABCB1 as a Possible Mechanism to Explain the Synergistic Effect of Hydroxychloroquine-Azithromycin Combination in COVID-19 Therapy. AAPS J. 2020;22(4):86. doi:10.1208/s12248-020-00465-w

747. Bauer R, Rauch I. The NAIP/NLRC4 inflammasome in infection and pathology. Mol Aspects Med. 2020;76:100863. doi:10.1016/j.mam.2020.100863

748. Hou Y, Cui Y, Zhou Z, Liu H, Zhang H, Ding Y, Nie H, Ji HL. Upregulation of the WNK4 Signaling Pathway Inhibits Epithelial Sodium Channels of Mouse Tracheal Epithelial Cells After Influenza A Infection. Front Pharmacol. 2019;10:12. doi:10.3389/fphar.2019.00012

749. Kutluay SB, Perez-Caballero D, Bieniasz PD. Fates of retroviral core components during unrestricted and TRIM5-restricted infection. PLoS Pathog. 2013;9(3):e1003214. doi:10.1371/journal.ppat.1003214

750. Wu J, Gao H, Rui H, Xu P, Ni L, Zhang J, Wang L.Exploring the role of YBX3 in PEDV infection through the utilization of YBX3 knockout and overexpression cell lines. Virus Genes. 2024. doi:10.1007/s11262-024-02109-z

751. Ma C, Li S, Yang F, Cao W, Liu H, Feng T, Zhang K, Zhu Z, Liu X, Hu Y, et al. FoxJ1 inhibits African swine fever virus replication and viral S273R protein decreases the expression of FoxJ1 to impair its antiviral effect. Virol Sin. 2022;37(3):445–454. doi:10.1016/j.virs.2022.04.008

752. Lee MT, Warren MK. CSF-1-induced resistance to viral infection in murine macrophages. J Immunol. 1987;138(9):3019–3022.

753. Cornet V, Khuyen TD, Mandiki SNM, Betoulle S, Bossier P, Reyes-López FE, Tort L, Kestemont P. GAS1: A New β-Glucan Immunostimulant Candidate to Increase Rainbow Trout (Oncorhynchus mykiss) Resistance to Bacterial Infections With Aeromonas salmonicida achromogenes. Front Immunol. 2021;12:693613. doi:10.3389/fimmu.2021.693613

754. Dong X, Hong Z, Chatterjee J, Kim S, Verma DP. Expression of callose synthase genes and its connection with Npr1 signaling pathway during pathogen infection. Planta. 2008;229(1):87–98. doi:10.1007/s00425-008-0812-3

755. Liu C, Zeng X, Yu S, Ren L, Sun X, Long Y, Wang X, Lu S, Song Y, et al. Up-regulated DNA-binding inhibitor Id3 promotes differentiation of regulatory T cell to influence antiviral immunity in chronic hepatitis B virus infection. Life Sci. 2021;285:119991. doi:10.1016/j.lfs.2021.119991

756. Mlambo ZP, Sebitloane M, Naicker T. Association of angiogenic factors (placental growth factor and soluble FMS-like tyrosine kinase-1) in preeclamptic women of African ancestry comorbid with HIV infection. Arch Gynecol Obstet. 2024. doi:10.1007/s00404-024-07590-3

757. Wu B, Ramaiah A, Garcia G Jr, Hasiakos S, Arumugaswami V, Srikanth S. ORAI1 Limits SARS-CoV-2 Infection by Regulating Tonic Type I IFN Signaling. J Immunol. 2022;208(1):74–84. doi:10.4049/jimmunol.2100742

758. Cao W, Oldstone MB, De La Torre JC. Viral persistent infection affects both transcriptional and posttranscriptional regulation of neuron-specific molecule GAP43. Virology. 1997;230(2):147–154. doi:10.1006/viro.1997.8458

759. Li Q, Wilkie AR, Weller M, Liu X, Cohen JI. THY-1 Cell Surface Antigen (CD90) Has an Important Role in the Initial Stage of Human Cytomegalovirus Infection. PLoS Pathog. 2015;11(7):e1004999. doi:10.1371/journal.ppat.1004999

760. Martins JSCC, Sousa TDC, Oliveira MLA, Gimba ERP, Siqueira MM, Matos ADR. Total Osteopontin and Its Isoform OPN4 Are Differently Expressed in Respiratory Samples during Influenza A(H1N1)pdm09 Infection and Progression. Microorganisms. 2023;11(5):1349.doi:10.3390/microorganisms11051349

761. Kundu S, Ramshankar V, Verma AK, Thangaraj SV, Krishnamurthy A, Kumar R, Kannan R, Ghosh SK. Association of DFNA5, SYK, and NELL1 variants along with HPV infection in oral cancer among the prolonged tobacco-chewers. Tumour Biol. 2018;40(8):1010428318793023. doi:10.1177/1010428318793023

762. Hiller BE, Yin Y, Perng YC, de Araujo Castro Í, Fox LE, Locke MC, Monte KJ, López CB, Ornitz DM, et al. Fibroblast growth factor-9 expression in airway epithelial cells amplifies the type I interferon response and alters influenza A virus pathogenesis. PLoS Pathog. 2022;18(6):e1010228. doi:10.1371/journal.ppat.1010228

763. Fatemi SH, Sidwell R, Kist D, Akhter P, Meltzer HY, Bailey K, Thuras P, Sedgwick J. Differential expression of synaptosome-associated protein 25 kDa [SNAP-25] in hippocampi of neonatal mice following exposure to human influenza virus in utero. Brain Res. 1998;800(1):1–9. doi:10.1016/s0006-8993(98)00450-8

764. de Antonellis P, Ferrucci V, Miceli M, Bibbo F, Asadzadeh F, Gorini F, Mattivi A, Boccia A, Russo R, Andolfo I, Lasorsa VA, et al. Targeting ATP2B1 impairs PI3K/Akt/FOXO signaling and reduces SARS-COV-2 infection and replication. EMBO Rep. 2024;25(7):2974–3007. doi:10.1038/s44319-024-00164-z

765. Heath RJ, Leong JM, Visegrády B, Machesky LM, Xavier RJ. Bacterial and host determinants of MAL activation upon EPEC infection: the roles of Tir, ABRA, and FLRT3. PLoS Pathog. 2011;7(4):e1001332. doi:10.1371/journal.ppat.1001332

766. Sun X, Wang T, Wang Y, Ai K, Pan G, Li Y, Zhou C, He S, Cong H. Downregulation of lncRNA-11496 in the Brain Contributes to Microglia Apoptosis via Regulation of Mef2c in Chronic T. gondii Infection Mice. Front Mol Neurosci. 2020;13:77. doi:10.3389/fnmol.2020.00077

767. Maruta H, He H. PAK1-blockers: Potential Therapeutics against COVID-19. Med Drug Discov. 2020;6:100039. doi:10.1016/j.medidd.2020.100039

768. Bhosle VK, Sun C, Patel S, Ho TWW, Westman J, Ammendolia DA, Langari FM, Fine N, Toepfner N, Li Z, et al. The chemorepellent, SLIT2, bolsters innate immunity against Staphylococcus aureus. Elife. 2023;12:e87392. doi:10.7554/eLife.87392

769. Zhu W, Zhu W, Wang S, Liu S, Zhang H. UCHL1 deficiency upon HCMV infection induces vascular endothelial inflammatory injury mediated by mitochondrial iron overload. Free Radic Biol Med. 2024;211:96–113. doi:10.1016/j.freeradbiomed.2023.12.002

770. Goto H, Kariya R, Kudo E, Katano H, Okada S. PAX5 functions as a tumor suppressor by RB-E2F-mediated cell cycle arrest in Kaposi sarcoma-associated herpesvirus-infected primary effusion lymphoma. Neoplasia. 2024;56:101035. doi:10.1016/j.neo.2024.101035

771. Weindel CG, Bell SL, Vail KJ, West KO, Patrick KL, Watson RO. LRRK2 maintains mitochondrial homeostasis and regulates innate immune responses to Mycobacterium tuberculosis. Elife. 2020;9:e51071. doi:10.7554/eLife.51071

772. Burberry A, Wells MF, Limone F, Couto A, Smith KS, Keaney J, Gillet G, van Gastel N, Wang JY, Pietilainen O, et al. C9orf72 suppresses systemic and neural inflammation induced by gut bacteria. Nature. 2020;582(7810):89–94. doi:10.1038/s41586-020-2288-7

773. Pisapia R, Capoluongo N, Palmiero G, Tascini C, Rescigno C. Relapsing Neurological Complications in a Child With ATP1A3 Gene Mutation and Influenza Infection: A Case Report. Front Neurol. 2021;12:774054. doi:10.3389/fneur.2021.774054

774. Xiong M, Liu X, Liang T, Ban Y, Liu Y, Zhang L, Xu Z, Song C. The Alpha-1 Subunit of the Na+/K+-ATPase (ATP1A1) Is a Host Factor Involved in the Attachment of Porcine Epidemic Diarrhea Virus. Int J Mol Sci. 2023;24(4):4000. doi:10.3390/ijms24044000

775. Beshara R, Sencio V, Soulard D, Barthélémy A, Fontaine J, Pinteau T, Deruyter L, Ismail MB, Paget C, Sirard JC, et al. Alteration of Flt3-Ligand-dependent de novo generation of conventional dendritic cells during influenza infection contributes to respiratory bacterial superinfection. PLoS Pathog. 2018;14(10):e1007360. doi:10.1371/journal.ppat.1007360

776. Takeuchi Y, Tsuge M, Tsushima K, Suehiro Y, Fujino H, Ono A, Yamauchi M, Makokha GN, Nakahara T, Murakami E, et al. Signal Activation of Hepatitis B Virus-Related Hepatocarcinogenesis by Up-regulation of SUV39h1. J Infect Dis. 2020;222(12):2061–2070. doi:10.1093/infdis/jiaa317

777. Wu J, Ma S, Sandhoff R, Ming Y, Hotz-Wagenblatt A, Timmerman V, Bonello-Palot N, Schlotter-Weigel B, Auer-Grumbach M, Seeman P, et al. Loss of Neurological Disease HSAN-I-Associated Gene SPTLC2 Impairs CD8+ T Cell Responses to Infection by Inhibiting T Cell Metabolic Fitness. Immunity. 2019;50(5):1218–1231.e5. doi:10.1016/j.immuni.2019.03.005

778. Mendonça VR, Souza LC, Garcia GC, Magalhães BM, Lacerda MV, Andrade BB, Gonçalves MS, Barral-Netto M. DDX39B (BAT1), TNF and IL6 gene polymorphisms and association with clinical outcomes of patients with Plasmodium vivax malaria. Malar J. 2014;13:278. doi:10.1186/1475-2875-13-278

779. Wang M, Yang XY, Zhang NZ, Zhang DL, Zhu XQ. Evaluation of Protective Immune Responses Induced by Recombinant TrxLp and ENO2 Proteins against Toxoplasma gondii Infection in BALB/c Mice. Biomed Res Int. 2016;2016:3571962. doi:10.1155/2016/3571962

780. Zankharia U, Yi Y, Lu F, Vladimirova O, Karisetty BC, Wikramasinghe J, Kossenkov A, Collman RG, Lieberman PM. HIV-induced RSAD2/Viperin supports sustained infection of monocyte-derived macrophages. J Virol. 2024;98(10):e0086324. doi:10.1128/jvi.00863-24

781. Schütz M, Cordsmeier A, Wangen C, Horn AHC, Wyler E, Ensser A, Sticht H, Marschall M. The Interactive Complex between Cytomegalovirus Kinase vCDK/pUL97 and Host Factors CDK7-Cyclin H Determines Individual Patterns of Transcription in Infected Cells. Int J Mol Sci. 2023;24(24):17421. doi:10.3390/ijms242417421

782. Ferreira P, Dâmaso F, Anastácio M, Pinto F, Lynce A. Immune-Mediated Necrotizing Myopathy With Anti-3-Hydroxy-3-Methylglutaryl-CoA Reductase (HMGCR) Antibodies Following Viral Infection and Without Association With Statin Use: A Case Report. Cureus. 2024;16(9):e70281. doi:10.7759/cureus.70281

783. Leber A, Bassaganya-Riera J, Tubau-Juni N, Zoccoli-Rodriguez V, Lu P, Godfrey V, Kale S, Hontecillas R. Lanthionine Synthetase C-Like 2 Modulates Immune Responses to Influenza Virus Infection. Front Immunol. 2017;8:178. doi:10.3389/fimmu.2017.00178

784. Mashayekhi F, Shabani S, Sasani ST, Salehi Z. The association of stem cell factor and soluble c-Kit (s-cKit) receptor serum concentrations with the severity and risk prediction of autism spectrum disorders. Metab Brain Dis. 2022;37(3):619–624. doi:10.1007/s11011-021-00883-5

785. Kameno Y, Iwata K, Matsuzaki H, Miyachi T, Tsuchiya KJ, Matsumoto K, Iwata Y, Suzuki K, Nakamura K, Maekawa M, et al. Serum levels of soluble platelet endothelial cell adhesion molecule-1 and vascular cell adhesion molecule-1 are decreased in subjects with autism spectrum disorder. Mol Autism. 2013;4(1):19. doi:10.1186/2040-2392-4-19

786. Rahbar MH, Samms-Vaughan M, Saroukhani S, Bressler J, Hessabi M, Grove ML, Shakspeare-Pellington S, Loveland KA, Beecher C, McLaughlin W. Associations of Metabolic Genes (GSTT1, GSTP1, GSTM1) and Blood Mercury Concentrations Differ in Jamaican Children with and without Autism Spectrum Disorder. Int J Environ Res Public Health. 2021;18(4):1377. doi:10.3390/ijerph18041377

787. Akköprü H, Alnak A, Karadoğan ZN, Çağlayan AO, Özçetin M, Coşkun M. Peripheral Expression of ADORA2A Is Increased and Is Correlated with Autism Spectrum Disorder Severity in a Sample of Turkish Children. Psychiatry Clin Psychopharmacol. 2023;33(1):14–19. doi:10.5152/pcp.2023.22509

788. Viggiano M, D’Andrea T, Cameli C, Posar A, Visconti P, Scaduto MC, Colucci R, Rochat MJ, Ceroni F, Milazzo G, et al. Contribution of CACNA1H Variants in Autism Spectrum Disorder Susceptibility. Front Psychiatry. 2022;13:858238. doi:10.3389/fpsyt.2022.858238

789. Kim YJ, Park JK, Kang WS, Kim SK, Park HJ, Nam M, Kim JW. LAMB1 polymorphism is associated with autism symptom severity in Korean autism spectrum disorder patients. Nord J Psychiatry. 2015;69(8):594–598. doi:10.3109/08039488.2015.102259

790. Szczurkowska J, Pischedda F, Pinto B, Managò F, Haas CA, Summa M, Bertorelli R, Papaleo F, Schäfer MK, Piccoli G, et al. NEGR1 and FGFR2 cooperatively regulate cortical development and core behaviours related to autism disorders in mice. Brain. 2018;141(9):2772–2794. doi:10.1093/brain/awy190

791. Bolognesi E, Guerini FR, Carta A, Chiappedi M, Sotgiu S, Mensi MM, Agliardi C, Zanzottera M, Clerici M. The Role of SNAP-25 in Autism Spectrum Disorders Onset Patterns. Int J Mol Sci. 2023;24(18):14042. doi:10.3390/ijms241814042

792. Li H, Yamagata T, Mori M, Momoi MY. Association of autism in two patients with hereditary multiple exostoses caused by novel deletion mutations of EXT1. J Hum Genet. 2002;47(5):262–265. doi:10.1007/s100380200036

793. Seiffert S, Pendziwiat M, Bierhals T, Goel H, Schwarz N, van der Ven A, Boßelmann CM, Lemke J, Syrbe S, Willemsen MH, et al. Modulating effects of FGF12 variants on NaV1.2 and NaV1.6 being associated with developmental and epileptic encephalopathy and Autism spectrum disorder: A case series. EBioMedicine. 2022;83:104234. doi:10.1016/j.ebiom.2022.104234

794. Basu S, Ro EJ, Liu Z, Kim H, Bennett A, Kang S, Suh H. The Mef2c Gene Dose-Dependently Controls Hippocampal Neurogenesis and the Expression of Autism-Like Behaviors. J Neurosci. 2024;44(5):e1058232023. doi:10.1523/JNEUROSCI.1058-23.2023

795. Floris C, Rassu S, Boccone L, Gasperini D, Cao A, Crisponi L. Two patients with balanced translocations and autistic disorder: CSMD3 as a candidate gene for autism found in their common 8q23 breakpoint area. Eur J Hum Genet. 2008;16(6):696–704. doi:10.1038/ejhg.2008.7

796. Alvarez Retuerto AI, Cantor RM, Gleeson JG, Ustaszewska A, Schackwitz WS, Pennacchio LA, Geschwind DH. Association of common variants in the Joubert syndrome gene (AHI1) with autism. Hum Mol Genet. 2008;17(24):3887–3896. doi:10.1093/hmg/ddn291

797. Kruth KA, Grisolano TM, Ahern CA, Williams AJ. SCN2A channelopathies in the autism spectrum of neuropsychiatric disorders: a role for pluripotent stem cells?. Mol Autism. 2020;11(1):23. doi:10.1186/s13229-020-00330-9

798. Jang WE, Park JH, Park G, Bang G, Na CH, Kim JY, Kim KY, Kim KP, Shin CY, An JY, et al. Cntnap2-dependent molecular networks in autism spectrum disorder revealed through an integrative multi-omics analysis. Mol Psychiatry. 2023;28(2):810–821. doi:10.1038/s41380-022-01822-1

799. Coskunpinar EM, Tur S, Cevher Binici N, Yazan Songür C. Association of GABRG3, GABRB3, HTR2A gene variants with autism spectrum disorder. Gene. 2023;870:147399. doi:10.1016/j.gene.2023.147399 800.

800. Papp-Hertelendi R, Tényi T, Hadzsiev K, Hau L, Benyus Z, Csábi G. First report on the association of SCN1A mutation, childhood schizophrenia and autism spectrum disorder without epilepsy. Psychiatry Res. 2018;270:1175–1176. doi:10.1016/j.psychres.2018.07.028

801. Matrone C, Ferretti G. Semaphorin 3A influences neuronal processes that are altered in patients with autism spectrum disorder: Potential diagnostic and therapeutic implications. Neurosci Biobehav Rev. 2023;153:105338. doi:10.1016/j.neubiorev.2023.105338

802. Çetin İ, Tezdiğ İ, Tarakçioğlu MC, Kadak MT, Demirel ÖF, Özer ÖF, Erdoğan F, Doğangün B. Do Low Serum UCH-L1 and TDP-43 Levels Indicate Disturbed Ubiquitin-Proteosome System in Autism Spectrum Disorder?. Noro Psikiyatr Ars. 2017;54(3):267–271. doi:10.5152/npa.2017.14873

803. Moreno-Ramos OA, Olivares AM, Haider NB, de Autismo LC, Lattig MC. Whole-Exome Sequencing in a South American Cohort Links ALDH1A3, FOXN1 and Retinoic Acid Regulation Pathways to Autism Spectrum Disorders. PLoS One. 2015;10(9):e0135927. doi:10.1371/journal.pone.0135927

804. Kaiser FMP, Gruenbacher S, Oyaga MR, Nio E, Jaritz M, Sun Q, van der Zwaag W, Kreidl E, Zopf LM, Dalm VASH, et al. Biallelic PAX5 mutations cause hypogammaglobulinemia, sensorimotor deficits, and autism spectrum disorder. J Exp Med. 2022;219(9):e20220498. doi:10.1084/jem.20220498

805. Pescucci C, Meloni I, Bruttini M, Ariani F, Longo I, Mari F, Canitano R, Hayek G, Zappella M, Renieri A. Chromosome 2 deletion encompassing the MAP2 gene in a patient with autism and Rett-like features. Clin Genet. 2003;64(6):497–501. doi:10.1046/j.1399-0004.2003.00176.x

806. Fang X, Fee T, Davis J, Stolerman ES, Caylor RC. Clinical case report: mosaic ANK3 pathogenic variant in a patient with autism spectrum disorder and neurodevelopmental delay. Cold Spring Harb Mol Case Stud. 2023;9(3):a006233.. doi:10.1101/mcs.a006233

807. Labonne JDJ, Driessen TM, Harris ME, Kong IK, Brakta S, Theisen J, Sangare M, Layman LC, Kim CH, Lim J, et al. Comparative Genomic Mapping Implicates LRRK2 for Intellectual Disability and Autism at 12q12, and HDHD1, as Well as PNPLA4, for X-Linked Intellectual Disability at Xp22.31. J Clin Med. 2020;9(1):274. doi:10.3390/jcm9010274

808. Cooper JN, Mittal J, Sangadi A, Klassen DL, King AM, Zalta M, Mittal R, Eshraghi AA. Landscape of NRXN1 Gene Variants in Phenotypic Manifestations of Autism Spectrum Disorder: A Systematic Review. J Clin Med. 2024;13(7):2067. doi:10.3390/jcm13072067

809. Yang W, Liu J, Zheng F, Jia M, Zhao L, Lu T, Ruan Y, Zhang J, Yue W, Zhang D, et al. The evidence for association of ATP2B2 polymorphisms with autism in Chinese Han population. PLoS One. 2013;8(4):e61021. doi:10.1371/journal.pone.0061021

810. Soueid J, Kourtian S, Makhoul NJ, Makoukji J, Haddad S, Ghanem SS, Kobeissy F, Boustany RM. RYR2, PTDSS1 and AREG genes are implicated in a Lebanese population-based study of copy number variation in autism. Sci Rep. 2016;6:19088. doi:10.1038/srep19088

811. Shin S, Santi A, Huang S. Conditional Pten knockout in parvalbumin- or somatostatin-positive neurons sufficiently leads to autism-related behavioral phenotypes. Mol Brain. 2021;14(1):24. doi:10.1186/s13041-021-00731-8

812. Janickova L, Rechberger KF, Wey L, Schwaller B. Absence of parvalbumin increases mitochondria volume and branching of dendrites in inhibitory Pvalb neurons in vivo: a point of convergence of autism spectrum disorder (ASD) risk gene phenotypes. Mol Autism. 2020;11(1):47. doi:10.1186/s13229-020-00323-8

813. Bitar T, Hleihel W, Marouillat S, Vonwill S, Vuillaume ML, Soufia M, Vourc’h P, Laumonnier F, Andres CR. Identification of rare copy number variations reveals PJA2, APCS, SYNPO, and TAC1 as novel candidate genes in Autism Spectrum Disorders. Mol Genet Genomic Med. 2019;7(8):e786. doi:10.1002/mgg3.786

814. Taketomi T, Tsuruta F. Mutations in Hevin/Sparcl1 and risk of autism spectrum disorder. Neural Regen Res. 2023;18(7):1499–1500. doi:10.4103/1673-5374.361543

815. Chien WH, Gau SS, Liao HM, Chiu YN, Wu YY, Huang YS, Tsai WC, Tsai HM, Chen CH. Deep exon resequencing of DLGAP2 as a candidate gene of autism spectrum disorders. Mol Autism. 2013;4(1):26. doi:10.1186/2040-2392-4-26

816. Torres A, Brownstein CA, Tembulkar SK, Graber K, Genetti C, Kleiman RJ, Sweadner KJ, Mavros C, Liu KX, Smedemark-Margulies N, et al. De novo ATP1A3 and compound heterozygous NLRP3 mutations in a child with autism spectrum disorder, episodic fatigue and somnolence, and muckle-wells syndrome. Mol Genet Metab Rep. 2018;16:23–29. doi:10.1016/j.ymgmr.2018.06.001

817. Sriwimol W, Limprasert P. Significant Changes in Plasma Alpha-Synuclein and Beta-Synuclein Levels in Male Children with Autism Spectrum Disorder. Biomed Res Int. 2018;2018:4503871. doi:10.1155/2018/4503871

818. Iwata K, Matsuzaki H, Tachibana T, Ohno K, Yoshimura S, Takamura H, Yamada K, Matsuzaki S, Nakamura K, Tsuchiya KJ, et al. N-ethylmaleimide-sensitive factor interacts with the serotonin transporter and modulates its trafficking: implications for pathophysiology in autism. Mol Autism. 2014;5:33. doi:10.1186/2040-2392-5-33

819. Zhubi A, Chen Y, Guidotti A, Grayson DR. Epigenetic regulation of RELN and GAD1 in the frontal cortex (FC) of autism spectrum disorder (ASD) subjects. Int J Dev Neurosci. 2017;62:63–72. doi:10.1016/j.ijdevneu.2017.02.003

820. Wang X, Sun Z, Yang T, Lin F, Ye S, Yan J, Li T, Chen J. Sodium butyrate facilitates CRHR2 expression to alleviate HPA axis hyperactivity in autism-like rats induced by prenatal lipopolysaccharides through histone deacetylase inhibition. mSystems. 2023;8(4):e0041523. doi:10.1128/msystems.00415-23

821. Orabona GM, Griesi-Oliveira K, Vadasz E, Bulcão VL, Takahashi VN, Moreira ES, Furia-Silva M, Ros-Melo AM, Dourado F, Matioli SR, et al. HTR1B and HTR2C in autism spectrum disorders in Brazilian families. Brain Res. 2009;1250:14–19. doi:10.1016/j.brainres.2008.11.007

822. Baris RO, Sahin N, Bilgic AD, Ozdemir C, Edgunlu TG. Molecular and in silico analyses of SYN III gene variants in autism spectrum disorder. Ir J Med Sci. 2023;192(6):2887–2895. doi:10.1007/s11845-023-03402-w

823. Dohrn MF, Bademci G, Rebelo AP, Jeanne M, Borja NA, Beijer D, Danzi MC, Bivona SA, Gueguen P, Zafeer MF; et al. Recurrent ATP1A1 variant Gly903Arg causes developmental delay, intellectual disability, and autism. Ann Clin Transl Neurol. 2024;11(4):1075–1079. doi:10.1002/acn3.51963

824. Ali G, Shin KC, Habbab W, Alkhadairi G, AbdelAleem A, AlShaban FA, Park Y, Stanton LW. Characterization of a loss-of-function NSF attachment protein beta mutation in monozygotic triplets affected with epilepsy and autism using cortical neurons from proband-derived and CRISPR-corrected induced pluripotent stem cell lines. Front Neurosci. 2024;17:1302470. doi:10.3389/fnins.2023.1302470

825. Pavinato L, Stanic J, Barzasi M, Gurgone A, Chiantia G, Cipriani V, Eberini I, Palazzolo L, Di Luca M, Costa A, et al. Missense variants in RPH3A cause defects in excitatory synaptic function and are associated with a clinically variable neurodevelopmental disorder. Genet Med. 2023;25(11):100922. doi:10.1016/j.gim.2023.100922

826. Soueid J, Hamze Z, Bedran J, Chahrour M, Boustany RM. A novel autism-associated UBLCP1 mutation impacts proteasome regulation/activity. Transl Psychiatry. 2023;13(1):404. doi:10.1038/s41398-023-02702-0

827. Wang M, Yang XY, Zhang NZ, Zhang DL, Zhu XQ. Evaluation of Protective Immune Responses Induced by Recombinant TrxLp and ENO2 Proteins against Toxoplasma gondii Infection in BALB/c Mice. Biomed Res Int. 2016;2016:3571962. doi:10.1155/2016/3571962

828. Aoki Y, Cortese S. Mitochondrial Aspartate/Glutamate Carrier SLC25A12 and Autism Spectrum Disorder: a Meta-Analysis. Mol Neurobiol. 2016;53(3):1579–1588. doi:10.1007/s12035-015-9116-3

829. Kantojärvi K, Kotala I, Rehnström K, Ylisaukko-Oja T, Vanhala R, von Wendt TN, von Wendt L, Järvelä I. Fine mapping of Xq11.1-q21.33 and mutation screening of RPS6KA6, ZNF711, ACSL4, DLG3, and IL1RAPL2 for autism spectrum disorders (ASD). Autism Res. 2011;4(3):228–233. doi:10.1002/aur.187

830. Kwon SJ, Hong KW, Choi S, Hong JS, Kim JW, Kim JW, Lee HJ, Jang HB, Yum KS. Association of 3-hydroxy-3-methylglutaryl-coenzyme A reductase gene polymorphism with obesity and lipid metabolism in children and adolescents with autism spectrum disorder. Metab Brain Dis. 2022;37(2):319–328. doi:10.1007/s11011-021-00877-3

831. Shimamoto C, Ohnishi T, Maekawa M, Watanabe A, Ohba H, Arai R, Iwayama Y, Hisano Y, Toyota T, Toyoshima M, et al. Functional characterization of FABP3, 5 and 7 gene variants identified in schizophrenia and autism spectrum disorder and mouse behavioral studies. Hum Mol Genet. 2014;23(24):6495–6511. doi:10.1093/hmg/ddu369

832. Corti S, Nizzardo M, Nardini M, Donadoni C, Salani S, Simone C, Falcone M, Riboldi G, Govoni A, Bresolin N, et al. Systemic transplantation of c-kit+ cells exerts a therapeutic effect in a model of amyotrophic lateral sclerosis. Hum Mol Genet. 2010;19(19):3782–3796. doi:10.1093/hmg/ddq293

833. Yamada S, Hashizume A, Hijikata Y, Ito D, Kishimoto Y, Iida M, Koike H, Hirakawa A, Katsuno M. Ratio of urinary N-terminal titin fragment to urinary creatinine is a novel biomarker for amyotrophic lateral sclerosis. J Neurol Neurosurg Psychiatry. 2021;92(10):1072–1079. doi:10.1136/jnnp-2020-324615

834. Capponi S, Geuens T, Geroldi A, Origone P, Verdiani S, Cichero E, Adriaenssens E, De Winter V, Bandettini di Poggio M, Barberis M, et al. Molecular Chaperones in the Pathogenesis of Amyotrophic Lateral Sclerosis: The Role of HSPB1. Hum Mutat. 2016;37(11):1202–1208. doi:10.1002/humu.23062

835. Ajroud-Driss S, Saeed M, Khan H, Siddique N, Hung WY, Sufit R, Heller S, Armstrong J, Casey P, Siddique T, et al. Riluzole metabolism and CYP1A1/2 polymorphisms in patients with ALS. Amyotroph Lateral Scler. 2007;8(5):305–309. doi:10.1080/17482960701500650

836. Fang L, Teuchert M, Huber-Abel F, Schattauer D, Hendrich C, Dorst J, Zettlmeissel H, Wlaschek M, Scharffetter-Kochanek K, Kapfer T, et al. MMP-2 and MMP-9 are elevated in spinal cord and skin in a mouse model of ALS. J Neurol Sci. 2010;294(1-2):51–56. doi:10.1016/j.jns.2010.04.005

837. Li X, Guan Y, Chen Y, Zhang C, Shi C, Zhou F, Yu L, Juan J, Wang X. Expression of Wnt5a and its receptor Fzd2 is changed in the spinal cord of adult amyotrophic lateral sclerosis transgenic mice. Int J Clin Exp Pathol. 2013;6(7):1245–1260.

838. Gupta PK, Prabhakar S, Abburi C, Sharma NK, Anand A. Vascular endothelial growth factor-A and chemokine ligand (CCL2) genes are upregulated in peripheral blood mononuclear cells in Indian amyotrophic lateral sclerosis patients. J Neuroinflammation. 2011;8:114. doi:10.1186/1742-2094-8-114

839. Milani M, Mammarella E, Rossi S, Miele C, Lattante S, Sabatelli M, Cozzolino M, D’Ambrosi N, Apolloni S. Targeting S100A4 with niclosamide attenuates inflammatory and profibrotic pathways in models of amyotrophic lateral sclerosis. J Neuroinflammation. 2021;18(1):132. doi:10.1186/s12974-021-02184-1

840. Si Y, Kazamel M, Kwon Y, Lee I, Anderson T, Zhou S, Bamman M, Wiggins D, Kwan T, King PH. The vitamin D activator CYP27B1 is upregulated in muscle fibers in denervating disease and can track progression in amyotrophic lateral sclerosis. J Steroid Biochem Mol Biol. 2020;200:105650. doi:10.1016/j.jsbmb.2020.105650

841. Qosa H, Lichter J, Sarlo M, Markandaiah SS, McAvoy K, Richard JP, Jablonski MR, Maragakis NJ, Pasinelli P, Trotti D. Astrocytes drive upregulation of the multidrug resistance transporter ABCB1 (P-Glycoprotein) in endothelial cells of the blood-brain barrier in mutant superoxide dismutase 1-linked amyotrophic lateral sclerosis. Glia. 2016;64(8):1298–1313. doi:10.1002/glia.23003

842. Rzhepetskyy Y, Lazniewska J, Blesneac I, Pamphlett R, Weiss N. CACNA1H missense mutations associated with amyotrophic lateral sclerosis alter Cav3.2 T-type calcium channel activity and reticular thalamic neuron firing. Channels (Austin). 2016;10(6):466–477. doi:10.1080/19336950.2016.1204497

843. Trias E, Kovacs M, King PH, Si Y, Kwon Y, Varela V, Ibarburu S, Moura IC, Hermine O, Beckman JS, et al. Schwann cells orchestrate peripheral nerve inflammation through the expression of CSF1, IL-34, and SCF in amyotrophic lateral sclerosis. Glia. 2020;68(6):1165–1181. doi:10.1002/glia.23768

844. Iłzecka J. Decreased serum endoglin level in patients with amyotrophic lateral sclerosis: a preliminary report. Scand J Clin Lab Invest. 2008;68(4):348–351. doi:10.1080/00365510701604628

845. Noda Y, Tanaka M, Nakamura S, Ito J, Kakita A, Hara H, Shimazawa M. Identification of VGF nerve growth factor inducible-producing cells in human spinal cords and expression change in patients with amyotrophic lateral sclerosis. Int J Med Sci. 2020;17(4):480–489. doi:10.7150/ijms.39101 846.

846. Ikemoto A, Hirano A, Akiguchi I. Increased expression of growth-associated protein 43 on the surface of the anterior horn cells in amyotrophic lateral sclerosis. Acta Neuropathol. 1999;98(4):367–373. doi:10.1007/s004010051096

847. Liu Y, Yan D, Yang L, Chen X, Hu C, Chen M. Stathmin 2 is a potential treatment target for TDP-43 proteinopathy in amyotrophic lateral sclerosis. Transl Neurodegener. 2024;13(1):20. doi:10.1186/s40035-024-00413-0

848. Arosio A, Sala G, Rodriguez-Menendez V, Grana D, Gerardi F, Lunetta C, Ferrarese C, Tremolizzo L. MEF2D and MEF2C pathways disruption in sporadic and familial ALS patients. Mol Cell Neurosci. 2016;74:10–17. doi:10.1016/j.mcn.2016.02.002

849. Birger A, Ottolenghi M, Perez L, Reubinoff B, Behar O. ALS-related human cortical and motor neurons survival is differentially affected by Sema3A. Cell Death Dis. 2018;9(3):256. doi:10.1038/s41419-018-0294-6

850. Li R, Wang J, Xie W, Liu J, Wang C. UCHL1 from serum and CSF is a candidate biomarker for amyotrophic lateral sclerosis. Ann Clin Transl Neurol. 2020;7(8):1420–1428. doi:10.1002/acn3.51141

851. Silvestri B, Mochi M, Mawrie D, de Turris V, Colantoni A, Borhy B, Medici M, Anderson EN, Garone MG, Zammerilla CP, et al. HuD (ELAVL4) gain-of-function impairs neuromuscular junctions and induces apoptosis in in vitro and in vivo models of amyotrophic lateral sclerosis. Preprint. bioRxiv. 2024;2023.08.22.554258. doi:10.1101/2023.08.22.554258

852. Whittle AJ, Ross OA, Naini A, Gordon P, Mistumoto H, Dächsel JC, Stone JT, Wszolek ZK, Farrer MJ, Przedborski S. Pathogenic Lrrk2 substitutions and Amyotrophic lateral sclerosis. J Neural Transm (Vienna). 2007;114(3):327–329. doi:10.1007/s00702-006-0525-3

853. van Doormaal PT, Ticozzi N, Gellera C, Ratti A, Taroni F, Chiò A, Calvo A, Mora G, Restagno G, Traynor BJ, et al. Analysis of the KIFAP3 gene in amyotrophic lateral sclerosis: a multicenter survival study. Neurobiol Aging. 2014;35(10):2420.e13-2420.e14. doi:10.1016/j.neurobiolaging.2014.04.014

854. Syeda SB, Lone MA, Mohassel P, Donkervoort S, Munot P, França MC Jr, Galarza-Brito JE, Eckenweiler M, Asamoah A, Gable K, et al. Recurrent de novo SPTLC2 variant causes childhood-onset amyotrophic lateral sclerosis (ALS) by excess sphingolipid synthesis. J Neurol Neurosurg Psychiatry. 2024;95(2):103–113. doi:10.1136/jnnp-2023-332132

855. Ruban A, Malina KC, Cooper I, Graubardt N, Babakin L, Jona G, Teichberg VI. Combined Treatment of an Amyotrophic Lateral Sclerosis Rat Model with Recombinant GOT1 and Oxaloacetic Acid: A Novel Neuroprotective Treatment. Neurodegener Dis. 2015;15(4):233–242. doi:10.1159/000382034

856. Jiang Y, Liang M, Chen L, Wang J, Huang Y, Huo H, Xiao D, Hu Y, Wang Z, Ji Q, et al. Myeloid SENP3 deficiency protects mice from diet and age-induced obesity via regulation of YAP1 SUMOylation. Cell Mol Life Sci. 2023;81(1):4. doi:10.1007/s00018-023-05050-w

857. Maehle BO, Tretli S, Thorsen T. The associations of obesity, lymph node status and prognosis in breast cancer patients: dependence on estrogen and progesterone receptor status. APMIS. 2004;112(6):349–357. doi:10.1111/j.1600-0463.2004.apm1120605.x

858. Deng Y, Qiu T, Zhang M, Wu J, Zhang X, Wang J, Chen K, Feng J, Ha X, Xie J, et al. High Level of Palmitic Acid Induced Over-Expressed Methyltransferase Inhibits Anti-Inflammation Factor KLF4 Expression in Obese Status. Inflammation. 2020;43(3):821–832. doi:10.1007/s10753-019-01168-x

859. Pan X, Chen X, Ren Q, Yue L, Niu S, Li Z, Zhu R, Chen X, Jia Z, Zhen R, et al. Single-cell transcriptomics identifies Col1a1 and Col1a2 as hub genes in obesity-induced cardiac fibrosis. Biochem Biophys Res Commun. 2022;618:30–37. doi:10.1016/j.bbrc.2022.06.018

860. Zhang W, Shi B, Li S, Liu Z, Li S, Dong S, Cheng Y, Zhu J, Zhang G, Zhong M. Sleeve gastrectomy improves lipid dysmetabolism by downregulating the USP20-HSPA2 axis in diet-induced obese mice. Front Endocrinol (Lausanne). 2022;13:1041027. doi:10.3389/fendo.2022.1041027861.

861. Pilch W, Wyrostek J, Major P, Zuziak R, Piotrowska A, Czerwińska-Ledwig O, Grzybkowska A, Zasada M, Ziemann E, Żychowska M. The effect of whole-body cryostimulation on body composition and leukocyte expression of HSPA1A, HSPB1, and CRP in obese men. Cryobiology. 2020;94:100–106. doi:10.1016/j.cryobiol.2020.04.002

862. Makey KL, Patterson SG, Robinson J, Loftin M, Waddell DE, Miele L, Chinchar E, Huang M, Smith AD, Weber M, et al. ncreased plasma levels of soluble vascular endothelial growth factor receptor 1 (sFlt-1) in women by moderate exercise and increased plasma levels of vascular endothelial growth factor in overweight/obese women. Eur J Cancer Prev. 2013;22(1):83–89. doi:10.1097/CEJ.0b013e328353ed81

863. Ye B, Chen X, Chen Y, Lin W, Xu D, Fang Z, Chattipakorn N, Huang W, Wang X, Wu G, et al. Inhibition of TAK1/TAB2 complex formation by costunolide attenuates obesity cardiomyopathy via the NF-κB signaling pathway. Phytomedicine. 2023;108:154523. doi:10.1016/j.phymed.2022.154523

864. Patel SJ, Liu N, Piaker S, Gulko A, Andrade ML, Heyward FD, Sermersheim T, Edinger N, Srinivasan H, Emont MP, et al. Hepatic IRF3 fuels dysglycemia in obesity through direct regulation of Ppp2r1b. Sci Transl Med. 2022;14(637):eabh3831. doi:10.1126/scitranslmed.abh3831

865. Hao J, Liu Z, Ju W, He F, Liu K, Wu J. Role and mechanism of FLT4 in high-fat diet-induced obesity in mice. Biochem Biophys Res Commun. 2023;675:61–70. doi:10.1016/j.bbrc.2023.06.025

866. Menghini R, Marchetti V, Cardellini M, Hribal ML, Mauriello A, Lauro D, Sbraccia P, Lauro R, Federici M. Phosphorylation of GATA2 by Akt increases adipose tissue differentiation and reduces adipose tissue-related inflammation: a novel pathway linking obesity to atherosclerosis. Circulation. 2005;111(15):1946–1953. doi:10.1161/01.CIR.0000161814.02942.B2

867. Wang Y, Cheng YS, Yin XQ, Yu G, Jia BL. Anxa2 gene silencing attenuates obesity-induced insulin resistance by suppressing the NF-κB signaling pathway. Am J Physiol Cell Physiol. 2019;316(2):C223–C234. doi:10.1152/ajpcell.00242.2018

868. DuBois BN, O’Tierney-Ginn P, Pearson J, Friedman JE, Thornburg K, Cherala G. Maternal obesity alters feto-placental cytochrome P4501A1 activity. Placenta. 2012;33(12):1045–1051. doi:10.1016/j.placenta.2012.09.008

869. Bai J, Cervantes C, Liu J, He S, Zhou H, Zhang B, Cai H, Yin D, Hu D, Li Z, et al. DsbA-L prevents obesity-induced inflammation and insulin resistance by suppressing the mtDNA release-activated cGAS-cGAMP-STING pathway. Proc Natl Acad Sci U S A. 2017;114(46):12196–12201. doi:10.1073/pnas.1708744114

870. Morgan AR, Han DY, Thompson JM, Mitchell EA, Ferguson LR. Analysis of MMP2 promoter polymorphisms in childhood obesity. BMC Res Notes. 2011;4:253. doi:10.1186/1756-0500-4-253

871. Cui Z, Liu Y, Wan W, Xu Y, Hu Y, Ding M, Dou X, Wang R, Li H, Meng Y, et al. Ethacrynic acid targets GSTM1 to ameliorate obesity by promoting browning of white adipocytes. Protein Cell. 2021;12(6):493–501. doi:10.1007/s13238-020-00717-7

872. Van de Pette M, Tunster SJ, John RM. Loss of Imprinting of Cdkn1c Protects against Age and Diet-Induced Obesity. Int J Mol Sci. 2018;19(9):2734.. Int J Mol Sci. 2018;19(9):2734. doi:10.3390/ijms19092734

873. Wu Y, Ma Y. CCL2-CCR2 signaling axis in obesity and metabolic diseases. J Cell Physiol. 2024;239(4):e31192. doi:10.1002/jcp.31192

874. Jamka M, Kaczmarek N, Mądry E, Krzyżanowska-Jankowska P, Bajerska J, Kręgielska-Narożna M, Bogdański P, Walkowiak J. Metabolic Health in Obese Subjects-Is There a Link to Lactoferrin and Lactoferrin Receptor-Related Gene Polymorphisms?. Nutrients. 2020;12(9):2843. doi:10.3390/nu12092843

875. Gómez-Úriz AM, Milagro FI, Mansego ML, Cordero P, Abete I, De Arce A, Goyenechea E, Blázquez V, Martínez-Zabaleta M, Martínez JA, et al. Obesity and ischemic stroke modulate the methylation levels of KCNQ1 in white blood cells. Hum Mol Genet. 2015;24(5):1432–1440. doi:10.1093/hmg/ddu559

876. Khadir A, Tiss A, Abubaker J, Abu-Farha M, Al-Khairi I, Cherian P, John J, Kavalakatt S, Warsame S, Al-Madhoun A, et al. MAP kinase phosphatase DUSP1 is overexpressed in obese humans and modulated by physical exercise. Am J Physiol Endocrinol Metab. 2015;308(1):E71–E83. doi:10.1152/ajpendo.00577.2013

877. Anguita-Ruiz A, Mendez-Gutierrez A, Ruperez AI, Leis R, Bueno G, Gil-Campos M, Tofe I, Gomez-Llorente C, Moreno LA, Gil Á, Aguilera CM. The protein S100A4 as a novel marker of insulin resistance in prepubertal and pubertal children with obesity. Metabolism. 2020;105:154187. doi:10.1016/j.metabol.2020.154187

878. AlSedairy SA, Al-Harbi LN, Binobead MA, Athinarayanan J, Arzoo S, Al-Tamimi DS, Shamlan G, Alshatwi AA, Periasamy VS. Association of CYP2R1 and CYP27B1 genes with the risk of obesity and vitamin D metabolism in Saudi women. J Genet Eng Biotechnol. 2023;21(1):59. doi:10.1186/s43141-023-00508-7

879. Justesen S, Bilde K, Olesen RH, Pedersen LH, Ernst E, Larsen A. ABCB1 expression is increased in human first trimester placenta from pregnant women classified as overweight or obese. Sci Rep. 2023;13(1):5175. doi:10.1038/s41598-023-31598-5

880. Sachan A, Aggarwal S, Pol MM, Singh A, Yadav R. Expression analysis of MMP14: Key enzyme action in modulating visceral adipose tissue plasticity in patients with obesity. Clin Obes. 2023;13(5):e12607. doi:10.1111/cob.12607

881. Takahashi D, Mori T, Sohara E, Tanaka M, Chiga M, Inoue Y, Nomura N, Zeniya M, Ochi H, Takeda S, et al. WNK4 is an Adipogenic Factor and Its Deletion Reduces Diet-Induced Obesity in Mice. EBioMedicine. 2017;18:118–127. doi:10.1016/j.ebiom.2017.03.011

882. Rivera-Gonzalez O, Mills MF, Konadu BD, Wilson NA, Murphy HA, Newberry MK, Hyndman KA, Garrett MR, Webb DJ, Speed JS. Adipocyte endothelin B receptor activation inhibits adiponectin production and causes insulin resistance in obese mice. Acta Physiol (Oxf). 2024;240(10):e14214. doi:10.1111/apha.14214

883. Pervanidou P, Chouliaras G, Akalestos A, Bastaki D, Apostolakou F, Papassotiriou I, Chrousos GP. Increased placental growth factor (PlGF) concentrations in children and adolescents with obesity and the metabolic syndrome. Hormones (Athens). 2014;13(3):369–374. doi:10.14310/horm.2002.1491

884. Pereira MJ, Vranic M, Kamble PG, Jernow H, Kristófi R, Holbikova E, Skrtic S, Kullberg J, Svensson MK, Hetty S, et al. CDKN2C expression in adipose tissue is reduced in type II diabetes and central obesity: impact on adipocyte differentiation and lipid storage?. Transl Res. 2022;242:105–121. doi:10.1016/j.trsl.2021.12.003

885. Folon L, Baron M, Toussaint B, Vaillant E, Boissel M, Scherrer V, Loiselle H, Leloire A, Badreddine A, Balkau B, et al. Contribution of heterozygous PCSK1 variants to obesity and implications for precision medicine: a case-control study. Lancet Diabetes Endocrinol. 2023;11(3):182–190. doi:10.1016/S2213-8587(22)00392-8

886. Hahm S, Fekete C, Mizuno TM, Windsor J, Yan H, Boozer CN, Lee C, Elmquist JK, Lechan RM, Mobbs CV, et al. VGF is required for obesity induced by diet, gold thioglucose treatment, and agouti and is differentially regulated in pro-opiomelanocortin-and neuropeptide Y-containing arcuate neurons in response to fasting. J Neurosci. 2002;22(16):6929–6938. doi:10.1523/JNEUROSCI.22-16-06929.2002

887. Bekar E, Altunkaynak BZ, Balcı K, Aslan G, Ayyıldız M, Kaplan S. Effects of high fat diet induced obesity on peripheral nerve regeneration and levels of GAP 43 and TGF-β in rats. Biotech Histochem. 2014;89(6):446–456. doi:10.3109/10520295.2014.894575

888. Al-Ameri HW, Shetty S, Rahman B, Gopalakrishnan ARK, Ismail AA, Acharya AB. Evaluation of salivary Thy-1 in health, periodontitis, and obesity. Oral Dis. 2024;30(4):2670–2677. doi:10.1111/odi.14652

889. Michaelides M, Miller ML, Egervari G, Primeaux SD, Gomez JL, Ellis RJ, Landry JA, Szutorisz H, Hoffman AF, Lupica CR, et al. Striatal Rgs4 regulates feeding and susceptibility to diet-induced obesity. Mol Psychiatry. 2020;25(9):2058–2069. doi:10.1038/s41380-018-0120-7

890. Sun Y, Wang R, Zhao S, Li W, Liu W, Tang L, Wang Z, Wang W, Liu R, Ning G, et al. FGF9 inhibits browning program of white adipocytes and associates with human obesity. J Mol Endocrinol. 2019;62(2):79–90. doi:10.1530/JME-18-0151

891. Huang T, Zhang X, Li Q, Li X, Yao J, Song J, Chen Y, Ye L, Li C, Xiran P, et al. The Association between Obesity Susceptibility and Polymorphisms of MC4R, SH2B1, and NEGR1 in Tibetans. Genet Test Mol Biomarkers. 2024;28(7):267–274. doi:10.1089/gtmb.2023.0546

892. Ceddia RP, Zurawski Z, Thompson Gray A, Adegboye F, McDonald-Boyer A, Shi F, Liu D, Maldonado J, Feng J, Li Y, et al. Gβγ-SNAP25 exocytotic brake removal enhances insulin action, promotes adipocyte browning, and protects against diet-induced obesity. J Clin Invest. 2023;133(19):e160617. doi:10.1172/JCI160617

893. Wang M, Li B, Qin F, Ye J, Jin L. Obesity induced Ext1 reduction mediates the occurrence of NAFLD. Biochem Biophys Res Commun. 2022;589:123–130. doi:10.1016/j.bbrc.2021.12.017

894. Huang L, Teng D, Wang H, Sheng G, Liu T. Association of copy number variation in the AHI1 gene with risk of obesity in the Chinese population. Eur J Endocrinol. 2012;166(4):727–734. doi:10.1530/EJE-11-0999

895. Yi H, Liu C, Shi J, Wang S, Zhang H, He Y, Tao J, Li S, Zhang R. GCG Alleviates Obesity-Induced Myocardial Fibrosis in Rats by Enhancing Expression of SCN5A. Front Cardiovasc Med. 2022;9:869279. doi:10.3389/fcvm.2022.869279

896. Naudhani S, Ahmad A, Khan Bazai F, Pervez MT, Zafar A, Shah SA, Raheem N, Baloch AH, Mushtaq M, Daud S. Missense Pathogenic Variant in a Conserved Region of CNTNAP2 Is Associated with Obesity, Seizures, and Language Impairment in a Pakistani Family. Mol Syndromol. 2023;14(4):293–302. doi:10.1159/000529427

897. Wang H, Wang Y, Zhang H, Liang Z, Hu W, Qiu S, Li K, Zhang L, Dai H, Yang M, et al. Hedgehog interacting protein as a circulating biomarker in women with obesity: a cross-sectional study and intervention studies. Ann Med. 2023;55(1):2206162. doi:10.1080/07853890.2023.2206162

898. Li Z, Shi B, Li N, Sun J, Zeng X, Huang R, Bok S, Chen X, Han J, Yallowitz AR, et al. Bone controls browning of white adipose tissue and protects from diet-induced obesity through Schnurri-3-regulated SLIT2 secretion. Nat Commun. 2024;15(1):6697. doi:10.1038/s41467-024-51155-6 899.

899. Chen X, Ruiz-Velasco A, Zou Z, Hille SS, Ross C, Fonseka O, Gare SR, Alatawi NHO, Raja R, Zhang J, et al. PAK3 Exacerbates Cardiac Lipotoxicity via SREBP1c in Obesity Cardiomyopathy. Diabetes. 2024;73(11):1805–1820. doi:10.2337/db24-0240

900. Zhang J, Zhang Y, Sun T, Guo F, Huang S, Chandalia M, Abate N, Fan D, Xin HB, Chen YE, et al. Dietary obesity-induced Egr-1 in adipocytes facilitates energy storage via suppression of FOXC2. Sci Rep. 2013;3:1476. doi:10.1038/srep01476

901. Zafar A, Ng HP, Chan ER, Dunwoodie SL, Mahabeleshwar GH. Myeloid-CITED2 Deficiency Exacerbates Diet-Induced Obesity and Pro-Inflammatory Macrophage Response. Cells. 2023;12(17):2136. doi:10.3390/cells12172136

902. Shintani T, Suzuki R, Takeuchi Y, Shirasawa T, Noda M. Deletion or inhibition of PTPRO prevents ectopic fat accumulation and induces healthy obesity with markedly reduced systemic inflammation. Life Sci. 2023;313:121292. doi:10.1016/j.lfs.2022.121292

903. Zhang T, Linghu KG, Tan J, Wang M, Chen D, Shen Y, Wu J, Shi M, Zhou Y, Tang L, Liu L, et al. TIGAR exacerbates obesity by triggering LRRK2-mediated defects in macroautophagy and chaperone-mediated autophagy in adipocytes. Autophagy. 2024;20(8):1741–1761. doi:10.1080/15548627.2024.2338576

904. Bai N, Lu X, Jin L, Alimujiang M, Ma J, Hu F, Xu Y, Sun J, Xu J, Zhang R, et al. CLSTN3 gene variant associates with obesity risk and contributes to dysfunction in white adipose tissue. Mol Metab. 2022;63:101531. doi:10.1016/j.molmet.2022.101531

905. Kumar U, Singh S. Role of Somatostatin in the Regulation of Central and Peripheral Factors of Satiety and Obesity. Int J Mol Sci. 2020;21(7):2568. doi:10.3390/ijms21072568

906. Prakash J, Mittal B, Awasthi S, Srivastava N. Association of the -243A>G, +61450C>A Polymorphisms of the Glutamate Decarboxylase 2 (GAD2) Gene with Obesity and Insulin Level in North Indian Population. Iran J Public Health. 2016;45(4):460–468.

907. Mazzarella L, Botteri E, Matthews A, Gatti E, Di Salvatore D, Bagnardi V, Breccia M, Montesinos P, Bernal T, Gil C, et al. Obesity is a risk factor for acute promyelocytic leukemia: evidence from population and cross-sectional studies and correlation with FLT3 mutations and polyunsaturated fatty acid metabolism. Haematologica. 2020;105(6):1559–1566. doi:10.3324/haematol.2019.223925

908. Costantino S, Paneni F, Virdis A, Hussain S, Mohammed SA, Capretti G, Akhmedov A, Dalgaard K, Chiandotto S, Pospisilik JA, et al. Interplay among H3K9-editing enzymes SUV39H1, JMJD2C and SRC-1 drives p66Shc transcription and vascular oxidative stress in obesity. Eur Heart J. 2019;40(4):383–391. doi:10.1093/eurheartj/ehx615

909. Nawrocki AR, Rodriguez CG, Toolan DM, Price O, Henry M, Forrest G, Szeto D, Keohane CA, Pan Y, Smith KM, et al. Genetic deletion and pharmacological inhibition of phosphodiesterase 10A protects mice from diet-induced obesity and insulin resistance. Diabetes. 2014;63(1):300–311. doi:10.2337/db13-0247

910. Aliasghari F, Nazm SA, Yasari S, Mahdavi R, Bonyadi M. Associations of the ANKK1 and DRD2 gene polymorphisms with overweight, obesity and hedonic hunger among women from the Northwest of Iran. Eat Weight Disord. 2021;26(1):305–312. doi:10.1007/s40519-020-00851-5

911. Ma WW, Ding BJ, Yuan LH, Zhao L, Yu HL, Xi YD, Xiao R. Neurocalcin-delta: a potential memory-related factor in hippocampus of obese rats induced by high-fat diet. Afr Health Sci. 2017;17(4):1211–1221. doi:10.4314/ahs.v17i4.32

912. Moreno-Navarrete JM, Moreno M, Ortega F, Sabater M, Xifra G, Ricart W, Fernández-Real JM. CISD1 in association with obesity-associated dysfunctional adipogenesis in human visceral adipose tissue. Obesity (Silver Spring). 2016;24(1):139–147. doi:10.1002/oby.21334

913. Shimada Y, Kuninaga S, Ariyoshi M, Zhang B, Shiina Y, Takahashi Y, Umemoto N, Nishimura Y, Enari H, Tanaka T. E2F8 promotes hepatic steatosis through FABP3 expression in diet-induced obesity in zebrafish. Nutr Metab (Lond). 2015;12:17. doi:10.1186/s12986-015-0012-7

914. Khalilian S, Hojati Z, Dehghanian F, Shaygannejad V, Imani SZH, Kheirollahi M, Khorrami M, Mirmosayyeb O. Gene expression profiles of YAP1, TAZ, CRB3, and VDR in familial and sporadic multiple sclerosis among an Iranian population. Sci Rep. 2021;11(1):7713. doi:10.1038/s41598-021-87131-z

915. Zarobkiewicz MK, Morawska I, Kowalska W, Halczuk P, Roliński J, Bojarska-Junak AA. PECAM-1 Is Down-Regulated in γδT Cells during Remission, but Up-Regulated in Relapse of Multiple Sclerosis. J Clin Med. 2022;11(11):3210. doi:10.3390/jcm11113210

916. Haupeltshofer S, Leichsenring T, Berg S, Pedreiturria X, Joachim SC, Tischoff I, Otte JM, Bopp T, Fantini MC, Esser C, et al. Smad7 in intestinal CD4+ T cells determines autoimmunity in a spontaneous model of multiple sclerosis. Proc Natl Acad Sci U S A. 2019;116(51):25860–25869. doi:10.1073/pnas.1905955116

917. Zare-Chahoki A, Ahmadi-Zeidabadi M, Azadarmaki S, Ghorbani S, Noorbakhsh F. Inflammation in an Animal Model of Multiple Sclerosis Leads to MicroRNA-25-3p Dysregulation Associated with Inhibition of Pten and Klf4. Iran J Allergy Asthma Immunol. 2021;20(3):314–325 doi:10.18502/ijaai.v20i3.6337

918. Azimi G, Ranjbaran F, Arsang-Jang S, Ghafouri-Fard S, Mazdeh M, Sayad A, Taheri M. Upregulation of VEGF-A and correlation between VEGF-A and FLT-1 expressions in Iranian multiple sclerosis patients. Neurol Sci. 2020;41(6):1459–1465. doi:10.1007/s10072-019-04234-2

919. Tezuka K, Suzuki M, Sato R, Kawarada S, Terasaki T, Uchida Y. Activation of Annexin A2 signaling at the blood-brain barrier in a mouse model of multiple sclerosis. J Neurochem. 2022;160(6):662–674. doi:10.1111/jnc.15578

920. Luo F, Tran AP, Xin L, Sanapala C, Lang BT, Silver J, Yang Y. Modulation of proteoglycan receptor PTPσ enhances MMP-2 activity to promote recovery from multiple sclerosis. Nat Commun. 2018;9(1):4126. doi:10.1038/s41467-018-06505-6

921. Shaker OG, Golam RM, Ayoub S, Daker LI, Elguaad MKA, Said ES, Khalil MAF. Correlation between LincR-Gng2-5’and LincR-Epas1-3’as with the severity of multiple sclerosis in Egyptian patients. Int J Neurosci. 2020;130(5):515–521. doi:10.1080/00207454.2019.1695610

922. Goris A, Williams-Gray CH, Foltynie T, Brown J, Maranian M, Walton A, Compston DA, Barker RA, Sawcer SJ. Investigation of TGFB2 as a candidate gene in multiple sclerosis and Parkinson’s disease. J Neurol. 2007;254(7):846–848. doi:10.1007/s00415-006-0414-6

923. Aliomrani M, Sahraian MA, Shirkhanloo H, Sharifzadeh M, Khoshayand MR, Ghahremani MH. Correlation between heavy metal exposure and GSTM1 polymorphism in Iranian multiple sclerosis patients. Neurol Sci. 2017;38(7):1271–1278. doi:10.1007/s10072-017-2934-5

924. Jaime-Pérez JC, Turrubiates-Hernández GA, López-Silva LJ, Salazar-Riojas R, Gómez-Almaguer D. Early changes in IL-21, IL-22, CCL2, and CCL4 serum cytokines after outpatient autologous transplantation for multiple sclerosis: A proof of concept study. Clin Transplant. 2020;34(12):e14114. doi:10.1111/ctr.14114

925. Abdelalim LR, Elnaggar YSR, Abdallah OY. Lactoferrin, chitosan double-coated oleosomes loaded with clobetasol propionate for remyelination in multiple sclerosis: Physicochemical characterization and in-vivo assessment in a cuprizone-induced demyelination model. Int J Biol Macromol. 2024;277(Pt 1):134144. doi:10.1016/j.ijbiomac.2024.134144

926. Čierny D, Lehotský J, Kantorová E, Sivák Š, Javor J, Kurča E, Dobrota D, Michalik J. The HLA-DRB1 and HLA-DQB1 alleles are associated with multiple sclerosis disability progression in Slovak population. Neurol Res. 2018;40(7):607–614. doi:10.1080/01616412.2018.1456711

927. Luo H, Broux B, Wang X, Hu Y, Ghannam S, Jin W, Larochelle C, Prat A, Wu J. EphrinB1 and EphrinB2 regulate T cell chemotaxis and migration in experimental autoimmune encephalomyelitis and multiple sclerosis. Neurobiol Dis. 2016;91:292–306. doi:10.1016/j.nbd.2016.03.013

928. Moll NM, Hong E, Fauveau M, Naruse M, Kerninon C, Tepavcevic V, Klopstein A, Seilhean D, Chew LJ, Gallo V, et al. SOX17 is expressed in regenerating oligodendrocytes in experimental models of demyelination and in multiple sclerosis. Glia. 2013;61(10):1659–1672. doi:10.1002/glia.22547

929. Lu M, Shi H, Taylor BV, Körner H. Alterations of subset and cytokine profile of peripheral T helper cells in PBMCs from Multiple Sclerosis patients or from individuals with MS risk SNPs near genes CYP27B1 and CYP24A1. Cytokine. 2022;153:155866. doi:10.1016/j.cyto.2022.155866

930. Avasarala JR, Jones JR, Rogers CR. Forkhead box C1 gene variant causing glaucoma and small vessel angiopathy can mimic multiple sclerosis. Mult Scler Relat Disord. 2018;22:157–160. doi:10.1016/j.msard.2018.04.004931.

931. Thöne J, Kleiter I, Stahl A, Ellrichmann G, Gold R, Hellwig K. Relevance of endoglin, IL-1α, IL-1β and anti-ovarian antibodies in females with multiple sclerosis. J Neurol Sci. 2016;362:240–243. doi:10.1016/j.jns.2016.01.057

932. Granström AL, Markljung E, Fink K, Nordenskjöld E, Nilsson D, Wester T, Nordenskjöld A. A novel stop mutation in the EDNRB gene in a family with Hirschsprung’s disease associated with multiple sclerosis. J Pediatr Surg. 2014;49(4):622–625. doi:10.1016/j.jpedsurg.2013.10.027

933. Rot U, Sandelius Å, Emeršič A, Zetterberg H, Blennow K. Cerebrospinal fluid GAP-43 in early multiple sclerosis. Mult Scler J Exp Transl Clin. 2018;4(3):2055217318792931. doi:10.1177/2055217318792931

934. Cunningham S, O’Doherty C, Patterson C, McDonnell G, Hawkins S, Marrosu MG, Vandenbroeck K. The neuropeptide genes TAC1, TAC3, TAC4, VIP and PACAP(ADCYAP1), and susceptibility to multiple sclerosis. J Neuroimmunol. 2007;183(1-2):208–213. doi:10.1016/j.jneuroim.2006.11.002

935. Thümmler K, Wrzos C, Franz J, McElroy D, Cole JJ, Hayden L, Arseni D, Schwarz F, Junker A, Edgar JM, et al. Fibroblast growth factor 9 (FGF9)-mediated neurodegeneration: Implications for progressive multiple sclerosis?. Neuropathol Appl Neurobiol. 2023;49(5):e12935. doi:10.1111/nan.12935

936. Brugger SW, Gardner MC, Beales JT, Briggs F, Davis MF. Depression in multiple sclerosis patients associated with risk variant near NEGR1. Mult Scler Relat Disord. 2020;46:102537. doi:10.1016/j.msard.2020.102537

937. Graves JS, Barcellos LF, Simpson S, Belman A, Lin R, Taylor BV, Ponsonby AL, Dwyer T, Krupp L, Waubant E, et al. The multiple sclerosis risk allele within the AHI1 gene is associated with relapses in children and adults. Mult Scler Relat Disord. 2018;19:161–165. doi:10.1016/j.msard.2017.10.008

938. ŞİmŞek F, ÖztÜrk N. DCLK-1 level in multiple sclerosis patients and its correlation with clinic. Mult Scler Relat Disord. 2020;43:102179. doi:10.1016/j.msard.2020.102179

939. Eiza N, Garty M, Staun-Ram E, Miller A, Vadasz Z. The possible involvement of sema3A and sema4A in the pathogenesis of multiple sclerosis. Clin Immunol. 2022;238:109017. doi:10.1016/j.clim.2022.109017

940. Duan J, Lv A, Guo Z, Liu Q, Tian C, Yang Y, Bi J, Yu X, Peng G, Luo B, et al. CX3CR1+/UCHL1+ microglial extracellular vesicles in blood: a potential biomarker for multiple sclerosis. J Neuroinflammation. 2024;21(1):254. doi:10.1186/s12974-024-03243-z

941. Shafit-Zagardo B, Kress Y, Zhao ML, Lee SC. A novel microtubule-associated protein-2 expressed in oligodendrocytes in multiple sclerosis lesions. J Neurochem. 1999;73(6):2531–2537. doi:10.1046/j.1471-4159.1999.0732531.x

942. Benítez-Fernández R, Josa-Prado F, Sánchez E, Lao Y, García-Rubia A, Cumella J, Martínez A, Palomo V, de Castro F. Efficacy of a benzothiazole-based LRRK2 inhibitor in oligodendrocyte precursor cells and in a murine model of multiple sclerosis. CNS Neurosci Ther. 2024;30(1):e14552. doi:10.1111/cns.14552

943. König T, Leutmezer F, Berger T, Zimprich A, Schmied C, Stögmann E, Zrzavy T. No Association of Multiple Sclerosis with C9orf72 Hexanucleotide Repeat Size in an Austrian Cohort. Int J Mol Sci. 2023;24(14):11254. doi:10.3390/ijms241411254

944. Basivireddy J, Somvanshi RK, Romero IA, Weksler BB, Couraud PO, Oger J, Kumar U. Somatostatin preserved blood brain barrier against cytokine induced alterations: possible role in multiple sclerosis. Biochem Pharmacol. 2013;86(4):497–507. doi:10.1016/j.bcp.2013.06.001

945. Ziccardi S, Tamanti A, Ruggieri C, Guandalini M, Marastoni D, Camera V, Montibeller L, Mazziotti V, Rossi S, Calderone M, Pizzini FB, et al. CSF Parvalbumin Levels at Multiple Sclerosis Diagnosis Predict Future Worse Cognition, Physical Disability, Fatigue, and Gray Matter Damage. Neurol Neuroimmunol Neuroinflamm. 2024;11(6):e200301. doi:10.1212/NXI.0000000000200301

946. Cunningham S, O’Doherty C, Patterson C, McDonnell G, Hawkins S, Marrosu MG, Vandenbroeck K. The neuropeptide genes TAC1, TAC3, TAC4, VIP and PACAP(ADCYAP1), and susceptibility to multiple sclerosis. J Neuroimmunol. 2007;183(1-2):208-213. doi:10.1016/j.jneuroim.2006.11.002

947. Bridel C, Koel-Simmelink MJA, Peferoen L, Derada Troletti C, Durieux S, Gorter R, Nutma E, Gami P, Iacobaeus E, Brundin L, et al. Brain endothelial cell expression of SPARCL-1 is specific to chronic multiple sclerosis lesions and is regulated by inflammatory mediators in vitro. Neuropathol Appl Neurobiol. 2018;44(4):404–416. doi:10.1111/nan.12412

948. Davies JL, Thompson S, Kaur-Sandhu H, Sawcer S, Coles A, Ban M, Jones J. Increased THEMIS First Exon Usage in CD4+ T-Cells Is Associated with a Genotype that Is Protective against Multiple Sclerosis. PLoS One. 2016;11(7):e0158327. doi:10.1371/journal.pone.0158327

949. Jazireian P, Sasani ST, Assarzadegan F, Azimian M. TRAILR1 (rs20576) and GRIA3 (rs12557782) are not associated with interferon-β response in multiple sclerosis patients. Mol Biol Rep. 2020;47(12):9659–9665. doi:10.1007/s11033-020-06026-w

950. Otaegui D, Zuriarrain O, Castillo-Triviño T, Aransay A, Ruíz-Martinez J, Olaskoaga J, Marti-Masso J, Lopez de Munain A. Association between synapsin III gene promoter SNPs and multiple sclerosis in Basque patients. Mult Scler. 2009;15(1):124–128. doi:10.1177/1352458508096682

951. Hughes AJ, Dunn KM, Chaffee T, Bhattarai JJ, Beier M. Diagnostic and Clinical Utility of the GAD-2 for Screening Anxiety Symptoms in Individuals With Multiple Sclerosis. Arch Phys Med Rehabil. 2018;99(10):2045–2049. doi:10.1016/j.apmr.2018.05.029

952. Wu Q, Du X, Cheng J, Qi X, Liu H, Lv X, Gong X, Shao C, Wang M, Yue L, et al. PECAM-1 drives β-catenin-mediated EndMT via internalization in colon cancer with diabetes mellitus. Cell Commun Signal. 2023;21(1):203. doi:10.1186/s12964-023-01193-2

953. Ferjeni Z, Bouzid D, Fourati H, Stayoussef M, Abida O, Kammoun T, Hachicha M, Penha-Gonçalves C, Masmoudi H. Association of TCR/CD3, PTPN22, CD28 and ZAP70 gene polymorphisms with type 1 diabetes risk in Tunisian population: family based association study. Immunol Lett. 2015;163(1):1–7. doi:10.1016/j.imlet.2014.11.005

954. Wang Y, Zhang R, Shen H, Kong J, Lv X. Pioglitazone protects blood vessels through inhibition of the apelin signaling pathway by promoting KLF4 expression in rat models of T2DM. Biosci Rep. 2019;39(12):BSR20190317. doi:10.1042/BSR20190317

955. Tokuda H, Kuroyanagi G, Tsujimoto M, Enomoto Y, Matsushima-Nishiwaki R, Onuma T, Kojima A, Doi T, Tanabe K, Akamatsu S, et al. Release of Phosphorylated HSP27 (HSPB1) from Platelets Is Accompanied with the Acceleration of Aggregation in Diabetic Patients. PLoS One. 2015;10(6):e0128977. doi:10.1371/journal.pone.0128977

956. Bayat M, Chien S, Chehelcheraghi F. Co-localization of Flt1 and tryptase of mast cells in skin wound of rats with type I diabetes: Initial studies. Acta Histochem. 2021;123(2):151680. doi:10.1016/j.acthis.2021.151680

957. Kosoy R, Concannon P. Functional variants in SUMO4, TAB2, and NFkappaB and the risk of type 1 diabetes. Genes Immun. 2005;6(3):231–235. doi:10.1038/sj.gene.6364174

958. Chuang PY, Dai Y, Liu R, He H, Kretzler M, Jim B, Cohen CD, He JC.Alteration of forkhead box O (foxo4) acetylation mediates apoptosis of podocytes in diabetes mellitus. PLoS One. 2011;6(8):e23566. doi:10.1371/journal.pone.0023566

959. Liu J, Cheng Q, Wu X, Zhu H, Deng X, Wang M, Yang S, Xu J, Chen Q, Li M, et al. Icariin Treatment Rescues Diabetes Induced Bone Loss via Scavenging ROS and Activating Primary Cilia/Gli2/Osteocalcin Signaling Pathway. Cells. 2022;11(24):4091. doi:10.3390/cells11244091

960. Wang XL, Greco M, Sim AS, Duarte N, Wang J, Wilcken DE. Effect of CYP1A1 MspI polymorphism on cigarette smoking related coronary artery disease and diabetes. Atherosclerosis. 2002;162(2):391–397. doi:10.1016/s0021-9150(01)00723-7

961. Gajewska B, Śliwińska-Mossoń M. Association of MMP-2 and MMP-9 Polymorphisms with Diabetes and Pathogenesis of Diabetic Complications. Int J Mol Sci. 2022;23(18):10571. doi:10.3390/ijms231810571

962. Li M, Huang S, Zhang Y, Song Z, Fu H, Lin Z, Huang X. Regulation of the unfolded protein response transducer IRE1α by SERPINH1 aggravates periodontitis with diabetes mellitus via prolonged ER stress. Cell Signal. 2022;91:110241. doi:10.1016/j.cellsig.2022.110241

963. Gao C, Lin X, Fan F, Liu X, Wan H, Yuan T, Zhao X, Luo Y. Status of higher TGF-β1 and TGF-β2 levels in the aqueous humour of patients with diabetes and cataracts. BMC Ophthalmol. 2022;22(1):156. doi:10.1186/s12886-022-02317-x

964. Yuan T, Annamalai K, Naik S, Lupse B, Geravandi S, Pal A, Dobrowolski A, Ghawali J, Ruhlandt M, Gorrepati KDD, et al. The Hippo kinase LATS2 impairs pancreatic β-cell survival in diabetes through the mTORC1-autophagy axis. Nat Commun. 2021;12(1):4928. doi:10.1038/s41467-021-25145-x

965. Stoian A, Bănescu C, Bălaşa RI, Moţăţăianu A, Stoian M, Moldovan VG, Voidăzan S, Dobreanu M. Influence of GSTM1, GSTT1, and GSTP1 Polymorphisms on Type 2 Diabetes Mellitus and Diabetic Sensorimotor Peripheral Neuropathy Risk. Dis Markers. 2015;2015:638693. doi:10.1155/2015/638693

966. Jin Q, Lin L, Zhao T, Yao X, Teng Y, Zhang D, Jin Y, Yang M. Overexpression of E3 ubiquitin ligase Cbl attenuates endothelial dysfunction in diabetes mellitus by inhibiting the JAK2/STAT4 signaling and Runx3-mediated H3K4me3. J Transl Med. 2021;19(1):469. doi:10.1186/s12967-021-03069-w

967. Kerns SL, Guevara-Aguirre J, Andrew S, Geng J, Guevara C, Guevara-Aguirre M, Guo M, Oddoux C, Shen Y, Zurita A, et al. A novel variant in CDKN1C is associated with intrauterine growth restriction, short stature, and early-adulthood-onset diabetes. J Clin Endocrinol Metab. 2014;99(10):E2117–E2122. doi:10.1210/jc.2014-1949

968. Mir MM, Alfaifi J, Sohail SK, Rizvi SF, Akhtar MT, Alghamdi MAA, Mir R, Wani JI, Sabah ZU, Alhumaydhi FA, et al. The Role of Pro-Inflammatory Chemokines CCL-1, 2, 4, and 5 in the Etiopathogenesis of Type 2 Diabetes Mellitus in Subjects from the Asir Region of Saudi Arabia: Correlation with Different Degrees of Obesity. J Pers Med. 2024;14(7):743. doi:10.3390/jpm14070743

969. Mohamed WA, Schaalan MF. Antidiabetic efficacy of lactoferrin in type 2 diabetic pediatrics; controlling impact on PPAR-γ, SIRT-1, and TLR4 downstream signaling pathway. Diabetol Metab Syndr. 2018;10:89. doi:10.1186/s13098-018-0390-x

970. Wang X, Zhu L, Liu J, Ma Y, Qiu C, Liu C, Gong Y, Yuwen Y, Guan G, Zhang Y, et al. Palmitic acid in type 2 diabetes mellitus promotes atherosclerotic plaque vulnerability via macrophage Dll4 signaling. Nat Commun. 2024;15(1):1281. doi:10.1038/s41467-024-45582-8

971. Guo Q, Wang W, Abboud R, Guo Z. Impairment of maturation of BMP-6 (35 kDa) correlates with delayed fracture healing in experimental diabetes. J Orthop Surg Res. 2020;15(1):186. doi:10.1186/s13018-020-01705-7

972. Preeti K, Fernandes V, Sood A, Khan I, Khatri DK, Singh SB. Necrostatin-1S mitigates type-2 diabetes-associated cognitive decrement and lipotoxicity-induced neuro-microglia changes through p-RIPK-RIPK3-p-MLKL axis. Metab Brain Dis. 2023;38(5):1581–1612. doi:10.1007/s11011-023-01185-8

973. Arner P, Petrus P, Esteve D, Boulomié A, Näslund E, Thorell A, Gao H, Dahlman I, Rydén M. Screening of potential adipokines identifies S100A4 as a marker of pernicious adipose tissue and insulin resistance. Int J Obes (Lond). 2018;42(12):2047–2056. doi:10.1038/s41366-018-0018-0

974. Hertle E, Arts IC, van der Kallen CJ, Feskens EJ, Schalkwijk CG, Hoffmann-Petersen IT, Thiel S, Stehouwer CD, van Greevenbroek MM. Distinct Longitudinal Associations of MBL, MASP-1, MASP-2, MASP-3, and MAp44 With Endothelial Dysfunction and Intima-Media Thickness: The Cohort on Diabetes and Atherosclerosis Maastricht (CODAM) Study. Arterioscler Thromb Vasc Biol. 2016;36(6):1278–1285. doi:10.1161/ATVBAHA.115.306552

975. Charles BA, Conley YP, Chen G, Miller RG, Dorman JS, Gorin MB, Ferrell RE, Sereika SM, Rotimi CN, Orchard TJ. Variants of the adenosine A(2A) receptor gene are protective against proliferative diabetic retinopathy in patients with type 1 diabetes. Ophthalmic Res. 2011;46(1):1–8. doi:10.1159/000317057

976. Hussein AG, Mohamed RH, Alghobashy AA. Synergism of CYP2R1 and CYP27B1 polymorphisms and susceptibility to type 1 diabetes in Egyptian children. Cell Immunol. 2012;279(1):42–45. doi:10.1016/j.cellimm.2012.08.006

977. Wu J, Wang X, Chen H, Yang R, Yu H, Wu Y, Hu Y. Type 2 Diabetes Risk and Lipid Metabolism Related to the Pleiotropic Effects of an ABCB1 Variant: A Chinese Family-Based Cohort Study. Metabolites. 2022;12(9):875. doi:10.3390/metabo12090875

978. Ghodsian N, Ismail P, Ahmadloo S, Heidari F, Haghvirdizadeh P, Ataollahi Eshkoor S, Etemad A. Novel Association of WNK4 Gene, Ala589Ser Polymorphism in Essential Hypertension, and Type 2 Diabetes Mellitus in Malaysia. J Diabetes Res. 2016;2016:8219543. doi:10.1155/2016/8219543

979. Chen K, Wu Q, Hu K, Yang C, Wu X, Cheung P, Williams KJ. Suppression of Hepatic FLOT1 (Flotillin-1) by Type 2 Diabetes Mellitus Impairs the Disposal of Remnant Lipoproteins via Syndecan-1. Arterioscler Thromb Vasc Biol. 2018;38(1):102–113. doi:10.1161/ATVBAHA.117.310358

980. Dikilitaş A, Karaaslan F, Evirgen Ş, Ertuğrul AS. Gingival crevicular fluid CSF-1 and IL-34 levels in patients with stage III grade C periodontitis and uncontrolled type 2 diabetes mellitus. J Periodontal Implant Sci. 2022;52(6):455–465. doi:10.5051/jpis.2106260313

981. Blázquez-Medela AM, García-Ortiz L, Gómez-Marcos MA, Recio-Rodríguez JI, Sánchez-Rodríguez A, López-Novoa JM, Martínez-Salgado C. Increased plasma soluble endoglin levels as an indicator of cardiovascular alterations in hypertensive and diabetic patients. BMC Med. 2010;8:86. doi:10.1186/1741-7015-8-86

982. Yin Y, Li L, Yu S, Xin Y, Zhu L, Hu X, Chen K, Gu W, Mu Y, Zang L, et al. The first compound heterozygous mutations in SLC12A3 and PDX1 genes: a unique presentation of Gitelman syndrome with distinct insulin resistance and familial diabetes insights. Front Endocrinol (Lausanne). 2024;14:1327729. doi:10.3389/fendo.2023.1327729

983. Gautheron J, Morisseau C, Chung WK, Zammouri J, Auclair M, Baujat G, Capel E, Moulin C, Wang Y, Yang J, et al. EPHX1 mutations cause a lipoatrophic diabetes syndrome due to impaired epoxide hydrolysis and increased cellular senescence. Elife. 2021;10:e68445. doi:10.7554/eLife.68445

984. Zhang S, Zang W. The Role and Mechanism of STIM1/Orai1-Regulated Ca2+ Influx in Myocardial Hypertrophy in Type 2 Diabetes Mellitus. Ann Clin Lab Sci. 2024;54(1):17–25.

985. Li Y, Ma WG, Li XC. Identification of Blood miR-216a, miR-377 and Their Target Genes ANGPTL4, GAP-43 and Serum of PPARG as Biomarkers for Diabetic Peripheral Neuropathy of Type 2 Diabetes. Clin Lab. 2021;67(4):https://10.7754/Clin.Lab.2020.191220. doi:10.7754/Clin.Lab.2020.191220

986. Zhang XL, Zhao X, Wu Y, Huang WQ, Chen JJ, Hu P, Liu W, Chen YW, Hao J, Xie RR, et al. Angiotensin(1-7) activates MAS-1 and upregulates CFTR to promote insulin secretion in pancreatic β-cells: the association with type 2 diabetes. Endocr Connect. 2022;11(1):e210357.doi:10.1530/EC-21-0357

987. Aparicio JM, Wakisaka A, Takada A, Matsuura N, Yoshiki T. Non-HLA genetic factors and insulin dependent diabetes mellitus in the Japanese: TCRA, TCRB and TCRG, INS, THY1, CD3D and ETS1. Dis Markers. 1990;8(5):283–294.

988. Demir I, Yilmaz I, Horoz E, Bozkaya G, Bilgir O. The relationship between stathmin-2 level and metabolic parameters in newly diagnosed type 2 diabetes mellitus patients. Am J Med Sci. 2024;368(1):25–32. doi:10.1016/j.amjms.2024.03.023

989. Zhang X, Lv H, Mei J, Ji B, Huang S, Li X. The Potential Role of R4 Regulators of G Protein Signaling (RGS) Proteins in Type 2 Diabetes Mellitus. Cells. 2022;11(23):3897. doi:10.3390/cells11233897

990. Gu HF. Genetic variation screening and association studies of the adenylate cyclase activating polypeptide 1 (ADCYAP1) gene in patients with type 2 diabetes. Hum Mutat. 2002;19(5):572–573. doi:10.1002/humu.9034

991. Wallner C, Schira J, Wagner JM, Schulte M, Fischer S, Hirsch T, Richter W, Abraham S, Kneser U, Lehnhardt M, et al. Application of VEGFA and FGF-9 enhances angiogenesis, osteogenesis and bone remodeling in type 2 diabetic long bone regeneration. PLoS One. 2015;10(3):e0118823. doi:10.1371/journal.pone.0118823

992. Ng MC, Tam CH, So WY, Ho JS, Chan AW, Lee HM, Wang Y, Lam VK, Chan JC, Ma RC. Implication of genetic variants near NEGR1, SEC16B, TMEM18, ETV5/DGKG, GNPDA2, LIN7C/BDNF, MTCH2, BCDIN3D/FAIM2, SH2B1, FTO, MC4R, and KCTD15 with obesity and type 2 diabetes in 7705 Chinese. J Clin Endocrinol Metab. 2010;95(5):2418–2425. doi:10.1210/jc.2009-2077

993. Al-Daghri NM, Costa AS, Alokail MS, Zanzottera M, Alenad AM, Mohammed AK, Clerici M, Guerini FR. Synaptosomal Protein of 25 kDa (Snap25) Polymorphisms Associated with Glycemic Parameters in Type 2 Diabetes Patients. J Diabetes Res. 2016;2016:8943092. doi:10.1155/2016/8943092

994. Alessi MC, Nicaud V, Scroyen I, Lange C, Saut N, Fumeron F, Marre M, Lantieri O, Fontaine-Bisson B, Juhan-Vague I, et al. Association of vitronectin and plasminogen activator inhibitor-1 levels with the risk of metabolic syndrome and type 2 diabetes mellitus. Results from the D.E.S.I.R. prospective cohort. Thromb Haemost. 2011;106(3):416–422. doi:10.1160/TH11-03-0179

995. Razeghi P, Young ME, Cockrill TC, Frazier OH, Taegtmeyer H. Downregulation of myocardial myocyte enhancer factor 2C and myocyte enhancer factor 2C-regulated gene expression in diabetic patients with nonischemic heart failure. Circulation. 2002;106(4):407–411. doi:10.1161/01.cir.0000026392.80723.dc

996. Veluthakal R, Chepurny OG, Leech CA, Schwede F, Holz GG, Thurmond DC. Restoration of Glucose-Stimulated Cdc42-Pak1 Activation and Insulin Secretion by a Selective Epac Activator in Type 2 Diabetic Human Islets. Diabetes. 2018;67(10):1999–2011. doi:10.2337/db17-1174

997. Holmkvist J, Anthonsen S, Wegner L, Andersen G, Jørgensen T, Borch-Johnsen K, Sandbaek A, Lauritzen T, Pedersen O, Hansen T. Polymorphisms in AHI1 are not associated with type 2 diabetes or related phenotypes in Danes: non-replication of a genome-wide association result. Diabetologia. 2008;51(4):609–614. doi:10.1007/s00125-008-0925-z

998. Ji L, Yang X, Jin Y, Li L, Yang B, Zhu W, Xu M, Wang Y, Wu G, Luo W, et al. Blockage of DCLK1 in cardiomyocytes suppresses myocardial inflammation and alleviates diabetic cardiomyopathy in streptozotocin-induced diabetic mice. Biochim Biophys Acta Mol Basis Dis. 2024;1870(1):166900. doi:10.1016/j.bbadis.2023.166900

999. Lin AC, Hung HC, Chen YW, Cheng KP, Li CH, Lin CH, Chang CJ, Wu HT, Ou HY. Elevated Hedgehog-Interacting Protein Levels in Subjects with Prediabetes and Type 2 Diabetes. J Clin Med. 2019;8(10):1635. doi:10.3390/jcm8101635

1000. Li P, Qin D, Chen T, Hou W, Song X, Yin S, Song M, Fernando WCHA, Chen X, Sun Y, et al. Dysregulated Rbfox2 produces aberrant splicing of CaV1.2 calcium channel in diabetes-induced cardiac hypertrophy. Cardiovasc Diabetol. 2023;22(1):168. doi:10.1186/s12933-023-01894-5

1001. Kang YE, Choung S, Lee JH, Kim HJ, Ku BJ. The Role of Circulating Slit2, the One of the Newly Batokines, in Human Diabetes Mellitus. Endocrinol Metab (Seoul). 2017;32(3):383–388. doi:10.3803/EnM.2017.32.3.383

1002. Qiao Q, Xu X, Song Y, Song S, Zhu W, Li F. Semaphorin 3A promotes osteogenic differentiation of BMSC from type 2 diabetes mellitus rats. J Mol Histol. 2018;49(4):369–376. doi:10.1007/s10735-018-9776-1

1003. Costes S, Huang CJ, Gurlo T, Daval M, Matveyenko AV, Rizza RA, Butler AE, Butler PC. β-cell dysfunctional ERAD/ubiquitin/proteasome system in type 2 diabetes mediated by islet amyloid polypeptide-induced UCH-L1 deficiency. Diabetes. 2011;60(1):227–238. doi:10.2337/db10-0522

1004. Ke RQ, Wang Y, Hong SH, Xiao LX. Anti-diabetic effect of quercetin in type 2 diabetes mellitus by regulating the microRNA-92b-3p/EGR1 axis. J Physiol Pharmacol. 2023;74(2):10.26402/jpp.2023.2.03. doi:10.26402/jpp.2023.2.03

1005. Jia G, Sowers JR. Targeting CITED2 for Angiogenesis in Obesity and Insulin Resistance. Diabetes. 2016;65(12):3535–3536. doi:10.2337/dbi16-0052

1006. Chang TJ, Chiu YF, Sheu WH, Shih KC, Hwu CM, Quertermous T, Jou YS, Kuo SS, Chang YC, Chuang LM. Genetic polymorphisms of PCSK2 are associated with glucose homeostasis and progression to type 2 diabetes in a Chinese population. Sci Rep. 2015;5:14380. doi:10.1038/srep14380

1007. Huang MH, Liu YF, Nfor ON, Hsu SY, Lin WY, Chang YS, Liaw YP. nteractive Association Between Intronic Polymorphism (rs10506151) of the LRRK2 Gene and Type 2 Diabetes on Neurodegenerative Diseases. Pharmgenomics Pers Med. 2021;14:839–847. doi:10.2147/PGPM.S316158

1008. Bansal V, Winkelmann BR, Dietrich JW, Boehm BO. Whole-exome sequencing in familial type 2 diabetes identifies an atypical missense variant in the RyR2 gene. Front Endocrinol (Lausanne). 2024;15:1258982. doi:10.3389/fendo.2024.1258982

1009. Bonner SM, Pietropaolo SL, Fan Y, Chang Y, Sethupathy P, Morran MP, Beems M, Giannoukakis N, Trucco G, Palumbo MO, et al. Sequence variation in promoter of Ica1 gene, which encodes protein implicated in type 1 diabetes, causes transcription factor autoimmune regulator (AIRE) to increase its binding and down-regulate expression. J Biol Chem. 2012;287(21):17882–17893. doi:10.1074/jbc.M111.319020

1010. Kwon YJ, Hong KW, Park BJ, Jung DH. Serotonin receptor 3B polymorphisms are associated with type 2 diabetes: The Korean Genome and Epidemiology Study. Diabetes Res Clin Pract. 2019;153:76–85. doi:10.1016/j.diabres.2019.05.032

1011. Li Q, Qiao ZR, Liu DB, Zeng JT, Zhang J, Bo Y, Zu HY, Hu Q, Wu X, Dong SS. Relationship between serum GAD-Ab and the genetic polymorphisms of GAD2 and type 2 diabetes mellitus. Genet Mol Res. 2015;14(2):3002–3009. doi:10.4238/2015.April.10.10

1012. Abosheasha MA, Zahran F, Bessa SS, El-Magd MA, Mohamed TM. Association between a novel G94A single nucleotide polymorphism in ATP1A1 gene and type 2 diabetes mellitus among Egyptian patients. J Res Med Sci. 2019;24:62. doi:10.4103/jrms.JRMS_975_18

1013. Díaz-García JD, Leyva-Leyva M, Sánchez-Aguillón F, de León-Bautista MP, Fuentes-Venegas A, Torres-Viloria A, Tenorio-Aguirre EK, Morales-Lázaro SL, Olivo-Díaz A, González-Ramírez R. Association Study of CACNA1D, KCNJ11, KCNQ1, and CACNA1E Single-Nucleotide Polymorphisms with Type 2 Diabetes Mellitus. Int J Mol Sci. 2024;25(17):9196. doi:10.3390/ijms25179196

1014. Goru SK, Kadakol A, Pandey A, Malek V, Sharma N, Gaikwad AB. Histone H2AK119 and H2BK120 mono-ubiquitination modulate SET7/9 and SUV39H1 in type 1 diabetes-induced renal fibrosis. Biochem J. 2016;473(21):3937–3949. doi:10.1042/BCJ20160595

1015. Wysocka-Mincewicz M, Groszek A, Ambrozkiewicz F, Paziewska A, Dąbrowska M, Rybak A, Konopka E, Ochocińska A, Żeber-Lubecka N, Karczmarski J, et al. Combination of HLA-DQ2/-DQ8 Haplotypes and a Single MSH5 Gene Variant in a Polish Population of Patients with Type 1 Diabetes as a First Line Screening for Celiac Disease?. J Clin Med. 2022;11(8):2223. doi:10.3390/jcm11082223

1016. Ramos-Lopez O, Mejia-Godoy R, Frías-Delgadillo KJ, Torres-Valadez R, Flores-García A, Sánchez-Enríquez S, Aguiar-García P, Martínez-López E, Zepeda-Carrillo EA. Interactions between DRD2/ANKK1 TaqIA Polymorphism and Dietary Factors Influence Plasma Triglyceride Concentrations in Diabetic Patients from Western Mexico: A Cross-sectional Study. Nutrients. 2019;11(12):2863. doi:10.3390/nu11122863

1017. Takei S, Nagashima S, Takei A, Yamamuro D, Wakabayashi T, Murakami A, Isoda M, Yamazaki H, Ebihara C, Takahashi M, et al. β-Cell-Specific Deletion of HMG-CoA (3-hydroxy-3-methylglutaryl-coenzyme A) Reductase Causes Overt Diabetes due to Reduction of β-Cell Mass and Impaired Insulin Secretion. Diabetes. 2020;69(11):2352–2363. doi:10.2337/db19-0996

1018. Liu F, Dong Y, Zhong F, Guo H, Dong P. CISD1 Is a Breast Cancer Prognostic Biomarker Associated with Diabetes Mellitus. Biomolecules. 2022;13(1):37. doi:10.3390/biom13010037

1019. Lu P, Bevan DR, Lewis SN, Hontecillas R, Bassaganya-Riera J. Molecular modeling of lanthionine synthetase component C-like protein 2: a potential target for the discovery of novel type 2 diabetes prophylactics and therapeutics. J Mol Model. 2011;17(3):543–553. doi:10.1007/s00894-010-0748-y

1020. Zbidi H, López JJ, Amor NB, Bartegi A, Salido GM, Rosado JA. Enhanced expression of STIM1/Orai1 and TRPC3 in platelets from patients with type 2 diabetes mellitus. Blood Cells Mol Dis. 2009;43(2):211–213. doi:10.1016/j.bcmd.2009.04.005.

1021. Rodríguez-Calvo R, Granado-Casas M, Pérez-Montes de Oca A, Julian MT, Domingo M, Codina P, Santiago-Vacas E, Cediel G, Julve J, Rossell J, et al. Fatty Acid Binding Proteins 3 and 4 Predict Both All-Cause and Cardiovascular Mortality in Subjects with Chronic Heart Failure and Type 2 Diabetes Mellitus. Antioxidants (Basel). 2023;12(3):645. doi:10.3390/antiox12030645

1022. Adeyemo AA, Zaghloul NA, Chen G, Doumatey AP, Leitch CC, Hostelley TL, Nesmith JE, Zhou J, Bentley AR, Shriner D, et al. ZRANB3 is an African-specific type 2 diabetes locus associated with beta-cell mass and insulin response. Nat Commun. 2019;10(1):3195. doi:10.1038/s41467-019-10967-7

1023. Zhao J, An K, Mao Z, Qu Y, Wang D, Li J, Min Z, Xue Z. CCL5 promotes LFA-1 expression in Th17 cells and induces LCK and ZAP70 activation in a mouse model of Parkinson’s disease. Front Aging Neurosci. 2023;15:1250685. doi:10.3389/fnagi.2023.1250685

1024. Tan JSY, Lee B, Lim J, Ma DR, Goh JX, Goh SY, Gulam MY, Koh SM, Lee WW, Feng L, et al. Parkinson’s Disease-Specific Autoantibodies against the Neuroprotective Co-Chaperone STIP1. Cells. 2022;11(10):1649. doi:10.3390/cells11101649

1025. Zamanian MY, Golmohammadi M, Amin RS, Bustani GS, Romero-Parra RM, Zabibah RS, Oz T, Jalil AT, Soltani A, Kujawska M. Therapeutic Targeting of Krüppel-Like Factor 4 and Its Pharmacological Potential in Parkinson’s Disease: a Comprehensive Review. Mol Neurobiol. 2024;61(6):3596–3606. doi:10.1007/s12035-023-03800-2

1026. Meng J, Fang J, Bao Y, Chen H, Hu X, Wang Z, Li M, Cheng Q, Dong Y, Yang X, et al. The biphasic role of Hspb1 on ferroptotic cell death in Parkinson’s disease. Theranostics. 2024;14(12):4643–4666. doi:10.7150/thno.98457

1027. Zhang B, Hu YB, Li G, Yu HX, Cui C, Han YY, Li HX, Li G. Itga5- PTEN signaling regulates striatal synaptic strength and motor coordination in Parkinson’s disease. Int J Biol Sci. 2024;20(9):3302–3316. doi:10.7150/ijbs.96116

1028. Kurzawski M, Białecka M, Sławek J, Kłodowska-Duda G, Droździk M. Association study of GATA-2 transcription factor gene (GATA2) polymorphism and Parkinson’s disease. Parkinsonism Relat Disord. 2010;16(4):284–287. doi:10.1016/j.parkreldis.2009.10.006

1029. Chan DK, Mellick GD, Buchanan DD, Hung WT, Ng PW, Woo J, Kay R. Lack of association between CYP1A1 polymorphism and Parkinson’s disease in a Chinese population. J Neural Transm (Vienna). 2002;109(1):35–39. doi:10.1007/s702-002-8234-8

1030. Ma C, Liu Y, Li S, Ma C, Huang J, Wen S, Yang S, Wang B. Microglial cGAS drives neuroinflammation in the MPTP mouse models of Parkinson’s disease. CNS Neurosci Ther. 2023;29(7):2018–2035. doi:10.1111/cns.14157

1031. Booth HDE, Wessely F, Connor-Robson N, Rinaldi F, Vowles J, Browne C, Evetts SG, Hu MT, Cowley SA, Webber C, et al. RNA sequencing reveals MMP2 and TGFB1 downregulation in LRRK2 G2019S Parkinson’s iPSC-derived astrocytes. Neurobiol Dis. 2019;129:56–66. doi:10.1016/j.nbd.2019.05.006

1032. Wang T, Wang B. Association between Glutathione S-transferase M1/Glutathione S-transferase T1 polymorphisms and Parkinson’s disease: a meta-analysis. J Neurol Sci. 2014;338(1-2):65–70. doi:10.1016/j.jns.2013.12.018

1033. Huang P, Wan Z, Qu S. Targeting the RUNX3-miR-186-3p-DAT- IGF1R axis as a therapeutic strategy in a Parkinson’s disease model. J Transl Med. 2024;22(1):719. doi:10.1186/s12967-024-05535-7

1034. Xiromerisiou G, Marogianni C, Lampropoulos IC, Dardiotis E, Speletas M, Ntavaroukas P, Androutsopoulou A, Kalala F, Grigoriadis N, Papoutsopoulou S. Peripheral Inflammatory Markers TNF-? and CCL2 Revisited: Association with Parkinson’s Disease Severity. Int J Mol Sci. 2022;24(1):264. doi:10.3390/ijms24010264

1035. Tang S, Wang A, Yan X, Chu L, Yang X, Song Y, Sun K, Yu X, Liu R, Wu Z, et al. rain-targeted intranasal delivery of dopamine with borneol and lactoferrin co-modified nanoparticles for treating Parkinson’s disease. Drug Deliv. 2019;26(1):700–707. doi:10.1080/10717544.2019.1636420

1036. Pandi S, Chinniah R, Sevak V, Ravi PM, Raju M, Vellaiappan NA, Karuppiah B. Association of HLA-DRB1, DQA1 and DQB1 alleles and haplotype in Parkinson’s disease from South India. Neurosci Lett. 2021;765:136296. doi:10.1016/j.neulet.2021.136296

1037. Deng H, Zhu SH, Le WD, Yang HR, Lv HW, Xu HB, Xie WJ, Jankovic J. Examination of the MSX1 gene in patients with Parkinson’s disease. Acta Neurol Scand. 2009;120(6):442–444. doi:10.1111/j.1600-0404.2009.01271.x

1038. Müller T. ABCB1: is there a role in the drug treatment of Parkinson’s disease?. Expert Opin Drug Metab Toxicol. 2018;14(2):127–129. doi:10.1080/17425255.2018.1416096

1039. Chang KH, Wu YR, Chen YC, Wu HC, Chen CM. Association between CSF1 and CSF1R Polymorphisms and Parkinson’s Disease in Taiwan. J Clin Med. 2019;8(10):1529. doi:10.3390/jcm8101529

1040. Ellioff KJ, Osting SMK, Lentine A, Welper AD, Burger C, Greenspan DS. Ablation of Mitochondrial RCC1-L Induces Nigral Dopaminergic Neurodegeneration and Parkinsonian-like Motor Symptoms. Preprint. bioRxiv. 2024;2023.12.01.567409. doi:10.1101/2023.12.01.567409

1041. Cocco C, Corda G, Lisci C, Noli B, Carta M, Brancia C, Manca E, Masala C, Marrosu F, Solla P, et al. VGF peptides as novel biomarkers in Parkinson’s disease. Cell Tissue Res. 2020;379(1):93–107. doi:10.1007/s00441-019-03128-1

1042. Saal KA, Galter D, Roeber S, Bähr M, Tönges L, Lingor P. Altered Expression of Growth Associated Protein-43 and Rho Kinase in Human Patients with Parkinson’s Disease. Brain Pathol. 2017;27(1):13–25. doi:10.1111/bpa.12346

1043. Torres ERS, Stanojlovic M, Zelikowsky M, Bonsberger J, Hean S, Mulligan C, Baldauf L, Fleming S, Masliah E, Chesselet MF, et al. Alpha-synuclein pathology, microgliosis, and parvalbumin neuron loss in the amygdala associated with enhanced fear in the Thy1-aSyn model of Parkinson’s disease. Neurobiol Dis. 2021;158:105478. doi:10.1016/j.nbd.2021.105478

1044. Kochmanski J, Kuhn NC, Bernstein AI. Parkinson’s disease-associated, sex-specific changes in DNA methylation at PARK7 (DJ-1), SLC17A6 (VGLUT2), PTPRN2 (IA-2?), and NR4A2 (NURR1) in cortical neurons. NPJ Parkinsons Dis. 2022;8(1):120. doi:10.1038/s41531-022-00355-2

1045. Shen Y, Cui X, Hu Y, Zhang Z, Zhang Z. LncRNA-MIAT regulates the growth of SHSY5Y cells by regulating the miR-34-5p-SYT1 axis and exerts a neuroprotective effect in a mouse model of Parkinson’s disease. Am J Transl Res. 2021;13(9):9993–10013.

1046. Patel S, Howard D, French L. A pH-eQTL Interaction at the RIT2- SYT4 Parkinson’s Disease Risk Locus in the Substantia Nigra. Front Aging Neurosci. 2021;13:690632. doi:10.3389/fnagi.2021.690632

1047. Shulskaya MV, Semenova EI, Rudenok MM, Partevian SA, Lukashevich MV, Karabanov AV, Fedotova EY, Illarioshkin SN, Slominsky PA, Shadrina MI, et al. Analysis of LRRN3, MEF2C, SLC22A, and P2RY12 Gene Expression in the Peripheral Blood of Patients in the Early Stages of Parkinson’s Disease. Biomedicines. 2024;12(7):1391. doi:10.3390/biomedicines12071391

1048. Yu J, Meng J, Qin Z, Yu Y, Liang Y, Wang Y, Min D. Dysbiosis of gut microbiota inhibits NMNAT2 to promote neurobehavioral deficits and oxidative stress response in the 6-OHDA-lesioned rat model of Parkinson’s disease. J Neuroinflammation. 2023;20(1):117. doi:10.1186/s12974-023-02782-1

1049. Bai X, Jin J, Li S, Wang H, Xie A. CUB and Sushi Multiple Domains (CSMD1) Gene Polymorphisms and Susceptibilities to Idiopathic Parkinson’s Disease in Northern Chinese Han Population: A Case-Control Study. Parkinsons Dis. 2021;2021:6661162. doi:10.1155/2021/6661162

1050. Wang J, Si YM, Liu ZL, Yu L. Cholecystokinin, cholecystokinin-A receptor and cholecystokinin-B receptor gene polymorphisms in Parkinson’s disease. Pharmacogenetics. 2003;13(6):365–369. doi:10.1097/00008571-200306000-00008

1051. Qi L, Tang YG, Wang L, He W, Pan HH, Nie RR, Can Y. ole of Rho- mediated ROCK-Semaphorin3A signaling pathway in the pathogenesis of Parkinson’s disease in a mouse model. J Neurol Sci. 2016;370:21–26. doi:10.1016/j.jns.2016.08.061

1052. Ham SJ, Lee D, Xu WJ, Cho E, Choi S, Min S, Park S, Chung J. Loss of UCHL1 rescues the defects related to Parkinson’s disease by suppressing glycolysis. Sci Adv. 2021;7(28):eabg4574. doi:10.1126/sciadv.abg4574

1053. DeStefano AL, Latourelle J, Lew MF, Suchowersky O, Klein C, Golbe LI, Mark MH, Growdon JH, Wooten GF, Watts R, et al. eplication of association between ELAVL4 and Parkinson disease: the GenePD study. Hum Genet. 2008;124(1):95–99. doi:10.1007/s00439-008-0526-4

1054. Guo S, Gao Y, Zhao Y. Neuroprotective microRNA-381 Binds to Repressed Early Growth Response 1 (EGR1) and Alleviates Oxidative Stress Injury in Parkinson’s Disease. ACS Chem Neurosci. 2023;14(11):1981–1991. doi:10.1021/acschemneuro.2c00724

1055. Sabbir MG, Speth RC, Albensi BC. Loss of Cholinergic Receptor Muscarinic 1 (CHRM1) Protein in the Hippocampus and Temporal Cortex of a Subset of Individuals with Alzheimer’s Disease, Parkinson’s Disease, or Frontotemporal Dementia: Implications for Patient Survival. J Alzheimers Dis. 2022;90(2):727–747. doi:10.3233/JAD-220766

1056. D’Andrea MR, Ilyin S, Plata-Salaman CR. Abnormal patterns of microtubule-associated protein-2 (MAP-2) immunolabeling in neuronal nuclei and Lewy bodies in Parkinson’s disease substantia nigra brain tissues. Neurosci Lett. 2001;306(3):137–140. doi:10.1016/s0304-3940(01)01811-0

1057. Tan S, Chi H, Wang P, Zhao R, Zhang Q, Gao Z, Xue H, Tang Q, Li G. Protein tyrosine phosphatase receptor type O serves as a key regulator of insulin resistance-induced ?-synuclein aggregation in Parkinson’s disease. Cell Mol Life Sci. 2024;81(1):403. doi:10.1007/s00018-024-05436-4

1058. Chang KH, Chen CM, Chen YC, Fung HC, Wu YR. Polymorphisms of ACMSD-TMEM163, MCCC1, and BCKDK-STX1B Are Not Associated with Parkinson’s Disease in Taiwan. Parkinsons Dis. 2019;2019:3489638. doi:10.1155/2019/3489638

1059. Lázaro-Figueroa A, Hernández-Medrano AJ, Ramírez-Pineda DB, Navarro Cadavid A, Makarious M, Foo JN, Alvarado CX, Bandres-Ciga S, Periñan MT; Global Parkinson’s Genetics Program (GP2). s SH3GL2 p.G276V the Causal Functional Variant Underlying Parkinson’s Disease Risk at this Locus?. Mov Disord. 2024;39(11):2117–2119. doi:10.1002/mds.29719

1060. Fong CS, Shyu HY, Shieh JC, Fu YP, Chin TY, Wang HW, Cheng CW. Association of MTHFR, MTR, and MTRR polymorphisms with Parkinson’s disease among ethnic Chinese in Taiwan. Clin Chim Acta. 2011;412(3-4):332–338. doi:10.1016/j.cca.2010.11.004

1061. Guo C, Zhu J, Wang J, Duan J, Ma S, Yin Y, Quan W, Zhang W, Guan Y, Ding Y, et al. Neuroprotective effects of protocatechuic aldehyde through PLK2/p-GSK3β/Nrf2 signaling pathway in both in vivo and in vitro models of Parkinson’s disease. Aging (Albany NY). 2019;11(21):9424–9441. doi:10.18632/aging.102394

1062. Theuns J, Verstraeten A, Sleegers K, Wauters E, Gijselinck I, Smolders S, Crosiers D, Corsmit E, Elinck E, Sharma M, et al. Global investigation and meta-analysis of the C9orf72 (G4C2)n repeat in Parkinson disease. Neurology. 2014;83(21):1906–1913. doi:10.1212/WNL.0000000000001012

1063. Strittmatter M, Hamann GF, Strubel D, Cramer H, Schimrigk K. Somatostatin-like immunoreactivity, its molecular forms and monoaminergic metabolites in aged and demented patients with Parkinson’s disease--effect of L-Dopa. J Neural Transm (Vienna). 1996;103(5):591–602. doi:10.1007/BF01273156

1064. Lanoue AC, Blatt GJ, Soghomonian JJ. Decreased parvalbumin mRNA expression in dorsolateral prefrontal cortex in Parkinson’s disease. Brain Res. 2013;1531:37–47. doi:10.1016/j.brainres.2013.07.025

1065. Chang CH, Lim KL, Foo JN. Synaptic Vesicle Glycoprotein 2C: a role in Parkinson’s disease. Front Cell Neurosci. 2024;18:1437144. doi:10.3389/fncel.2024.1437144

1066. Beyer K, Ispierto L, Latorre P, Tolosa E, Ariza A. Alpha- and beta-synuclein expression in Parkinson disease with and without dementia. J Neurol Sci. 2011;310(1-2):112–117. doi:10.1016/j.jns.2011.05.049

1067. Tanji K, Kamitani T, Mori F, Kakita A, Takahashi H, Wakabayashi K. TRIM9, a novel brain-specific E3 ubiquitin ligase, is repressed in the brain of Parkinson’s disease and dementia with Lewy bodies. Neurobiol Dis. 2010;38(2):210–218. doi:10.1016/j.nbd.2010.01.007

1068. Nedic Erjavec G, Grubor M, Zivkovic M, Bozina N, Sagud M, Nikolac Perkovic M, Mihaljevic-Peles A, Pivac N, Svob Strac D. SLC6A3, HTR2C and HTR6 Gene Polymorphisms and the Risk of Haloperidol-Induced Parkinsonism. Biomedicines. 2022;10(12):3237.doi:10.3390/biomedicines10123237

1069. Faustini G, Longhena F, Masato A, Bassareo V, Frau R, Klingstedt T, Shirani H, Brembati V, Parrella E, Vezzoli M, et al. Synapsin III gene silencing redeems alpha-synuclein transgenic mice from Parkinson’s disease-like phenotype. Mol Ther. 2022;30(4):1465–1483. doi:10.1016/j.ymthe.2022.01.021

1070. Sukhanov I, Dorotenko A, Fesenko Z, Savchenko A, Efimova EV, Mor MS, Belozertseva IV, Sotnikova TD, Gainetdinov RR. Inhibition of PDE10A in a New Rat Model of Severe Dopamine Depletion Suggests New Approach to Non-Dopamine Parkinson’s Disease Therapy. Biomolecules. 2022;13(1):9. doi:10.3390/biom13010009

1071. Pérez-Santamarina E, García-Ruiz P, Martínez-Rubio D, Ezquerra M, Pla-Navarro I, Puente J, Martí MJ, Palau F, Hoenicka J. Author Correction: Regulatory rare variants of the dopaminergic gene ANKK1 as potential risk factors for Parkinson’s disease. Sci Rep. 2022;12(1):12719. doi:10.1038/s41598-022-17301-0

1072. Turcotte MA, Garant JM, Cossette-Roberge H, Perreault JP. Guanine Nucleotide-Binding Protein-Like 1 (GNL1) binds RNA G-quadruplex structures in genes associated with Parkinson’s disease. RNA Biol. 2021;18(9):1339–1353. doi:10.1080/15476286.2020.1847866

1073. Verbovaya ER, Kadnikov IA, Logvinov IO, Antipova TA, Voronin MV, Seredenin SB. In vitro modelling of Parkinson’s disease using 6-OHDA is associated with increased NQO2 activity. Toxicol In Vitro. 2024;101:105940. doi:10.1016/j.tiv.2024.105940

1074. Geldenhuys WJ, Benkovic SA, Lin L, Yonutas HM, Crish SD, Sullivan PG, Darvesh AS, Brown CM, Richardson JR. MitoNEET (CISD1) Knockout Mice Show Signs of Striatal Mitochondrial Dysfunction and a Parkinson’s Disease Phenotype. ACS Chem Neurosci. 2017;8(12):2759–2765. doi:10.1021/acschemneuro.7b00287

1075. Fu Y, Pickford R, Galper J, Phan K, Wu P, Li H, Kim YB, Dzamko N, Halliday GM, Kim WS. A protective role of ABCA5 in response to elevated sphingomyelin levels in Parkinson’s disease. NPJ Parkinsons Dis. 2024;10(1):20. doi:10.1038/s41531-024-00632-2

1076. Haga H, Yamada R, Izumi H, Shinoda Y, Kawahata I, Miyachi H, Fukunaga K. Novel fatty acid-binding protein 3 ligand inhibits dopaminergic neuronal death and improves motor and cognitive impairments in Parkinson’s disease model mice. Pharmacol Biochem Behav. 2020;191:172891. doi:10.1016/j.pbb.2020.172891

1077. Liu W, Shi R, Yang W, Zhao N, Du Y, Zou Y, Yu W. Synchronous alteration pattern between serine-threonine kinase receptor-associated protein and Smad7 in pilocarpine-induced rats of epilepsy. Synapse. 2014;68(6):275–282. doi:10.1002/syn.21739

1078. Chen Y, Chen J, Chen Y, Li Y. miR-146a/KLF4 axis in epileptic mice: A novel regulator of synaptic plasticity involving STAT3 signaling. Brain Res. 2022;1790:147988. doi:10.1016/j.brainres.2022.147988

1079. Jin X, Liao X, Wu L, Huang J, Li Z, Li Y, Guo F. FOXO4 alleviates hippocampal neuronal damage in epileptic mice via the miR-138-5p/ROCK2 axis. Am J Med Genet B Neuropsychiatr Genet. 2022;189(7-8):271–284. doi:10.1002/ajmg.b.32904

1080. Ma L, Wu Q, Yuan J, Wang Y, Zhang P, Liu Q, Tan D, Liang M, Chen Y. Inhibition of ANXA2 activity attenuates epileptic susceptibility and GluA1 phosphorylation. CNS Neurosci Ther. 2023;29(11):3644–3656. doi:10.1111/cns.14295

1081. Talwar P, Kanojia N, Mahendru S, Baghel R, Grover S, Arora G, Grewal GK, Parween S, Srivastava A, Singh M, et al. Genetic contribution of CYP1A1 variant on treatment outcome in epilepsy patients: a functional and interethnic perspective. Pharmacogenomics J. 2017;17(3):242–251. doi:10.1038/tpj.2016.1

1082. Zhai J, Wang C, Jin L, Liu F, Xiao Y, Gu H, Liu M, Chen Y. Gut Microbiota Metabolites Mediate Bax to Reduce Neuronal Apoptosis via cGAS/STING Axis in Epilepsy. Mol Neurobiol. 2023. doi:10.1007/s12035-023-03545-y

1083. Wang R, Zeng GQ, Tong RZ, Zhou D, Hong Z. Serum matrix metalloproteinase-2: A potential biomarker for diagnosis of epilepsy. Epilepsy Res. 2016;122:114–119. doi:10.1016/j.eplepsyres.2016.02.009

1084. Ercegovac M, Jovic N, Sokic D, Savic-Radojevic A, Coric V, Radic T, Nikolic D, Kecmanovic M, Matic M, Simic T, et al. GSTA1, GSTM1, GSTP1 and GSTT1 polymorphisms in progressive myoclonus epilepsy: A Serbian case-control study. Seizure. 2015;32:30–36. doi:10.1016/j.seizure.2015.08.010

1085. Česká K, Papež J, Ošlejšková H, Slabý O, Radová L, Loja T, Libá Z, Svěráková A, Brázdil M, Aulická Š., et al. CCL2/MCP-1, interleukin-8, and fractalkine/CXC3CL1: Potential biomarkers of epileptogenesis and pharmacoresistance in childhood epilepsy. Eur J Paediatr Neurol. 2023;46:48–54. doi:10.1016/j.ejpn.2023.06.001

1086. Tiron C, Campuzano O, Pérez-Serra A, Mademont I, Coll M, Allegue C, Iglesias A, Partemi S, Striano P, Oliva A, et al. Further evidence of the association between LQT syndrome and epilepsy in a family with KCNQ1 pathogenic variant. Seizure. 2015;25:65–67. doi:10.1016/j.seizure.2015.01.003

1087. Horta WG, Paradela E, Figueiredo A, Meira ID, Pereira VC, Rego CC, Oliveira R, Andraus ME, de Lacerda GC, Moura P, et al. Genetic association study of the HLA class II alleles DRB1, DQA1, and DQB1 in patients with pharmacoresistant temporal lobe epilepsy associated with mesial hippocampal sclerosis. Seizure. 2015;31:7–11. doi:10.1016/j.seizure.2015.06.005

1088. Shao LL, Gao MM, Gong JX, Yang LY. DUSP1 regulates hippocampal damage in epilepsy rats via ERK1/2 pathway. J Chem Neuroanat. 2021;118:102032. doi:10.1016/j.jchemneu.2021.102032

1089. Zan X, Yue G, Hao Y, Sima X. A systematic review and meta-analysis of the association of ABCC2/ABCG2 polymorphisms with antiepileptic drug responses in epileptic patients. Epilepsy Res. 2021;175:106678. doi:10.1016/j.eplepsyres.2021.106678

1090. Shi NR, Wang Q, Liu J, Zhang JZ, Deng BL, Hu XM, Yang J, Wang X, Chen X, Zuo YQ, et al. Association of the ADORA2A receptor and CD73 polymorphisms with epilepsy. Front Pharmacol. 2023;14:1152667. doi:10.3389/fphar.2023.1152667

1091. Heinrich A, Zhong XB, Rasmussen TP. Variability in expression of the human MDR1 drug efflux transporter and genetic variation of the ABCB1 gene: implications for drug-resistant epilepsy. Curr Opin Toxicol. 2018;11–12:35–42. doi:10.1016/j.cotox.2018.12.004

1092. Algahtani HA, Shirah BH, Samman A, Alhazmi A. Epilepsy and Hearing Loss in a Patient with a Rare Heterozygous Variant in the CACNA1H Gene. J Epilepsy Res. 2022;12(1):33–35. doi:10.14581/jer.22006

1093. Chen Z, Luo S, Liu ZG, Deng YC, He SL, Liu XR, Yi YH, Wang J, Gao LD, Li BM, et al. CELSR1 variants are associated with partial epilepsy of childhood. Am J Med Genet B Neuropsychiatr Genet. 2022;189(7-8):247–256. doi:10.1002/ajmg.b.32916

1094. Hu T, Zeng X, Tian T, Liu J. Association of EPHX1 polymorphisms with plasma concentration of carbamazepine in epileptic patients: Systematic review and meta-analysis. J Clin Neurosci. 2021;91:159–171. doi:10.1016/j.jocn.2021.07.009

1095. Majewski L, Wojtas B, Maciąg F, Kuznicki J. Changes in Calcium Homeostasis and Gene Expression Implicated in Epilepsy in Hippocampi of Mice Overexpressing ORAI1. Int J Mol Sci. 2019;20(22):5539. doi:10.3390/ijms20225539

1096. Blümcke I, Schewe JC, Normann S, Brüstle O, Schramm J, Elger CE, Wiestler OD. Increase of nestin-immunoreactive neural precursor cells in the dentate gyrus of pediatric patients with early-onset temporal lobe epilepsy. Hippocampus. 2001;11(3):311–321. doi:10.1002/hipo.1045

1097. Nemes AD, Ayasoufi K, Ying Z, Zhou QG, Suh H, Najm IM. Growth Associated Protein 43 (GAP-43) as a Novel Target for the Diagnosis, Treatment and Prevention of Epileptogenesis. Sci Rep. 2017;7(1):17702. doi:10.1038/s41598-017-17377-z

1098. Hong W, Haviland I, Pestana-Knight E, Weisenberg JL, Demarest S, Marsh ED, Olson HE. CDKL5 Deficiency Disorder-Related Epilepsy: A Review of Current and Emerging Treatment. CNS Drugs. 2022;36(6):591–604. doi:10.1007/s40263-022-00921-5

1099. Liu Y, Cai N, Xu F, Shi Y, Wang Z, Wang N, Chen W, Yang K. Deep brain stimulation in progressive myoclonus epilepsy with SERPINI1 mutation. Parkinsonism Relat Disord. 2024;127:107085. doi:10.1016/j.parkreldis.2024.107085

1100. Guo M, Cui C, Song X, Jia L, Li D, Wang X, Dong H, Ma Y, Liu Y, Cui Z, et al. Deletion of FGF9 in GABAergic neurons causes epilepsy. Cell Death Dis. 2021;12(2):196. doi:10.1038/s41419-021-03478-1

1101. Wang P, Zhang Y, Wang Z, Yang A, Li Y, Zhang Q. miR-128 regulates epilepsy sensitivity in mice by suppressing SNAP-25 and SYT1 expression in the hippocampus. Biochem Biophys Res Commun. 2021;545:195–202. doi:10.1016/j.bbrc.2021.01.079

1102. Klöckner C, Sticht H, Zacher P, Popp B, Babcock HE, Bakker DP, Barwick K, Bonfert MV, Bönnemann CG, Brilstra EH; et al. De novo variants in SNAP25 cause an early-onset developmental and epileptic encephalopathy. Genet Med. 2021;23(4):653–660. doi:10.1038/s41436-020-01020-w

1103. Zhang Q, Forster-Gibson C, Bercovici E, Bernardo A, Ding F, Shen W, Langer K, Rex T, Kang JQ. Epilepsy plus blindness in microdeletion of GABRA1 and GABRG2 in mouse and human. Exp Neurol. 2023;369:114537. doi:10.1016/j.expneurol.2023.114537

1104. Ohori S, Miyauchi A, Osaka H, Lourenco CM, Arakaki N, Sengoku T, Ogata K, Honjo RS, Kim CA, Mitsuhashi S, et al. Biallelic structural variations within FGF12 detected by long-read sequencing in epilepsy. Life Sci Alliance. 2023;6(8):e202302025. doi:10.26508/lsa.202302025

1105. Zheng D, Li M, Li G, Hu J, Jiang X, Wang Y, Sun Y. Circular RNA circ_DROSHA alleviates the neural damage in a cell model of temporal lobe epilepsy through regulating miR-106b-5p/MEF2C axis. Cell Signal. 2021;80:109901. doi:10.1016/j.cellsig.2020.109901

1106. Bleakley LE, Reid CA. HCN1 epilepsy: From genetics and mechanisms to precision therapies. J Neurochem. 2023. doi:10.1111/jnc.15928

1107. Shimizu A, Asakawa S, Sasaki T, Yamazaki S, Yamagata H, Kudoh J, Minoshima S, Kondo I, Shimizu N. A novel giant gene CSMD3 encoding a protein with CUB and sushi multiple domains: a candidate gene for benign adult familial myoclonic epilepsy on human chromosome 8q23.3-q24.1. Biochem Biophys Res Commun. 2003;309(1):143–154. doi:10.1016/s0006-291x(03)01555-9

1108. Panneerselvam S, Wang J, Zhu W, Dai H, Pappas JG, Rabin R, Low KJ, Rosenfeld JA, Emrick L, Xiao R, et al. PPP3CA truncating variants clustered in the regulatory domain cause early-onset refractory epilepsy. Clin Genet. 2021;100(2):227–233. doi:10.1111/cge.13979

1109. Zeng Q, Yang Y, Duan J, Niu X, Chen Y, Wang D, Zhang J, Chen J, Yang X, Li J, et al. CN2A-Related Epilepsy: The Phenotypic Spectrum, Treatment and Prognosis. Front Mol Neurosci. 2022;15:809951. doi:10.3389/fnmol.2022.809951

1110. Remme CA. SCN5A channelopathy: arrhythmia, cardiomyopathy, epilepsy and beyond. Philos Trans R Soc Lond B Biol Sci. 2023;378(1879):20220164. doi:10.1098/rstb.2022.0164

1111. Ul Mudassir B, Agha Z. Novel and known minor alleles of CNTNAP2 gene variants are associated with comorbidity of intellectual disability and epilepsy phenotypes: a case-control association study reveals potential biomarkers. Mol Biol Rep. 2024;51(1):276. doi:10.1007/s11033-023-09176-9

1112. Yang Y, Zeng Q, Cheng M, Niu X, Xiangwei W, Gong P, Li W, Ma J, Zhang X, Yang X, et al. GABRB3-related epilepsy: novel variants, clinical features and therapeutic implications. J Neurol. 2022;269(5):2649–2665. doi:10.1007/s00415-021-10834-w

1113. Brunklaus A, Brünger T, Feng T, Fons C, Lehikoinen A, Panagiotakaki E, Vintan MA, Symonds J, Andrew J, Arzimanoglou A, et al. The gain of function SCN1A disorder spectrum: novel epilepsy phenotypes and therapeutic implications. Brain. 2022;145(11):3816–3831. doi:10.1093/brain/awac210

1114. Fang M, Liu GW, Pan YM, Shen L, Li CS, Xi ZQ, Xiao F, Wang L, Chen D, Wang XF. Abnormal expression and spatiotemporal change of Slit2 in neurons and astrocytes in temporal lobe epileptic foci: A study of epileptic patients and experimental animals. Brain Res. 2010;1324:14–23. doi:10.1016/j.brainres.2010.02.007

1115. Baglietto MG, Caridi G, Gimelli G, Mancardi M, Prato G, Ronchetto P, Cuoco C, Tassano E. RORB gene and 9q21.13 microdeletion: report on a patient with epilepsy and mild intellectual disability. Eur J Med Genet. 2014;57(1):44–46. doi:10.1016/j.ejmg.2013.12.001

1116. Vega-García A, Orozco-Suárez S, Villa A, Rocha L, Feria-Romero I, Alonso Vanegas MA, Guevara-Guzmán R. Cortical expression of IL1-?, Bcl-2, Caspase-3 and 9, SEMA-3a, NT-3 and P-glycoprotein as biological markers of intrinsic severity in drug-resistant temporal lobe epilepsy. Brain Res. 2021;1758:147303. doi:10.1016/j.brainres.2021.147303

1117. Aksoy HU, Yılmaz C, Orak SA, Ayça S, Polat M. Evaluation of GFAP, S100B, and UCHL-1 Levels in Children With Refractory Epilepsy. J Child Neurol. 2024;39(9-10):317–323. doi:10.1177/08830738241273339

1118. Gurnett CA, Veile R, Zempel J, Blackburn L, Lovett M, Bowcock A. Disruption of sodium bicarbonate transporter SLC4A10 in a patient with complex partial epilepsy and mental retardation. Arch Neurol. 2008;65(4):550–553. doi:10.1001/archneur.65.4.550

1119. Koskinen LL, Seppälä EH, Weissl J, Jokinen TS, Viitmaa R, Hänninen RL, Quignon P, Fischer A, André C, Lohi H. ADAM23 is a common risk gene for canine idiopathic epilepsy. BMC Genet. 2017;18(1):8. doi:10.1186/s12863-017-0478-6

1120. Dong Z, Min F, Zhang S, Zhang H, Zeng T. EGR1-Driven METTL3 Activation Curtails VIM-Mediated Neuron Injury in Epilepsy. Neurochem Res. 2023;48(11):3349–3362. doi:10.1007/s11064-023-03950-8

1121. Srivastava S, Cohen J, Pevsner J, Aradhya S, McKnight D, Butler E, Johnston M, Fatemi A. A novel variant in GABRB2 associated with intellectual disability and epilepsy. Am J Med Genet A. 2014;164A(11):2914–2921. doi:10.1002/ajmg.a.36714

1122. Marcé-Grau A, Elorza-Vidal X, Pérez-Rius C, Ruiz-Nel Lo A, Sala- Coromina J, Gabau E, Estévez R, Macaya A. Muscarinic acetylcholine receptor M1 mutations causing neurodevelopmental disorder and epilepsy. Hum Mutat. 2021;42(10):1215–1220. doi:10.1002/humu.24252

1123. Xue J, Song Z, Ma S, Yi Z, Yang C, Li F, Liu K, Zhang Y. A Novel De Novo TUBB3 Variant Causing Developmental Delay, Epilepsy and Mild Ophthalmological Symptoms in a Chinese Child. J Mol Neurosci. 2022;72(1):37–44. doi:10.1007/s12031-021-01909-4

1124. Westphal DS, Andres S, Makowski C, Meitinger T, Hoefele J. MAP2 - A Candidate Gene for Epilepsy, Developmental Delay and Behavioral Abnormalities in a Patient With Microdeletion 2q34. Front Genet. 2018;9:99. doi:10.3389/fgene.2018.00099

1125. Villacres JE, Riveira N, Kim S, Colgin LL, Noebels JL, Lopez AY. Abnormal patterns of sleep and waking behaviors are accompanied by neocortical oscillation disturbances in an Ank3 mouse model of epilepsy- bipolar disorder comorbidity. Transl Psychiatry. 2023;13(1):403. doi:10.1038/s41398-023-02700-2

1126. Ke P, Gu J, Liu J, Liu Y, Tian X, Ma Y, Meng Y, Xiao F. Syntabulin regulates neuronal excitation/inhibition balance and epileptic seizures by transporting syntaxin 1B. Cell Death Discov. 2023;9(1):187. doi:10.1038/s41420-023-01461-7

1127. Desprairies C, Valence S, Maurey H, Helal SI, Weckhuysen S, Soliman H, Mefford HC, Spentchian M, Héron D, Leguern E, et al. Three novel patients with epileptic encephalopathy due to biallelic mutations in the PLCB1 gene. Clin Genet. 2020;97(3):477–482. doi:10.1111/cge.13696

1128. Rochtus AM, Trowbridge S, Goldstein RD, Sheidley BR, Prabhu SP, Haynes R, Kinney HC, Poduri AH. Mutations in NRXN1 and NRXN2 in a patient with early-onset epileptic encephalopathy and respiratory depression. Cold Spring Harb Mol Case Stud. 2019;5(1):a003442. doi:10.1101/mcs.a003442

1129. Xing Y, Cui T, Sun F. A novel RyR2 mutation associated with co- morbid catecholaminergic polymorphic ventricular tachycardia (CPVT) and benign epilepsy with centrotemporal spikes (BECTS). J Electrocardiol. 2024;84:75–80. doi:10.1016/j.jelectrocard.2024.03.013

1130. Luo FM, Deng MX, Yu R, Liu L, Fan LL. Case Report: Chorea-Acanthocytosis Presents as Epilepsy in a Consanguineous Family With a Nonsense Mutation of in VPS13A. Front Neurosci. 2021;15:604715. doi:10.3389/fnins.2021.604715

1131. Komulainen-Ebrahim J, Saastamoinen E, Rahikkala E, Helander H, Hinttala R, Risteli L, Rantala H, Uusimaa J. Intractable Epilepsy due to MTR Deficiency: Importance of Homocysteine Analysis. Neuropediatrics. 2017;48(6):467–472. doi:10.1055/s-0037-1603976

1132. Cai S, Li J, Wu Y, Jiang Y. De novo mutations of TUBB2A cause infantile-onset epilepsy and developmental delay. J Hum Genet. 2020;65(7):601–608. doi:10.1038/s10038-020-0739-5

1133. Muroni A, Floris G, Polizzi L, Fadda L, Piga G, Primicerio G, Rocchi L, Defazio G. Does epilepsy contribute to the clinical phenotype of C9orf72 mutation in fronto-temporal dementia?. Epilepsy Behav. 2022;133:108783. doi:10.1016/j.yebeh.2022.108783

1134. Wengert ER, Miralles RM, Wedgwood KCA, Wagley PK, Strohm SM, Panchal PS, Idrissi AM, Wenker IC, Thompson JA, Gaykema RP, et al. Somatostatin-Positive Interneurons Contribute to Seizures in SCN8A Epileptic Encephalopathy. J Neurosci. 2021;41(44):9257–9273. doi:10.1523/JNEUROSCI.0718-21.2021

1135. Wang Y, Liang J, Chen L, Shen Y, Zhao J, Xu C, Wu X, Cheng H, Ying X, Guo Y, et al. Pharmaco-genetic therapeutics targeting parvalbumin neurons attenuate temporal lobe epilepsy. Neurobiol Dis. 2018;117:149–160. doi:10.1016/j.nbd.2018.06.006

1136. Ranta S, Zhang Y, Ross B, Takkunen E, Hirvasniemi A, de la Chapelle A, Gilliam TC, Lehesjoki AE. Positional cloning and characterisation of the human DLGAP2 gene and its exclusion in progressive epilepsy with mental retardation. Eur J Hum Genet. 2000;8(5):381–384. doi:10.1038/sj.ejhg.5200440

1137. Tian MQ, Li RK, Yang F, Shu XM, Li J, Chen J, Peng LY, Yu XH, Yang CJ. Phenotypic expansion of KCNH1-associated disorders to include isolated epilepsy and its associations with genotypes and molecular sub- regional locations. CNS Neurosci Ther. 2023;29(1):270–281. doi:10.1111/cns.14001

1138. Hayashi T, Yano N, Kora K, Yokoyama A, Maizuru K, Kayaki T, Nishikawa K, Osawa M, Niwa A, Takenouchi T, et al. Involvement of mTOR pathway in neurodegeneration in NSF-related developmental and epileptic encephalopathy. Hum Mol Genet. 2023;32(10):1683–1697. doi:10.1093/hmg/ddad008

1139. Mertens A, Papadopoulou MT, Papathanasiou Terzi MA, Lesca G, Biela M, Smigiel R, Panagiotakaki E. Epilepsy with eyelid myoclonia in a patient with ATP1A3-related neurologic disorder. Epileptic Disord. 2024. doi:10.1002/epd2.20272

1140. Okano S, Makita Y, Miyamoto A, Taketazu G, Kimura K, Fukuda I, Tanaka H, Yanagi K, Kaname T. GRIA3 p.Met661Thr variant in a female with developmental epileptic encephalopathy. Hum Genome Var. 2023;10(1):4. doi:10.1038/s41439-023-00232-1

1141. Neuray C, Maroofian R, Scala M, Sultan T, Pai GS, Mojarrad M, Khashab HE, deHoll L, Yue W, Alsaif HS, et al. Early-infantile onset epilepsy and developmental delay caused by bi-allelic GAD1 variants. Brain. 2020;143(8):2388–2397. doi:10.1093/brain/awaa178

1142. Pazarlar BA, Aripaka SS, Petukhov V, Pinborg L, Khodosevich K, Mikkelsen JD. Expression profile of synaptic vesicle glycoprotein 2A, B, and C paralogues in temporal neocortex tissue from patients with temporal lobe epilepsy (TLE). Mol Brain. 2022;15(1):45. doi:10.1186/s13041-022-00931-w

1143. Massey CA, Thompson SJ, Ostrom RW, Drabek J, Sveinsson OA, Tomson T, Haas EA, Mena OJ, Goldman AM, Noebels JL. X-linked serotonin 2C receptor is associated with a non-canonical pathway for sudden unexpected death in epilepsy. Brain Commun. 2021;3(3):fcab149. doi:10.1093/braincomms/fcab149

1144. Sheng W, Wang P, Cai Y, Zhai C, Wang H, Zhou F, Liu X, Wang L, Li D, Shu J, et al. Epilepsy due to potential loss of ATP6V1B2 function with mechanistic insight by a Drosophila Vha55 model. Clin Genet. 2024;106(6):702–712. doi:10.1111/cge.14600

1145. Carvill GL. Calcium Channel Dysfunction in Epilepsy: Gain of CACNA1E. Epilepsy Curr. 2019;19(3):199–201. doi:10.1177/1535759719845324

1146. Alsemari A, Al-Younes B, Goljan E, Jaroudi D, BinHumaid F, Meyer BF, Arold ST, Monies D. Recessive VARS2 mutation underlies a novel syndrome with epilepsy, mental retardation, short stature, growth hormone deficiency, and hypogonadism. Hum Genomics. 2017;11(1):28. doi:10.1186/s40246-017-0124-4

1147. Polani S, Dean M, Lichter-Peled A, Hendrickson S, Tsang S, Fang X, Feng Y, Qiao W, Avni G, Kahila Bar-Gal G. Sequence Variant in the TRIM39-RPP21 Gene Readthrough is Shared Across a Cohort of Arabian Foals Diagnosed with Juvenile Idiopathic Epilepsy. J Genet Mutat Disord. 2022;1(1):103.

1148. Jiang J, Feng Y, Tang Q, Zhao C, Guo M, Wu J, Guo R, Lu H, Sun X, Gao J, et al. Novel IARS1 variants cause syndromic developmental disorder with epilepsy in a Chinese patient and the literature review. Mol Genet Genomic Med. 2024;12(1):e2326. doi:10.1002/mgg3.2326

1149. Li H, Wang X, Zhou Y, Ni G, Su Q, Chen Z, Chen Z, Li J, Chen X, Hou X, et al. Association of LEPR and ANKK1 Gene Polymorphisms with Weight Gain in Epilepsy Patients Receiving Valproic Acid. Int J Neuropsychopharmacol. 2015;18(7):pyv021.doi:10.1093/ijnp/pyv021

1150. Zeng C, Zhou P, Jiang T, Yuan C, Ma Y, Feng L, Liu R, Tang W, Long X, Xiao B, et al. Upregulation and Diverse Roles of TRPC3 and TRPC6 in Synaptic Reorganization of the Mossy Fiber Pathway in Temporal Lobe Epilepsy. Mol Neurobiol. 2015;52(1):562–572. doi:10.1007/s12035-014-8871-x

1151. Miller JP, Yates BE, Al-Ramahi I, Berman AE, Sanhueza M, Kim E, de Haro M, DeGiacomo F, Torcassi C, Holcomb J, et al. A genome-scale RNA-interference screen identifies RRAS signaling as a pathologic feature of Huntington’s disease. PLoS Genet. 2012;8(11):e1003042. doi:10.1371/journal.pgen.1003042

1152. Sharma M, Rajendrarao S, Shahani N, Ramírez-Jarquín UN, Subramaniam S. Cyclic GMP-AMP synthase promotes the inflammatory and autophagy responses in Huntington disease. Proc Natl Acad Sci U S A. 2020;117(27):15989–15999. doi:10.1073/pnas.2002144117

1153. Taylor DM, Moser R, Régulier E, Breuillaud L, Dixon M, Beesen AA, Elliston L, Silva Santos Mde F, Kim J, Jones L, et al. MAP kinase phosphatase 1 (MKP-1/DUSP1) is neuroprotective in Huntington’s disease via additive effects of JNK and p38 inhibition. J Neurosci. 2013;33(6):2313–2325. doi:10.1523/JNEUROSCI.4965-11.2013

1154. Taherzadeh-Fard E, Saft C, Wieczorek S, Epplen JT, Arning L. Age at onset in Huntington’s disease: replication study on the associations of ADORA2A, HAP1 and OGG1. Neurogenetics. 2010;11(4):435–439. doi:10.1007/s10048-010-0248-3

1155. Elbaz EM, Helmy HS, El-Sahar AE, Saad MA, Sayed RH. Lercanidipine boosts the efficacy of mesenchymal stem cell therapy in 3- NP-induced Huntington’s disease model rats via modulation of the calcium/calcineurin/NFATc4 and Wnt/?-catenin signalling pathways. Neurochem Int. 2019;131:104548. doi:10.1016/j.neuint.2019.104548

1156. Noda Y, Shimazawa M, Tanaka H, Tamura S, Inoue T, Tsuruma K, Hara H. VGF and striatal cell damage in in vitro and in vivo models of Huntington’s disease. Pharmacol Res Perspect. 2015;3(3):e00140. doi:10.1002/prp2.140

1157. Yusuf IO, Chen HM, Cheng PH, Chang CY, Tsai SJ, Chuang JI, Wu CC, Huang BM, Sun HS, Chen CM, et al. FGF9 induces neurite outgrowth upon ERK signaling in knock-in striatal Huntington’s disease cells. Life Sci. 2021;267:118952. doi:10.1016/j.lfs.2020.118952

1158. Smith R, Klein P, Koc-Schmitz Y, Waldvogel HJ, Faull RL, Brundin P, Plomann M, Li JY. Loss of SNAP-25 and rabphilin 3a in sensory-motor cortex in Huntington’s disease. J Neurochem. 2007;103(1):115–123. doi:10.1111/j.1471-4159.2007.04703.x

1159. Xu EH, Tang Y, Li D, Jia JP. Polymorphism of HD and UCHL-1 genes in Huntington’s disease. J Clin Neurosci. 2009;16(11):1473–1477. doi:10.1016/j.jocn.2009.03.027

1160. Lee J, Hwang YJ, Shin JY, Lee WC, Wie J, Kim KY, Lee MY, Hwang D, Ratan RR, Pae AN, et al. Epigenetic regulation of cholinergic receptor M1 (CHRM1) by histone H3K9me3 impairs Ca(2+) signaling in Huntington’s disease. Acta Neuropathol. 2013;125(5):727–739. doi:10.1007/s00401-013-1103-z

1161. Cabrera JR, Lucas JJ. MAP2 Splicing is Altered in Huntington’s Disease. Brain Pathol. 2017;27(2):181–189. doi:10.1111/bpa.12387

1162. Cho K, Kim GW. Neurexin1 level in Huntington’s Disease and decreased Neurexin1 in disease progression. Neurosci Res. 2024. doi:10.1016/j.neures.2024.10.006

1163. García-García E, Carreras-Caballé M, Coll-Manzano A, Ramón- Lainez A, Besa-Selva G, Pérez-Navarro E, Malagelada C, Alberch J, Masana M, Rodríguez MJ. reserved VPS13A distribution and expression in Huntington’s disease: divergent mechanisms of action for similar movement disorders?. Front Neurosci. 2024;18:1394478. doi:10.3389/fnins.2024.1394478

1164. Alva-Diaz C, Alarcon-Ruiz CA, Pacheco-Barrios K, Mori N, Pacheco-Mendoza J, Traynor BJ, Rivera-Valdivia A, Lertwilaiwittaya P, Bird TD, Cornejo-Olivas M. C9orf72 Hexanucleotide Repeat in Huntington-Like Patients: Systematic Review and Meta-Analysis. Front Genet. 2020;11:551780. doi:10.3389/fgene.2020.551780

1165. Aronin N, Cooper PE, Lorenz LJ, Bird ED, Sagar SM, Leeman SE, Martin JB. omatostatin is increased in the basal ganglia in Huntington disease. Ann Neurol. 1983;13(5):519–526. doi:10.1002/ana.410130508

1166. Beaumont V, Zhong S, Lin H, Xu W, Bradaia A, Steidl E, Gleyzes M, Wadel K, Buisson B, Padovan-Neto FE, et al. Phosphodiesterase 10A Inhibition Improves Cortico-Basal Ganglia Function in Huntington’s Disease Models. Neuron. 2016;92(6):1220–1237. doi:10.1016/j.neuron.2016.10.064

1167. Santos R, Linker SB, Stern S, Mendes APD, Shokhirev MN, Erikson G, Randolph-Moore L, Racha V, Kim Y, Kelsoe JR, et al. Deficient LEF1 expression is associated with lithium resistance and hyperexcitability in neurons derived from bipolar disorder patients. Mol Psychiatry. 2021;26(6):2440–2456. doi:10.1038/s41380-020-00981-3

1168. Mohammadynejad P, Saadat I, Ghanizadeh A, Saadat M. Bipolar disorder and polymorphisms of glutathione S-transferases M1 (GSTM1) and T1 (GSTT1). Psychiatry Res. 2011;186(1):144–146. doi:10.1016/j.psychres.2010.06.017

1169. Ghoryani M, Faridhosseini F, Talaei A, Faridhosseini R, Tavakkol-Afshari J, Dadgar Moghaddam M, Azim P, Salimi Z, Marzouni HZ, Mohammadi M. Gene expression pattern of CCL2, CCL3, and CXCL8 in patients with bipolar disorder. J Res Med Sci. 2019;24:45. doi:10.4103/jrms.JRMS_763_18

1170. Le Clerc S, Lombardi L, Baune BT, Amare AT, Schubert KO, Hou L, Clark SR, Papiol S, Cearns M, Heilbronner U, et al. HLA-DRB1 and HLA- DQB1 genetic diversity modulates response to lithium in bipolar affective disorders. Sci Rep. 2021;11(1):17823. doi:10.1038/s41598-021-97140-7

1171. Zhang T, Rao Q, Lin K, He Y, Cai J, Yang M, Xu Y, Hou L, Lin Y, Liu H. CYP2C19-rs4986893 confers risk to major depressive disorder and bipolar disorder in the Han Chinese population whereas ABCB1-rs1045642 acts as a protective factor. BMC Psychiatry. 2023;23(1):69. doi:10.1186/s12888-022-04514-w

1172. Li J, Liu J, Feng G, Li T, Zhao Q, Li Y, Hu Z, Zheng L, Zeng Z, He L, et al. The MDGA1 gene confers risk to schizophrenia and bipolar disorder. Schizophr Res. 2011;125(2-3):194–200. doi:10.1016/j.schres.2010.11.002

1173. Chen S, Jiang H, Hou Z, Yue Y, Zhang Y, Zhao F, Xu Z, Li Y, Mou X, Li L, et al. Higher serum VGF protein levels discriminate bipolar depression from major depressive disorder. J Neurosci Res. 2019;97(5):597–606. doi:10.1002/jnr.2437

1174. Wang Y, Zhang J, Li X, Ji J, Yang F, Wan C, Feng G, Wan P, He L, He G. SCN8A as a novel candidate gene associated with bipolar disorder in the Han Chinese population. Prog Neuropsychopharmacol Biol Psychiatry. 2008;32(8):1902–1904. doi:10.1016/j.pnpbp.2008.09.003

1175. Li Y, Zhao Q, Zhang Z, Wang P, Che R, Tang W, Feng G, Lindpaintner K, He L, Shi Y. Association study between RGS4 and bipolar disorder in the Chinese Han population. Psychiatr Genet. 2010;20(3):130–132. doi:10.1097/YPG.0b013e32833a2009

1176. Lucash J, Hong W, Swanson L, Maski K, Urion D, Kim J, Leonard H, Olson H. Bipolar Disorder in a female with CDKL5 Deficiency Disorder: A Case Report. Preprint. Res Sq. 2024;rs.3.rs-4851179. doi:10.21203/rs.3.rs-4851179/v1

1177. Lohoff FW, Bloch PJ, Weller AE, Ferraro TN, Berrettini WH. Association analysis of the pituitary adenylate cyclase-activating polypeptide (PACAP/ADCYAP1) gene in bipolar disorder. Psychiatr Genet. 2008;18(2):53–58. doi:10.1097/YPG.0b013e3282f60320

1178. Houenou J, Boisgontier J, Henrion A, d’Albis MA, Dumaine A, Linke J, Wessa M, Daban C, Hamdani N, Delavest M, et al. A Multilevel Functional Study of a SNAP25 At-Risk Variant for Bipolar Disorder and Schizophrenia. J Neurosci. 2017;37(43):10389–10397. doi:10.1523/JNEUROSCI.1040-17.2017

1179. Fatjó-Vilas M, Prats C, Pomarol-Clotet E, Lázaro L, Moreno C, González-Ortega I, Lera-Miguel S, Miret S, Muñoz MJ, Ibáñez I, et al. Involvement of NRN1 gene in schizophrenia-spectrum and bipolar disorders and its impact on age at onset and cognitive functioning. World J Biol Psychiatry. 2016;17(2):129–139. doi:10.3109/15622975.2015.1093658

1180. Slonimsky A, Levy I, Kohn Y, Rigbi A, Ben-Asher E, Lancet D, Agam G, Lerer B. Lymphoblast and brain expression of AHI1 and the novel primate-specific gene, C6orf217, in schizophrenia and bipolar disorder. Schizophr Res. 2010;120(1-3):159–166. doi:10.1016/j.schres.2010.03.041

1181. Xu W, Cohen-Woods S, Chen Q, Noor A, Knight J, Hosang G, Parikh SV, De Luca V, Tozzi F, Muglia P, et al. Genome-wide association study of bipolar disorder in Canadian and UK populations corroborates disease loci including SYNE1 and CSMD1. BMC Med Genet. 2014;15:2. doi:10.1186/1471-2350-15-2

1182. McGrath CL, Glatt SJ, Sklar P, Le-Niculescu H, Kuczenski R, Doyle AE, Biederman J, Mick E, Faraone SV, Niculescu AB, et al. Evidence for genetic association of RORB with bipolar disorder. BMC Psychiatry. 2009;9:70. doi:10.1186/1471-244X-9-70

1183. Chen J, Tsang SY, Zhao CY, Pun FW, Yu Z, Mei L, Lo WS, Fang S, Liu H, Stöber G, et al. GABRB2 in schizophrenia and bipolar disorder: disease association, gene expression and clinical correlations. Biochem Soc Trans. 2009;37(Pt 6):1415–1418. doi:10.1042/BST0371415

1184. Maheshwari H, Garg P, Srivastava P. In silico analysis predicts mutational consequences of CITED2, NUDT4, and Ar18B in patients with bipolar disorder. Behav Brain Res. 2025;476:115257. doi:10.1016/j.bbr.2024.115257

1185. Yosifova A, Mushiroda T, Stoianov D, Vazharova R, Dimova I, Karachanak S, Zaharieva I, Milanova V, Madjirova N, Gerdjikov I, et al. Case-control association study of 65 candidate genes revealed a possible association of a SNP of HTR5A to be a factor susceptible to bipolar disease in Bulgarian population. J Affect Disord. 2009;117(1-2):87–97. doi:10.1016/j.jad.2008.12.021

1186. Gaynor LS, Yadollahikhales G, Tsoy E, Hall M, Boxer AL, Miller BL, Grinberg LT. C9orf72 Repeat Expansion Initially Presenting as Late-Onset Bipolar Disorder With Psychosis. Neurologist. 2024;29(2):109–112. doi:10.1097/NRL.0000000000000527

1187. Pantazopoulos H, Wiseman JT, Markota M, Ehrenfeld L, Berretta S. Decreased Numbers of Somatostatin-Expressing Neurons in the Amygdala of Subjects With Bipolar Disorder or Schizophrenia: Relationship to Circadian Rhythms. Biol Psychiatry. 2017;81(6):536–547. doi:10.1016/j.biopsych.2016.04.006

1188. Philibert RA, Cheung D, Welsh N, Damschroder-Williams P, Thiel B, Ginns EI, Gershenfeld HK. Absence of a significant linkage between Na(+),K(+)-ATPase subunit (ATP1A3 and ATP1B3) genotypes and bipolar affective disorder in the old-order Amish. Am J Med Genet. 2001;105(3):291–294. doi:10.1002/ajmg.1322

1189. Gray L, Scarr E, Dean B. N-Ethylmaleimide sensitive factor in the cortex of subjects with schizophrenia and bipolar I disorder. Neurosci Lett. 2006;391(3):112–115. doi:10.1016/j.neulet.2005.08.051

1190. Gécz J, Barnett S, Liu J, Hollway G, Donnelly A, Eyre H, Eshkevari HS, Baltazar R, Grunn A, Nagaraja R, et al. Characterization of the human glutamate receptor subunit 3 gene (GRIA3), a candidate for bipolar disorder and nonspecific X-linked mental retardation. Genomics. 1999;62(3):356–368. doi:10.1006/geno.1999.6032

1191. Anjanappa RM, Nayak S, Moily NS, Manduva V, Nadella RK, Viswanath B, Reddy YCJ, Jain S, Anand A. A linkage and exome study implicates rare variants of KANK4 and CAP2 in bipolar disorder in a multiplex family. Bipolar Disord. 2020;22(1):70–78. doi:10.1111/bdi.12815

1192. Sitbon J, Nestvogel D, Kappeler C, Nicolas A, Maciuba S, Henrion A, Troudet R, Courtois E, Grannec G, Latapie V, et al. CADPS functional mutations in patients with bipolar disorder increase the sensitivity to stress. Mol Psychiatry. 2022;27(2):1145–1157. doi:10.1038/s41380-021-01151-9

1193. Arrúe A, González-Torres MA, Basterreche N, Arnaiz A, Olivas O, Zamalloa MI, Erkoreka L, Catalán A, Zumárraga M. GAD1 gene polymorphisms are associated with bipolar I disorder and with blood homovanillic acid levels but not with plasma GABA levels. Neurochem Int. 2019;124:152–161. doi:10.1016/j.neuint.2019.01.004

1194. Levchenko A, Vyalova NM, Nurgaliev T, Pozhidaev IV, Simutkin GG, Bokhan NA, Ivanova SA. NRG1, PIP4K2A, and HTR2C as Potential Candidate Biomarker Genes for Several Clinical Subphenotypes of Depression and Bipolar Disorder. Front Genet. 2020;11:936. doi:10.3389/fgene.2020.00936

1195. Jian J, Li C, Xu J, Qiao D, Mi G, Chen X, Tang M. Associations of serotonin receptor gene HTR3A, HTR3B, and HTR3A haplotypes with bipolar disorder in Chinese patients. Genet Mol Res. 2016;15(3):10.4238/gmr.15038671. doi:10.4238/gmr.15038671

1196. MacMullen CM, Vick K, Pacifico R, Fallahi-Sichani M, Davis RL. Novel, primate-specific PDE10A isoform highlights gene expression complexity in human striatum with implications on the molecular pathology of bipolar disorder. Transl Psychiatry. 2016;6(2):e742. doi:10.1038/tp.2016.3

1197. Lee SY, Chen SL, Chang YH, Chen SH, Chu CH, Huang SY, Tzeng NS, Wang CL, Lee IH, Yeh TL, et al. The DRD2/ANKK1 gene is associated with response to add-on dextromethorphan treatment in bipolar disorder. J Affect Disord. 2012;138(3):295–300. doi:10.1016/j.jad.2012.01.024

1198. Roedding AS, Tong SY, Au-Yeung W, Li PP, Warsh JJ. Chronic oxidative stress modulates TRPC3 and TRPM2 channel expression and function in rat primary cortical neurons: relevance to the pathophysiology of bipolar disorder. Brain Res. 2013;1517:16–27. doi:10.1016/j.brainres.2013.04.025

1199. Kowalczyk M, Kucia K, Owczarek A, Suchanek-Raif R, Paul-Samojedny M, Choreza P, Kowalski J. HSPB1 Gene Variants and Schizophrenia: A Case-Control Study in a Polish Population. Dis Markers. 2022;2022:4933011. doi:10.1155/2022/493301

1200. Greenbaum L, Alkelai A, Rigbi A, Kohn Y, Lerer B. Evidence for association of the GLI2 gene with tardive dyskinesia in patients with chronic schizophrenia. Mov Disord. 2010;25(16):2809–2817. doi:10.1002/mds.23377

1201. Li X, Wu X, Li W, Yan Q, Zhou P, Xia Y, Yao W, Zhu F. HERV-W ENV Induces Innate Immune Activation and Neuronal Apoptosis via linc01930/cGAS Axis in Recent-Onset Schizophrenia. Int J Mol Sci. 2023;24(3):3000. doi:10.3390/ijms24033000

1202. Yang H, Peng R, Yang M, Zhang J, Shi Z, Zhang X. Association between elevated serum matrix metalloproteinase-2 and tumor necrosis factor-?, and clinical symptoms in male patients with treatment-resistant and chronic medicated schizophrenia. BMC Psychiatry. 2024;24(1):173. doi:10.1186/s12888-024-05621-6

1203. Liu X, Low SK, Atkins JR, Wu JQ, Reay WR, Cairns HM, Green MJ, Schall U, Jablensky A, Mowry B, et al. Wnt receptor gene FZD1 was associated with schizophrenia in genome-wide SNP analysis of the Australian Schizophrenia Research Bank cohort. Aust N Z J Psychiatry. 2020;54(9):902–908. doi:10.1177/0004867419885443

1204. Pinheiro DS, Santos RDS, de Brito RB, Cruz AHDS, Ghedini PC, Reis AAS. GSTM1/GSTT1 double-null genotype increases risk of treatment-resistant schizophrenia: A genetic association study in Brazilian patients. PLoS One. 2017;12(8):e0183812. doi:10.1371/journal.pone.0183812

1205. Xiong Y, Wei Z, Huo R, Wu X, Shen L, Li Y, Gong X, Wu Z, Feng G, Li W, et al. A pharmacogenetic study of risperidone on chemokine (C-C motif) ligand 2 (CCL2) in Chinese Han schizophrenia patients. Prog Neuropsychopharmacol Biol Psychiatry. 2014;51:153–158. doi:10.1016/j.pnpbp.2014.01.017

1206. Seshasubramanian V, Raghavan V, SathishKannan AD, Naganathan C, Ramachandran A, Arasu P, Rajendren P, John S, Mowry B, Rangaswamy T, et al. Association of HLA-A, -B, -C, -DRB1 and -DQB1 alleles at amino acid level in individuals with schizophrenia: A study from South India. Int J Immunogenet. 2020;47(6):501–511. doi:10.1111/iji.12507

1207. Sun L, Qiu Q, Ban C, Fan S, Xiao S, Li X. Decrease levels of bone morphogenetic protein 6 and noggin in chronic schizophrenia elderly. Cogn Neurodyn. 2023;17(3):695–701. doi:10.1007/s11571-022-09855-6

1208. Foldager L, Steffensen R, Thiel S, Als TD, Nielsen HJ, Nordentoft M, Mortensen PB, Mors O, Jensenius JC. MBL and MASP-2 concentrations in serum and MBL2 promoter polymorphisms are associated to schizophrenia. Acta Neuropsychiatr. 2012;24(4):199–207. doi:10.1111/j.1601-5215.2011.00618.x

1209. Miao J, Liu L, Yan C, Zhu X, Fan M, Yu P, Ji K, Huang Y, Wang Y, Zhu G. Association between ADORA2A gene polymorphisms and schizophrenia in the North Chinese Han population. Neuropsychiatr Dis Treat. 2019;15:2451–2458. doi:10.2147/NDT.S205014

1210. Hattori S, Suda A, Miyauchi M, Shiraishi Y, Saeki T, Fukushima T, Fujibayashi M, Tsujita N, Ishii C, Ishii N, et al. The association of genetic polymorphisms in CYP1A2, UGT1A4, and ABCB1 with autonomic nervous system dysfunction in schizophrenia patients treated with olanzapine. BMC Psychiatry. 2020;20(1):72. doi:10.1186/s12888-020-02492-5

1211. Gross J, Grimm O, Ortega G, Teuber I, Lesch KP, Meyer J. Mutational analysis of the neuronal cadherin gene CELSR1 and exclusion as a candidate for catatonic schizophrenia in a large family. Psychiatr Genet. 2001;11(4):197–200. doi:10.1097/00041444-200112000-00003

1212. Hatayama M, Ishiguro A, Iwayama Y, Takashima N, Sakoori K, Toyota T, Nozaki Y, Odaka YS, Yamada K, Yoshikawa T, et al. Zic2 hypomorphic mutant mice as a schizophrenia model and ZIC2 mutations identified in schizophrenia patients. Sci Rep. 2011;1:16. doi:10.1038/srep00016

1213. Hsiung A, Naya FJ, Chen X, Shiang R. A schizophrenia associated CMYA5 allele displays differential binding with desmin. J Psychiatr Res. 2019;111:8–15. doi:10.1016/j.jpsychires.2019.01.007

1214. Mizoguchi T, Hara H, Shimazawa M. VGF has Roles in the Pathogenesis of Major Depressive Disorder and Schizophrenia: Evidence from Transgenic Mouse Models. Cell Mol Neurobiol. 2019;39(6):721–727. doi:10.1007/s10571-019-00681-9

1215. Shen YC, Tsai HM, Cheng MC, Hsu SH, Chen SF, Chen CH. Genetic and functional analysis of the gene encoding GAP-43 in schizophrenia. Schizophr Res. 2012;134(2-3):239–245. doi:10.1016/j.schres.2011.11.016

1216. Xu FL, Yao J, Wang BJ. Association between RGS4 gene polymorphisms and schizophrenia: A protocol for systematic review and meta-analysis. Medicine (Baltimore). 2021;100(44):e27607. doi:10.1097/MD.0000000000027607

1217. Koga M, Ishiguro H, Horiuchi Y, Inada T, Ujike H, Itokawa M, Otowa T, Watanabe Y, Someya T, Arinami T. Replication study of association between ADCYAP1 gene polymorphisms and schizophrenia. Psychiatr Genet. 2010;20(3):123–125. doi:10.1097/YPG.0b013e32833a1f52

1218. Li XL, Yu Y, Hu Y, Wu HT, Li XS, Chen GY, Cheng Y. Fibroblast Growth Factor 9 as a Potential Biomarker for Schizophrenia. Front Psychiatry. 2022;13:788677. doi:10.3389/fpsyt.2022.788677

1219. Ma Y, Gao K, Sun X, Wang J, Yang Y, Wu J, Chai A, Yao L, Liu N, Yu H, et al. STON2 variations are involved in synaptic dysfunction and schizophrenia-like behaviors by regulating Syt1 trafficking. Sci Bull (Beijing). 2024;69(10):1458–1471. doi:10.1016/j.scib.2024.02.013

1220. Alshammari TK, Alshammari MA, Nenov MN, Hoxha E, Cambiaghi M, Marcinno A, James TF, Singh P, Labate D, Li J, et al. Genetic deletion of fibroblast growth factor 14 recapitulates phenotypic alterations underlying cognitive impairment associated with schizophrenia. Transl Psychiatry. 2016;6(5):e806. doi:10.1038/tp.2016.66

1221. Ali D, Laighneach A, Corley E, Patlola SR, Mahoney R, Holleran L, McKernan DP, Kelly JP, Corvin AP, Hallahan B, et al. Direct targets of MEF2C are enriched for genes associated with schizophrenia and cognitive function and are involved in neuron development and mitochondrial function. PLoS Genet. 2024;20(9):e1011093. doi:10.1371/journal.pgen.1011093

1222. Zai CC, Zai GC, Tiwari AK, Manchia M, de Luca V, Shaikh SA, Strauss J, Kennedy JL. Association study of GABRG2 polymorphisms with suicidal behaviour in schizophrenia patients with alcohol use disorder. Neuropsychobiology. 2014;69(3):154–158. doi:10.1159/000358839

1223. Chen X, Zhang Q, Su Y, Zhao W, Li Y, Du B, Deng X, Ji F, Dong Q, Chen C, et al. Evidence for the contribution of HCN1 gene polymorphism (rs1501357) to working memory at both behavioral and neural levels in schizophrenia patients and healthy controls. Schizophrenia (Heidelb). 2022;8(1):66. doi:10.1038/s41537-022-00271-7

1224. Bray NJ, Kirov G, Owen RJ, Jacobsen NJ, Georgieva L, Williams HJ, Norton N, Spurlock G, Jones S, Zammit S, et al. Screening the human protocadherin 8 (PCDH8) gene in schizophrenia. Genes Brain Behav. 2002;1(3):187–191. doi:10.1034/j.1601-183x.2002.10307.x

1225. Jiang J, Long J, Ling W, Huang G, Su L. Genetic variation in the 3’- untranslated region of PAK1 influences schizophrenia susceptibility. Exp Ther Med. 2017;13(3):1101–1108. doi:10.3892/etm.2017.4039

1226. Blennow K, Bogdanovic N, Heilig M, Grenfeldt B, Karlsson I, Davidsson P. Reduction of the synaptic protein rab3a in the thalamus and connecting brain regions in post-mortem schizophrenic brains. J Neural Transm (Vienna). 2000;107(8-9):1085–1097. doi:10.1007/s007020070054

1227. Liu Y, Fu X, Tang Z, Li C, Xu Y, Zhang F, Zhou D, Zhu C. Altered expression of the CSMD1 gene in the peripheral blood of schizophrenia patients. BMC Psychiatry. 2019;19(1):113. doi:10.1186/s12888-019-2089-4

1228. Suddaby JS, Silver J, So J. Understanding the schizophrenia phenotype in the first patient with the full SCN2A phenotypic spectrum. Psychiatr Genet. 2019;29(3):91–94. doi:10.1097/YPG.0000000000000219

1229. Roberts E. GABAergic malfunction in the limbic system resulting from an aboriginal genetic defect in voltage-gated Na+-channel SCN5A is proposed to give rise to susceptibility to schizophrenia. Adv Pharmacol. 2006;54:119–145. doi:10.1016/s1054-3589(06)54006-2

1230. Håvik B, Degenhardt FA, Johansson S, Fernandes CP, Hinney A, Scherag A, Lybæk H, Djurovic S, Christoforou A, Ersland KM, et al. DCLK1 variants are associated across schizophrenia and attention deficit/hyperactivity disorder. PLoS One. 2012;7(4):e35424. doi:10.1371/journal.pone.0035424

1231. Hattori E, Yamada K, Toyota T, Yoshitsugu K, Toru M, Shibuya H, Yoshikawa T. Association studies of the CT repeat polymorphism in the 5’ upstream region of the cholecystokinin B receptor gene with panic disorder and schizophrenia in Japanese subjects. Am J Med Genet. 2001;105(8):779–782. doi:10.1002/ajmg.10043

1232. Liu Y, Ding M, Liu YP, Zhang XC, Xing JX, Xuan JF, Xia X, Yao J, Wang BJ. Functional analysis of haplotypes and promoter activity at the 5’ region of the human GABRB3 gene and associations with schizophrenia. Mol Genet Genomic Med. 2019;7(5):e652. doi:10.1002/mgg3.652

1233. Kim MJ, Biag J, Fass DM, Lewis MC, Zhang Q, Fleishman M, Gangwar SP, Machius M, Fromer M, Purcell SM, et al. Functional analysis of rare variants found in schizophrenia implicates a critical role for GIT1- PAK3 signaling in neuroplasticity. Mol Psychiatry. 2017;22(3):417–429. doi:10.1038/mp.2016.98

1234. Demirel ÖF, Cetin İ, Turan Ş, Sağlam T, Yıldız N, Duran A. Decreased Expression of α-Synuclein, Nogo-A and UCH-L1 in Patients with Schizophrenia: A Preliminary Serum Study. Psychiatry Investig. 2017;14(3):344–349. doi:10.4306/pi.2017.14.3.344

1235. Hu TM, Chen SJ, Hsu SH, Cheng MC. Functional analyses and effect of DNA methylation on the EGR1 gene in patients with schizophrenia. Psychiatry Res. 2019;275:276–282. doi:10.1016/j.psychres.2019.03.044

1236. Boiko AS, Ivanova SA, Pozhidaev IV, Freidin MB, Osmanova DZ, Fedorenko OY, Semke AV, Bokhan NA, Wilffert B, Loonen AJM. Pharmacogenetics of tardive dyskinesia in schizophrenia: The role of CHRM1 and CHRM2 muscarinic receptors. World J Biol Psychiatry. 2020;21(1):72–77. doi:10.1080/15622975.2018.1548780

1237. Grubisha MJ, Sun X, MacDonald ML, Garver M, Sun Z, Paris KA, Patel DS, DeGiosio RA, Lewis DA, Yates NA, et al. MAP2 is differentially phosphorylated in schizophrenia, altering its function. Mol Psychiatry. 2021;26(9):5371–5388. doi:10.1038/s41380-021-01034-z

1238. Guo X, Zhang Y, Du J, Yang H, Ma Y, Li J, Yan M, Jin T, Liu X. Association analysis of ANK3 gene variants with schizophrenia in a northern Chinese Han population. Oncotarget. 2016;7(52):85888–85894. doi:10.18632/oncotarget.13043

1239. Guan F, Lin H, Chen G, Li L, Chen T, Liu X, Han J, Li T. Evaluation of association of common variants in HTR1A and HTR5A with schizophrenia and executive function. Sci Rep. 2016;6:38048. doi:10.1038/srep38048

1240. Li X, Zhang W, Zhang C, Gong W, Tang J, Yi Z, Wang D, Lu W, Fang Y, Chen X, et al. No association between genetic variants of the LRRK2 gene and schizophrenia in Han Chinese. Neurosci Lett. 2014;566:210–215. doi:10.1016/j.neulet.2014.03.006

1241. Udawela M, Scarr E, Boer S, Um JY, Hannan AJ, McOmish C, Felder CC, Thomas EA, Dean B. Isoform specific differences in phospholipase C beta 1 expression in the prefrontal cortex in schizophrenia and suicide. NPJ Schizophr. 2017;3:19. doi:10.1038/s41537-017-0020-x

1242. Ishizuka K, Yoshida T, Kawabata T, Imai A, Mori H, Kimura H, Inada T, Okahisa Y, Egawa J, Usami M, et al. Functional characterization of rare NRXN1 variants identified in autism spectrum disorders and schizophrenia. J Neurodev Disord. 2020;12(1):25.:10.1186/s11689-020-09325-2

1243. Morris JA, Kandpal G, Ma L, Austin CP. DISC1 (Disrupted-In-Schizophrenia 1) is a centrosome-associated protein that interacts with MAP1A, MIPT3, ATF4/5 and NUDEL: regulation and loss of interaction with mutation. Hum Mol Genet. 2003;12(13):1591–1608. doi:10.1093/hmg/ddg162

1244. Xu K, Zheng P, Zhao S, Feng J, Pu J, Wang J, Zhao S, Wang H, Chen J, Xie P. Altered MANF and RYR2 concentrations associated with hypolipidemia in the serum of patients with schizophrenia. J Psychiatr Res. 2023;163:142–149. doi:10.1016/j.jpsychires.2023.05.044

1245. Burhan AM, Anazodo UC, Marlatt NM, Palaniyappan L, Blair M, Finger E. Schizophrenia syndrome due to C9ORF72 mutation case report: a cautionary tale and role of hybrid brain imaging!. BMC Psychiatry. 2021;21(1):331. doi:10.1186/s12888-021-03341-9

1246. Chung Y, Dienel S, Belch M, Fish K, Ermentrout G, Lewis D, Chung D. Altered Rbfox1-Vamp1 pathway and prefrontal cortical dysfunction in schizophrenia. Preprint. Res Sq. 2023;rs.3.rs-2944372. doi:10.21203/rs.3.rs-2944372/v1

1247. Dienel SJ, Dowling KF, Barile Z, Bazmi HH, Liu A, Vespoli JC, Fish KN, Lewis DA. Diagnostic Specificity and Association With Cognition of Molecular Alterations in Prefrontal Somatostatin Neurons in Schizophrenia. JAMA Psychiatry. 2023;80(12):1235–1245. doi:10.1001/jamapsychiatry.2023.2972

1248. Reynolds GP. The neurochemical pathology of schizophrenia: post-mortem studies from dopamine to parvalbumin. J Neural Transm (Vienna). 2022;129(5-6):643–647. doi:10.1007/s00702-021-02453-6

1249. Li Y, Wang K, Zhang P, Huang J, An H, Wang N, De Yang F, Wang Z, Tan S, Chen S, et al. Quantitative DNA Methylation Analysis of DLGAP2 Gene using Pyrosequencing in Schizophrenia with Tardive Dyskinesia: A Linear Mixed Model Approach. Sci Rep. 2018;8(1):17466. doi:10.1038/s41598-018-35718-4

1250. Chaumette B, Ferrafiat V, Ambalavanan A, Goldenberg A, Dionne-Laporte A, Spiegelman D, Dion PA, Gerardin P, Laurent C, Cohen D, Rapoport J, et al. Missense variants in ATP1A3 and FXYD gene family are associated with childhood-onset schizophrenia. Mol Psychiatry. 2020;25(4):821–830. doi:10.1038/s41380-018-0103-8

1251. Shen Q, Zhang J, Wang Y, Liu B, Li X, Zhao Q, Chen S, Ji J, Yang F, Wan C, et al. No association between the KCNH1, KCNJ10 and KCNN3 genes and schizophrenia in the Han Chinese population. Neurosci Lett. 2011;487(1):61–65. doi:10.1016/j.neulet.2010.09.074

1252. Noori-Daloii MR, Kheirollahi M, Mahbod P, Mohammadi F, Astaneh AN, Zarindast MR, Azimi C, Mohammadi MR. Alpha- and beta-synucleins mRNA expression in lymphocytes of schizophrenia patients. Genet Test Mol Biomarkers. 2010;14(5):725–729. doi:10.1089/gtmb.2010.0050

1253. Kordi-Tamandani DM, Dahmardeh N, Torkamanzehi A. Evaluation of hypermethylation and expression pattern of GMR2, GMR5, GMR8, and GRIA3 in patients with schizophrenia. Gene. 2013;515(1):163–166. doi:10.1016/j.gene.2012.10.075

1254. Miyazawa A, Kanahara N, Kogure M, Otsuka I, Okazaki S, Watanabe Y, Yamasaki F, Nakata Y, Oda Y, Hishimoto A, et al. A preliminary genetic association study of GAD1 and GABAB receptor genes in patients with treatment-resistant schizophrenia. Mol Biol Rep. 2022;49(3):2015–2024. doi:10.1007/s11033-021-07019-z

1255. Grubor M, Zivkovic M, Sagud M, Nikolac Perkovic M, Mihaljevic-Peles A, Pivac N, Muck-Seler D, Svob Strac D. TR1A, HTR1B, HTR2A, HTR2C and HTR6 Gene Polymorphisms and Extrapyramidal Side Effects in Haloperidol-Treated Patients with Schizophrenia. Int J Mol Sci. 2020;21(7):2345. doi:10.3390/ijms21072345

1256. Chen Q, Che R, Wang X, O’Neill FA, Walsh D, Tang W, Shi Y, He L, Kendler KS, Chen X. Association and expression study of synapsin III and schizophrenia. Neurosci Lett. 2009;465(3):248–251. doi:10.1016/j.neulet.2009.09.032

1257. Gupta M, Jain S, Moily NS, Kaur H, Jajodia A, Purushottam M, Kukreti R. Genetic studies indicate a potential target 5-HTR(3B) for drug therapy in schizophrenia patients. Am J Med Genet B Neuropsychiatr Genet. 2012;159B(8):1006-1008. doi:10.1002/ajmg.b.32105

1258. Pozhidaev IV, Boiko AS, Loonen AJM, Paderina DZ, Fedorenko OY, Tenin G, Kornetova EG, Semke AV, Bokhan NA, Wilffert B, et al. Association of Cholinergic Muscarinic M4 Receptor Gene Polymorphism with Schizophrenia. Appl Clin Genet. 2020;13:97–105. doi:10.2147/TACG.S247174

1259. Davis KN, Tao R, Li C, Gao Y, Gondré-Lewis MC, Lipska BK, Shin JH, Xie B, Ye T, Weinberger DR, et al. Alternative Transcripts in the Human Prefrontal Cortex, and in Schizophrenia and Affective Disorders. PLoS One. 2016;11(2):e0148558. doi:10.1371/journal.pone.0148558

1260. Li JM, Lu CL, Cheng MC, Luu SU, Hsu SH, Chen CH. Genetic analysis of the DLGAP1 gene as a candidate gene for schizophrenia. Psychiatry Res. 2013;205(1-2):13–17. doi:10.1016/j.psychres.2012.08.014

1261. Braunewell KH, Dwary AD, Richter F, Trappe K, Zhao C, Giegling I, Schönrath K, Rujescu D. Association of VSNL1 with schizophrenia, frontal cortical function, and biological significance for its gene product as a modulator of cAMP levels and neuronal morphology. Transl Psychiatry. 2011;1(7):e22. doi:10.1038/tp.2011.20

1262. Marques TR, Natesan S, Niccolini F, Politis M, Gunn RN, Searle GE, Howes O, Rabiner EA, Kapur S. Phosphodiesterase 10A in Schizophrenia: A PET Study Using [(11)C]IMA107. Am J Psychiatry. 2016;173(7):714–721. doi:10.1176/appi.ajp.2015.15040518

1263. Habibzadeh P, Nemati A, Dastsooz H, Taghipour-Sheshdeh A, Mariam Paul P, Sahraian A, Ali Faghihi M. Investigating the association between common DRD2/ANKK1 genetic polymorphisms and schizophrenia: a meta-analysis. J Genet. 2021;100:59.

1264. Gardella R, Sacchetti E, Legati A, Magri C, Traversa M, Gennarelli M. Compound heterozygosity for a hemizygous rare missense variant (rs141999351) and a large CNV deletion affecting the FSTL5 gene in a patient with schizophrenia. Psychiatry Res. 2017;258:598–599. doi:10.1016/j.psychres.2016.10.057

1265. Harada S, Tachikawa H, Kawanishi Y. A possible association between an insertion/deletion polymorphism of the NQO2 gene and schizophrenia. Psychiatr Genet. 2003;13(4):205–209. doi:10.1097/00041444-200312000-00003

1266. Hartmann D, Fiedler J, Sonnenschein K, Just A, Pfanne A, Zimmer K, Remke J, Foinquinos A, Butzlaff M, Schimmel K, et al. MicroRNA-Based Therapy of GATA2-Deficient Vascular Disease. Circulation. 2016;134(24):1973–1990. doi:10.1161/CIRCULATIONAHA.116.022478

1267. de Jager SCA, Hoefer IE. Beyond the matrix: MMP2 as critical regulator of inflammation-mediated vascular dysfunction. Cardiovasc Res. 2017;113(14):1705–1707. doi:10.1093/cvr/cvx202

1268. Lawrie A, Spiekerkoetter E, Martinez EC, Ambartsumian N, Sheward WJ, MacLean MR, Harmar AJ, Schmidt AM, Lukanidin E, Rabinovitch M. Interdependent serotonin transporter and receptor pathways regulate S100A4/Mts1, a gene associated with pulmonary vascular disease. Circ Res. 2005;97(3):227–235. doi:10.1161/01.RES.0000176025.57706.1e

1269. Mazzuca MQ, Khalil RA. Vascular endothelin receptor type B: structure, function and dysregulation in vascular disease. Biochem Pharmacol. 2012;84(2):147–162. doi:10.1016/j.bcp.2012.03.020

1270. Van Bergen T, Etienne I, Cunningham F, Moons L, Schlingemann RO, Feyen JHM, Stitt AW. The role of placental growth factor (PlGF) and its receptor system in retinal vascular diseases. Prog Retin Eye Res. 2019;69:116–136. doi:10.1016/j.preteyeres.2018.10.006

1271. Saint-Martin Willer A, Montani D, Capuano V, Antigny F. Orai1/STIMs modulators in pulmonary vascular diseases. Cell Calcium. 2024;121:102892. doi:10.1016/j.ceca.2024.102892

1272. Kuriakose J, Montezano AC, Touyz RM. ACE2/Ang-(1-7)/Mas1 axis and the vascular system: vasoprotection to COVID-19-associated vascular disease. Clin Sci (Lond). 2021;135(2):387–407. doi:10.1042/CS20200480

1273. Wang Y, Xu J, Chen J, Fan X, Zhang Y, Yu W, Liu J, Hui R. Promoter variants of VTN are associated with vascular disease. Int J Cardiol. 2013;168(1):163–168. doi:10.1016/j.ijcard.2012.09.100

1274. Fu Y, Liu JW, Wu J, Wu ZX, Li J, Ji HF, Liang NP, Zhang HJ, Lai ZQ, Dong YF. Inhibition of semaphorin-3a alleviates lipopolysaccharide-induced vascular injury. Microvasc Res. 2022;142:104346. doi:10.1016/j.mvr.2022.104346

1275. Guglielmotto M, Monteleone D, Vasciaveo V, Repetto IE, Manassero G, Tabaton M, Tamagno E. The Decrease of Uch-L1 Activity Is a Common Mechanism Responsible for A? 42 Accumulation in Alzheimer’s and Vascular Disease. Front Aging Neurosci. 2017;9:320. doi:10.3389/fnagi.2017.00320

1276. Wang X, Athayde N, Trudinger B. Egr-1 transcription activation exists in placental endothelium when vascular disease is present. BJOG. 2006;113(6):683–687. doi:10.1111/j.1471-0528.2006.00928.x

1277. Zhang J, Chen J, Yang J, Xu C, Hu Q, Wu H, Cai W, Guo Q, Gao W, He C, et al. Suv39h1 downregulation inhibits neointimal hyperplasia after vascular injury. Atherosclerosis. 2019;288:76–84. doi:10.1016/j.atherosclerosis.2019.06.909

1278. Luo L, Cai Y, Zhang Y, Hsu CG, Korshunov VA, Long X, Knight PA, Berk BC, Yan C. Role of PDE10A in vascular smooth muscle cell hyperplasia and pathological vascular remodelling. Cardiovasc Res. 2022;118(12):2703–2717.

1279. Gauthier-Kemper A, Weissmann C, Golovyashkina N, Sebö-Lemke Z, Drewes G, Gerke V, Heinisch JJ, Brandt R. The frontotemporal dementia mutation R406W blocks tau’s interaction with the membrane in an annexin A2-dependent manner. J Cell Biol. 2011;192(4):647–661. doi:10.1083/jcb.201007161

1280. Tuna G, Yener GG, Oktay G, İşlekel GH, Kİrkalİ FG. Evaluation of Matrix Metalloproteinase-2 (MMP-2) and -9 (MMP-9) and Their Tissue Inhibitors (TIMP-1 and TIMP-2) in Plasma from Patients with Neurodegenerative Dementia. J Alzheimers Dis. 2018;66(3):1265–1273. doi:10.3233/JAD-180752

1281. Dhillon NK, Peng F, Bokhari S, Callen S, Shin SH, Zhu X, Kim KJ, Buch SJ. Cocaine-mediated alteration in tight junction protein expression and modulation of CCL2/CCR2 axis across the blood-brain barrier: implications for HIV-dementia. J Neuroimmune Pharmacol. 2008;3(1):52–56. doi:10.1007/s11481-007-9091-1

1282. Yaowaluk T, Senanarong V, Limwongse C, Boonprasert R, Kijsanayotin P. Influence of CYP2D6, CYP3A5, ABCB1, APOE polymorphisms and nongenetic factors on donepezil treatment in patients with Alzheimer’s disease and vascular dementia. Pharmgenomics Pers Med. 2019;12:209–224.

1283. Hansson O, Santillo AF, Meeter LH, Nilsson K, Landqvist Waldö M, Nilsson C, Blennow K, van Swieten JC, Janelidze S. SF placental growth factor - a novel candidate biomarker of frontotemporal dementia. Ann Clin Transl Neurol. 2019;6(5):863–872. doi:10.1002/acn3.763

1284. van Steenoven I, Noli B, Cocco C, Ferri GL, Oeckl P, Otto M, Koel-Simmelink MJA, Bridel C, van der Flier WM, Lemstra AW, et al. VGF Peptides in Cerebrospinal Fluid of Patients with Dementia with Lewy Bodies. Int J Mol Sci. 2019;20(19):4674. doi:10.3390/ijms20194674

1285. Ariaei A, Ghorbani A, Habibzadeh E, Moghaddam N, Chegeni Nezhad N, Abdoli A, Mazinanian S, Sadeghi M, Mayeli M. Investigating the association between the GAP-43 concentration with diffusion tensor imaging indices in Alzheimer’s dementia continuum. BMC Neurol. 2024;24(1):397. doi:10.1186/s12883-024-03904-9

1286. Bentivenga GM, Baiardi S, Mastrangelo A, Zenesini C, Mammana A, Polischi B, Capellari S, Parchi P. Diagnostic and prognostic value of cerebrospinal fluid SNAP-25 and neurogranin in Creutzfeldt-Jakob disease in a clinical setting cohort of rapidly progressive dementias. Alzheimers Res Ther. 2023;15(1):150. doi:10.1186/s13195-023-01300-y

1287. Crocco P, De Rango F, Bruno F, Malvaso A, Maletta R, Bruni AC, Passarino G, Rose G, Dato S. Genetic variability of FOXP2 and its targets CNTNAP2 and PRNP in frontotemporal dementia: A pilot study in a southern Italian population. Heliyon. 2024;10(11):e31624. doi:10.1016/j.heliyon.2024.e31624

1288. Shibata N, Motoi Y, Tomiyama H, Ohnuma T, Kuerban B, Tomson K, Komatsu M, Hattori N, Arai H. Lack of genetic association of the UCHL1 gene with Alzheimer’s disease and Parkinson’s disease with dementia. Dement Geriatr Cogn Disord. 2012;33(4):250–254. doi:10.1159/000339357

1289. Xie C, Miyasaka T. The Role of the Carboxyl-Terminal Sequence of Tau and MAP2 in the Pathogenesis of Dementia. Front Mol Neurosci. 2016;9:158. doi:10.3389/fnmol.2016.00158

1290. Yang D, Xie H, Wu S, Ying C, Chen Y, Ge Y, Yao R, Li K, Jiang Z, Chen G. Neurofilament light chain as a mediator between LRRK2 mutation and dementia in Parkinson’s disease. NPJ Parkinsons Dis. 2023;9(1):132. doi:10.1038/s41531-023-00572-3

1291. Strittmatter M, Cramer H, Reuner C, Strubel D, Hamann G, Schimrigk K. Molecular forms of somatostatin-like immunoreactivity in the cerebrospinal fluid of patients with senile dementia of the Alzheimer type. Biol Psychiatry. 1997;41(11):1124–1130. doi:10.1016/S0006-3223(96)00211-9

1292. Zagórska A, Bucki A, Partyka A, Jastrzębska-Więsek M, Siwek A, Głuch-Lutwin M, Mordyl B, Jaromin A, Walczak M, Wesołowska A, et al. Design, synthesis, and behavioral evaluation of dual-acting compounds as phosphodiesterase type 10A (PDE10A) inhibitors and serotonin ligands targeting neuropsychiatric symptoms in dementia. Eur J Med Chem. 2022;233:114218. doi:10.1016/j.ejmech.2022.114218

1293. Chiasserini D, Biscetti L, Eusebi P, Salvadori N, Frattini G, Simoni S, De Roeck N, Tambasco N, Stoops E, Vanderstichele H, et al. Differential role of CSF fatty acid binding protein 3, ?-synuclein, and Alzheimer’s disease core biomarkers in Lewy body disorders and Alzheimer’s dementia. Alzheimers Res Ther. 2017;9(1):52. doi:10.1186/s13195-017-0276-4

1294. Li R, Sano T, Mizokami A, Fukuda T, Shinjo T, Iwashita M, Yamashita A, Sanui T, Nakatsu Y, Sotomaru Y, et al. miR-582-5p targets Skp1 and regulates NF-?B signaling-mediated inflammation. Arch Biochem Biophys. 2023;734:109501. doi:10.1016/j.abb.2022.109501

1295. Pilar AV, Reid-Yu SA, Cooper CA, Mulder DT, Coombes BK. GogB is an anti-inflammatory effector that limits tissue damage during Salmonella infection through interaction with human FBXO22 and Skp1. PLoS Pathog. 2012;8(6):e1002773. doi:10.1371/journal.ppat.1002773

1296. Salas-Leal AC, Salas-Pacheco SM, Hernández-Cosaín EI, Vélez- Vélez LM, Antuna-Salcido EI, Castellanos-Juárez FX, Méndez-Hernández EM, Llave-León O, Quiñones-Canales G, Arias-Carrión O, et al. Differential expression of PSMC4, SKP1, and HSPA8 in Parkinson’s disease: insights from a Mexican mestizo population. Front Mol Neurosci. 2023;16:1298560. doi:10.3389/fnmol.2023.1298560

1297. Kumar P, Dezso Z, MacKenzie C, Oestreicher J, Agoulnik S, Byrne M, Bernier F, Yanagimachi M, Aoshima K, Oda Y. irculating miRNA biomarkers for Alzheimer’s disease. PLoS One. 2013;8(7):e69807. doi:10.1371/journal.pone.0069807

1298. Gámez-Valero A, Campdelacreu J, Vilas D, Ispierto L, Reñé R, Álvarez R, Armengol MP, Borràs FE, Beyer K. Exploratory study on microRNA profiles from plasma-derived extracellular vesicles in Alzheimer’s disease and dementia with Lewy bodies. Transl Neurodegener. 2019;8:31. doi:10.1186/s40035-019-0169-5

1299. Li Y, Shaw CA, Sheffer I, Sule N, Powell SZ, Dawson B, Zaidi SN, Bucasas KL, Lupski JR, Wilhelmsen KC, Doody R, et al. Integrated copy number and gene expression analysis detects a CREB1 association with Alzheimer’s disease. Transl Psychiatry. 2012;2(11):e192. doi:10.1038/tp.2012.119

1300. Shibata N, Ohnuma T, Higashi S, Higashi M, Usui C, Ohkubo T, Watanabe T, Kawashima R, Kitajima A, Ueki A, et al. Genetic association between USF 1 and USF 2 gene polymorphisms and Japanese Alzheimer’s disease. J Gerontol A Biol Sci Med Sci. 2006;61(7):660–662. doi:10.1093/gerona/61.7.660

1301. Li H, Wang F, Guo X, Jiang Y. Decreased MEF2A Expression Regulated by Its Enhancer Methylation Inhibits Autophagy and May Play an Important Role in the Progression of Alzheimer’s Disease. Front Neurosci. 2021;15:682247. doi:10.3389/fnins.2021.682247

1302. Yao Y, Kang SS, Xia Y, Wang ZH, Liu X, Muller T, Sun YE, Ye K. A delta-secretase-truncated APP fragment activates CEBPB, mediating Alzheimer’s disease pathologies. Brain. 2021;144(6):1833–1852. doi:10.1093/brain/awab062

1303. Lingam I, Avdic-Belltheus A, Meehan C, Martinello K, Ragab S, Peebles D, Barkhuizen M, Tann CJ, Tachtsidis I, Wolfs TGAM, et al. Serial blood cytokine and chemokine mRNA and microRNA over 48 h are insult specific in a piglet model of inflammation-sensitized hypoxia-ischaemia. Pediatr Res. 2021;89(3):464–475. doi:10.1038/s41390-020-0986-3

1304. Han K, Wang FR, Yu MQ. MicroRNA-21-5p promotes the inflammatory response after spinal cord injury by targeting PLAG1. Eur Rev Med Pharmacol Sci. 2020;24(11):5878–5885. doi:10.26355/eurrev_202006_21480

1305. Jia P, Zhang W, Shi Y. NFIC attenuates rheumatoid arthritis-induced inflammatory response in mice by regulating PTEN/SENP8 transcription. Tissue Cell. 2023;81:102013. doi:10.1016/j.tice.2023.102013

1306. Chen J, Bai Y, Xue K, Li Z, Zhu Z, Li Q, Yu C, Li B, Shen S, Qiao P, et al. CREB1-driven CXCR4hi neutrophils promote skin inflammation in mouse models and human patients. Nat Commun. 2023;14(1):5894. doi:10.1038/s41467-023-41484-3

1307. Xiong Y, Wang L, Jiang W, Pang L, Liu W, Li A, Zhong Y, Ou W, Liu B, Liu SM. MEF2A alters the proliferation, inflammation-related gene expression profiles and its silencing induces cellular senescence in human coronary endothelial cells. BMC Mol Biol. 2019;20(1):8. doi:10.1186/s12867-019-0125-z

1308. van der Krieken SE, Popeijus HE, Mensink RP, Plat J. CCAAT/enhancer binding proteinin relation to ER stress, inflammation, and metabolic disturbances. Biomed Res Int. 2015;2015:324815. doi:10.1155/2015/324815

1309. Cao Q, Guo Z, Du S, Ling H, Song C. Circular RNAs in the pathogenesis of atherosclerosis. Life Sci. 2020;255:117837. doi:10.1016/j.lfs.2020.117837

1310. Ye Y, Jin Q, Gong Q, Li A, Sun M, Jiang S, Jin Y, Zhang Z, He J, Zhuang L. Bioinformatics and Experimental Analyses Reveal NFIC as an Upstream Transcriptional Regulator for Ischemic Cardiomyopathy. Genes (Basel). 2022;13(6):1051. doi:10.3390/genes13061051

1311. Zhang T, Ge J. Mechanism of CREB1 in cardiac function of rats with heart failure via regulating the microRNA-376a-3p/TRAF6 axis. Mamm Genome. 2022;33(3):490–501. doi:10.1007/s00335-022-09947-y

1312. Pu Y, Zhao Q, Men X, Jin W, Yang M. MicroRNA-325 facilitates atherosclerosis progression by mediating the SREBF1/LXR axis via KDM1A. Life Sci. 2021;277:119464. doi:10.1016/j.lfs.2021.119464

1313. Muhammad Sulaiman K. A Study of transcription factor MEF2A gene polymorphisms in patients with coronary artery disease. Cell Mol Biol (Noisy-le-grand). 2024;70(1):80–86. doi:10.14715/cmb/2024.70.1.11

1314. Satoh T, Wang L, Espinosa-Diez C, Wang B, Hahn SA, Noda K, Rochon ER, Dent MR, Levine AR, Baust JJ, et al. Metabolic Syndrome Mediates ROS-miR-193b-NFYA-Dependent Downregulation of Soluble Guanylate Cyclase and Contributes to Exercise-Induced Pulmonary Hypertension in Heart Failure With Preserved Ejection Fraction. Circulation. 2021;144(8):615–637. doi:10.1161/CIRCULATIONAHA.121.053889

1315. Liu C, Gao Q, Dong J, Cai H. Usf2 Deficiency Promotes Autophagy to Alleviate Cerebral Ischemia-Reperfusion Injury Through Suppressing YTHDF1-m6A-Mediated Cdc25A Translation. Mol Neurobiol. 2024;61(5):2556–2568. doi:10.1007/s12035-023-03735-8

1316. Cortes-Canteli M, Luna-Medina R, Sanz-Sancristobal M, Alvarez-Barrientos A, Santos A, Perez-Castillo A. CCAAT/enhancer binding protein beta deficiency provides cerebral protection following excitotoxic injury. J Cell Sci. 2008;121(Pt 8):1224–1234. doi:10.1242/jcs.025031

1317. Li J, Lv H, Che Y. microRNA-381-3p Confers Protection Against Ischemic Stroke Through Promoting Angiogenesis and Inhibiting Inflammation by Suppressing Cebpb and Map3k8. Cell Mol Neurobiol. 2020;40(8):1307–1319. doi:10.1007/s10571-020-00815-4

1318. Kim HK, Tyryshkin K, Elmi N, Dharsee M, Evans KR, Good J, Javadi M, McCormack S, Vaccarino AL, Zhang X, et al. Plasma microRNA expression levels and their targeted pathways in patients with major depressive disorder who are responsive to duloxetine treatment. J Psychiatr Res. 2019;110:38–44. doi:10.1016/j.jpsychires.2018.12.007

1319. Wang P, Yang Y, Yang X, Qiu X, Qiao Z, Wang L, Zhu X, Sui H, Ma J. CREB1 gene polymorphisms combined with environmental risk factors increase susceptibility to major depressive disorder (MDD). Int J Clin Exp Pathol. 2015;8(1):906–913.

1320. Resende LS, Amaral CE, Soares RB, Alves AS, Alves-Dos-Santos L, Britto LR, Chiavegatto S. Social stress in adolescents induces depression and brain-region-specific modulation of the transcription factor MAX. Transl Psychiatry. 2016;6(10):e914. doi:10.1038/tp.2016.202

1321. Cao D, Wang J, Ji Z, Shangguan Y, Guo W, Feng X, Xu K, Yang J. Profiling the mRNA and miRNA in Peripheral Blood Mononuclear Cells in Subjects with Active Tuberculosis. Infect Drug Resist. 2020;13:4223–4234. doi:10.2147/IDR.S278705

1322. de Klerk JA, Beulens JWJ, Bijkerk R, van Zonneveld AJ, Elders PJM, ’t Hart LM, Slieker R. Circulating small non-coding RNAs are associated with the insulin-resistant and obesity-related type 2 diabetes clusters. Diabetes Obes Metab. 2024;26(10):4375–4385. doi:10.1111/dom.15786

1323. Kuryłowicz A, Wicik Z, Owczarz M, Jonas MI, Kotlarek M, Świerniak M, Lisik W, Jonas M, Noszczyk B, Puzianowska-Kuźnicka M. NGS Reveals Molecular Pathways Affected by Obesity and Weight Loss-Related Changes in miRNA Levels in Adipose Tissue. Int J Mol Sci. 2017;19(1):66. doi:10.3390/ijms19010066

1324. Czogała W, Czogała M, Strojny W, Wątor G, Wołkow P, Wójcik M, Bik Multanowski M, Tomasik P, Wędrychowicz A, Kowalczyk W, et al. Methylation and Expression of FTO and PLAG1 Genes in Childhood Obesity: Insight into Anthropometric Parameters and Glucose-Lipid Metabolism. Nutrients. 2021;13(5):1683. doi:10.3390/nu13051683

1325. Sukhatme MG, Kar A, Arasu UT, Lee SHT, Alvarez M, Garske KM, Gelev KZ, Rajkumar S, Das SS, Kaminska D, et al. ntegration of single cell omics with biobank data discovers trans effects of SREBF1 abdominal obesity risk variants on adipocyte expression of more than 100 genes. Preprint. medRxiv. 2024;2024.11.22.24317804. doi:10.1101/2024.11.22.24317804

1326. Bennett CE, Nsengimana J, Bostock JA, Cymbalista C, Futers TS, Knight BL, McCormack LJ, Prasad UK, Riches K, Rolton D, et al. CCAAT/enhancer binding protein alpha, beta and delta gene variants: associations with obesity related phenotypes in the Leeds Family Study. Diab Vasc Dis Res. 2010;7(3):195–203. doi:10.1177/1479164110366274

1327. Groen K, Maltby VE, Scott RJ, Tajouri L, Lechner-Scott J. Erythrocyte microRNAs show biomarker potential and implicate multiple sclerosis susceptibility genes. Clin Transl Med. 2020;10(1):74–90. doi:10.1002/ctm2.22

1328. Salemi M, Marchese G, Lanza G, Cosentino FII, Salluzzo MG, Schillaci FA, Ventola GM, Cordella A, Ravo M, Ferri R. Role and Dysregulation of miRNA in Patients with Parkinson’s Disease. Int J Mol Sci. 2022;24(1):712. doi:10.3390/ijms24010712

1329. Liu T, Li G. miR-15b-5p transcription mediated by CREB1 protects against inflammation and apoptosis in Parkinson disease models by inhibiting AXIN2 and activating Wnt/?-catenin. J Neuropathol Exp Neurol. 2023;82(12):995–1009. doi:10.1093/jnen/nlad084

1330. Yuan X, Cao B, Wu Y, Chen Y, Wei Q, Ou R, Yang J, Chen X, Zhao B, Song W, et al. Association analysis of SNP rs11868035 in SREBF1 with sporadic Parkinson’s disease, sporadic amyotrophic lateral sclerosis and multiple system atrophy in a Chinese population. Neurosci Lett. 2018;664:128–132. doi:10.1016/j.neulet.2017.11.015

1331. Casado Gama H, Amorós MA, Andrade de Araújo M, Sha CM, Vieira MPS, Torres RGD, Souza GF, Junkes JA, Dokholyan NV, Leite Góes Gitaí D, et al. Systematic review and meta-analysis of dysregulated microRNAs derived from liquid biopsies as biomarkers for amyotrophic lateral sclerosis. Noncoding RNA Res. 2024;9(2):523–535. doi:10.1016/j.ncrna.2024.02.006

1332. Joilin G, Gray E, Thompson AG, Bobeva Y, Talbot K, Weishaupt J, Ludolph A, Malaspina A, Leigh PN, Newbury SF, et al. Identification of a potential non-coding RNA biomarker signature for amyotrophic lateral sclerosis. Brain Commun. 2020;2(1):fcaa053. doi:10.1093/braincomms/fcaa053

1333. Zhao G, Fu Y, Yang C, Yang X, Hu X. Exploring the pathogenesis linking traumatic brain injury and epilepsy via bioinformatic analyses. Front Aging Neurosci. 2022;14:1047908. doi:10.3389/fnagi.2022.1047908

1334. Wang X, Sun Y, Tan Z, Che N, Ji A, Luo X, Sun X, Li X, Yang K, Wang G, et al. Serum MicroRNA-4521 is a Potential Biomarker for Focal Cortical Dysplasia with Refractory Epilepsy. Neurochem Res. 2016;41(4):905–912. doi:10.1007/s11064-015-1773-0

1335. Huang Y, Wu X, Guo J, Yuan J. Myocyte-specific enhancer binding factor 2A expression is downregulated during temporal lobe epilepsy. Int J Neurosci. 2016;126(9):786–796. doi:10.3109/00207454.2015.1062003

1336. Mirra P, Desiderio A, Spinelli R, Nigro C, Longo M, Parrillo L, D’Esposito V, Carissimo A, Hedjazifar S, Smith U, et al. Adipocyte precursor cells from first degree relatives of type 2 diabetic patients feature changes in hsa-mir-23a-5p, -193a-5p, and -193b-5p and insulin-like growth factor 2 expression. FASEB J. 2021;35(4):e21357. doi:10.1096/fj.202002156RRR

1337. Shahin NN, Shaker OG, Mahmoud MO. GOAT rs10096097 and CREB1 rs6740584 single nucleotide polymorphisms are associated with type 2 diabetes mellitus in Egyptians. Arch Pharm (Weinheim). 2024;357(8):e2400011. doi:10.1002/ardp.202400011

1338. Grarup N, Stender-Petersen KL, Andersson EA, Jørgensen T, Borch-Johnsen K, Sandbaek A, Lauritzen T, Schmitz O, Hansen T, Pedersen O. Association of variants in the sterol regulatory element-binding factor 1 (SREBF1) gene with type 2 diabetes, glycemia, and insulin resistance: a study of 15,734 Danish subjects. Diabetes. 2008;57(4):1136–1142. doi:10.2337/db07-1534

1339. Lobanov SV, McAllister B, McDade-Kumar M, Landwehrmeyer GB, Orth M, Rosser AE; REGISTRY Investigators of the European Huntington’s disease network; Paulsen JS; PREDICT-HD Investigators of the Huntington Study Group; Lee JM, et al. Huntington’s disease age at motor onset is modified by the tandem hexamer repeat in TCERG1. NPJ Genom Med. 2022;7(1):53. doi:10.1038/s41525-022-00317-w

1340. McCourt AC, Parker J, Silajdžić E, Haider S, Sethi H, Tabrizi SJ, Warner TT, Björkqvist M. Analysis of White Adipose Tissue Gene Expression Reveals CREB1 Pathway Altered in Huntington’s Disease. J Huntingtons Dis. 2015;4(4):371–382. doi:10.3233/JHD-150172

1341. Reynolds CA, Hong MG, Eriksson UK, Blennow K, Wiklund F, Johansson B, Malmberg B, Berg S, Alexeyenko A, Grönberg H, et al. nalysis of lipid pathway genes indicates association of sequence variation near SREBF1/TOM1L2/ATPAF2 with dementia risk. Hum Mol Genet. 2010;19(10):2068–2078. doi:10.1093/hmg/ddq079

1342. Wang X, Zhang G, Lu W, Zhang Y, Fan W, Tang W, Zhang C. Common variants in CREB1 gene confer risk for bipolar disorder in Han Chinese. Asian J Psychiatr. 2021;59:102648. doi:10.1016/j.ajp.2021.102648

1343. Pan B, Han B, Zhu X, Wang Y, Ji H, Weng J, Liu Y. Dysfunctional microRNA-144-3p/ZBTB20/ERK/CREB1 signalling pathway is associated with MK-801-induced schizophrenia-like abnormalities. Brain Res. 2023;1798:148153. doi:10.1016/j.brainres.2022.148153

1344. Yang L, Chen J, Li Y, Wang Y, Liang S, Shi Y, Shi S, Xu Y. Association between SCAP and SREBF1 gene polymorphisms and metabolic syndrome in schizophrenia patients treated with atypical antipsychotics. World J Biol Psychiatry. 2016;17(6):467–474. doi:10.3109/15622975.2016.1165865

